# ELAVL1 and ELAVL4 are required for Musashi-dependent translational activation

**DOI:** 10.64898/2026.07.01.735911

**Authors:** Katherine Bronson, Milla M. Reddick, Kenzie B. MacNicol, Cole Bolen, Linda Hardy, Alex Lagasse, Angela K. Odle, Gwen V. Childs, Melanie C. MacNicol, Angus M. MacNicol

## Abstract

The RNA-binding proteins Musashi1 and Musashi2 (MSI1 and MSI2) regulate stem cell function and tissue plasticity by modulating mRNA translation. While typically known as translational repressors, the MSI1 and MSI2 proteins can also act as context-dependent activators of mRNA translation, although the mechanism of MSI-mediated translational activation are unknown. Here, we identify Embryonic Lethal Abnormal Vision-like (ELAVL) proteins as essential co-regulators of MSI1-dependent translational activation. In *Xenopus laevis* oocytes, antisense oligonucleotide knockdown of *Elavl4* inhibited progesterone-stimulated maturation and blocked polyadenylation and translation of key MSI target mRNAs, including the *Mos* and *Cyclin B5* mRNAs. Exogenous expression of ELAVL4 rescued these defects, confirming its necessity for maturation and cell cycle progression. Mechanistically, we determined that the ELAVL4 C-terminal domain interacts with the N-terminal RNA recognition motifs of MSI1 in an RNA-independent manner. Mass spectrometry and functional assays revealed this interaction is evolutionarily conserved: mouse ELAVL1 interacts with MSI1 in the pituitary, and human ELAVL1 rescues *Elavl4*-depleted *Xenopus* oocytes. Furthermore, knockdown of *Elavl1* in a mammalian cell line abrogated MSI-dependent translational activation of a pituitary *Prop1* 3’ UTR mRNA reporter. Our results establish a conserved mechanism where ELAVL family members interact with MSI to promote MSI-dependent mRNA translational activation.

## Introduction

The Musashi (MSI) family of evolutionarily conserved, sequence-specific mRNA binding proteins (MSI1 and MSI2) has been shown to regulate the translation of target transcripts involved in the promotion of stem cell self-renewal and the suppression of cell differentiation.^1^ Dysregulated MSI activity is associated with a number of pathologies including neurodegeneration, cancer, and Zika virus-mediated microcephaly.^1–5^ Both MSI paralogs – MSI1 and MSI2 – are conserved within vertebrates and despite apparent functional redundancy, show distinct tissue-specific patterns of expression.^4,6–8^ While originally considered a repressor of target mRNA translation^9^, our group identified an alternate requirement for MSI in the translational activation of target mRNAs in the progesterone-stimulated maturation of *Xenopus* oocytes^10,11^ and also in mammalian cell reporter assays^12^, establishing that MSI acts in a context-dependent manner to either repress or to activate target mRNA translation.^12–16^ Recently, a novel truncated form of MSI1 has been reported (C-MSI1), that drives embryonic stem cell pluripotency through novel mechanism(s) distinct from the canonical MSI1 protein.^17^

The vertebrate pituitary functions as a master endocrine gland to coordinate systemic hormonal activity and has the highest tissue expression levels of both *Msi1* and *Msi2* isoforms other than the gonads.^15^ Our previous work has implicated MSI function in pituitary cell fate plasticity, in which the pituitary remodels its hormone-producing cell populations in response to changing organismal needs. We have shown that MSI targets a range of key pituitary mRNAs, repressing translation of some targets, e.g. the *Pou1f1/Pit1*, *Gnrhr*, *Fsh*, *Prl*, and *Tshb* mRNAs^14–16,18^, while activating translation of others, e.g. the *Prop1*, *Gata2*, and *Nr5a1* (*SF-1*) mRNAs.^12^ Moreover, we have shown that *Msi1* and *Msi2* are necessary *in vivo* for cycling female pituitary gonadotropes to produce normal levels of luteinizing hormone (LH) secretion.^14^

The mechanisms by which MSI function distinguishes between mRNA target repression versus mRNA target translational activation, are unknown. Both categories of MSI target mRNAs are recognized via the highly conserved RNA recognition motifs (RRMs) in the N-terminal of MSI, that recognize canonical MSI-binding elements (MBEs, (A/G)U_1-3_AG)) located within the target mRNA 3’ untranslated region (3’ UTR).^1,19^ However, RNA binding alone is not sufficient to activate translation. Our earlier work revealed that MSI complexes undergo dynamic remodeling in *Xenopus laevis* oocytes after progesterone stimulation which may serve as a functional switch between target repression and target activation.^20^ Indeed, GLD2, ePABP/PABPC1 and LSM14B protein associations with MSI1 are required for translation of the *Mos* and *Cyclin B5* mRNAs as well as to promote *Xenopus* oocyte progesterone-stimulated maturation^13,20,21^ and noteably, LSM14B is also required for MSI1-dependent translational activation of the *Prop1* mRNA in mammalian cells.^13^ An additional 45 MSI1-associated proteins have been identified in mature *Xenopus* oocytes, including three members of the Embryonic Lethal Abnormal Vision-like (ELAVL) protein family, ELAVL1 (HuR), ELAVL2 (HuB) and ELAVL4 (HuD) which are also termed ElrA, ElrB and ElrD respectively in *Xenopus*.^20^

The ELAVL protein family are RNA-binding proteins with established roles as regulators of mRNA translation.^22,23^ ELAVL1 is widely expressed, while ELAVL2, ELAVL3, and ELAVL4 are associated with neurodevelopment and are highly expressed in neural tissue.^23–25^ ELAVL family members bind to AU-rich element (ARE)-containing mRNA targets and generally promote cytoplasmic localization, transcript stabilization, and mRNA translation.^26^ ELAVL4 has been detected *via* auto-antibodies in patients with paraneoplastic encephalomyelitis and sensory neuropathy and has since been implicated in axonal growth, cranial nerve development, the establishment of hippocampal circuits, and the differentiation of neural stem cells in the subventricular zone.^27–30^ Similar to MSI, ELAVL4 contains N-terminal RRMs; the first two of which bind to target ARE regions, while the third RRM binds to target mRNA poly[A] tails and to other proteins.^31–33^ It has been reported that ELAVL4 and MSI1 associate with common mRNA targets, including *p21^WAF^*^1^ and *Myc*, although it has not been elucidated whether they directly cooperate or antagonize each other when occupying the same mRNA 3’ UTR.^34–38^ Here, we have identified a role for ELAVL proteins in promoting MSI mRNA translational activation function.

In the current study, we report that knockdown of ELAVL4 attenuates progesterone-stimulated *Xenopus* oocyte maturation and MSI-mediated translation of target mRNAs. Notably, ectopic expression of mRNA encoding either ELAVL4 or of ELAVL1 could rescue progesterone-dependent maturation in ELAVL4-knockdown oocytes, demonstrating redundancy of ELAVL family members to promote oocyte maturation and MSI-dependent mRNA translation. Interestingly, while ectopic MSI1 could rescue maturation in ELAVL4-knockdown oocytes, ectopic ELAVL4 could not rescue maturation in MSI-knockdown *Xenopus* oocytes. Both *Xenopus* ELAVL4 and mammalian ELAVL1 interact with MSI1 in an RNA-independent manner. Further, ELAVL1 interacts with MSI in mouse pituitaries and is required for MSI1-dependent translational activation directed by the pituitary *Prop1* mRNA 3’ UTR in a luciferase reporter assay. Our results reveal a conserved requirement for the interaction of MSI1 with ELAVL proteins for MSI-dependent mRNA translational activation.

## Methods

### Oocyte Culture and Microinjections

Dumont Stage VI immature Xenopus laevis oocytes were isolated and cultured as previously described.^39^ Oocytes were microinjected using a Nanoject II Auto-Nanoliter Injector (Drummond Scientific). mRNA for oocyte injection was made by linearization of the expression plasmids and *in vitro* transcription using SP6 (Promega) RNA polymerase. Oocytes were stimulated to mature with 2 µg/ml progesterone. The appearance of a white spot on the animal pole was used to score the rate of oocyte maturation as it indicates germinal vesicle (nuclear) breakdown (GVBD). Where indicated, progesterone-stimulated oocytes were segregated when 50% of the oocytes completed GVBD (GVBD50) into those that had not (-) or had (+) completed GVBD. Animal protocols were approved by the UAMS Institutional Animal Care and Use Committee, in accordance with federal regulations.

### Oocyte Lysis and Sample Preparation

For protein:protein co-association experiments, oocytes were lysed in 10 µL/oocyte of ice cold NP40 lysis buffer (1% NP40, 20mM Tris pH 8.0, 137mM NaCl, 10% glycerol, 2mM EDTA, 50mM NaF, 10mM Sodium Pyrophosphate, 1mM PMSF, 1x Halt Protease Inhibitor Cocktail (ThermoFisher Scientific)). For western blot analysis and pulldowns from Xenopus oocytes, yolk and cell debris were removed by centrifugation at 14,500+ RCF for 10 minutes in a refrigerated tabletop centrifuge at 4°C. For each sample, half oocyte equivalents of lysate were prepared in NuPAGE sample loading buffer and electrophoresed through a 10% NuPAGE gel (ThermoFisher).

For RNA extraction and polyadenylation assays, media-free oocytes were treated with 800 µL STAT-60 (Tel-Test, Inc) for RNA extraction using the manufacture’s protocol followed by a subsequent purification by precipitation in 4M LiCl at -20°C overnight and centrifugation at 14,500 RCF for 10 minutes in a refrigerated tabletop centrifuge.

### Pulldown and RNase Treatment

Oocytes were injected with 4.4 µg/µL each in vitro transcribed mRNA and incubated for 16h at 18°C. Lysates were prepared as described above. 300 µL of oocyte lysate was added to 440 µL ice cold NP40 lysis buffer and incubated with 60 µL of 50% glutathione sepharose conjugated bead slurry (GE) at 4°C for 5h with gentle rotation. Beads were then gently pelleted by centrifugation at 20 RCF for 5 minutes; the supernatant was removed and replaced with 500 µL fresh NP40 lysis buffer and the beads inverted to mix, and the buffer then removed. This process was repeated 3 times at 4°C. On the third wash, 200U of RNase1 (Ambion) was added and incubated for 15 minutes at 37°C. Following final centrifugation, all NP40 was removed and 50 µL of NuPAGE sample loading buffer (ThermoFisher) was added. The samples were incubated at 4° with gentle rotation overnight. The next day, the beads were incubated for 10 minutes at 70°C, then crushed by centrifugation at 14,500 RCF for 10 minutes. Finally, 30 µL of the sample was loaded per each lane of a 10% NuPAGE gel (ThermoFisher) and electrophoresed.

### Western Blotting

NuPAGE gels were transferred to a 0.2 µm-pore-size nitrocellulose membrane (Protran; Midwest Scientific) after electrophoresis. The membrane was blocked with 5% non-fat dried milk in TBST (20mM Tris pH 7.5, 150mM NaCl, 0.05% Tween20) for 60 min at room temperature, or overnight at 4°C. Following incubation with primary antibody and 10% block by volume, filters were washed 6x5 minutes in TBST, incubated with horseradish peroxidase-conjugated secondary antibody for two hours then washed 6x5 minutes in TBST. Blots were developed using enhanced chemiluminescence in a FluorChem M Imager system (ProteinSimple Bio-techne).

### Antibodies

The antibodies used for immuno-detection were Anti-GAPDH (1:5000, 14C10, Cell Signaling), Anti-GST (1:1000, ab19256, Abcam; 1:1000, A5800, Invitrogen), Anti-GFP (1:5000, A11122, Invitrogen), Anti-Tubulin (1:5000, ab7291, AbCam), Anti-Mos^xe^ (C237) (1:100, sc-86, Santa Cruz), HRP-Conjugated Rabbit (1:5000 or 1:10,000 as indicated, W401B, Promega), and HRP-Conjugated Mouse (1:10,000, W402B, Promega). All working antibody preparations were made in TBST + 0.5% non-fat milk or 0.2% BSA.

### Polyadenylation Assays

cDNAs for polyadenylation assays were synthesized using RNA ligation-coupled PCR as described.^40^ The increase in PCR product length is specifically due to extension of the poly[A] tail.^10,41,42^ The same reverse primer P1’ was used for all reactions and has the sequence: 5’-GCTTCAGATCAAGGTGACCTTTTT-3’. The Mos forward primer has the sequence: 5’-GCAAGGATATGAAAAAAAGATTTC-3’. The Cyclin B1 forward primer has the sequence: 5 - GTGGCATTCCAATTGTGTATTGTT-3. The Cyclin B5 forward primer has the sequence 5’-AATAACTTTTTAAGTAGTCGCTGG-3’.

### Antisense Oligodeoxynucleotide Injections

Antisense oligodeoxynucleotides targeting the Xenopus MSI1 (5’-GCGCTTCTGTCTCCATTCGGTCTCT-3’) and MSI2 (5’- CCCATCTGCCTCCATAGCCTTCTC-3’) mRNAs have been described previously.^10^ Antisense oligodeoxynucleotides targeting *Xenopus Elavl1, Elavl2* and *Elavl4* were designed to span the ATG start codon (bold): for *Elavl1* mRNA (5’-CCGTTAGA**CAT**CTTATATTACC-3’); for *Elavl2* mRNA (5’-CACACAGTCTGACTGC**CAT**GTTAG-3’); and for *Elavl4* mRNA (5’-GCCATTCCACTC**CAT**CTGTGACTATGG-3’). Control oocytes were injected with a scrambled oligonucleotide with the sequence 5’-TAGAGAAGATAATCGTCATCTTA-3’.^10^ A total of 100 ng of antisense oligonucleotides was injected for each condition and oocytes were incubated at 18°C for 16 hours. Following overnight incubation, oocytes were stimulated with progesterone and the extent of GVBD scored.

### Plasmids and Plasmid Construction

The following plasmids have been described previously: pXen^43^, pXen-GFP,^21^ pXen-GFP-xMsi1^11^, pXen-GFP-xMSI2^21^, pmiRGLO mProp1 last 195 nucleotide 3’ UTR^12^, peGFPN1 and peGFPN1 MSI1.^44^ Antisense resistant (“wobble”) pXen-XeELAVL4 was constructed using a GeneBlock fragment purchased from IDT containing XhoI and BamHI restriction sites for the 5’ and 3’ ends respectively, the *Xenopus Elavl4* coding sequence (NM_001087440) containing an altered 5’ starting sequence (ATGGAATGGAACGGA instead of wild-type ATGGAGTGGAATGGC to retain the same amino acid sequence but disrupt targeting by the *Elavl4* antisense oligonucleotide). After using restriction enzymes to digest the XeElavl4 GeneBlock and the pXen2 vector^43^, they were ligated using NEB Quick Ligase (M2200) following the manufacturer’s protocol. This generated an N-terminal GST tagged *Elavl4* expression plasmid, designated pXen-GST-XeELAVL4. PCR primers were designed to insert either a XhoI site or a 5’ STOP codon followed by a 3’ BamHI site between nucleotides 753 and 754 of the 1203 nucleotide XeELAVL4 in pXen-XeELAVL4. pXen-XeELAVL4 with the added STOP codon and BamHI site was digested with BamHI to generated a truncated N-terminal XeELAVL4 variant in pXen, while pXen-XeELAVL4 with the added XhoI site was digested with XhoI to generate a truncated C-terminal XeELAVL4 variant in pXen. The resulting plasmids were sequence verified and designated pXen-GST N-XeELAVL4 (which spans AA 1-251 or nucleotides 1-753 of full-length XeELAVL4) and pXen-GST-C-XeELAVL4 (which spans AA 252-400 or nucleotides 754-1203 of full-length XeELAVL4). The XeELAVL4 fragment was isolated from pXen-GST-XeELAVL4 via digestion with XhoI and BamHI and inserted into pXen-GFP^21^ digested with XhoI and BamHI, to generate an N-terminal GFP tagged *Elavl4* designated peGFP-XeELAVL4. A Geneblock oligonucleotide (IDT) was designed to encode the human ELAVL1 protein (NM_001419) flanked by 5’ Cla1 and 3’ Xho1 restriction sites and ligated into ClaI/XhoI digested pXen2^43^ and the resulting plasmid was sequence verified and designated pXen-GST-huELAVL1.

### Animals

The Institutional Animal Care and Use Committee approved all animal care protocols. All mice belonged to the sighted FVB.129P hybrid strain, from Jackson Laboratories (FVB.129P2-Pde6b+ Tyrc-ch/Ant) and are referred to as FVB mice. *Xenopus laevis* adult (> 1 year old) females were purchased as needed from Xenopus 1 (Michigan, USA) and housed on a 12 h light/12 h dark cycle in dechlorinated freshwater tanks maintained at 20 C°.

### Immunoprecipitation and Mass Spectrometry

The Dynabeads Protein A Immunoprecipitation Kit (#10006D) was used for the immunoprecipitations, and the manufacturer’s protocols were followed. Six whole male pituitaries were collected, pooled, and flash frozen in liquid nitrogen until use. After adding 250 µL RIPA to the pituitaries, they were homogenized using a syringe and then centrifuged at 4°C for 10 minutes at 19,700 RCF. The supernatant was moved to a new Eppendorf tube and sat on ice for up to 1 hour and centrifuged for 10 minutes at 4°C at 14,500 RCF. 100 µL of the lysate was added to the Dynabeads bound by MSI1/MSI2 or IgG antibodies per manufacturer’s protocol. On the final wash of the protocol, 1 µL of RQ DNase and 1 µL RNase 1 was added and the immunoprecipitation samples were incubated for 30 minutes then washed. After the last wash, the supernatant was removed, and the samples were washed twice with 1 mL fresh cold 100 mM ammonium hydrogen carbonate (AMBIC) for a mass spectrometry-compatible buffer. On the second wash the beads were transferred to a new Eppendorf tube, and the samples were spun down to remove the supernatant. This was repeated on three separate occasions (n=3, 18 whole male pituitaries total). Supernatant samples were submitted to the IdEA Proteomics Core at the University of Arkansas for Medical Sciences.

Protein samples were reduced, alkylated, and digested on-bead using filter-aided sample preparation^45^ with sequencing grade modified porcine trypsin (Promega). Tryptic peptides were then separated by reverse phase XSelect CSH C18 2.5 um resin (Waters) on an in-line 150 x 0.075 mm column using an UltiMate 3000 RSLCnano system (Thermo). Peptides were eluted using a 60 min gradient from 98:2 to 65:35 buffer A:B ratio (Buffer A = 0.1% formic acid, 0.5% acetonitrile; Buffer B = 0.1% formic acid, 99.9% acetonitrile). Eluted peptides were ionized by electrospray (2.2 kV) followed by mass spectrometric analysis on an Orbitrap Exploris 480 mass spectrometer (Thermo). To assemble a chromatogram library, six gas-phase fractions were acquired on the Orbitrap Exploris with 4 m/z DIA spectra (4 m/z precursor isolation windows at 30,000 resolution, normalized AGC target 100%, maximum inject time 66 ms) using a staggered window pattern from narrow mass ranges using optimized window placements. Precursor spectra were acquired after each DIA duty cycle, spanning the m/z range of the gas-phase fraction (i.e., 496-602 m/z, 60,000 resolution, normalized AGC target 100%, maximum injection time 50 ms). For wide-window acquisitions, the Orbitrap Exploris was configured to acquire a precursor scan (385-1015 m/z, 60,000 resolution, normalized AGC target 100%, maximum injection time 50 ms) followed by 50x 12 m/z DIA spectra (12 m/z precursor isolation windows at 15,000 resolution, normalized AGC target 100%, maximum injection time 33 ms) using a staggered window pattern with optimized window placements. Precursor spectra were acquired after each DIA duty cycle.

Following data acquisition, data were searched using an empirically corrected library against the UniProt *Mus musculus* database and a quantitative analysis was performed to obtain a comprehensive proteomic profile. Proteins were identified and quantified using EncyclopeDIA^46^ and visualized with Scaffold DIA using 1% false discovery thresholds at both the protein and peptide level. Protein exclusive intensity values were assessed for quality using ProteiNorm.^47^ The data was normalized using cyclic loess^48^ and statistical analysis was performed using linear models for microarray data (limma) with empirical Bayes (eBayes) smoothing to the standard errors.^48^ Proteins with an FDR adjusted p-value < 0.055 and a positive fold change ≥ 2 were considered significant. The datasets from these MS analyses have been deposited in MassIVE (PXD080073).

### Luciferase reporter assays

For siRNA knockdown of ELAVL1 experiments, NIH3T3 cells (ATCC CRL-1658) were seeded in a 12 well plate at 1 x 10^5^ cells/well and transfected with either a scrambled control siRNA or siRNA targeting murine ELAVL1 24 h post-seeding using the TriFECTa DsiRNA kit (IDT, mm.RI.Elavl1.13). Two of the three siRNAs targeting ELAVL1, mm.RI.Elavl1.13.1 and mm.RI.Elavl1.13.3, were found to efficiently knock down ELAVL1 protein levels. After 48 hr incubation with siRNA, cells were then co-transfected with the pmiRGLO mProp1 last 195 mucleotide 3′ UTR plasmid along with either murine MSI1-eGFP or eGFP control plasmids as described previously.^44,49^ Cells were incubated for 18–24 h after transfection, and luciferase activity was determined in quadruplicate using the Dual-Luciferase Reporter Assay System (Promega, E1910) and Turner Biosystems luminometer (Promega) according to the supplier’s protocol. Data are expressed as relative luciferase activity (FLuc/Renilla luciferase) in arbitrary units. The experiments were repeated on 3 separate occasions.

## Results

### ELAVL4 is required for *Xenopus* oocyte maturation

ELAVL1, ELAVL2, and ELAVL4 proteins have been identified as MSI1-associated proteins by mass spectrometry in both immature and mature *Xenopus* oocytes.^20^ To determine if members of the ELAVL family are necessary for *Xenopus laevis* oocyte maturation, individual DNA antisense oligonucleotides were designed to knockdown *Elavl1*, *Elavl2*, and *Elavl4*. The specific antisense oligonucleotides were separately microinjected into oocytes, which were cultured overnight and subsequently treated with progesterone to prompt oocyte maturation. Successful oocyte maturation can be tracked by the appearance of a white spot on the animal pole of the oocyte as they undergo germinal vesicle (nuclear) breakdown (GVBD). The proportion of oocytes undergoing maturation after specific antisense injections was statistically analyzed and compared to oocytes injected with non-specific scrambled DNA oligonucleotides at two time points: 50% GVBD, where 50% of scrambled control oocytes had matured (Fig. 1A), and 100% GVBD, where 100% of scrambled control oocytes had matured (Fig. 1B) to distinguish between a delay or a block to maturation. Antisense oligonucleotides targeting knockdown of both MSI gene isoforms (*Msi1* and *Msi2)* have been previously shown to block oocyte maturation at both 50% GVBD and 100% GVBD and were used here as a technical control.^10,11^ Injection of *Elavl1* and *Elavl2* antisense oligonucleotides each resulted in a partial delay to GVBD. By contrast, injection of *Elavl4* antisense oligonucleotides robustly inhibited GVBD at both 50% GVBD (Fig. 1A) and 100% GVBD (Fig. 1B) when compared to scramble injected oocytes, an effect that was similar to the inhibition seen with combined MSI1 and MSI2 knockdown (Msi1/2 AS). Given the similarity of the effect of knockdown of ELAVL4 to knockdown of MSI, we next determined if MSI and ELAVL4 were part of a common pathway.

**Figure 1.**
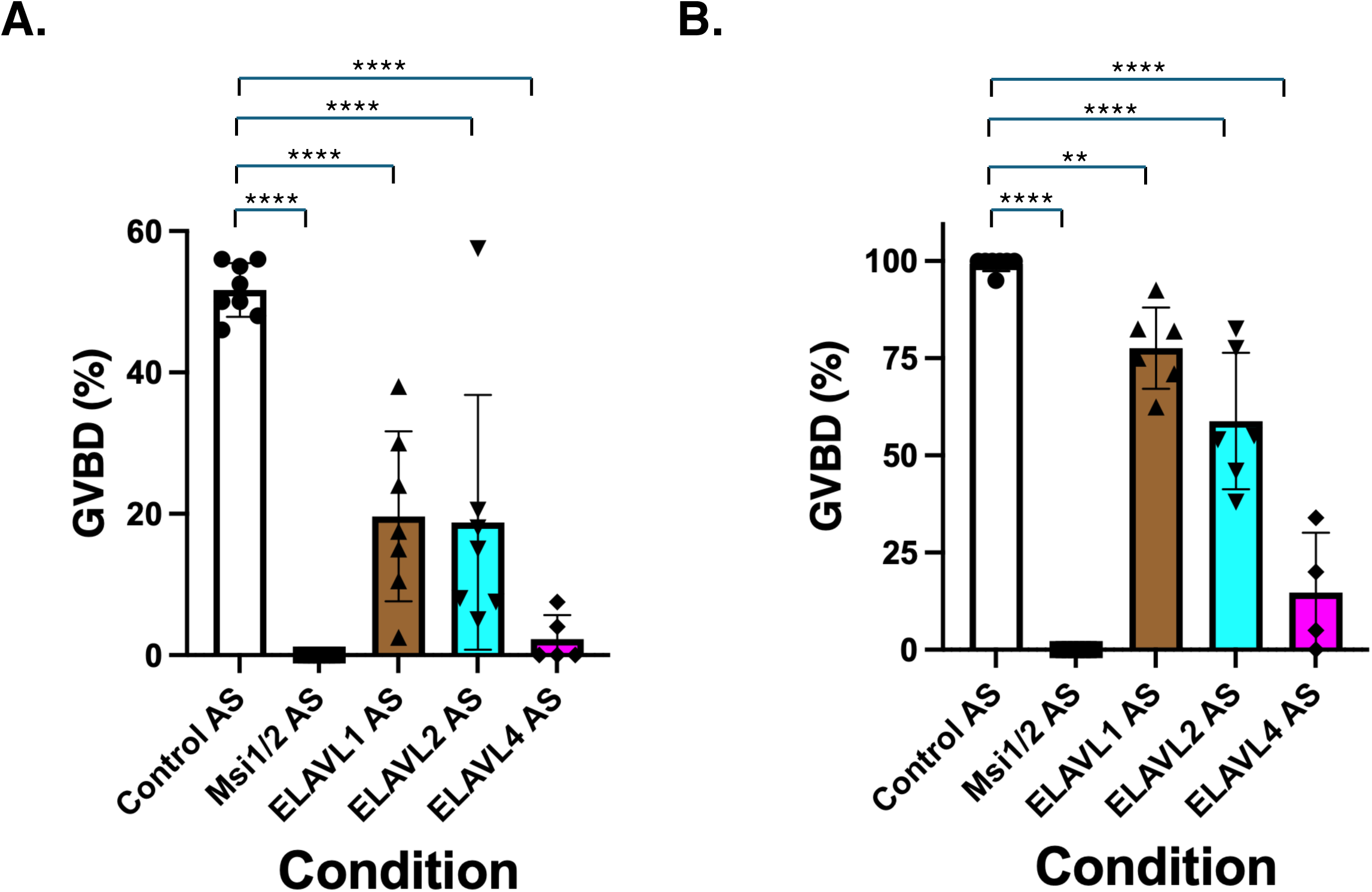
Knockdown of ELAVL4 attenuates *Xenopus* oocyte maturation. (A) Immature oocytes were microinjected with antisense oligonucleotides targeting *Elavl4*, both *Msi1* and *Msi2* (Msi1/2AS), or scrambled control (Control AS) as indicated. Following incubation overnight, oocytes were stimulated with progesterone. At the timepoint when 50% (A) or 100% (B) of Control (scrambled antisense-injected) oocytes had undergone germinal vesicle breakdown (GVBD), the degree of maturation (GVBD) in the experimental groups was assessed. The statistical significance over multiple independent experiments is indicated: **, p<0.005, ****, p<0.0001.

### ELAVL4 is required for MSI1-mediated target mRNA translation

MSI1 is necessary for the polyadenylation and translation of the *Mos* and *Cyclin B5* mRNAs, that ocurrs immediately following progesterone stimulation.^10,11^ RNA ligation-coupled PCR analysis using primers specific to the endogenous *Mos* mRNA revealed that ELAVL4 knockdown inhibited the polyadenylation of the *Mos* transcript (Fig. 2A) with a correlated inhibitory effect upon Mos protein accumulation (Fig. 2B). These effects mirrored those of MSI1 and MSI2 knockdown which also showed a block to maturation, *Mos* mRNA polyadenylation and Mos protein accumulation.^10,11^

**Figure 2.**
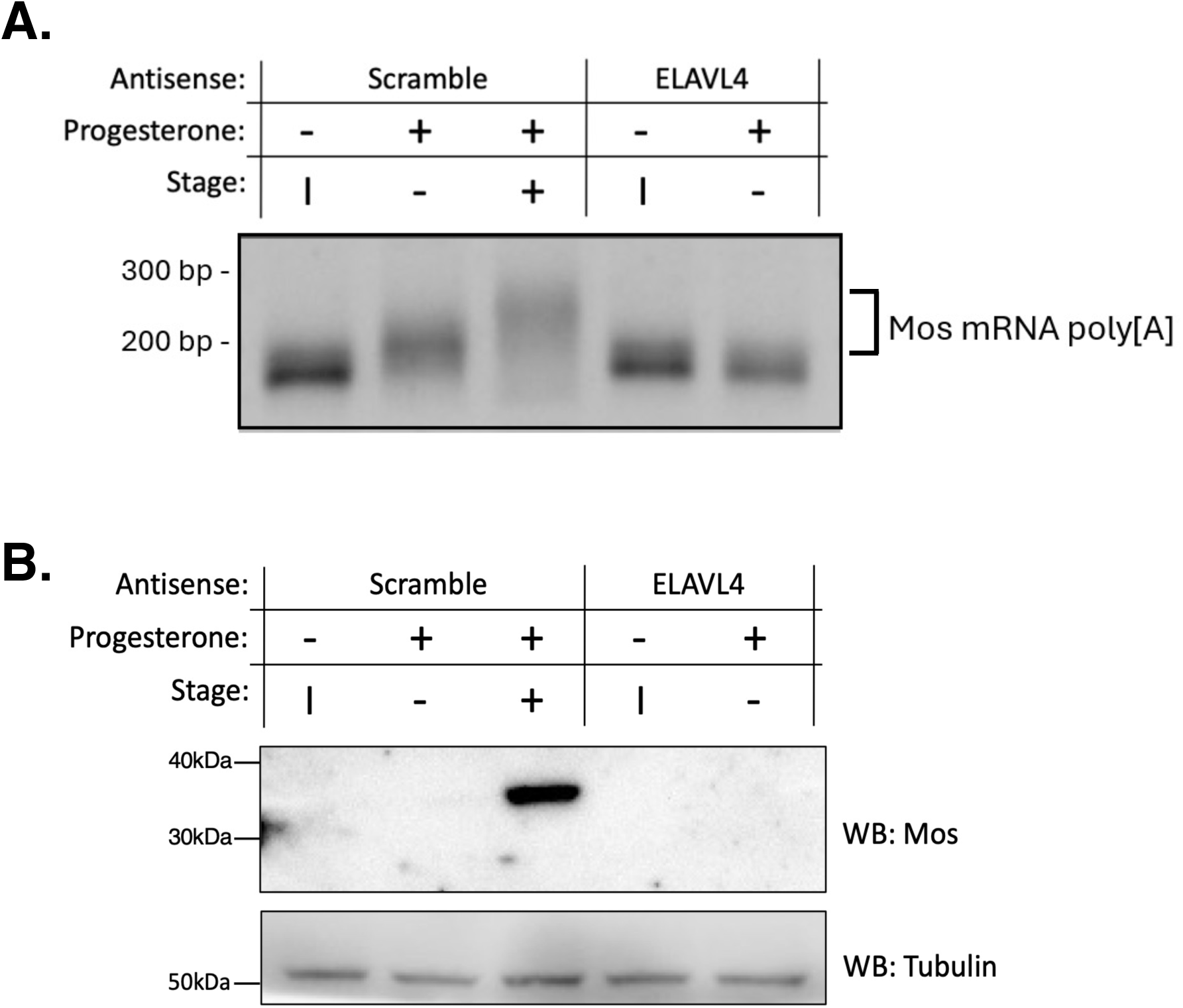
Knockdown of *Elavl4* ablates *Mos* mRNA polyadenylation and translation. (A) Immature oocytes were microinjected with control (scrambled) or *Elavl4* antisense oligonucleotides as indicated. Following incubation overnight, oocytes were (+) or were not (-) stimulated with progesterone. The progesterone-dependent polyadenylation of the endogenous *Mos* mRNA was assessed by PCR, where an increase in PCR product indicates polyadenylation. In control (scramble) oocytes, Mos mRNA polyadenyation (bracket) is observed in progesterone stimulated oocytes prior to GVBD (Stage -) and more robustly after GVBD (Stage +) relative to immature oocytes (Stage I). By contrast, *Elavl4* antisense oligonucleotide injection attenuated the polyadenylation of *Mos* mRNA and prevented GVBD (Stage -). (B) *Elavl4* antisense oligonucleotide injection blocked progesterone-dpendent Mos protein synthesis, as determined by western blot (WB: Mos). Tubulin levels in the same protein lysates serve as lane loading controls (WB: Tubulin)

To verify the specificity of the ELAVL4 antisense knockdown and rule out possible off-target effects of the ELAVL4 antisense oligonucleotide, the oocytes were co-injected with the ELAVL4 antisense and exogenous RNA encoding GST-tagged ELAVL4. The *Elav4* mRNA expression construct was designed to contain point mutations in the third “wobble” nucleotide position in the codons that are targeted by the *Elavl4* antisense oligonucleotide, to prevent antisense targeting and mRNA knockdown. Co-injection of *Elavl4* antisense with RNA encoding the wobble rescue ELAVL4 protein (GST-XeELAVL4ΔWobble) was able to rescue oocyte maturation in a statistically significant manner, albeit not to the extent seen in the scrambled control antisense samples (Fig. 3A). The GST-ELAVL4 expression construct also resulted in recovery of *Mos, Cyclin B5, and Cyclin B1* polyadenylation (Fig. 3B) and accumulation of Mos protein levels (Fig. 3C). The results indicate that ELAVL4 is necessary for MSI1-mediated translational activation of target mRNAs (*Mos and Cyclin B5*) and the downstream CPEB-target mRNA, *Cyclin B1*.^50^

**Figure 3.**
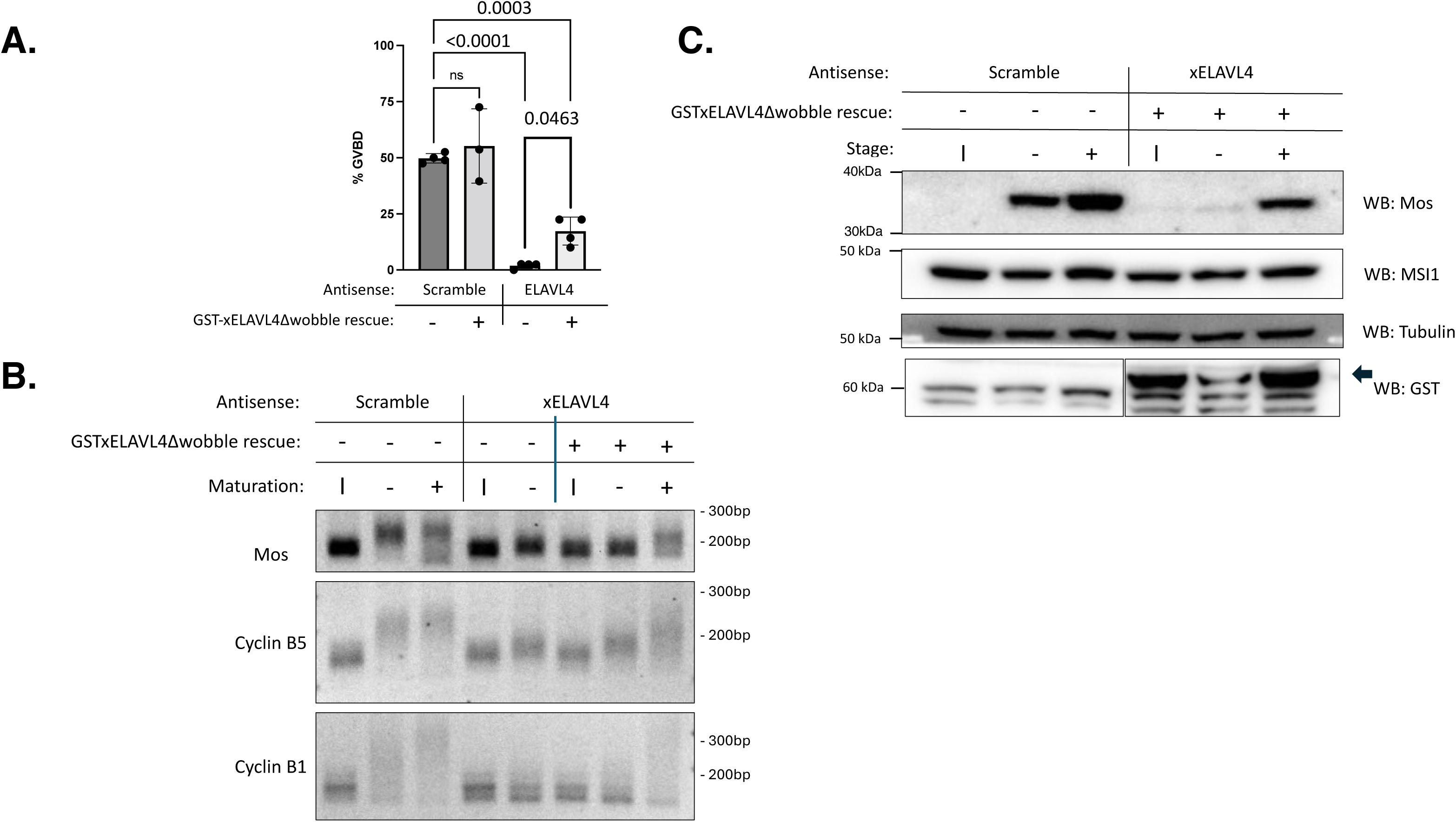
ELAVL4 knockdown phenotype is reversed by ELAVL4Δwobble expression. (A) Immature oocytes were injected with the indicated antisense oligonucleotides and either water (-) or RNA encoding a GST tagged ELAVL4 rescue contruct (GST-xELAVL4Δwobble) as indicated. Following overnight incubation, oocytes were stimulated with progesterone and the extent of maturation (GVBD) assessed when 50% of control oocyte reached GVBD. Statistically significant p-values are as indicated. Not significant = ns. (B) The oocytes injected in (A) were assessed for the progesterone-dependent polyadenyation of the early class *Mos* and *Cyclin B5* mRNAs as well as the late class, *Cyclin B1* mRNA. An increase in PCR product size over the size in immature oocytes is indicative of polyadenylation. (C) The oocytes injected in (A) were analysed by western blot (WB) for endogenous Mos, MSI1 and Tubulin levels as indicated, as well as levels of the injected GST tagged xELAVL4Δwobble protein (arrowhead). Stage refers to whether oocytes had completed GVBD. I = immature oocytes.

### ELAVL4 co-associates with MSI in an RNA-independent manner

To determine if ELAVL4 binding to MSI1 was direct or indirectly involved co-occupancy on the same target mRNAs, RNA encoding the GST-tagged, Xenopus ELAVL4 protein (GST-XeELAVL4) was co-injected with RNA encoding GFP-tagged, Xenopus MSI1 (GFP-XeMSI1) or GFP-tagged, Xenopus MSI2 (GFP-XeMSI2) proteins into immature *Xenopus* oocytes. Following overnight incubation, GST-xELAVL4 protein was captured with glutathione Sepharose beads. During the bead wash steps, the samples were incubated with RNase1 to degrade endogenous co-associated RNAs. After further wash steps, MSI protein associated with the GST-ELAVL4 protein complex was identified by gel electrophoresis and Western blotting. GFP-tagged and GST-tagged proteins were also identified in the input *Xenopus* lysates. The Western blot of the elute from the GST pulldown revealed that ELAVL4 specifically associates with MSI1, and to a lesser extent MSI2, and that these associations are preserved in the absence of RNA (Figure 4, upper panel).

**Figure 4.**
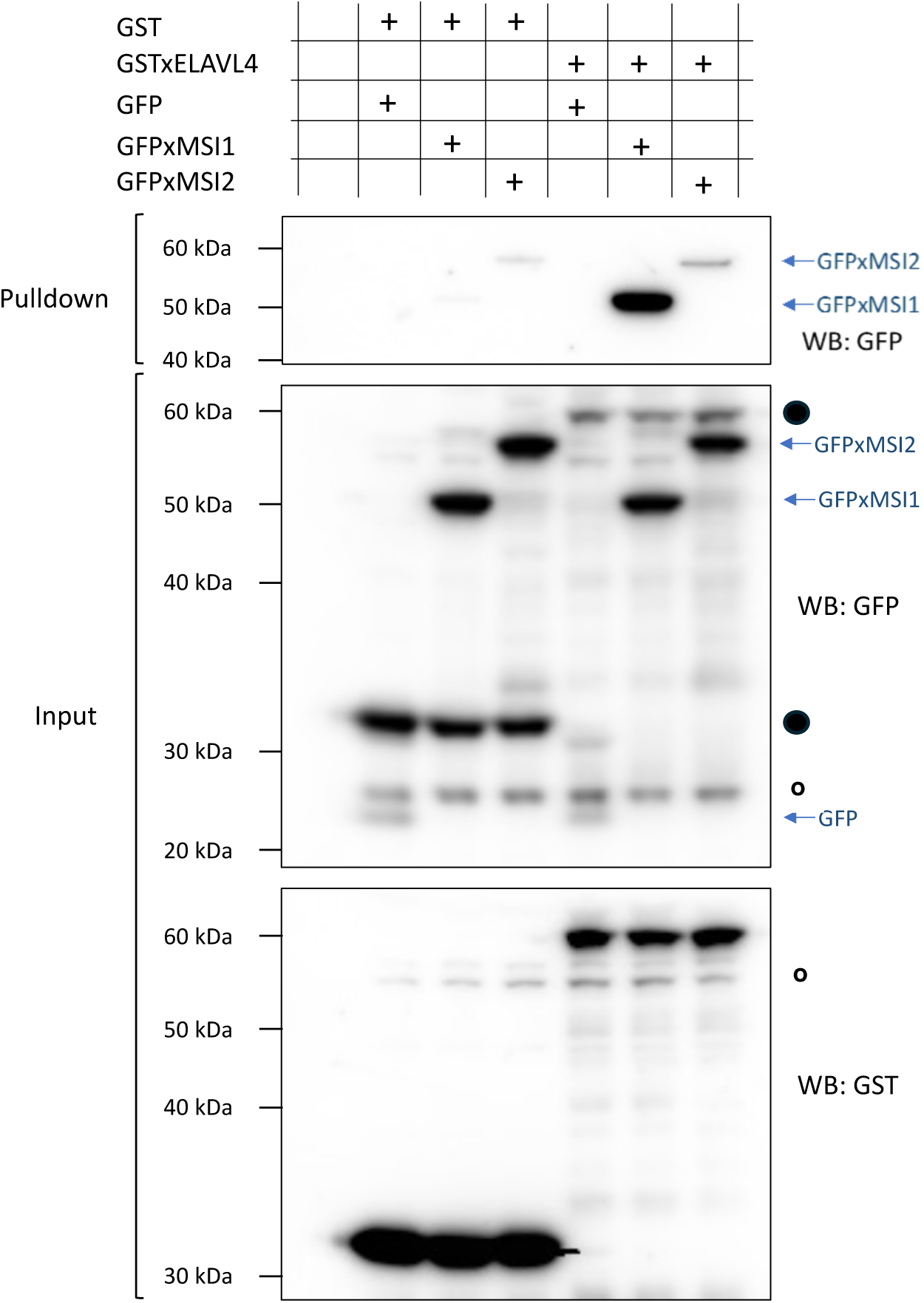
ELAVL4 association with MSI1 and MSI2 is independent of RNA binding. Oocytes were co-injected with RNA encoding either the GST moiety alone (GST) or GST-tagged xELAVL4 and either RNA encoding the GFP moiety alone, GFP-tagged xMSI1 or xMSI2 and incubated overnight. The next morning, the oocytes were lysed and GST proteins recovered using Glutathione-sepharose in the presence of RNase A to degrade RNA. The samples were analyzed by western blot for association of GFP-tagged proteins (Pulldown). The input lysates were analyzed for expression of the tagged protein contructs by first performing a GST western blot (lower Inout panel) and subsequently washed and re-probed for GFP protein expression. Black circles indicate residual GST signals, open circles indicate non-specific bands.

### The N-terminal domain of MSI1 functionally interacts with the C-terminal domain of ELAVL4

We next identified the regions of MSI and of ELAVL4 that mediated the protein-protein interaction, using GST- or GFP-tagged truncated versions of XeELAVL4, XeMSI and mouse MSI 1 (mMSI1) (Fig. 5). ELAVL4 contains three RRMs which are highly homologous between ELAVL family members.^51^ RRM1 and RRM2 are located in the N-terminal of the protein, while RRM3 and a hinge region are in the C-terminal region (Fig. 5A).^23,24^ RRM1 and RRM2 of ELAVL family members have been shown to interact directly with target RNAs, while RRM3 has been shown to stabilize protein-protein interactions in concert with the hinge region between RRM2 and RRM3.^24,52^ The N-terminal domains of MSI1 and MSI2 contain highly conserved RNA-recognition motifs (RRMs) which allow MSI to bind to target mRNAs in a sequence-specific manner, whereas the MSI1 and MSI2 C-terminal domains contain intrinsically disordered regions and are less conserved (Fig. 5B).^53,54^

**Figure 5.**
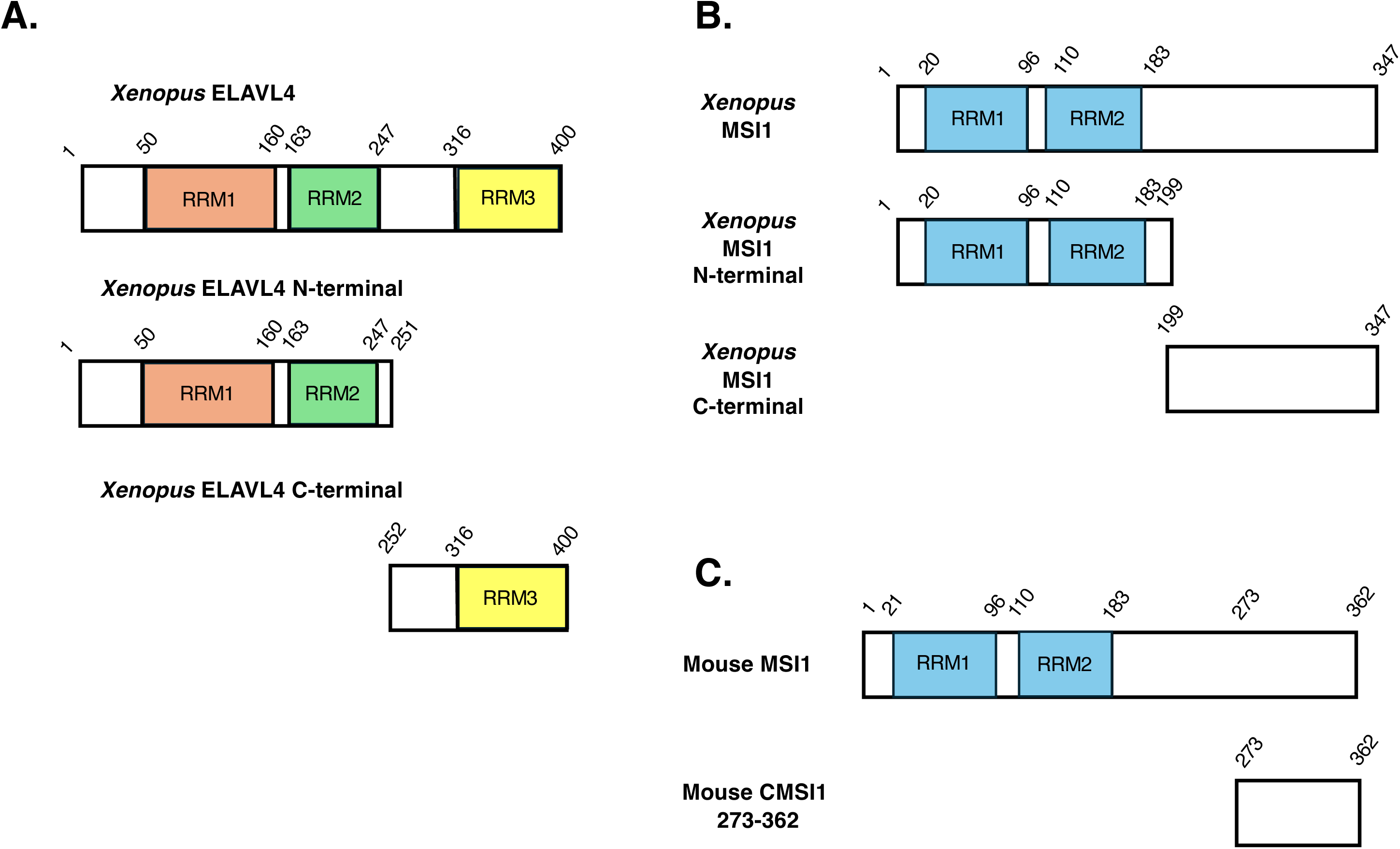
Schematic representations of the (A) *Xenopus* ELAVL4 and (B) *Xenopus* MSI1, and (C) Mouse MSI1 constructs used in association tests. Amino acid numbers are shown above each construct. (A) The relative positions of the three Xenopus ELAVL4 RNA recognition motifs (RRM1, RRM2 and RRM3) are shown in the full length protein (amino acids 1 to 400), as well as in the N-terminal and C-terminal deletion mutants (amino acids 1 to 251 and 252 to 400, respectively). (B) The relative positions of the two Xenopus MSI1 RNA recognition motifs (RRM1 and RRM2) are shown in the full length protein (amino acids 1 to 347), as well as in the N-terminal and C-terminal deletion mutants (amino acids 1 to 199 and 199 to 347, respectively). (C) The relative positions of the two mouse MSI1 RNA recognition motifs (RRM1 and RRM2) are shown in the full length protein (amino acids 1 to 362). The C-terminal C-MSI1 protein lacks the RRMs (amino acids 273 to 362).

ELAVL4 interacted with the N-terminal domain of MSI1 and had negligible interaction with the C-terminal domain of MSI1 (Fig. 6 A-D) . To further validate the observed ELAVL4 interaction with the N-terminal domain of MSI1, we tested association with the naturally ocurring trucated MSI1 isoform, C-MSI1 (C-MSI1), which has an identical amino acid sequence in both mice and humans. C-MSI1 was originally identified in mouse and human embryonic stem cells^17^ where it is necessary for embryonic stem cell pluripotency. C-MSI1 does not contain the MSI N-terminal RRM domains, and consistent with a requirement for the MSI N-terminal domain for ELAV4 association, C-MSI1 did not interact with ELAVL4 (Fig. 6 E-H).

**Figure 6.**
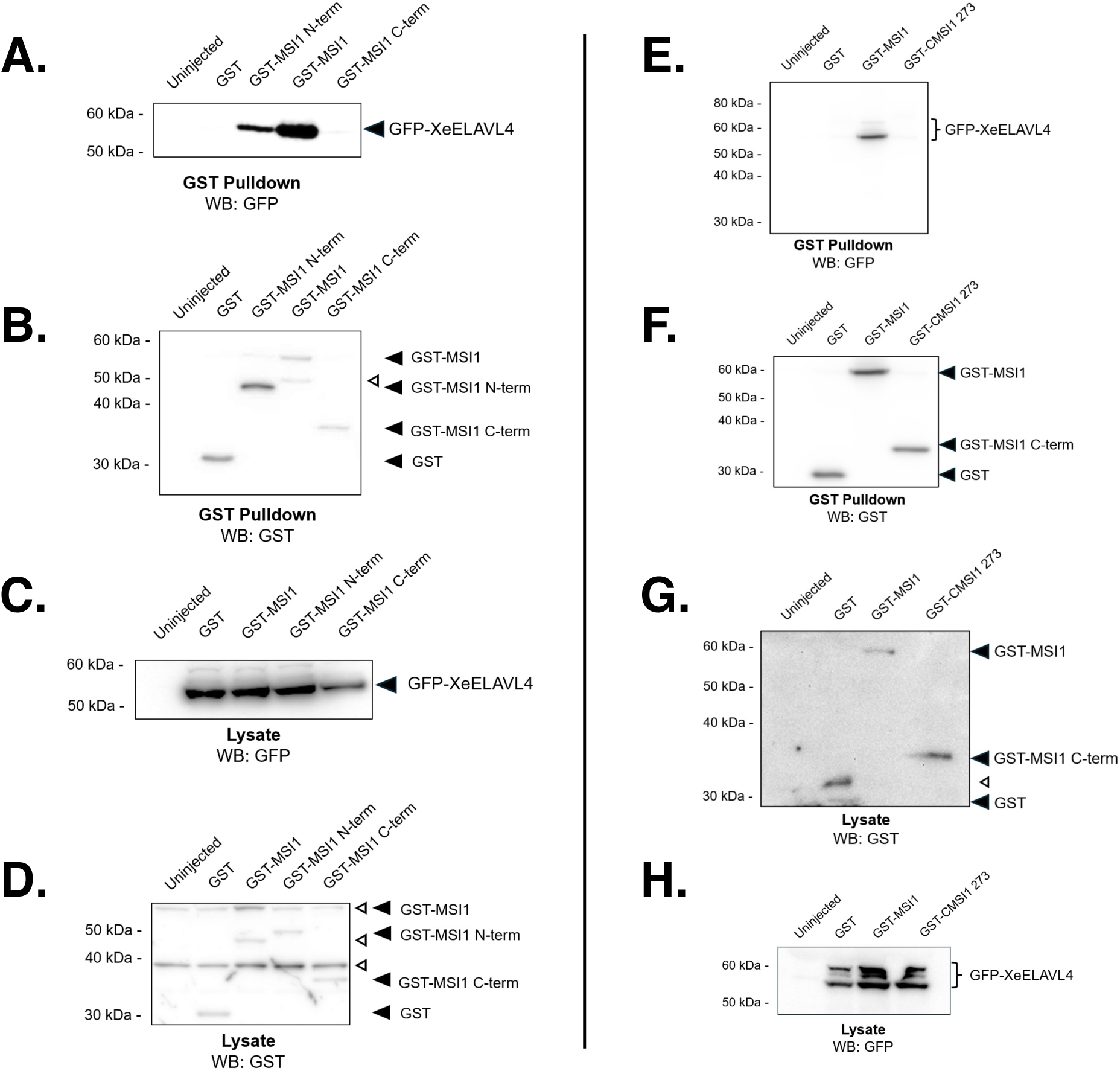
ELAVL4 associates with the N-terminal domain of MSI1. (A) Immature oocytes were co-injected with RNA encoding GFP-tagged Xenopus ELAVL4 (GFP-XeELAVL4) and either RNA encoding the GST moiety alone (GST), GST-tagged *Xenopus* MSI1 (GST-MSI1), GST-tagged *Xenopus* N-terminal MSI1 (GST-MSI N-term), or GST-tagged *Xenopus* C-terminal MSI1 (GST-MSI1 C-term) and incubated overnight. The next morning, the oocytes were lysed and GST proteins recovered using Glutathione-sepharose in the presence of RNase A to degrade RNA. The samples were analyzed by western blot for association of GFP-tagged XeELAVL4 protein (GST pulldown). (B) The blot in (A) was re-probed for GST to quantitate the level of recovery of the expressed GST-tagged proteins (marked with closed arrowheads). The open arrowhead indicates a non-specific band. (C) A portion of the input lysates used for the pulldown in panels (A) and (B) were analyzed by western blot for GFP-XeELAVL4 expression. (D) The input lysate blot from panel (C) was re-probed for relative GST protein expression (closed arrowheads). Non-specific bands are indicated by open arrowheads. (E-H) Immature oocytes were separately co-injected with RNA encoding the GFP-XeELAVL4 and either RNA encoding the GST moiety, GST-tagged mouse MSI1 (GST-MSI1), or GST-tagged mouse C-MSI1 273-362 (GST-CMSI1 273-362) and analyzed as above (A-D).

For the reciprocal analyses, we injected immature oocytes with RNA encoding GST-XeMSI1 and either the GFP moiety alone, GFP-XeELAVL4 full length, GFP-XeELAVL4-N-terminal or GFP-XeELAVL4-C-terminal proteins. Following GST pulldown and RNase1 treatment, the full length GFP-xELAVL4 and the C-terminal GFP-xELAVL4-C were both detected in complex with full-length GST-MSI1 (Fig. 7). Negligible interaction was detected between the N-terminal domain of ELAVL4 and GST-MSI1 (Fig. 7). We conclude that ELAVL4 interacts with MSI1 primarily via the ELAVL4 C-terminal domain, which contains the hinge region and RRM3.

**Figure 7.**
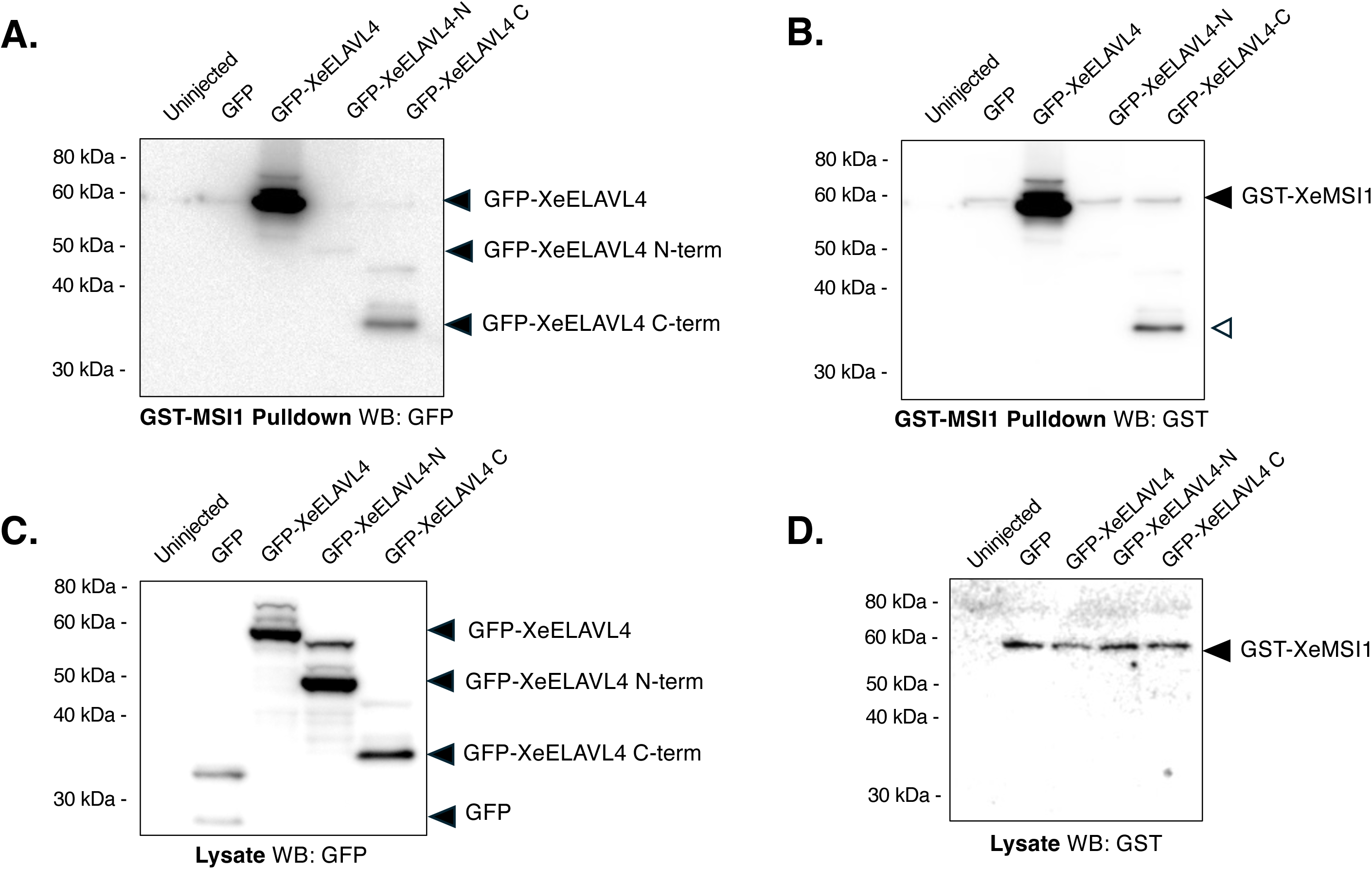
MSI1 associates with the C-terminal domain of ELAVL4. (A) Immature oocytes were co-injected with RNA encoding GST-XeMSI1 and RNA encoding either the GFP moeity alone (GFP), GFP-tagged *Xenopus* ELAVL4 (GFP-ELAVL4), GFP-tagged *Xenopus* N-terminal ELAVL4 (GFP-ELAVL4-N), or GFP-tagged *Xenopus* C-terminal ELAVL4 (GFP-ELAVL4-C) and incubated overnight. The oocytes were then lysed and subjected to glutathione Sepharose pulldown (essentially as described in the legend to Figs 4 and 6). The pulldown samples were analyzed by western blot for co-association of GFP-tagged proteins (A) and the blot re-probed to show expression of GST-XeMSI (B). Note the GST-XeMSI1 protein runs at a similar size as the GFP-XeELAVL4 protein. A residual GFP-XeXeELAVL4 C-term signal is also observed (open arrowhead). A portion of the input lysates used for the pulldown in panels (A) and (B) were analyzed by western blot for GFP tagged protein expression (C) and re-probed to detect the relative expression levels of GST-XeMSI1 (D).

### MSI associates with over 200 proteins in the anterior pituitary

In earlier work, we identified the MSI-associated proteins in immature and mature *Xenopus* oocytes.^20^ Here we identified MSI-associated proteins in the mouse pituitary, where we have previously determined that MSI1 is required for robust gonadotrope function and LH surge levels^20^ (Figure 8A). Mouse pituitaries were collected, pooled and then split into two aliquots. One aliquot was immunoprecipitated with IgG control antibodies, while the other aliquot was immunoprecipitated with a cocktail of MSI1 and MSI2 antibodies. The experiment was repeated three times and the recovered protein samples analyzed by mass spectrometry (Supplementary Table 1). We identified 256 proteins specifically enriched for interaction with MSI over the IgG control, with a log fold change greater than or equal to 2 (Supplementary Table 2). Associated proteins included previously identified MSI-interacting partner proteins e.g. AGO2^55^ and PABP4 (PABPC4)^20^, and revealed that the ELAVL1 protein associates with MSI1 and MSI2 in mouse pituitary (Fig 8B). In addition, other previously identified MSI-interacting proteins from our study in *Xenopus* oocytes were also found to interact with MSI in the mouse pituitary, e.g. PURA, UPF1, hnRNPC, RPS27, MOV10 (Fig. 8B). Gene ontology (GO) analysis of co-associated proteins, implicated a number of cellular processes including cytoplasmic mRNA translation, vesicle function and cell cycle progression (Fig. 8C).

**Figure 8.**
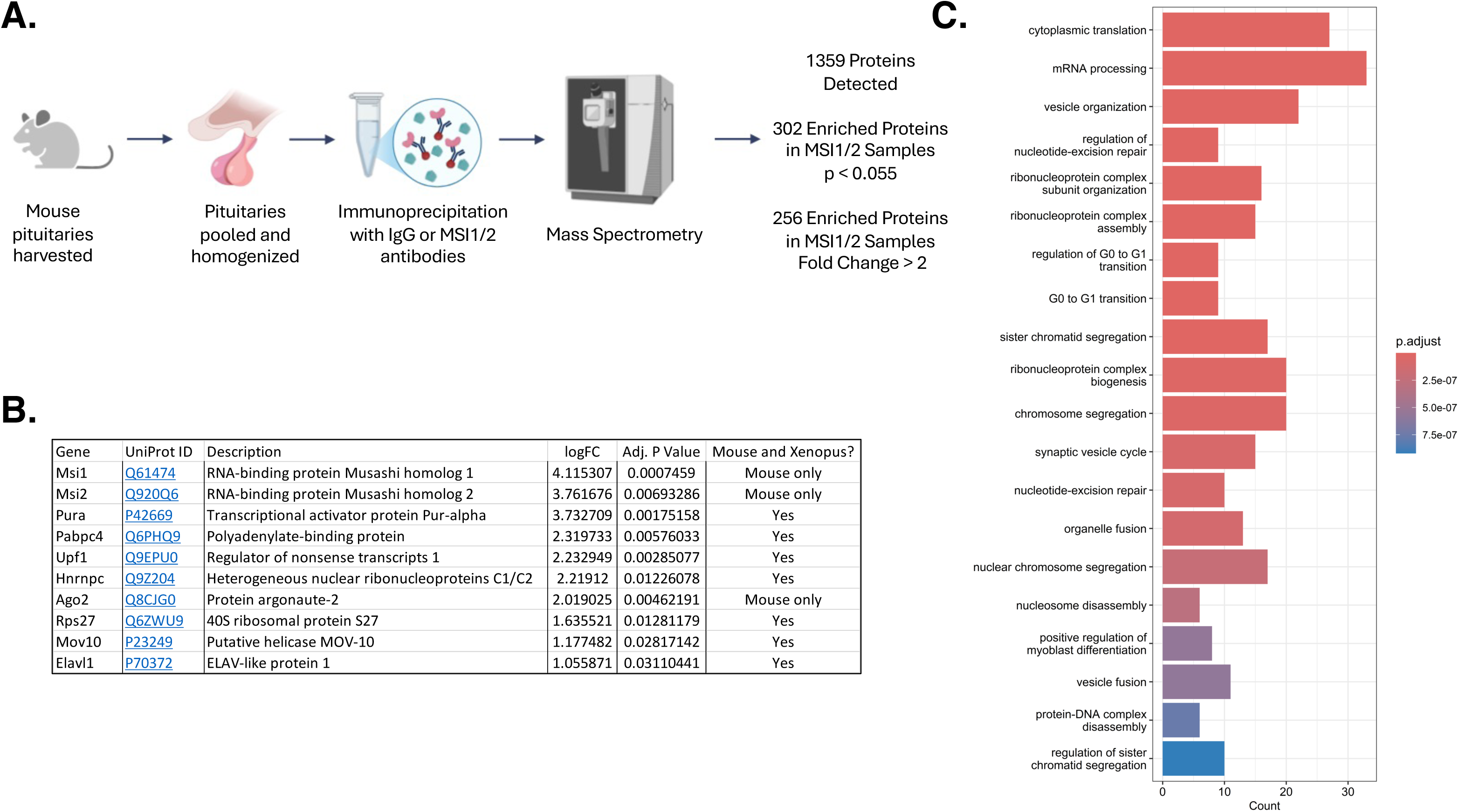
The MSI1 and MSI2 protein interactome in the mouse pituitary. (A) Schematic representation of the experimental strategy employed. Mouse pituitary lysates (see Methods) were incubated with either control IgG or a combination of anti-MSI1 and anti-MSI2 antibodies. The immunoprecipitated protein complex was quantified by mass spectrometry. Of the total 1359 proteins detected, 302 proteins were significantly enriched in the anti-MSI1 and anti-MSI2 complex compared to the IgG control complex (p < 0.055). Of these proteins, 256 were enriched at fold change ≥ 2. (B) An number of known MSI1 and MSI2-associated proteins were detected, including some that were previously identified as part of the MSI1 and MSI2 interactome in *Xenopus* oocytes. (C) Gene Ontology analysis of MSI1 and MSI2 co-associated proteins in the mouse pituitary.

### Mammalian ELAVL1 interacts with MSI1 in an RNA-independent manner and can functionally compensate for ELAVL4 knockdown in *Xenopus* oocytes

To confirm an interaction of mammalian ELAVL1 with mammalian MSI1, we co-injected *Xenopus* oocytes with RNA encoding GFP-human MSI1 (GFP-hMSI1) with RNA encoding either the GST moiety alone or GST-human ELAVL1 (GST-hELAVL1). Following GST pulldown and treatment with RNase1, retained proteins were analyzed by western blotting. A specific, RNA-independent interaction was detected between human ELAVL1 and human MSI1 (Fig. 9A).

**Figure 9.**
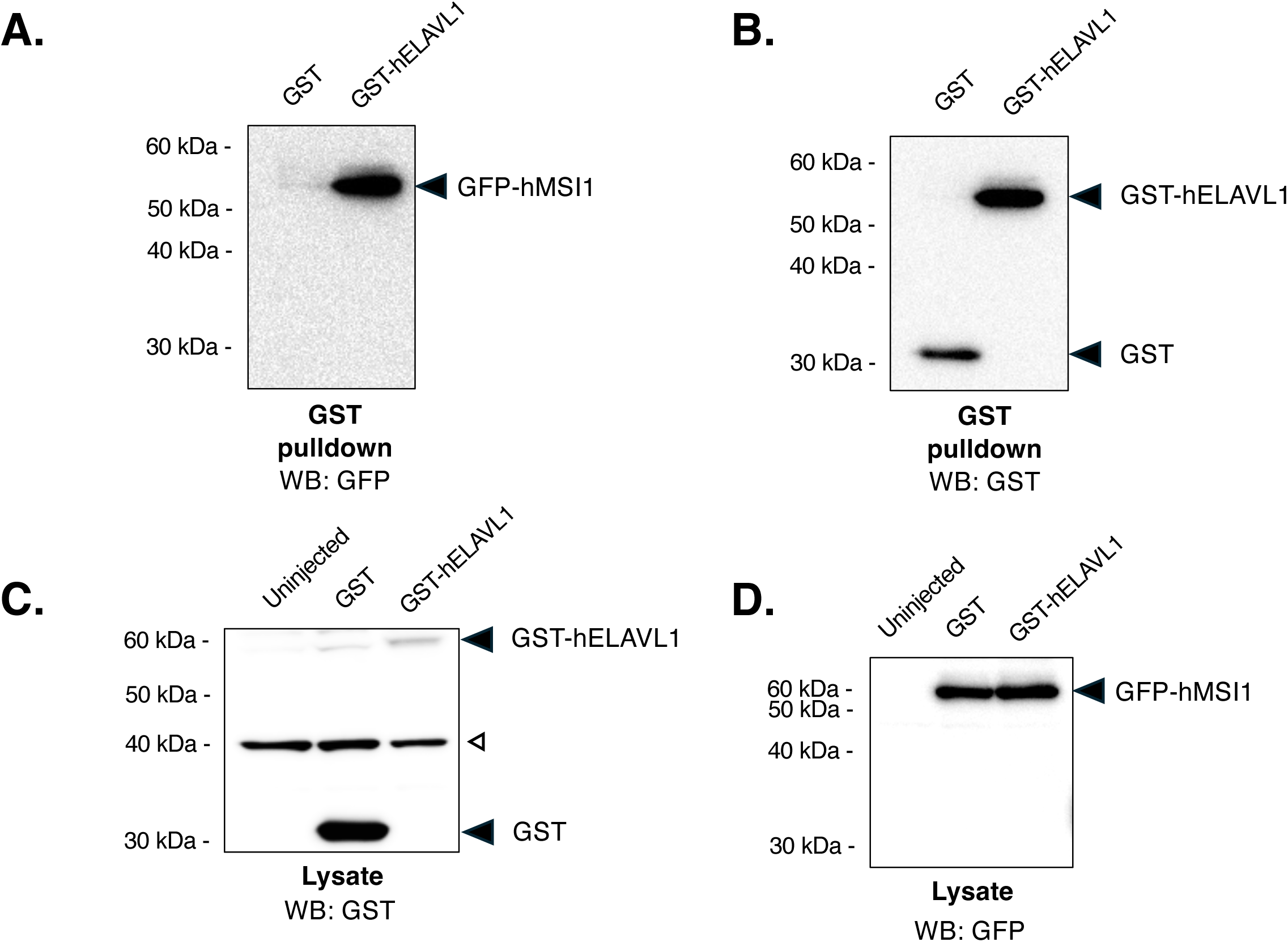
Human MSI1 associates with Human ELAVL1. (A-D) Immature oocytes were coinjected with RNA encoding GFP-tagged human MSI1 (hMSI1) and either RNA encoding the GST moiety (GST) or GST-tagged human ELAVL1 (GST-hELAVL1), and incubated overnight. The oocytes were then lysed and subjected to glutathione Sepharose pulldown (essentially as described in the legend to Figs 4,6 and 7). The pulldown samples were analyzed by western blot for co-association of GFP-hMSI1 (A) and the blot re-probed to show expression of GST and GST-hELAVL1 (B). A portion of the input lysates used for the pulldown in panels (A) and (B) were analyzed by western blot for GST-tagged protein expression (C) and re-probed to detect the relative expression levels of GFP-hMSI1 (D).

We next wanted to know if human ELAVL1 could functionally compensate for ELAVL4 in *Xenopus* oocytes. Oocytes were co-injected with ELAVL4 antisense oligonucleotides and either RNA encoding GST-XeELAVL4, GST-huELAVL1 or the GST moiety alone. The next morning, the oocytes were stimulated with progesterone and the ability to undergo oocyte maturation assessed. The maturation of oocytes treated with *Elavl4* antisense could be rescued in a statistically significant manner by either the addition of *Xenopus* ELAVL4 or human ELAVL1 (Fig. 10A).

**Figure 10.**
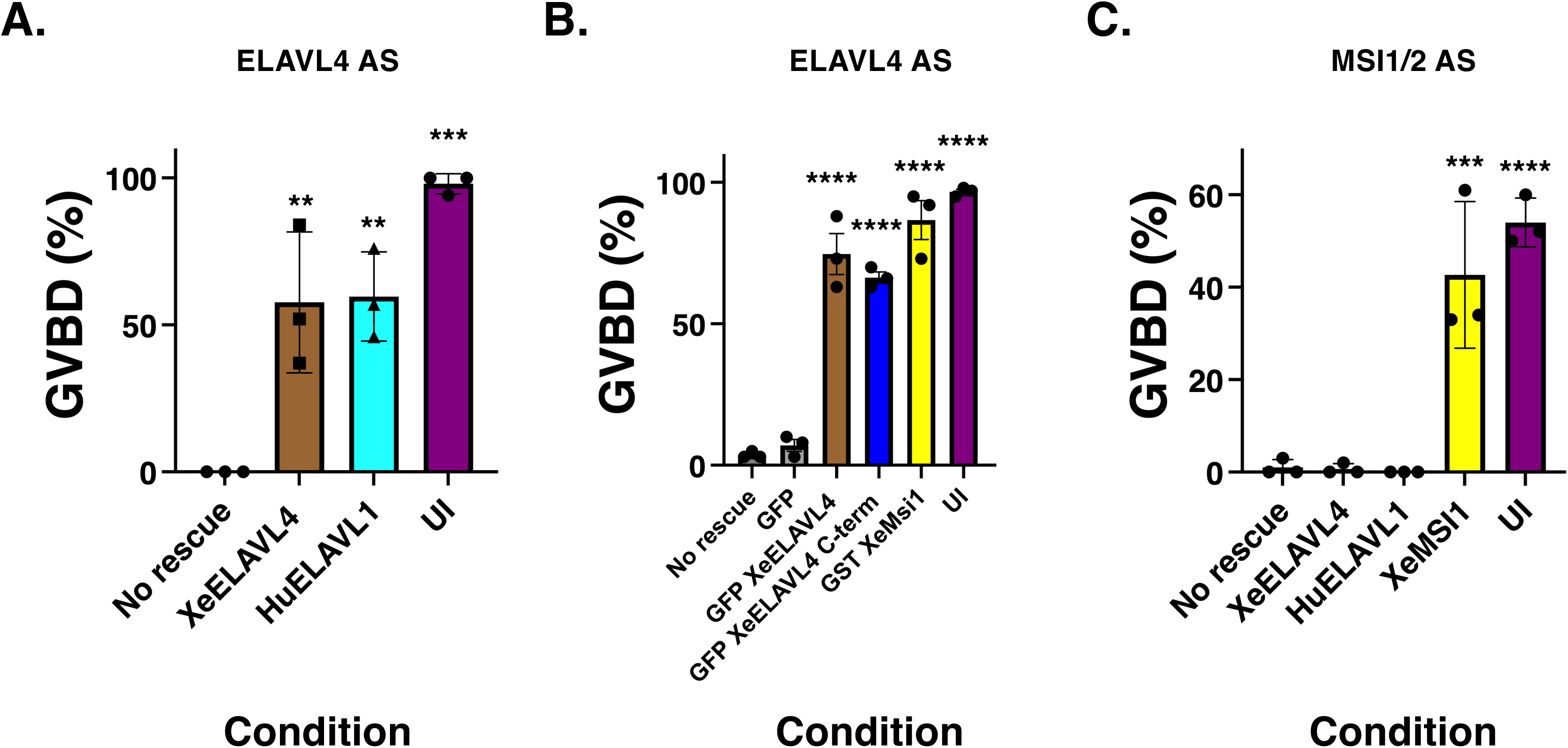
Human ELAVL1 substitutes for Xenopus ELAVL4 in oocyte maturation, and also requires MSI1. (A) Immature oocytes were microinjected with antisense oligonucleotides targeting Elavl4 and either water (No rescue) or RNA encoding GST-XeELAVL4 or GST-hELAVL1 as indicated. Following overnight incubation, oocytes were stimulated with progesterone and the extent of maturation (GVBD) assessed when 100% of control (UI) oocytes completed GVBD. Uninjected oocytes served as an internal control for progesterone stimulation (UI). Statistically significant p-values relative to the no rescue control are indicated. (B) Similar to (A), immature oocytes were microinjected with antisense oligonucleotides targeting Elavl4 and either water (No rescue) or RNA encoding the indicated GFP- or GST-tagged proteins. Following overnight incubation, oocytes were stimulated with progesterone and the extent of maturation (GVBD) assessed when 100% of control (UI) oocytes completed GVBD. Uninjected oocytes served as an internal control for progesterone stimulation (UI). Statistically significant p-values relative to the no rescue control are indicated. (C) Immature oocytes were microinjected with antisense oligonucleotides targeting Msi1 and Msi2 (MSI1/2 AS) and either water (No rescue) or RNA encoding the indicated GST-tagged proteins. Following overnight incubation, oocytes were stimulated with progesterone and the extent of maturation (GVBD) assessed when 50% of control (UI) oocytes completed GVBD. Uninjected oocytes served as an internal control for progesterone stimulation (UI). Statistically significant p-values relative to the no rescue control are indicated. The statistical significance over multiple independent experiments is indicated: **, p<0.005, ***, p<0.001, ****, p<0.0001.

Since MSI1 interacts primarily with the C-terminal domain of ELAVL4 (Fig. 7), we were curious if ectopic expression of the ELAVL4 C-terminal domain alone would be sufficient to effect rescue of ELAVL4 antisense oligonucleotide-injected. To this end, oocytes were co-injected with ELAVL4 antisense oligonucleotides and either RNA encoding the GFP moiety alone, GFP-XeELAVL4, GFP-XeELAVL1 C-terminal domain or GFP-XeMSI1. The next morning, the oocytes were stimulated with progesterone and the ability to undergo oocyte maturation assessed. Similar to the full length XeELAVL4 protein, expression of the XeELAVL4-C-terminal domain was able to exert rescue of maturation, consistent with a MSI-dependent interaction and recruitment to target mRNAs. Moreover, ectopic MSI1 expression was able to rescue oocyte maturation in *Elavl4* antisense-injected oocytes (Fig. 10B). As an additional specificity control, we injected oocytes with MSI1/2 antisense oligonucleotides alone or co-injected oocytes with RNA encoding Xenopus ELAVL4, human ELAVL1, or *Xenopus* MSI1 and cultured overnight. Following progesterone stimulation the next morning, ELAVL1 and ELAVL4 were unable to rescue maturation. By contrast, *Xenopus* MSI1 was able to robustly rescue oocyte maturation as expected (Fig. 10C). Together, these results suggest a dependence on MSI for ELAVL family members to mediate progesterone-stimulated oocyte maturation.

We have recently reported that MSI activates the translation of a subset of target mRNAs in the pituitary, including the mRNA encoding the pituitary lineage specification factor PROP1.^12^ Western blot analyses of endogenous ELAVL1 revealed expression in the mouse NIH3T3 cell line used for our mRNA translation reporter assays (Figure 11). We next wanted to assess a possible requirement for ELAVL1 in mammalian MSI1-dependent translational activation of a luciferase reporter mRNA under the control of the murine *Prop1* mRNA 3’ UTR. The ability of MSI1 to promote luciferase-*Prop1* reporter mRNA translation was determined in the presence of either a control scramble siRNA (scr) or two independent siRNAs targeting the endogenous *Elavl1* mRNA (ELAVL1.1 or ELAVL1.3 siRNA). Expression of MSI1-GFP in the presence of the control siRNA increased translation of the *Prop1* reporter mRNA by an average of 1.85 (Figure 11A, scr MSI WT upper graph) compared to expression of the GFP moiety alone (scr + empty vector). However, expression of MSI1-GFP with either of the ELAVL1 siRNAs ablated the MSI1-dependent increase in firefly luciferase activity (Figure 11A). Western blots prepared from the same lysates used in the reporter assays confirmed that ELAVL1.1 or ELAVL1.3 siRNAs attenuated levels of endogenous ELAVL1 protein (70% reduction and 80% reduction, respectively), while the control siRNA had no effect on ELAVL1 protein levels (Figure 11B). GAPDH served as a loading control in the western blot experiments. We conclude that ELAVL1 is necessary for MSI1-dependent translational activation controlled by the mouse *Prop1* mRNA 3’ UTR.

**Figure 11.**
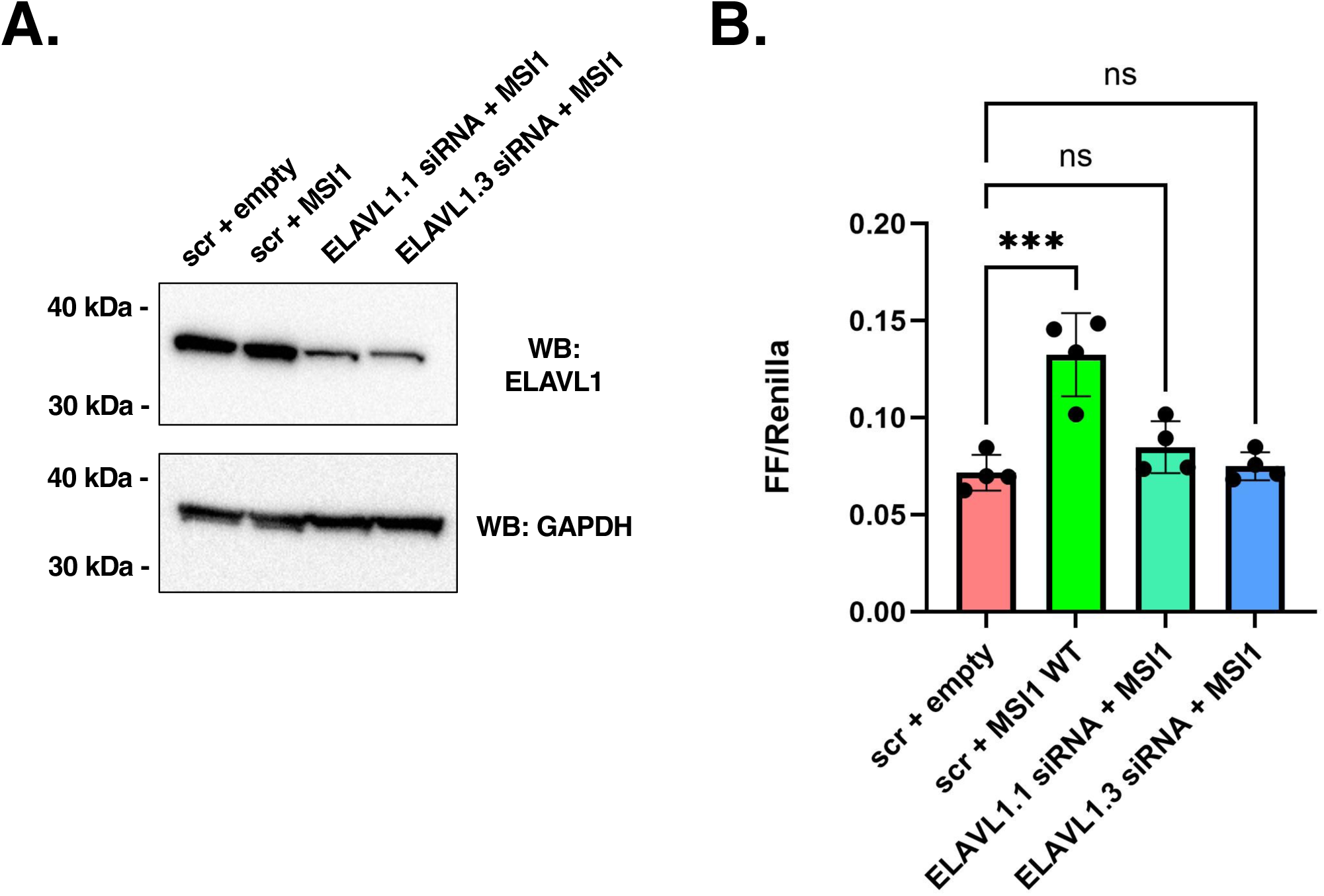
Mammalian ELAVL1 is required for MSI1-dependent mRNA translational activation of a pituitary *Prop1* 3’ UTR reporter mRNA in NIH3T3 cells. (A) NIH3T3 cells were initially transfected with either scrambled control siRNA or two separate siRNAs (mm.RI.Elavl1.13.1 and mm.RI.Elavl1.13.3) targeting the murine Elavl1 mRNA. After 48 hours, the cells were transfected with a luciferase reporter mRNA under the control of a Prop1 3’ UTR along with either the empty pEGFPN1 vector or the peGFPN1 vector containing murine Msi1. Cells were lysed and the relative levels of endogenous ELAVL1 protein determined by western blot in cells transfected with the scrambled siRNA and empty pEGFPN1 vector (scr + empty), scrambled siRNA and the peGFPN1 vector containing murine Msi1 (scr + MSI1), or cells expressing MSI1 and one of the two siRNAs targeting Elavl1 as indicated. Endogenous levels of ELAVL1 (upper panel) and GAPDH as a loading control (lower panel) are shown. A representative experiment is shown, (B) Cells transfected as indicated in (A) were assessed for Prop1 3’ UTR reporter mRNA luciferase activity as indicated. Reporter assays were repeated on three separate occasions and statistical significance assessed by one way ANOVA: ****, p<0.001; ns, not significant.

## Discussion

Here, we have identified a requirement for ELAVL4 in mediating MSI-dependent polyadenylation and translational activation of the key developmental target mRNAs *Mos* and *Cyclin B5*. We further determined that ELAVL4, like MSI, is required for oocyte maturation. ELAVL4 antisense oligonucleotide knockdown abrogated the oocyte’s ability to mature in response to progesterone (Figure 1) and prevented MSI-dependent polyadenylation and translational activation of target mRNAs (Figure 2). These effects were rescued after co-injection of RNA encoding an antisense-resistant ELAVL4 expression construct, increasing polyadenylation of *Mos* and *Cyclin B5*, translation of MOS protein, in addition to recovering a statistically significant amount of oocyte maturation (Figure 3). Functional redundancy within the ELAVL family members has been previously reported.^24,56,57^ Consistent with this redundancy, expression of antisense-resistant human ELAVL1 could rescue oocyte maturation with similar efficacy to antisense-resistant *Xenopus* ELAVL4 in oocytes injected with *Elavl4* antisense oligonucleotides. A prior study has demonstrated an RNA-independent interaction between ELAVL1 and MSI2 in the control of miR- 7 biogenesis^58^, and our results extend these findings by revealing an RNA-independent interaction between ELAVL1 and MSI1 (Fig. 9). Notably, neither ectopic expression of *Xenopus* ELAVL4 or human ELAVL1 were able to rescue oocyte maturation after *Msi1/Msi2* antisense injection (Figure 10C), indicating that the ability of ELAVL4 or ELAVL1 to enable oocyte maturation is dependent upon the presence of MSI. By contrast, ectopic expression of MSI1 rescued maturation in *Elavl4* antisense-injected oocytes (Fig. 10B), suggesting that MSI was able to exert maturation independently of ELAVL4 and/or could be utilizing ELAVL1 and/or ELAVL2 in place of ELAVL4.

ELAVL4 expression has been identified as elevated in bovine oocytes relative to immature blastocysts,^59^ however, the function of ELAVL4 in oocytes has not been previously determined. During *Xenopus* oogenesis, ELAVL1 and ELAVL2 have been implicated in the localization of the *Vg1* transcript, encoding a TGFβ family signaling molecule, during oogenesis.^60,61^ ELAVL2 has been proposed to repress *Vg1* mRNA translation in stage III oocytes via self-oligomerization.^60,62,63^ However, there is also evidence that the ELAVL proteins can function *via* interaction with distinct co-associated proteins, as indicated by the absence of an oligomerization domain in the ELAVL family members, ELAVL1, ELAVL3 and ELAVL4.^62^

We determined that ELAVL4 specifically associates with MSI1 and to a lesser extent, MSI2, and that the association ocurrs independently of RNA, excluding a simple mRNA co-occupancy mechanism for the observed association (Fig. 4). We were somewhat surprised to find that ELAVL4 interacts with the N-terminal RRM-containing domain of MSI1, as most characterized protein associations ocurr with the C-terminal, regulatory domain of MSI1, with the exception of LSM14B.^13^ Reciprocally, ELAVL4 association with MSI1 oocurs via the C-terminal domain of ELAVL4, which contains the RRM3 and the hinge region between RRM2 and RRM3, that has been previously identified as mediating protein-protein interactions in ELAVL family members.^24,33,52,64,65^ Consistent with a functional role for the ELAVL4-MSI1 interaction, we observed that expression of the isolated ELAVL4-C-terminal domain was sufficient to mediate oocyte maturation in the absence of full-length ELAVL4 (Fig. 10B). These findings support a model in which ELAVL4 and MSI1 cooperate to mediate oocyte maturation.

MSI1 and MSI2 have been implicated as key post-transcriptional regulators of hormone production in the anterior pituitary.^14–16,66^ While MSI was first implicated as a repressor of pituitary target mRNA translation, our recent studies have identified the *Prop1* mRNA as being a target of MSI-dependent translational activation.^12^ Here, our *in vitro* mRNA reporter assays revealed that MSI1-dependent translational activation *via* the *Prop1* 3’ UTR was attenuated when cells were co-transfected with siRNA targeting the *Elavl1* mRNA (Figure 11). These results support the conclusion that ELAVL1 cooperates with MSI1 to promote translational activation in mammalian cells.

Our findings implicating both ELAVL1 and ELAVL4 in promoting translation of MSI1-target mRNAs are at odds with a several prior studies. Functional antagonism between MSI1 and ELAVL4 has been reported for the regulation of production of the cell cyle control protein, p21^WAF1^ (encoded by the *Cdkn1a* gene), with MSI1 repressing translation of the *Cdkn1a* mRNA and ELAVL4 promoting translation of the *Cdkn1a* mRNA.^34,36,67^ Similarly, ELAVL1 and MSI1 appear to exert opposite control over translation of the mRNA encoding gonadotropin releasing hormone receptor (GnRHR).^68^ ELAVL1 has been shown to stabilizes the *Gnrhr* mRNA in the LβT2 gonadotrope-like cell line, to promote translation.^69^ Conversely, we have reported that MSI1 and MSI2 repress translation of the *Gnrhr* mRNA *in vivo*.^14^

Several possibilities can be considered to potentially reconcile the cooperative action of ELAVL family members to promote MSI-dependent mRNA translation reported here and the reported functional antagonism reported in other studies. MSI can function to either repress translation, or to promote translation of target mRNAs. The mechanism underlying the switch in functional activity is unknown, although we have identified a role for regulation by phosphosphorylation of conserved MSI C-terminal serine residues.^70,71^ Consequently, the ability of MSI1 to oppose ELAVL family members may be abrogated by MSI1 phosphorylation on shared target mRNAs in a context-dependent manner. Alternatively, it is possible that these distinct functions occur when ELAVL1 or ELAVL4 associate with MSI1 independently within a target mRNA 3’ UTR versus ELAVL1 or ELAVL4 recruitment to a MSI1 regulatory complex independently of RNA targeting. Interestingly, we detected ELAVL1 and ELAVL4 interaction with MSI1 in immature *Xenopus* oocytes, as well as in the progesterone-stimulated oocytes where MSI undergoes regulatory phosphorylation, indicating that MSI phosphorylation does not control co-association with ELAVL1 or ELAVL4.^20^

Further studies will be required to determine what features of the target mRNAs distinguish between cooperative interactions and antagonistic activity between MSI and ELAVL family members. It also will prove informative to determine if ELAVL4-MSI1 associations ocurr in mammalian tissues and whether ELAVL1 is required for regulation of other identified MSI-dependent pituitary mRNAs.^12,14–16,18^ Given the involvement of both MSI and ELAVL1 in cancer progression^72–74^, the precedent of ELAVL1-MSI2 complexes controlling miR-7 biogenesis^58^ and shared mRNA targets of both ELAVL proteins and MSI^14,34,35,44,68,69^, our results position ELAVL-MSI protein interactions as novel therapeutic targets to modulate mRNA translational control of proteostasis in disease states.

In conclusion, we report a novel dependency of progesterone-stimulated maturation on ELAVL4 and further show that ELAVL4 interacts with MSI1 to promote MSI-dependent mRNA translational activation. This interaction appears to be conserved with the related ELAVL1 protein. Our study suggests a novel combinatorial regulation of target mRNA translation through an mRNP composed in part of MSI1 and ELAVL1 or ELAVL4 proteins. Future experiments designed to disrupt MSI and ELAVL protein interactions will reveal the extent to which these proteins co-regulate target mRNA translation in physiological and pathological contexts.

## ACKNOWLEDGEMENTS

We thank Dr. Duah Alkam for proteomic data processing and dataset upload to MassIVE. This work was supported by the National Institutes of Health RO1DK139476 (to A.M.M., G.V.C., and M.C.M.), a Barton Pilot Award (A.M.M., and M.C.M.), a UAMS TRI Data Scholar award (M.C.M., UL1 TR003107, KL2 TR003108 and TL1 TR003109).

