## Supplementary Tables for "ELAVL1 and ELAVL4 are required for Musashi-dependent translational activation"

### Supplementary Data Table Legends:

#### Supplementary Table 1

An excel file listing all proteins identified in MSI1/2 immunoprecipitation vs control IgG immunoprecipitations from mouse whole pituitaries. The data includes 3 independent replicates for each immunoprecipitation condition.

#### Supplementary Table 2

An excel file showing only proteins from Supplemental Table 1 which showed enrichment in MSI1/2 over control IgG immunoprecipitations with a FDR p-value <0.055 and a positive log fold change > 2.

| Protein Annotation | | | | | | | | | | Log2 Cyclic Loess Normalized Intensities | | | | | | Musashi_vs_IgG | | | | | | | | | |
|---|---|---|---|---|---|---|---|---|---|---|---|---|---|---|---|---|---|---|---|---|---|---|---|---|---|
| id | rotein Nam | ession Num | lecular Wei | ein Group S | ied Peptide | Exclusivity | UniProt ID | Gene name | Description | E1_MSI12 | E1_IgG | E2_MSI12 | E2_IgG | E3_MSI12 | E3_IgG | logFC | CI.L | CI.R | AveExpr | t | B | P.Value | adj.P.Val | sig.PVal | sig.FDR |
| 1223 | sp\|Q9JKC8 | AP3M1_MC | 47 kDa | 0.997625 | 2 | 100% | Q9JKC8 | Ap3m1 | AP-3 comp | 20.76077 | 27.76044 | 21.21283 | 27.62662 | 20.90359 | 27.16517 | -6.55835 | -7.27387 | -5.84282 | 24.23824 | -21.8157 | 7.658529 | 1.55E-07 | 0.000211 | -1 | -1 |
| 27 | sp\|Q9Z1G4 | VPP1_MOU | 96 kDa | 0.999864 | 25 | 92% | Q9Z1G4 | Atp6v0a1 | V-type prot | 24.21201 | 30.41103 | 24.84352 | 30.34598 | 24.00178 | 30.14157 | -5.94709 | -6.70844 | -5.18574 | 27.32598 | -18.5917 | 6.892845 | 4.53E-07 | 0.000237 | -1 | -1 |
| 146 | sp\|P51863 | VA0D1_MC | 40 kDa | 0.999576 | 12 | 100% | P51863 | Atp6v0d1 | V-type prot | 23.90194 | 29.86094 | 24.62866 | 29.91128 | 24.0949 | 29.59593 | -5.58089 | -6.31142 | -4.85036 | 26.99894 | -18.183 | 6.779771 | 5.25E-07 | 0.000237 | -1 | -1 |
| 815 | sp\|P97873 | LOXL1_MO | 67 kDa | 0.999425 | 3 | 100% | P97873 | Loxl1 | Lysyl oxida | 19.32217 | 24.25602 | 18.69365 | 23.90748 | 19.21276 | 23.77751 | -4.90414 | -5.63738 | -4.1709 | 21.52826 | -15.9191 | 6.072755 | 1.27E-06 | 0.000425 | -1 | -1 |
| 150 | sp\|Q6IRU5 | CLCB_MOU | 25 kDa | 0.999464 | 12 | 92% | Q6IRU5 | Cltb | Clathrin lig | 28.38085 | 23.80329 | 28.405 | 24.14557 | 28.20823 | 23.45666 | 4.529515 | 3.830334 | 5.228696 | 26.0666 | 15.41923 | 5.895842 | 1.57E-06 | 0.000425 | 1 | 1 |
| 965 | sp\|Q9ET30 | TM9S3_MC | 68 kDa | 0.997681 | 3 | 100% | Q9ET30 | Tm9sf3 | Transmemb | 21.73077 | 25.95239 | 21.51526 | 25.49191 | 21.92613 | 25.78442 | -4.01885 | -4.69071 | -3.34699 | 23.73348 | -14.2371 | 5.442526 | 2.65E-06 | 0.000427 | -1 | -1 |
| 1269 | sp\|Q8CHP0 | ZC3H3_MO | 103 kDa | 0.996205 | 2 | 100% | Q8CHP0 | Zc3h3 | Zinc finger | 17.30309 | 21.83121 | 16.87701 | 21.70069 | 16.17289 | 21.75948 | -4.97947 | -5.82331 | -4.13562 | 19.27406 | -14.0552 | 5.341256 | 2.98E-06 | 0.000427 | -1 | -1 |
| 346 | sp\|P70662 | LDB1_MOU | 47 kDa | 0.999359 | 7 | 86% | P70662 | Ldb1 | LIM domain | 26.67237 | 21.65568 | 26.89319 | 21.90304 | 26.29582 | 22.43223 | 4.62348 | 3.83559 | 5.41137 | 24.30872 | 13.96704 | 5.331501 | 3.01E-06 | 0.000427 | 1 | 1 |
| 778 | sp\|P01811 | HVM41_M | 13 kDa | 0.995278 | 4 | 25% | P01811 | NA | Ig heavy ch | 21.20257 | 14.84614 | 21.92793 | 15.22002 | 20.71727 | 15.79102 | 5.996865 | 4.988373 | 7.005358 | 18.28416 | 14.19065 | 5.325797 | 3.03E-06 | 0.000427 | 1 | 1 |
| 25 | sp\|O35245 | PKD2_MOU | 109 kDa | 0.999878 | 26 | 100% | O35245 | Pkd2 | Polycystin- | 26.92774 | 23.18959 | 26.6901 | 22.84655 | 26.98119 | 22.60046 | 3.98748 | 3.303276 | 4.671684 | 24.8726 | 13.87119 | 5.291385 | 3.15E-06 | 0.000427 | 1 | 1 |
| 1015.2 | tr\|E9Q035 | E9Q035_M | 108 kDa | 0.99986 | 2 | 100% | E9Q035 | Gm20425 | Predicted g | 24.19942 | 18.47307 | 24.71601 | 18.60751 | 23.98793 | 19.52478 | 5.432666 | 4.499976 | 6.365356 | 21.58479 | 13.90037 | 5.207377 | 3.46E-06 | 0.000427 | 1 | 1 |
| 821 | sp\|P80315 | TCPD_MOU | 58 kDa | 0.999397 | 3 | 67% | P80315 | Cct4 | T-complex | 22.14219 | 26.46881 | 22.63716 | 25.86335 | 22.41899 | 26.38492 | -3.83958 | -4.56915 | -3.11001 | 24.31923 | -12.5262 | 4.68612 | 6.15E-06 | 0.000695 | -1 | -1 |
| 89 | sp\|P60879 | SNP25_MO | 23 kDa | 0.999799 | 16 | 94% | P60879 | Snap25 | Synaptosor | 28.13807 | 22.63727 | 27.60588 | 23.42141 | 27.93553 | 23.73367 | 4.629044 | 3.744928 | 5.513159 | 25.57864 | 12.48178 | 4.611365 | 6.68E-06 | 0.000697 | 1 | 1 |
| 131 | sp\|Q61474 | MSI1H_MO | 39 kDa | 0.999636 | 13 | 92% | Q61474 | Msi1 | RNA-bindir | 28.00456 | 24.28889 | 27.83918 | 23.26068 | 27.77549 | 23.72374 | 4.115307 | 3.306103 | 4.924512 | 25.81542 | 12.10503 | 4.477346 | 7.7E-06 | 0.000746 | 1 | 1 |
| 222 | sp\|O09044 | SNP23_MO | 23 kDa | 0.999882 | 9 | 89% | O09044 | Snap23 | Synaptosor | 24.79802 | 21.10309 | 24.14536 | 20.64175 | 24.43624 | 21.11742 | 3.505791 | 2.767889 | 4.243692 | 22.70698 | 11.30807 | 4.061053 | 1.2E-05 | 0.000976 | 1 | 1 |
| 817 | tr\|A0A0B4. | A0A0B4J1N | 11 kDa | 0.999419 | 3 | 33% | A0A0B4J1N | Ighv1-77 | Immunoglo | 19.53749 | 16.75791 | 19.94923 | 16.80292 | 19.96815 | 16.76371 | 3.043447 | 2.402736 | 3.684159 | 18.29657 | 11.30587 | 4.059855 | 1.2E-05 | 0.000976 | 1 | 1 |
| 1184 | sp\|Q8K386 | RAB15_MO | 24 kDa | 0.998107 | 2 | 50% | Q8K386 | Rab15 | Ras-related | 18.95745 | 15.91514 | 19.35923 | 16.2916 | 19.33877 | 16.0425 | 3.135401 | 2.470756 | 3.800046 | 17.65078 | 11.22805 | 4.017113 | 1.25E-05 | 0.000976 | 1 | 1 |
| 1 | sp\|Q68FD5 | CLH1_MOU | 192 kDa | 0.999927 | 86 | 100% | Q68FD5 | Cltc | Clathrin he | 33.45827 | 28.44516 | 32.44402 | 27.97304 | 33.37165 | 28.92852 | 4.642404 | 3.66433 | 5.620478 | 30.77011 | 11.32718 | 3.990771 | 1.3E-05 | 0.000976 | 1 | 1 |
| 35 | sp\|Q6NZC7 | S23IP_MOU | 111 kDa | 0.999696 | 23 | 100% | Q6NZC7 | Sec23ip | SEC23-inte | 28.70469 | 24.29238 | 27.76828 | 24.26505 | 28.49033 | 24.75437 | 3.883829 | 3.049447 | 4.718211 | 26.37918 | 11.07943 | 3.933021 | 1.37E-05 | 0.000976 | 1 | 1 |
| 465 | sp\|O08585 | CLCA_MOU | 26 kDa | 0.998487 | 6 | 83% | O08585 | Clta | Clathrin lig | 26.99857 | 21.91196 | 26.85232 | 23.2787 | 26.83168 | 22.39684 | 4.365024 | 3.419942 | 5.310106 | 24.71168 | 11.02218 | 3.823632 | 1.54E-05 | 0.001047 | 1 | 1 |
| 673 | sp\|P53395 | ODB2_MOU | 53 kDa | 0.999094 | 4 | 100% | P53395 | Dbt | Lipoamide | 17.56244 | 21.5222 | 18.35403 | 21.60027 | 17.99298 | 21.1373 | -3.45011 | -4.21641 | -2.6838 | 19.69487 | -10.716 | 3.726752 | 1.69E-05 | 0.001058 | -1 | -1 |
| 305 | sp\|Q60749 | KHDR1_MC | 48 kDa | 0.999885 | 7 | 86% | Q60749 | Khdrbs1 | KH domain | 26.58346 | 21.36758 | 25.47485 | 20.85161 | 27.02601 | 20.11857 | 5.582188 | 4.353376 | 6.811 | 23.57035 | 10.84101 | 3.721701 | 1.72E-05 | 0.001058 | 1 | 1 |
| 877 | sp\|P46412 | GPX3_MOU | 25 kDa | 0.998944 | 3 | 100% | P46412 | Gpx3 | Glutathion | 18.0483 | 24.1013 | 19.86396 | 23.92534 | 18.39102 | 24.25832 | -5.32723 | -6.52138 | -4.13308 | 21.43137 | -10.6462 | 3.609801 | 1.93E-05 | 0.001124 | -1 | -1 |
| 273 | sp\|O35737 | HNRH1_M | 49 kDa | 0.999513 | 8 | 38% | O35737 | Hnrnph1 | Heterogene | 23.74254 | 26.19736 | 23.47067 | 26.27192 | 23.46819 | 26.46083 | -2.74957 | -3.37576 | -2.12338 | 24.93525 | -10.451 | 3.570037 | 1.99E-05 | 0.001124 | -1 | -1 |
| 1061 | sp\|O09108 | LSHB_MOU | 15 kDa | 0.999248 | 2 | 100% | O09108 | Lhb | Lutropin su | 19.02823 | NA | 18.45906 | 13.25611 | 19.67979 | 13.71208 | 5.571598 | 4.430488 | 6.712707 | 16.82705 | 12.10273 | 3.290413 | 2.8E-05 | 0.001518 | 1 | 1 |
| 329 | sp\|P06879 | PRL_MOUS | 25 kDa | 0.999529 | 7 | 100% | P06879 | Prl | Prolactin | 30.24519 | 26.1948 | 30.41172 | 26.66332 | 29.35326 | 26.185 | 3.655683 | 2.766844 | 4.544523 | 28.17555 | 9.804813 | 3.125505 | 3.15E-05 | 0.001631 | 1 | 1 |
| 593 | tr\|A0A075 | A0A075B5 | 13 kDa | 0.999779 | 4 | 50% | A0A075B5 | Ighv5-15 | Immunoglo | 20.77639 | 23.62668 | 20.93077 | 23.51604 | 20.3823 | 23.2847 | -2.77932 | -3.46261 | -2.09603 | 22.08615 | -9.68131 | 3.08767 | 3.25E-05 | 0.001631 | -1 | -1 |
| 18 | sp\|P63017 | HSP7C_MO | 71 kDa | 0.999881 | 31 | 65% | P63017 | Hspa8 | Heat shock | 27.85382 | 24.30508 | 27.7775 | 24.49842 | 27.83724 | 25.23323 | 3.143945 | 2.361162 | 3.926729 | 26.25088 | 9.559479 | 3.007402 | 3.52E-05 | 0.001704 | 1 | 1 |
| 266 | sp\|Q8BMF4 | ODP2_MOU | 68 kDa | 0.999605 | 8 | 100% | Q8BMF4 | Dlat | Dihydrolip | 22.12227 | 25.28007 | 22.79759 | 25.53783 | 22.747 | 25.6216 | -2.92421 | -3.65684 | -2.19158 | 24.01772 | -9.50006 | 2.967848 | 3.66E-05 | 0.001704 | -1 | -1 |
| 611 | sp\|Q9CYZ8 | SSBP2_MO | 38 kDa | 0.999605 | 4 | 75% | Q9CYZ8 | Ssbp2 | Single-stra | 23.22381 | 17.48517 | 23.58941 | 19.2692 | 22.92989 | 18.70953 | 4.759738 | 3.573528 | 5.945949 | 20.86784 | 9.575731 | 2.949857 | 3.77E-05 | 0.001704 | 1 | 1 |
| 415 | sp\|P42669 | PURA_MOU | 35 kDa | 0.999562 | 6 | 100% | P42669 | Pura | Transcripti | 27.2554 | 23.15779 | 26.17552 | 23.10798 | 27.42142 | 23.38845 | 3.732709 | 2.792645 | 4.672772 | 25.08443 | 9.475831 | 2.884084 | 4.03E-05 | 0.001752 | 1 | 1 |
| 45 | sp\|P28740 | KIF2A_MO | 80 kDa | 0.999855 | 21 | 100% | P28740 | Kif2a | Kinesin-like | 27.45602 | 23.07958 | 26.23326 | 23.07752 | 27.59822 | 23.29447 | 3.94531 | 2.947576 | 4.943043 | 25.12318 | 9.436634 | 2.85807 | 4.13E-05 | 0.001752 | 1 | 1 |
| 401 | sp\|Q9CX30 | YIF1B_MOU | 34 kDa | 0.999669 | 6 | 100% | Q9CX30 | Yif1b | Protein YIF | 25.40531 | 22.23832 | 25.71674 | 23.02212 | 25.38173 | 22.52984 | 2.904495 | 2.157756 | 3.651234 | 24.04901 | 9.257689 | 2.803678 | 4.32E-05 | 0.001774 | 1 | 1 |
| 185 | sp\|Q7TSH6 | SCAF4_MO | 129 kDa | 0.999718 | 10 | 70% | Q7TSH6 | Scaf4 | SR-related | 26.36227 | 22.25709 | 25.69055 | 23.08979 | 26.06716 | 21.89685 | 3.625415 | 2.680733 | 4.570096 | 24.22728 | 9.158468 | 2.670024 | 4.99E-05 | 0.001989 | 1 | 1 |
| 336 | sp\|Q8VCM7 | FIBG_MOU | 49 kDa | 0.999459 | 7 | 100% | Q8VCM7 | Fgg | Fibrinogen | 24.21079 | 21.07079 | 23.39936 | 21.09125 | 23.81612 | 20.64447 | 2.873253 | 2.09987 | 3.646637 | 22.37213 | 8.842596 | 2.511556 | 5.77E-05 | 0.002206 | 1 | 1 |
| 869 | sp\|Q61543 | GSLG1_MO | 134 kDa | 0.998993 | 3 | 100% | Q61543 | Glg1 | Golgi appa | 22.33651 | 19.98973 | 22.35442 | 19.45564 | 22.00334 | 19.49077 | 2.58604 | 1.888317 | 3.283763 | 20.9384 | 8.821714 | 2.496481 | 5.86E-05 | 0.002206 | 1 | 1 |
| 960 | sp\|Q8CC88 | VWA8_MO | 213 kDa | 0.997749 | 3 | 100% | Q8CC88 | Vwa8 | von Willeb | 19.3021 | 21.82637 | 18.8084 | 22.5329 | 19.51113 | 22.76163 | -3.16642 | -4.03557 | -2.29728 | 20.79042 | -8.68502 | 2.359742 | 6.75E-05 | 0.002473 | -1 | -1 |
| 725 | tr\|A0A075 | A0A075B5 | 12 kDa | 0.998596 | 4 | 100% | A0A075B5 | Igkv12-38 | Immunoglo | 22.06952 | 25.55634 | 21.10743 | 25.18654 | 22.49077 | 25.58051 | -3.55189 | -4.53411 | -2.56966 | 23.66519 | -8.62976 | 2.295263 | 7.22E-05 | 0.002578 | -1 | -1 |
| 32 | sp\|Q6PDG5 | SMRC2_MC | 133 kDa | 0.999852 | 24 | 67% | Q6PDG5 | Smarcc2 | SWI/SNF co | 25.9951 | 22.71468 | 25.24442 | 22.74158 | 26.21342 | 23.12373 | 2.957648 | 2.127008 | 3.788287 | 24.33882 | 8.47531 | 2.239788 | 7.54E-05 | 0.002622 | 1 | 1 |
| 317 | sp\|Q8BP99 | CA216_MO | 25 kDa | 0.999646 | 7 | 100% | Q8BP99 | NA | UPF0500 p | 22.22913 | 17.70089 | 22.15634 | 19.02669 | 22.08825 | 19.02444 | 3.573903 | 2.569788 | 4.578018 | 20.37096 | 8.493954 | 2.195138 | 7.97E-05 | 0.002702 | 1 | 1 |
| 10 | sp\|Q9EPU0 | RENT1_MO | 124 kDa | 0.999794 | 40 | 100% | Q9EPU0 | Upf1 | Regulator c | 26.88991 | 24.32191 | 26.69588 | 24.45162 | 26.57692 | 24.69033 | 2.232949 | 1.592188 | 2.873711 | 25.60443 | 8.294372 | 2.10313 | 8.62E-05 | 0.002851 | 1 | 1 |
| 760 | sp\|Q8R3N6 | THOC1_MC | 75 kDa | 0.997356 | 4 | 100% | Q8R3N6 | Thoc1 | THO compl | 21.25154 | 18.81515 | 20.96518 | 18.36662 | 21.37676 | 19.01436 | 2.465782 | 1.747415 | 3.184149 | 19.96494 | 8.169757 | 2.006487 | 9.47E-05 | 0.003058 | 1 | 1 |
| 171 | sp\|Q99KN9 | EPN4_MOU | 69 kDa | 0.999326 | 11 | 100% | Q99KN9 | Clint1 | Clathrin int | 24.55638 | 21.62096 | 24.17584 | 21.75137 | 23.7416 | 21.67437 | 2.47571 | 1.745107 | 3.206314 | 22.92009 | 8.065269 | 1.924321 | 0.000103 | 0.003176 | 1 | 1 |
| 614 | sp\|O70439 | STX7_MOU | 30 kDa | 0.999563 | 4 | 100% | O70439 | Stx7 | Syntaxin-7 | 24.47173 | 21.79934 | 24.09237 | 22.1601 | 24.28612 | 21.40442 | 2.495454 | 1.75852 | 3.232387 | 23.03568 | 8.059757 | 1.919958 | 0.000103 | 0.003176 | 1 | 1 |
| 700 | sp\|P01672 | KV3AK_MO | 12 kDa | 0.998891 | 4 | 25% | P01672 | NA | Ig kappa ch | 17.85217 | 25.33286 | 16.11599 | 23.82975 | 19.55036 | 25.45224 | -7.03211 | -9.17916 | -4.88507 | 21.35556 | -7.81619 | 1.670113 | 0.000133 | 0.003972 | -1 | -1 |
| 672 | sp\|P62814 | VATB2_MO | 57 kDa | 0.999109 | 4 | 100% | P62814 | Atp6v1b2 | V-type prot | 21.42942 | 23.58816 | 21.8876 | 23.90719 | 21.62955 | 23.90907 | -2.15262 | -2.81644 | -1.4888 | 22.72516 | -7.71826 | 1.643795 | 0.000135 | 0.003972 | -1 | -1 |
| 842 | sp\|P35278 | RAB5C_MO | 23 kDa | 0.999231 | 3 | 33% | P35278 | Rab5c | Ras-related | 23.52915 | 25.74374 | 23.58007 | 25.43355 | 23.42921 | 25.40566 | -2.01484 | -2.63907 | -1.39061 | 24.52023 | -7.68242 | 1.614133 | 0.000139 | 0.004 | -1 | -1 |
| 1006 | sp\|Q8R1F6 | HID1_MOU | 89 kDa | 0.995228 | 3 | 100% | Q8R1F6 | Hid1 | Protein HID | 21.51472 | 18.28828 | 21.00575 | 17.14254 | 22.13313 | 16.37584 | 4.282311 | 2.959029 | 5.605593 | 19.41004 | 7.722831 | 1.594312 | 0.000143 | 0.004042 | 1 | 1 |
| 1020 | sp\|Q91VK4 | ITM2C_MO | 30 kDa | 0.99973 | 2 | 100% | Q91VK4 | Itm2c | Integral me | 22.43251 | 24.79777 | 22.86348 | 25.01832 | 22.11566 | 24.62815 | -2.3442 | -3.08138 | -1.60702 | 23.64265 | -7.56874 | 1.519172 | 0.000152 | 0.004094 | -1 | -1 |
| 1021 | tr\|A0A075 | A0A075B5 | 13 kDa | 0.999721 | 2 | 50% | A0A075B5 | Ighv8-11 | Immunoglo | 22.43251 | 24.79777 | 22.86348 | 25.01832 | 22.11566 | 24.62815 | -2.3442 | -3.08138 | -1.60702 | 23.64265 | -7.56874 | 1.519172 | 0.000152 | 0.004094 | -1 | -1 |
| 487 | sp\|P29699 | FETUA_MO | 37 kDa | 0.99975 | 5 | 100% | P29699 | Ahsg | Alpha-2-HS | 25.11261 | 22.42983 | 25.37017 | 22.68068 | 24.46618 | 22.30354 | 2.511637 | 1.720119 | 3.303154 | 23.72717 | 7.552614 | 1.505597 | 0.000154 | 0.004094 | 1 | 1 |
| 16 | sp\|P14873 | MAP1B_M | 270 kDa | 0.999829 | 33 | 100% | P14873 | Map1b | Microtubu | 23.45623 | 25.67341 | 23.8141 | 25.9599 | 23.4985 | 25.48232 | -2.1156 | -2.78703 | -1.44416 | 24.64741 | -7.49944 | 1.460631 | 0.000161 | 0.004193 | -1 | -1 |
| 149 | sp\|P46460 | NSF_MOUS | 83 kDa | 0.999474 | 12 | 100% | P46460 | Nsf | Vesicle-fusi | 23.79325 | 25.98677 | 24.10546 | 25.83201 | 24.08428 | 26.0159 | -1.95057 | -2.57399 | -1.32714 | 24.96961 | -7.44693 | 1.415927 | 0.000168 | 0.004295 | -1 | -1 |
| 1167 | sp\|P11438 | LAMP1_M | 44 kDa | 0.998266 | 2 | 100% | P11438 | Lamp1 | Lysosome-a | 19.02603 | 22.54742 | 18.41984 | 21.2358 | 19.0289 | 21.83544 | -3.04796 | -4.03173 | -2.0642 | 20.3489 | -7.39381 | 1.320057 | 0.000186 | 0.004586 | -1 | -1 |
| 1034 | tr\|A0A075 | A0A075B6 | 10 kDa | 0.999481 | 2 | 100% | A0A075B6 | Igkv4-53 | Immunoglo | 22.30582 | 18.99434 | 21.35155 | 19.49426 | 22.41964 | 18.67668 | 2.970574 | 2.010291 | 3.930856 | 20.54038 | 7.382304 | 1.310256 | 0.000188 | 0.004586 | 1 | 1 |
| 202 | sp\|Q61235 | SNTB2_MO | 56 kDa | 0.99945 | 10 | 90% | Q61235 | Sntb2 | Beta-2-synt | 23.90047 | 20.00991 | 23.21943 | 18.6166 | 23.04242 | 20.27871 | 3.752367 | 2.538014 | 4.966719 | 21.51126 | 7.374136 | 1.303293 | 0.000189 | 0.004586 | 1 | 1 |
| 240 | sp\|Q9WTY1 | PDCD7_M | 54 kDa | 0.999289 | 9 | 100% | Q9WTY1 | Pdcd7 | Programme | 24.95165 | 20.8272 | 23.6166 | 21.2312 | 25.04715 | 21.50362 | 3.351132 | 2.257518 | 4.444745 | 22.8629 | 7.312712 | 1.250699 | 0.000199 | 0.004622 | 1 | 1 |
| 932 | sp\|Q6P5F9 | XPO1_MOU | 123 kDa | 0.998194 | 3 | 100% | Q6P5F9 | Xpo1 | Exportin-1 | 24.83588 | 26.61359 | 24.54679 | 26.46385 | 24.7107 | 26.59012 | -1.85806 | -2.47004 | -1.24609 | 25.62682 | -7.22649 | 1.225039 | 0.000202 | 0.004622 | -1 | -1 |
| 113 | tr\|Q3UJB0 | Q3UJB0_M | 98 kDa | 0.999862 | 14 | 100% | Q3UJB0 | Sf3b2 | Splicing fac | 24.56331 | 21.89792 | 23.82033 | 22.09233 | 24.76927 | 21.53842 | 2.541412 | 1.698843 | 3.383981 | 23.1136 | 7.184315 | 1.174536 | 0.000212 | 0.004622 | 1 | 1 |
| 895 | tr\|A0A075 | A0A075B5 | 13 kDa | 0.998746 | 3 | 33% | A0A075B5 | Ighv1-54 | Immunoglo | 19.32216 | 22.55592 | 20.29939 | 22.08272 | 19.97117 | 22.64287 | -2.56293 | -3.41249 | -1.71337 | 21.1457 | -7.19014 | 1.167838 | 0.000214 | 0.004622 | -1 | -1 |
| 867.1 | tr\|A0A075 | A0A075B5 | 13 kDa | 0.999017 | 3 | 100% | A0A075B5 | Igkv6-29 | Immunoglo | 20.24114 | 23.03438 | 20.98652 | 23.70762 | 19.87035 | 22.95128 | -2.86509 | -3.81424 | -1.91594 | 21.79855 | -7.2037 | 1.156352 | 0.000218 | 0.004622 | -1 | -1 |
| 967 | sp\|Q99LX0 | PARK7_MO | 20 kDa | 0.997612 | 3 | 100% | Q99LX0 | Park7 | Protein/nu | 19.73811 | 17.72553 | 20.29779 | 17.60711 | 19.48512 | 17.55125 | 2.212375 | 1.473917 | 2.950833 | 18.73415 | 7.130727 | 1.140447 | 0.000219 | 0.004622 | 1 | 1 |
| 128 | sp\|Q8CJG0 | AGO2_MOU | 97 kDa | 0.999715 | 13 | 85% | Q8CJG0 | Ago2 | Protein arg | 23.84731 | 21.8418 | 23.43332 | 21.78657 | 23.93438 | 21.52956 | 2.019025 | 1.344763 | 2.693288 | 22.72882 | 7.127113 | 1.137234 | 0.000219 | 0.004622 | 1 | 1 |
| 224 | sp\|Q3UH66 | WNK2_MO | 228 kDa | 0.999818 | 9 | 89% | Q3UH66 | Wnk2 | Serine/thre | 21.59838 | 18.0727 | 21.81789 | 19.49968 | 21.87457 | 18.92093 | 2.932511 | 1.959063 | 3.90596 | 20.29736 | 7.189146 | 1.143654 | 0.000221 | 0.004622 | 1 | 1 |
| 1135 | sp\|Q61771 | KIF3B_MOU | 85 kDa | 0.998572 | 2 | 100% | Q61771 | Kif3b | Kinesin-like | 20.20314 | 22.59185 | 19.91895 | 21.91601 | 19.55877 | 22.09827 | -2.30842 | -3.08057 | -1.53628 | 21.04783 | -7.11573 | 1.127105 | 0.000222 | 0.004622 | -1 | -1 |
| 51 | sp\|Q8BZ98 | DYN3_MOU | 97 kDa | 0.999347 | 20 | 65% | Q8BZ98 | Dnm3 | Dynamin-3 | 25.00528 | 21.075 | 23.35635 | 19.66307 | 25.03548 | 20.22352 | 4.145173 | 2.745449 | 5.544897 | 22.39312 | 7.067259 | 1.03641 | 0.000245 | 0.004931 | 1 | 1 |
| 257 | sp\|Q8VDM4 | PSMD2_M | 100 kDa | 0.999764 | 8 | 100% | Q8VDM4 | Psmd2 | 26S protea | 22.18327 | 24.19029 | 22.36904 | 24.03655 | 22.13198 | 23.91414 | -1.81889 | -2.43739 | -1.2004 | 23.13754 | -6.99963 | 1.022977 | 0.000245 | 0.004931 | -1 | -1 |
| 47 | sp\|Q9QX47 | SON_MOUS | 266 kDa | 0.999742 | 20 | 100% | Q9QX47 | Son | Protein SON | 23.98495 | 21.62267 | 23.9458 | 21.34901 | 24.5157 | 22.22077 | 2.417995 | 1.5945 | 3.24149 | 22.93982 | 6.989019 | 1.012516 | 0.000247 | 0.004931 | 1 | 1 |
| 1122 | tr\|B2RXQ2 | B2RXQ2_M | 143 kDa | 0.998694 | 2 | 100% | B2RXQ2 | Ppfia1 | Ppfia1 prot | 20.92404 | 19.22578 | 20.93676 | 18.99896 | 20.78955 | 18.725 | 1.900201 | 1.251107 | 2.549294 | 19.93335 | 6.967754 | 0.994115 | 0.000252 | 0.004945 | 1 | 1 |
| 1201 | sp\|Q9D883 | U2AF1_MO | 28 kDa | 0.997941 | 2 | 100% | Q9D883 | U2af1 | Splicing fac | 13.95881 | 20.42201 | 13.85657 | 18.76473 | 15.7423 | 18.70523 | -4.7781 | -6.41853 | -3.13767 | 16.90828 | -6.95101 | 0.932566 | 0.00027 | 0.005213 | -1 | -1 |
| 133 | sp\|Q8C3F2 | F120C_MO | 120 kDa | 0.999553 | 13 | 85% | Q8C3F2 | Fam120c | Constitutiv | 24.45541 | 21.95124 | 24.25257 | 21.71945 | 23.83938 | 22.35075 | 2.175304 | 1.422157 | 2.928451 | 23.0948 | 6.874494 | 0.909007 | 0.000273 | 0.005213 | 1 | 1 |
| 1200 | sp\|O35226 | PSMD4_M | 41 kDa | 0.997946 | 2 | 100% | O35226 | Psmd4 | 26S protea | 19.63834 | 18.04042 | 19.6948 | 17.72898 | 19.696 | 17.97699 | 1.760916 | 1.148333 | 2.373499 | 18.79592 | 6.841865 | 0.87899 | 0.000281 | 0.00529 | 1 | 1 |
| 1001 | sp\|Q9DC51 | GNAI3_MO | 41 kDa | 0.995483 | 3 | 33% | Q9DC51 | Gnai3 | Guanine nu | 20.16763 | 22.66421 | 21.0293 | 22.78926 | 20.4766 | 22.65797 | -2.14597 | -2.8958 | -1.39614 | 21.63083 | -6.81184 | 0.851257 | 0.000288 | 0.005357 | -1 | -1 |
| 660 | sp\|P56395 | CYB5_MOU | 15 kDa | 0.999229 | 4 | 100% | P56395 | Cyb5a | Cytochrom | 21.33571 | 25.39062 | 22.28798 | 24.08911 | 21.39577 | 24.7135 | -3.05792 | -4.1267 | -1.98914 | 23.20212 | -6.82791 | 0.820907 | 0.000301 | 0.005507 | -1 | -1 |
| 488 | tr\|A0A0A6 | A0A0A6YYE | 13 kDa | 0.999748 | 5 | 80% | A0A0A6YYE | Igkv4-57 | Immunoglo | 22.71461 | 25.18804 | 23.24824 | 25.04631 | 22.70693 | 24.64219 | -2.06892 | -2.80411 | -1.33374 | 23.92439 | -6.69807 | 0.745204 | 0.000319 | 0.00576 | -1 | -1 |
| 158 | sp\|P27659 | RL3_MOUS | 46 kDa | 0.999783 | 11 | 100% | P27659 | Rpl3 | 60S riboso | 25.38364 | 23.54433 | 25.15199 | 23.32146 | 25.12776 | 23.56829 | 1.7431 | 1.121216 | 2.364985 | 24.34958 | 6.671347 | 0.720072 | 0.000327 | 0.00576 | 1 | 1 |
| 505 | sp\|Q9CQF3 | CPSF5_MO | 26 kDa | 0.999523 | 5 | 100% | Q9CQF3 | Nudt21 | Cleavage a | 23.55598 | 21.18985 | 22.98271 | 21.12811 | 23.0107 | 21.40734 | 1.94136 | 1.245526 | 2.637194 | 22.21245 | 6.640507 | 0.690958 | 0.000336 | 0.00576 | 1 | 1 |
| 209 | tr\|Q6PHQ9 | Q6PHQ9_M | 72 kDa | 0.999289 | 10 | 80% | Q6PHQ9 | Pabpc4 | Polyadenyl | 22.35413 | 19.69856 | 21.93965 | 19.30669 | 21.99438 | 20.32371 | 2.319733 | 1.48701 | 3.152457 | 20.93619 | 6.630695 | 0.680866 | 0.000339 | 0.00576 | 1 | 1 |
| 162 | sp\|P47911 | RL6_MOUS | 34 kDa | 0.999527 | 11 | 100% | P47911 | Rpl6 | 60S riboso | 25.64704 | 24.0894 | 25.82642 | 24.06906 | 25.67027 | 23.73458 | 1.750233 | 1.118812 | 2.381655 | 24.83946 | 6.597471 | 0.650136 | 0.000349 | 0.00576 | 1 | 1 |
| 72 | sp\|Q9D8E6 | RL4_MOUS | 47 kDa | 0.999669 | 17 | 100% | Q9D8E6 | Rpl4 | 60S riboso | 25.47186 | 23.96585 | 25.47296 | 23.57612 | 25.53497 | 23.76307 | 1.724916 | 1.102302 | 2.347531 | 24.63081 | 6.594014 | 0.646847 | 0.00035 | 0.00576 | 1 | 1 |
| 100.1 | sp\|P17879 | HS71B_MO | 70 kDa | 0.999699 | 15 | 73% | P17879 | Hspa1b | Heat shock | 24.99751 | 22.79349 | 24.59787 | 22.00965 | 24.82205 | 22.90713 | 2.23572 | 1.426672 | 3.044768 | 23.68795 | 6.577565 | 0.630384 | 0.000356 | 0.00576 | 1 | 1 |
| 100.2 | sp\|Q61696 | HS71A_MO | 70 kDa | 0.999699 | 15 | 73% | Q61696 | Hspa1a | Heat shock | 24.99751 | 22.79349 | 24.59787 | 22.00965 | 24.82205 | 22.90713 | 2.23572 | 1.426672 | 3.044768 | 23.68795 | 6.577565 | 0.630384 | 0.000356 | 0.00576 | 1 | 1 |
| 474 | sp\|P06330 | HVM51_M | 13 kDa | 0.997551 | 6 | 33% | P06330 | NA | Ig heavy ch | 22.48901 | 24.47297 | 22.66889 | 24.13055 | 22.60441 | 24.30527 | -1.71549 | -2.3366 | -1.09438 | 23.44518 | -6.57383 | 0.627613 | 0.000357 | 0.00576 | -1 | -1 |
| 534 | sp\|Q91YN9 | BAG2_MOU | 23 kDa | 0.999217 | 5 | 100% | Q91YN9 | Bag2 | BAG family | 22.41719 | 20.56972 | 22.06817 | 20.48345 | 22.55459 | 19.98094 | 2.001945 | 1.276328 | 2.727562 | 21.34568 | 6.566671 | 0.620783 | 0.000359 | 0.00576 | 1 | 1 |
| 390 | sp\|Q02105 | C1QC_MOU | 26 kDa | 0.999859 | 6 | 100% | Q02105 | C1qc | Compleme | 25.20597 | 27.0774 | 25.37386 | 27.64586 | 25.31825 | 26.98727 | -1.93749 | -2.64036 | -1.23461 | 26.2681 | -6.56089 | 0.615261 | 0.000361 | 0.00576 | -1 | -1 |
| 1207 | sp\|Q9CR67 | TMM33_M | 28 kDa | 0.997827 | 2 | 100% | Q9CR67 | Tmem33 | Transmemb | 19.33726 | 20.98385 | 18.95207 | 21.01609 | 18.36109 | 21.07694 | -2.14216 | -2.92678 | -1.35754 | 19.95455 | -6.49818 | 0.555091 | 0.000382 | 0.006028 | -1 | -1 |
| 603 | sp\|Q91V92 | ACLY_MOU | 120 kDa | 0.999696 | 4 | 100% | Q91V92 | Acly | ATP-citrate | 19.25311 | 21.09801 | 18.97263 | 21.2785 | 19.51354 | 20.91574 | -1.85099 | -2.53071 | -1.17127 | 20.17192 | -6.48145 | 0.538951 | 0.000388 | 0.006051 | -1 | -1 |
| 888 | sp\|Q9CYK2 | QPCT_MOU | 41 kDa | 0.998803 | 3 | 100% | Q9CYK2 | Qpct | Glutaminyl | 22.55832 | 20.3088 | 22.12631 | 20.06781 | 21.77316 | 20.14065 | 1.98018 | 1.250027 | 2.710333 | 21.16251 | 6.454927 | 0.513303 | 0.000398 | 0.006063 | 1 | 1 |
| 201 | sp\|P12970 | RL7A_MOU | 30 kDa | 0.999456 | 10 | 100% | P12970 | Rpl7a | 60S riboso | 24.97129 | 23.25296 | 25.23147 | 23.27719 | 24.75899 | 23.24403 | 1.729186 | 1.091552 | 2.36682 | 24.12266 | 6.454625 | 0.51301 | 0.000398 | 0.006063 | 1 | 1 |
| 325 | sp\|P46062 | SIPA1_MO | 112 kDa | 0.999569 | 7 | 100% | P46062 | Sipa1 | Signal-indu | 22.8695 | 19.37953 | 21.36005 | 19.55026 | 22.09384 | 19.09797 | 2.765208 | 1.743169 | 3.787247 | 20.72519 | 6.456705 | 0.473304 | 0.000418 | 0.006259 | 1 | 1 |
| 12 | sp\|Q8CGB6 | TNS2_MOU | 152 kDa | 0.999844 | 38 | 97% | Q8CGB6 | Tns2 | Tensin-2 | 27.48814 | 25.23302 | 27.2899 | 24.28074 | 27.30562 | 25.42792 | 2.380662 | 1.497997 | 3.263327 | 26.17089 | 6.429779 | 0.463807 | 0.00042 | 0.006259 | 1 | 1 |
| 620 | sp\|Q9CR57 | RL14_MOU | 24 kDa | 0.999496 | 4 | 100% | Q9CR57 | Rpl14 | 60S riboso | 25.19446 | 23.55456 | 25.51297 | 23.76991 | 25.40258 | 23.79248 | 1.664354 | 1.038076 | 2.290632 | 24.53783 | 6.325271 | 0.386636 | 0.000449 | 0.006612 | 1 | 1 |
| 313 | sp\|Q9CR62 | M2OM_MO | 34 kDa | 0.999669 | 7 | 100% | Q9CR62 | Slc25a11 | Mitochond | 23.14452 | 24.76875 | 23.08397 | 25.90765 | 22.56156 | 25.32734 | -2.40457 | -3.3112 | -1.49793 | 24.1323 | -6.32262 | 0.359554 | 0.000464 | 0.006761 | -1 | -1 |
| 863 | sp\|P60867 | RS20_MOU | 13 kDa | 0.99904 | 3 | 100% | P60867 | Rps20 | 40S riboso | 22.76709 | 20.86503 | 23.24579 | 21.0859 | 22.65141 | 21.19691 | 1.838817 | 1.141489 | 2.536146 | 21.96869 | 6.276274 | 0.338213 | 0.00047 | 0.006775 | 1 | 1 |
| 442 | sp\|P21981 | TGM2_MO | 77 kDa | 0.999192 | 6 | 100% | P21981 | Tgm2 | Protein-glu | 23.24223 | 24.71714 | 23.06424 | 24.9141 | 23.33857 | 24.83918 | -1.60846 | -2.22203 | -0.9949 | 24.01924 | -6.23955 | 0.301716 | 0.000486 | 0.006871 | -1 | -1 |
| 762 | sp\|Q8CJG1 | AGO1_MOU | 97 kDa | 0.997303 | 4 | 50% | Q8CJG1 | Ago1 | Protein arg | 25.07083 | 26.84661 | 25.39802 | 27.50542 | 25.40617 | 27.12042 | -1.86581 | -2.57759 | -1.15402 | 26.22458 | -6.23905 | 0.301218 | 0.000486 | 0.006871 | -1 | -1 |
| 121 | sp\|O70133 | DHX9_MOU | 149 kDa | 0.999414 | 14 | 100% | O70133 | Dhx9 | ATP-depen | 24.84958 | 22.61174 | 24.1873 | 22.58325 | 24.62655 | 22.89707 | 1.857126 | 1.147273 | 2.566979 | 23.62592 | 6.226926 | 0.289132 | 0.000492 | 0.006878 | 1 | 1 |
| 227 | sp\|Q9QZM0 | UBQL2_MC | 67 kDa | 0.999751 | 9 | 78% | Q9QZM0 | Ubqln2 | Ubiquilin-2 | 25.1567 | 22.82399 | 26.28683 | 22.33402 | 24.68113 | 22.3235 | 2.88105 | 1.784037 | 3.978064 | 23.93436 | 6.267431 | 0.289529 | 0.000498 | 0.006888 | 1 | 1 |
| 164 | sp\|Q920Q6 | MSI2H_MO | 37 kDa | 0.999498 | 11 | 91% | Q920Q6 | Msi2 | RNA-bindir | 30.20477 | 26.58534 | 29.58572 | 24.87125 | 30.10317 | 27.15205 | 3.761676 | 2.325269 | 5.198084 | 28.08372 | 6.249632 | 0.272013 | 0.000506 | 0.006933 | 1 | 1 |
| 137 | sp\|Q3UZ01 | RNPC3_MC | 58 kDa | 0.999248 | 13 | 100% | Q3UZ01 | Rnpc3 | RNA-bindir | 25.61458 | 20.83318 | 24.21064 | 22.35387 | 25.69598 | 21.84229 | 3.497288 | 2.152342 | 4.842235 | 23.42509 | 6.205503 | 0.22841 | 0.000527 | 0.007135 | 1 | 1 |
| 934 | sp\|Q3U9G9 | LBR_MOUS | 71 kDa | 0.998165 | 3 | 100% | Q3U9G9 | Lbr | Delta(14)-s | 19.24133 | 23.56489 | 20.08595 | 21.79438 | 19.18778 | 22.75413 | -3.19945 | -4.43145 | -1.96745 | 21.10474 | -6.19747 | 0.220444 | 0.000531 | 0.007135 | -1 | -1 |
| 1033.1 | sp\|O08583 | THOC4_MC | 27 kDa | 0.999499 | 2 | 100% | O08583 | Alyref | THO compl | 21.36412 | 24.28091 | 20.31773 | 25.11617 | 18.88521 | 23.73803 | -4.18935 | -5.81467 | -2.56403 | 22.2837 | -6.15116 | 0.174372 | 0.000555 | 0.007302 | -1 | -1 |
| 253 | sp\|O54774 | AP3D1_MC | 135 kDa | 0.997664 | 9 | 100% | O54774 | Ap3d1 | AP-3 comp | 18.61988 | 20.98259 | 18.38482 | 20.68194 | 18.8474 | 20.1498 | -1.98741 | -2.76301 | -1.21182 | 19.61107 | -6.09892 | 0.160342 | 0.000556 | 0.007302 | -1 | -1 |
| 180 | sp\|Q80U40 | RIMB2_MC | 118 kDa | 0.998865 | 11 | 100% | Q80U40 | Rimbp2 | RIMS-bindi | 22.50782 | 19.61629 | 22.41039 | 18.36859 | 23.01815 | 20.31138 | 3.213367 | 1.9648 | 4.461934 | 21.03877 | 6.14185 | 0.165072 | 0.00056 | 0.007302 | 1 | 1 |
| 119 | sp\|Q8VEK3 | HNRPU_M | 88 kDa | 0.999549 | 14 | 100% | Q8VEK3 | Hnrnpu | Heterogene | 26.3786 | 24.6574 | 26.38156 | 24.48236 | 26.37727 | 25.01479 | 1.660958 | 1.008343 | 2.313572 | 25.54866 | 6.057632 | 0.118341 | 0.000578 | 0.007469 | 1 | 1 |
| 825 | tr\|A0A075 | A0A075B5 | 11 kDa | 0.999375 | 3 | 33% | A0A075B5 | Ighv1-56 | Immunoglo | 23.47433 | 20.17281 | 22.34031 | 20.90487 | 23.77537 | 20.27581 | 2.745506 | 1.665334 | 3.825678 | 21.82392 | 6.065691 | 0.088595 | 0.000602 | 0.00764 | 1 | 1 |
| 511 | sp\|Q3TWN | CNNM2_M | 97 kDa | 0.999457 | 5 | 100% | Q3TWN3 | Cnnm2 | Metal trans | 20.05472 | 17.92137 | 19.41725 | 17.7696 | 20.64281 | 17.37158 | 2.350741 | 1.425285 | 3.276196 | 18.86289 | 6.060599 | 0.086212 | 0.000603 | 0.00764 | 1 | 1 |
| 1041 | sp\|Q8VHH5 | AGAP3_MC | 98 kDa | 0.999401 | 2 | 100% | Q8VHH5 | Agap3 | Arf-GAP wi | 21.07145 | 23.06676 | 21.07973 | 23.7278 | 21.0453 | 22.63263 | -2.0769 | -2.90335 | -1.25046 | 22.10394 | -5.98172 | 0.039839 | 0.000623 | 0.007788 | -1 | -1 |
| 1040 | sp\|P62267 | RS23_MOU | 16 kDa | 0.99942 | 2 | 100% | P62267 | Rps23 | 40S riboso | 24.25166 | 22.62279 | 24.47838 | 22.71259 | 24.08159 | 22.75469 | 1.573851 | 0.946897 | 2.200804 | 23.48362 | 5.974877 | 0.033478 | 0.000627 | 0.007788 | 1 | 1 |
| 199 | sp\|Q7TNC4 | LC7L2_MO | 47 kDa | 0.999471 | 10 | 100% | Q7TNC4 | Luc7l2 | Putative RN | 23.05372 | 24.66801 | 22.94288 | 24.84157 | 23.44575 | 24.92716 | -1.6648 | -2.32891 | -1.00068 | 23.97985 | -5.9665 | 0.024836 | 0.000632 | 0.007788 | -1 | -1 |
| 203 | sp\|O08917 | FLOT1_MO | 48 kDa | 0.9994 | 10 | 100% | O08917 | Flot1 | Flotillin-1 | 21.59521 | 23.52845 | 21.92339 | 24.57669 | 22.24405 | 24.3758 | -2.23943 | -3.13842 | -1.34045 | 23.0406 | -5.93855 | -0.02629 | 0.000668 | 0.008158 | -1 | -1 |
| 331 | sp\|Q99KP6 | PRP19_MO | 55 kDa | 0.999488 | 7 | 100% | Q99KP6 | Prpf19 | Pre-mRNA- | 24.82743 | 22.70567 | 24.27015 | 23.05885 | 25.17441 | 22.69926 | 1.93607 | 1.146878 | 2.725263 | 23.78929 | 5.839006 | -0.10786 | 0.000716 | 0.008616 | 1 | 1 |
| 485 | sp\|Q9D0L4 | ADCK1_MC | 60 kDa | 0.999797 | 5 | 100% | Q9D0L4 | Adck1 | Uncharacte | 22.14353 | 19.38703 | 21.67331 | 19.53578 | 22.35148 | 20.44388 | 2.267211 | 1.346956 | 3.187467 | 20.9225 | 5.877701 | -0.09952 | 0.000718 | 0.008616 | 1 | 1 |
| 710 | sp\|P08226 | APOE_MOU | 36 kDa | 0.998824 | 4 | 100% | P08226 | Apoe | Apolipopro | 21.16605 | 19.66038 | 21.29676 | 19.95859 | 21.53739 | 19.48436 | 1.632292 | 0.96341 | 2.301174 | 20.51726 | 5.808305 | -0.14014 | 0.000738 | 0.008781 | 1 | 1 |
| 235 | sp\|Q9Z2N8 | ACL6A_MO | 47 kDa | 0.999542 | 9 | 89% | Q9Z2N8 | Actl6a | Actin-like p | 24.21382 | 21.73671 | 23.18929 | 21.35708 | 24.51204 | 21.71926 | 2.367363 | 1.399225 | 3.335502 | 22.78803 | 5.8355 | -0.14725 | 0.000752 | 0.008839 | 1 | 1 |
| 405 | sp\|P62754 | RS6_MOUS | 29 kDa | 0.999619 | 6 | 100% | P62754 | Rps6 | 40S riboso | 25.81162 | 24.16112 | 26.02414 | 23.63338 | 25.38644 | 24.0423 | 1.795134 | 1.055764 | 2.534505 | 24.84316 | 5.778772 | -0.17132 | 0.00076 | 0.008839 | 1 | 1 |
| 1086 | sp\|P39654 | LOX15_MO | 75 kDa | 0.999073 | 2 | 100% | P39654 | Alox15 | Arachidona | 24.30309 | 26.42581 | 24.06726 | 25.96152 | 24.83022 | 26.41315 | -1.86664 | -2.63589 | -1.09739 | 25.33351 | -5.77558 | -0.1747 | 0.000763 | 0.008839 | -1 | -1 |
| 928 | sp\|P01831 | THY1_MOU | 18 kDa | 0.998266 | 3 | 100% | P01831 | Thy1 | Thy-1 mem | 18.01021 | 16.79774 | 18.45987 | 15.83689 | 18.24632 | 15.81 | 2.090589 | 1.221915 | 2.959262 | 17.19351 | 5.73728 | -0.23635 | 0.000814 | 0.009355 | 1 | 1 |
| 281 | sp\|Q61103 | REQU_MOU | 44 kDa | 0.999382 | 8 | 88% | Q61103 | Dpf2 | Zinc finger | 24.10588 | 21.64723 | 23.5233 | 21.22568 | 24.4496 | 22.29356 | 2.304103 | 1.347004 | 3.261203 | 22.87421 | 5.74507 | -0.24186 | 0.000822 | 0.009368 | 1 | 1 |
| 315 | sp\|Q8R081 | HNRPL_MC | 64 kDa | 0.999659 | 7 | 100% | Q8R081 | Hnrnpl | Heterogene | 23.52547 | 21.8458 | 23.09379 | 21.48003 | 23.40808 | 21.87588 | 1.608544 | 0.935155 | 2.281933 | 22.53818 | 5.685484 | -0.27059 | 0.000835 | 0.009433 | 1 | 1 |
| 535 | sp\|P50544 | ACADV_MC | 71 kDa | 0.999215 | 5 | 100% | P50544 | Acadvl | Very long-c | 18.81377 | 20.64122 | 18.5118 | 20.44809 | 19.08922 | 21.41604 | -2.03018 | -2.88479 | -1.17558 | 19.82002 | -5.66193 | -0.31351 | 0.000875 | 0.009789 | -1 | -1 |
| 170 | sp\|P48962 | ADT1_MOU | 33 kDa | 0.999351 | 11 | 55% | P48962 | Slc25a4 | ADP/ATP tr | 24.79143 | 23.2315 | 24.44439 | 22.75918 | 24.49906 | 23.01531 | 1.576296 | 0.909271 | 2.243321 | 23.79015 | 5.624665 | -0.33597 | 0.000888 | 0.009789 | 1 | 1 |
| 437 | sp\|O55203 | LDB2_MOU | 43 kDa | 0.999263 | 6 | 83% | O55203 | Ldb2 | LIM domain | 23.60646 | 21.88231 | 23.91088 | 20.79993 | 23.29214 | 21.68058 | 2.148885 | 1.242273 | 3.055498 | 22.52872 | 5.650481 | -0.32866 | 0.000888 | 0.009789 | 1 | 1 |
| 191 | sp\|Q3U0V1 | FUBP2_MC | 77 kDa | 0.999648 | 10 | 90% | Q3U0V1 | Khsrp | Far upstrea | 24.72273 | 21.66067 | 24.14036 | 21.8274 | 23.23654 | 21.44241 | 2.389714 | 1.380986 | 3.398441 | 22.83835 | 5.653567 | -0.33875 | 0.000901 | 0.009848 | 1 | 1 |
| 204 | sp\|Q60634 | FLOT2_MO | 47 kDa | 0.999346 | 10 | 100% | Q60634 | Flot2 | Flotillin-2 | 21.85661 | 23.9754 | 21.5404 | 23.67719 | 22.37026 | 23.52969 | -1.805 | -2.57745 | -1.03256 | 22.82493 | -5.56175 | -0.40414 | 0.000947 | 0.010239 | -1 | -1 |
| 1057 | sp\|A2AFS3 | K1324_MO | 111 kDa | 0.999283 | 2 | 100% | A2AFS3 | Kiaa1324 | UPF0577 p | 14.40482 | 16.87575 | 13.91451 | 16.51142 | 15.23356 | 16.78388 | -2.20605 | -3.14634 | -1.26577 | 15.62066 | -5.59896 | -0.39713 | 0.000951 | 0.010239 | -1 | -1 |
| 489 | sp\|P47963 | RL13_MOU | 24 kDa | 0.99974 | 5 | 100% | P47963 | Rpl13 | 60S riboso | 25.0958 | 23.73636 | 25.33176 | 23.65812 | 24.96917 | 23.67106 | 1.443731 | 0.824413 | 2.06305 | 24.41038 | 5.548471 | -0.41861 | 0.00096 | 0.010246 | 1 | 1 |
| 562 | sp\|P84751 | HVM63_M | 25 kDa | 0.998735 | 5 | 40% | P84751 | NA | Ig heavy ch | 17.50556 | 21.33888 | 17.75594 | 21.7733 | 16.89312 | 19.47553 | -3.4777 | -4.97845 | -1.97695 | 19.12372 | -5.53011 | -0.47133 | 0.00102 | 0.01075 | -1 | -1 |

| | | | | | | | | | | | | | | | | | | | | | | | | | |
|---|---|---|---|---|---|---|---|---|---|---|---|---|---|---|---|---|---|---|---|---|---|---|---|---|---|
| 1140 | sp\|Q8K2T1 | NMRL1_M | 34 kDa | 0.998537 | 2 | 100% | Q8K2T1 | Nmral1 | NmrA-like f | 18.37238 | 20.24967 | 17.58994 | 19.46955 | 18.02624 | 20.04258 | -1.92441 | -2.75928 | -1.08953 | 18.95839 | -5.48652 | -0.487 | 0.001024 | 0.01075 | -1 | -1 |
| 1164 | sp\|Q9JMD | ZN207_MO | 53 kDa | 0.998285 | 2 | 100% | Q9JMD0 | Znf207 | BUB3-inter | 19.74728 | 21.67862 | 19.75813 | 21.20502 | 19.82974 | 21.09754 | -1.54867 | -2.22141 | -0.87594 | 20.55272 | -5.47921 | -0.49445 | 0.001031 | 0.01075 | -1 | -1 |
| 553 | sp\|P62141 | PP1B_MOU | 37 kDa | 0.998911 | 5 | 40% | P62141 | Ppp1cb | Serine/thre | 22.1122 | 19.15572 | 21.42328 | 20.32593 | 22.05817 | 19.44851 | 2.221164 | 1.258729 | 3.183599 | 20.75397 | 5.507569 | -0.49578 | 0.001044 | 0.010776 | 1 | 1 |
| 491 | sp\|Q99LX5 | MMTA2_M | 29 kDa | 0.999696 | 5 | 100% | Q99LX5 | Mmtag2 | Multiple m | 22.0584 | 23.60252 | 22.02218 | 23.40948 | 21.98945 | 23.27442 | -1.40546 | -2.01824 | -0.79269 | 22.72608 | -5.4591 | -0.51659 | 0.001052 | 0.010776 | -1 | -1 |
| 1261 | sp\|P51660 | DHB4_MOU | 79 kDa | 0.996427 | 2 | 100% | P51660 | Hsd17b4 | Peroxisoma | 20.02291 | 18.00302 | 19.22207 | 17.81911 | 20.25533 | 17.99475 | 1.894478 | 1.067929 | 2.721027 | 18.8862 | 5.455607 | -0.52104 | 0.001057 | 0.010776 | 1 | 1 |
| 441 | sp\|P62717 | RL18A_MO | 21 kDa | 0.999204 | 6 | 100% | P62717 | Rpl18a | 60S ribosor | 23.75002 | 22.17157 | 23.98249 | 22.46415 | 23.73768 | 22.51228 | 1.440731 | 0.805961 | 2.0755 | 23.10303 | 5.402163 | -0.57961 | 0.001117 | 0.011262 | 1 | 1 |
| 787 | sp\|P00920 | CAH2_MOU | 29 kDa | 0.999666 | 3 | 100% | P00920 | Ca2 | Carbonic ar | 22.45722 | 24.61984 | 23.44527 | 24.91472 | 22.92938 | 24.74197 | -1.81489 | -2.61509 | -1.01468 | 23.8514 | -5.39817 | -0.58405 | 0.001121 | 0.011262 | -1 | -1 |
| 255 | sp\|P63038 | CH60_MOU | 61 kDa | 0.99996 | 8 | 100% | P63038 | Hspd1 | 60 kDa heat | 24.20734 | 25.35854 | 24.24799 | 25.71248 | 23.98566 | 25.76242 | -1.46415 | -2.11229 | -0.81602 | 24.87907 | -5.37678 | -0.60786 | 0.001147 | 0.011298 | -1 | -1 |
| 1227 | sp\|Q3UMR | MCU_MOU | 40 kDa | 0.997551 | 2 | 100% | Q3UMR5 | Mcu | Calcium un | 19.99982 | 22.60322 | 21.23592 | 22.51478 | 20.10491 | 22.5854 | -2.12092 | -3.05604 | -1.1858 | 21.50734 | -5.41262 | -0.59952 | 0.001151 | 0.011298 | -1 | -1 |
| 916.8 | sp\|C0HKE8 | H2A1O_M | 14 kDa | 0.998436 | 3 | 33% | C0HKE8 | Hist1h2ao | Histone H2 | 22.23442 | 23.69836 | 22.50782 | 23.97068 | 22.58551 | 24.18567 | -1.50899 | -2.17846 | -0.83952 | 23.19708 | -5.36487 | -0.62115 | 0.001161 | 0.011298 | -1 | -1 |
| 1093.2 | sp\|Q9CQL1 | MGN2_MO | 17 kDa | 0.999012 | 2 | 100% | Q9CQL1 | Magohb | Protein ma | 17.28198 | 20.77473 | 18.80031 | 21.09546 | 18.0593 | 20.05533 | -2.59464 | -3.74123 | -1.44806 | 19.34452 | -5.40035 | -0.61302 | 0.001165 | 0.011298 | -1 | -1 |
| 57 | sp\|Q8CG14 | CS1A_MOU | 77 kDa | 0.999695 | 19 | 53% | Q8CG14 | C1sa | Compleme | 25.23354 | 26.80708 | 25.39255 | 27.05831 | 24.99033 | 26.46789 | -1.57228 | -2.2704 | -0.87417 | 25.99162 | -5.36049 | -0.62604 | 0.001166 | 0.011298 | -1 | -1 |
| 452.1 | sp\|P62270 | RS18_MOU | 18 kDa | 0.999008 | 6 | 100% | P62270 | Rps18 | 40S ribosor | 24.85654 | 23.21343 | 24.95486 | 23.24909 | 24.33298 | 22.92725 | 1.584867 | 0.875634 | 2.294099 | 23.92236 | 5.318691 | -0.67285 | 0.001219 | 0.011522 | 1 | 1 |
| 452.2 | tr\|A0A1Y7 | A0A1Y7VKY | 18 kDa | 0.999008 | 6 | 100% | A0A1Y7VKY | Gm11361 | MCG11667 | 24.85654 | 23.21343 | 24.95486 | 23.24909 | 24.33298 | 22.92725 | 1.584867 | 0.875634 | 2.294099 | 23.92236 | 5.318691 | -0.67285 | 0.001219 | 0.011522 | 1 | 1 |
| 452.3 | tr\|F6YVP7\| | F6YVP7_M | 18 kDa | 0.999008 | 6 | 100% | F6YVP7 | Gm10260 | Predicted g | 24.85654 | 23.21343 | 24.95486 | 23.24909 | 24.33298 | 22.92725 | 1.584867 | 0.875634 | 2.294099 | 23.92236 | 5.318691 | -0.67285 | 0.001219 | 0.011522 | 1 | 1 |
| 777 | sp\|Q62159 | RHOC_MOU | 22 kDa | 0.99532 | 4 | 25% | Q62159 | Rhoc | Rho-related | 23.62226 | 22.13382 | 23.3445 | 21.15459 | 23.28133 | 21.55788 | 1.8006 | 0.994191 | 2.607008 | 22.51573 | 5.314763 | -0.67783 | 0.001225 | 0.011522 | 1 | 1 |
| 1172 | sp\|Q9QXB9 | DRG2_MOU | 41 kDa | 0.998227 | 2 | 100% | Q9QXB9 | Drg2 | Developme | 18.50962 | NA | 15.50291 | 25.30819 | 16.69878 | NA | -8.40442 | -11.5833 | -5.22555 | 19.00488 | -6.92992 | -0.19366 | 0.001232 | 0.011522 | -1 | -1 |
| 655 | sp\|P14206 | RSSA_MOU | 33 kDa | 0.999272 | 4 | 100% | P14206 | Rpsa | 40S ribosor | 21.95529 | 23.99335 | 22.54462 | 23.8496 | 22.24228 | 23.57012 | -1.55696 | -2.25852 | -0.8554 | 23.02587 | -5.28218 | -0.71396 | 0.001267 | 0.011767 | -1 | -1 |
| 109 | sp\|P62702 | RS4X_MOU | 30 kDa | 0.999356 | 15 | 100% | P62702 | Rps4x | 40S ribosor | 24.89577 | 23.38949 | 24.89357 | 23.57903 | 24.68421 | 23.46402 | 1.346999 | 0.738328 | 1.95567 | 24.15102 | 5.267271 | -0.7308 | 0.001287 | 0.011873 | 1 | 1 |
| 546 | sp\|P98086 | C1QA_MOU | 26 kDa | 0.999049 | 5 | 100% | P98086 | C1qa | Compleme | 25.07402 | 26.66377 | 25.41641 | 27.02354 | 24.78715 | 26.46682 | -1.62552 | -2.36253 | -0.88851 | 25.90529 | -5.24954 | -0.75086 | 0.001312 | 0.012017 | -1 | -1 |
| 104 | sp\|Q6A068 | CDC5L_MO | 92 kDa | 0.9996 | 15 | 27% | Q6A068 | Cdc5l | Cell divisio | 22.38783 | 20.79496 | 22.74659 | 20.95253 | 22.79606 | 21.42558 | 1.585804 | 0.865296 | 2.306312 | 21.85059 | 5.238554 | -0.76332 | 0.001327 | 0.012077 | 1 | 1 |
| 75 | sp\|P16546 | SPTN1_MO | 285 kDa | 0.999629 | 17 | 100% | P16546 | Sptan1 | Spectrin alp | 24.12955 | 25.25185 | 23.58235 | 25.19174 | 23.77182 | 25.49079 | -1.48355 | -2.16013 | -0.80698 | 24.56969 | -5.21903 | -0.7855 | 0.001355 | 0.012249 | -1 | -1 |
| 176 | sp\|Q9Z204 | HNRPC_M | 34 kDa | 0.999252 | 11 | 100% | Q9Z204 | Hnrnpc | Heterogene | 25.77075 | 23.35799 | 24.80237 | 23.26727 | 26.29288 | 23.58338 | 2.21912 | 1.21016 | 3.228079 | 24.51244 | 5.248771 | -0.78161 | 0.001365 | 0.012261 | 1 | 1 |
| 87 | sp\|P61979 | HNRPK_M | 51 kDa | 0.999832 | 16 | 100% | P61979 | Hnrnpk | Heterogene | 25.68902 | 24.44393 | 25.63567 | 24.37872 | 25.64861 | 24.2448 | 1.301951 | 0.705108 | 1.898794 | 25.00679 | 5.192008 | -0.8163 | 0.001395 | 0.012336 | 1 | 1 |
| 7 | sp\|P70336 | ROCK2_M | 161 kDa | 0.99975 | 49 | 100% | P70336 | Rock2 | Rho-associa | 27.49656 | 24.87803 | 28.27345 | 24.56162 | 26.65873 | 25.46058 | 2.509501 | 1.362953 | 3.656049 | 26.22149 | 5.223309 | -0.81026 | 0.001403 | 0.012336 | 1 | 1 |
| 386 | sp\|Q6NXH9 | K2C73_M | 59 kDa | 0.997663 | 7 | 29% | Q6NXH9 | Krt73 | Keratin, typ | 19.95852 | 21.63969 | 18.08747 | 21.57577 | 19.05922 | 22.74447 | -2.95158 | -4.30129 | -1.60187 | 20.51086 | -5.21873 | -0.81543 | 0.001409 | 0.012336 | -1 | -1 |
| 574 | sp\|P35276 | RAB3D_M | 24 kDa | 0.998133 | 5 | 20% | P35276 | Rab3d | Ras-related | 23.05199 | 21.57264 | 22.76164 | 21.55718 | 22.76453 | 21.38654 | 1.353929 | 0.73204 | 1.975819 | 22.18242 | 5.181839 | -0.82792 | 0.00141 | 0.012336 | 1 | 1 |
| 482 | sp\|O35295 | PURB_MOU | 34 kDa | 0.99993 | 5 | 100% | O35295 | Purb | Transcripti | 22.71308 | 20.3186 | 21.9463 | 20.82005 | 22.24894 | 20.75416 | 1.671833 | 0.901958 | 2.441707 | 21.46685 | 5.168606 | -0.84306 | 0.00143 | 0.012407 | 1 | 1 |
| 541 | sp\|P18529 | HVM58_M | 13 kDa | 0.999137 | 5 | 20% | P18529 | NA | Ig heavy ch | 20.90458 | 22.80126 | 21.44474 | 23.44875 | 20.59252 | 22.53165 | -1.9466 | -2.84138 | -1.05183 | 21.95392 | -5.1863 | -0.84052 | 0.001437 | 0.012407 | -1 | -1 |
| 1133 | sp\|Q9ERU9 | RBP2_MOU | 341 kDa | 0.998587 | 2 | 100% | Q9ERU9 | Ranbp2 | E3 SUMO-p | 20.797 | 19.66775 | 21.43199 | 19.14767 | 20.93915 | 19.54311 | 1.603206 | 0.861802 | 2.34461 | 20.25445 | 5.146776 | -0.8681 | 0.001464 | 0.012567 | 1 | 1 |
| 277 | sp\|P62281 | RS11_MOU | 18 kDa | 0.999465 | 8 | 100% | P62281 | Rps11 | 40S ribosor | 24.70272 | 23.4546 | 24.58657 | 22.66491 | 24.48314 | 22.88109 | 1.590609 | 0.853852 | 2.327365 | 23.79551 | 5.138543 | -0.87756 | 0.001477 | 0.012599 | 1 | 1 |
| 347 | sp\|P01837 | IGKC_MOU | 12 kDa | 0.999358 | 7 | 100% | P01837 | Igkc | Immunoglo | 25.62897 | 27.77326 | 26.25782 | 28.12965 | 25.62317 | 27.19193 | -1.86163 | -2.72373 | -0.99952 | 26.76746 | -5.14784 | -0.88434 | 0.001497 | 0.012686 | -1 | -1 |
| 685 | sp\|Q6ZWU | RS27_MOU | 9 kDa | 0.999015 | 4 | 50% | Q6ZWU9 | Rps27 | 40S ribosor | 21.56806 | 19.69359 | 21.59014 | 20.16681 | 21.07589 | 19.46714 | 1.63552 | 0.873956 | 2.397085 | 20.59361 | 5.111524 | -0.90867 | 0.001521 | 0.012812 | 1 | 1 |
| 607 | sp\|P62751 | RL23A_MO | 18 kDa | 0.999674 | 4 | 100% | P62751 | Rpl23a | 60S ribosor | 24.08806 | 22.72058 | 24.21192 | 22.90146 | 24.01876 | 22.82157 | 1.291708 | 0.688635 | 1.89478 | 23.46039 | 5.097951 | -0.92435 | 0.001544 | 0.012892 | 1 | 1 |
| 833 | sp\|P01029 | CO4B_MOU | 193 kDa | 0.99931 | 3 | 100% | P01029 | C4b | Compleme | 25.37734 | 26.83135 | 24.74714 | 26.71139 | 25.56214 | 27.20751 | -1.68787 | -2.47646 | -0.89929 | 26.07281 | -5.0944 | -0.92845 | 0.00155 | 0.012892 | -1 | -1 |
| 1142.1 | sp\|P47915 | RL29_MOU | 18 kDa | 0.998518 | 2 | 100% | P47915 | Rpl29 | 60S ribosor | 18.46726 | 20.34786 | 18.94775 | 21.21091 | 17.87443 | 20.19413 | -2.15449 | -3.15835 | -1.15062 | 19.50706 | -5.12177 | -0.92546 | 0.001563 | 0.012896 | -1 | -1 |
| 1169 | sp\|Q7TQA1 | IGSF1_MO | 147 kDa | 0.998241 | 2 | 100% | Q7TQA1 | Igsf1 | Immunoglo | 20.38809 | 23.01611 | 21.21515 | 22.93423 | 20.39119 | 22.10245 | -2.01945 | -2.96112 | -1.07778 | 21.67454 | -5.11784 | -0.92995 | 0.001569 | 0.012896 | -1 | -1 |
| 699 | sp\|P33267 | CP2F2_MO | 56 kDa | 0.998901 | 4 | 100% | P33267 | Cyp2f2 | Cytochrom | 19.63472 | 17.50575 | 19.24635 | 17.57778 | 19.99771 | 18.32354 | 1.823905 | 0.969059 | 2.678751 | 18.71431 | 5.085218 | -0.95373 | 0.001596 | 0.013037 | 1 | 1 |
| 1060.2 | tr\|A2A4X6 | A2A4X6_M | 19 kDa | 0.999248 | 2 | 100% | A2A4X6 | Gm12355 | MCG21910 | 20.09105 | 21.88482 | 20.20941 | 21.18869 | 19.83026 | 21.66737 | -1.53672 | -2.2627 | -0.81074 | 20.81193 | -5.03813 | -0.99375 | 0.001648 | 0.013378 | -1 | -1 |
| 592 | sp\|Q9D1G1 | RAB1B_MO | 22 kDa | 0.999802 | 4 | 25% | Q9D1G1 | Rab1b | Ras-related | 19.76854 | 18.12916 | 20.66956 | 18.64778 | 19.87042 | 18.24636 | 1.761741 | 0.926592 | 2.596891 | 19.22197 | 5.02111 | -1.0141 | 0.00168 | 0.01351 | 1 | 1 |
| 916.15 | tr\|Q8CGP4 | Q8CGP4_M | 14 kDa | 0.998436 | 3 | 33% | Q8CGP4 | Hist1h2aa | Histone H2 | 22.70511 | 24.71866 | 21.19408 | 24.28151 | 22.63311 | 24.54156 | -2.33648 | -3.4401 | -1.23286 | 23.34567 | -5.05234 | -1.00511 | 0.001684 | 0.01351 | -1 | -1 |
| 142 | sp\|Q8BG05 | ROA3_MOU | 40 kDa | 0.999911 | 12 | 75% | Q8BG05 | Hnrnpa3 | Heterogene | 25.09629 | 23.31561 | 24.26269 | 23.01576 | 24.88287 | 23.11078 | 1.599898 | 0.839059 | 2.360738 | 23.94733 | 5.004955 | -1.03247 | 0.001709 | 0.013629 | 1 | 1 |
| 719 | sp\|Q02819 | NUCB1_M | 53 kDa | 0.998696 | 4 | 100% | Q02819 | Nucb1 | Nucleobind | 20.65448 | 21.99998 | 20.00173 | 22.41841 | 20.34966 | 21.62265 | -1.67839 | -2.47787 | -0.87891 | 21.17448 | -4.99675 | -1.04207 | 0.001724 | 0.013654 | -1 | -1 |
| 1250 | sp\|Q9D1R9 | RL34_MOU | 13 kDa | 0.996907 | 2 | 100% | Q9D1R9 | Rpl34 | 60S ribosor | 18.36554 | 20.49861 | 16.84509 | 19.59845 | 16.29315 | 20.05903 | -2.88411 | -4.25345 | -1.51476 | 18.60998 | -5.02631 | -1.03516 | 0.001732 | 0.013654 | -1 | -1 |
| 1259 | sp\|Q9CU62 | SMC1A_M | 143 kDa | 0.996524 | 2 | 100% | Q9CU62 | Smc1a | Structural | 19.69182 | 17.35776 | 20.37165 | 18.03578 | 19.11579 | 16.78584 | 2.333295 | 1.22361 | 3.44298 | 18.55977 | 5.017885 | -1.0449 | 0.001748 | 0.013699 | 1 | 1 |
| 23 | sp\|Q61768 | KINH_MOU | 110 kDa | 0.999746 | 27 | 81% | Q61768 | Kif5b | Kinesin-1 h | 24.23015 | 22.50157 | 24.08387 | 22.4242 | 24.27546 | 23.13422 | 1.509827 | 0.785926 | 2.233728 | 23.44158 | 4.964192 | -1.08027 | 0.001787 | 0.013927 | 1 | 1 |
| 111 | sp\|Q62095 | DDX3Y_M | 73 kDa | 0.997675 | 15 | 13% | Q62095 | Ddx3y | ATP-depen | 19.10992 | 17.52939 | 19.08762 | 16.97791 | 18.737 | 17.74815 | 1.559697 | 0.805989 | 2.313406 | 18.19833 | 4.925359 | -1.12604 | 0.001865 | 0.014455 | 1 | 1 |
| 788 | sp\|P16125 | LDHB_MOU | 37 kDa | 0.999654 | 3 | 67% | P16125 | Ldhb | L-lactate de | 21.37743 | 17.29409 | 19.77552 | 17.52084 | 22.76833 | 18.14281 | 3.654516 | 1.893459 | 5.415572 | 19.47984 | 4.952303 | -1.12114 | 0.001877 | 0.014463 | 1 | 1 |
| 765 | sp\|P12382 | PFKAL_MO | 85 kDa | 0.997215 | 4 | 50% | P12382 | Pfkl | ATP-depen | 20.07549 | 21.14724 | 20.12847 | 21.53602 | 19.63317 | 21.39185 | -1.41266 | -2.0977 | -0.72762 | 20.65204 | -4.90822 | -1.14632 | 0.001901 | 0.014566 | -1 | -1 |
| 140 | sp\|Q6ZQ38 | CAND1_M | 136 kDa | 0.999055 | 13 | 100% | Q6ZQ38 | Cand1 | Cullin-asso | 23.07882 | 25.01108 | 23.75891 | 24.58636 | 23.03663 | 25.13218 | -1.61842 | -2.40482 | -0.83203 | 24.10066 | -4.89837 | -1.15798 | 0.001922 | 0.014644 | -1 | -1 |
| 472 | tr\|A0A075 | A0A075B5 | 13 kDa | 0.997707 | 6 | 17% | A0A075B5 | Ighv1-22 | Immunoglo | 15.67546 | 17.83639 | 16.93668 | 19.07442 | 16.01395 | 18.7235 | -2.33607 | -3.46928 | -1.20286 | 17.37673 | -4.91956 | -1.15945 | 0.001946 | 0.014741 | -1 | -1 |
| 1098 | sp\|Q8CH18 | CCAR1_M | 132 kDa | 0.99899 | 2 | 100% | Q8CH18 | Ccar1 | Cell divisio | 16.3035 | 21.98187 | 16.40305 | 19.46667 | 17.0631 | 19.23739 | -3.63876 | -5.41676 | -1.86076 | 18.40926 | -4.88395 | -1.20129 | 0.002024 | 0.015179 | -1 | -1 |
| 430 | sp\|P01810 | HVM40_M | 13 kDa | 0.99932 | 6 | 17% | P01810 | NA | Ig heavy ch | 23.84835 | 25.8581 | 24.30025 | 25.89394 | 23.23519 | 25.21181 | -1.86002 | -2.77044 | -0.94959 | 24.7246 | -4.87054 | -1.20704 | 0.002026 | 0.015179 | -1 | -1 |
| 503 | sp\|Q9JM93 | AR6P4_MO | 26 kDa | 0.999531 | 5 | 100% | Q9JM93 | Arl6ip4 | ADP-ribosy | 22.14555 | 23.97164 | 22.74135 | 24.49789 | 23.04196 | 24.73911 | -1.75993 | -2.62535 | -0.89451 | 23.52292 | -4.848 | -1.23353 | 0.002077 | 0.015474 | -1 | -1 |
| 616 | sp\|P62821 | RAB1A_MO | 23 kDa | 0.999554 | 4 | 25% | P62821 | Rab1A | Ras-related | 19.98859 | 18.9982 | 20.65602 | 18.91909 | 20.51377 | 18.20002 | 1.680356 | 0.845515 | 2.515197 | 19.54595 | 4.790927 | -1.28672 | 0.00217 | 0.016076 | 1 | 1 |
| 631 | sp\|Q924W | RBTN1_MO | 18 kDa | 0.999432 | 4 | 25% | Q924W9 | Lmo1 | Rhombotin | 20.35864 | 15.1481 | 20.83886 | 17.96954 | 19.56281 | 16.95335 | 3.563107 | 1.795188 | 5.331027 | 18.47189 | 4.809689 | -1.28917 | 0.002197 | 0.016125 | 1 | 1 |
| 524 | sp\|Q9CZM2 | RL15_MOU | 24 kDa | 0.999297 | 5 | 100% | Q9CZM2 | Rpl15 | 60S ribosor | 24.96063 | 23.24249 | 24.88872 | 23.63036 | 24.9648 | 23.87104 | 1.356754 | 0.680046 | 2.033462 | 24.25967 | 4.772004 | -1.30903 | 0.002215 | 0.016125 | 1 | 1 |
| 741 | sp\|P49615 | CDK5_MOU | 33 kDa | 0.9981 | 4 | 100% | P49615 | Cdk5 | Cyclin-depe | 20.73597 | 19.32378 | 21.71029 | 19.15244 | 20.60443 | 19.31223 | 1.754077 | 0.880937 | 2.627218 | 20.13986 | 4.789152 | -1.30366 | 0.002218 | 0.016125 | 1 | 1 |
| 1114 | sp\|Q9DCX2 | ATP5H_MO | 19 kDa | 0.998808 | 2 | 100% | Q9DCX2 | Atp5pd | ATP syntha | 24.55405 | 25.93757 | 23.79019 | 25.79254 | 24.63254 | 25.86288 | -1.53873 | -2.30744 | -0.77003 | 25.09496 | -4.76438 | -1.31822 | 0.002234 | 0.016125 | -1 | -1 |
| 328 | sp\|Q8BRM | GORAB_M | 41 kDa | 0.999541 | 7 | 100% | Q8BRM2 | Gorab | RAB6-inter | 24.13988 | 21.38348 | 22.70421 | 21.12528 | 23.93925 | 21.7973 | 2.159091 | 1.084359 | 3.233824 | 22.5149 | 4.794256 | -1.30754 | 0.002236 | 0.016125 | 1 | 1 |
| 812 | sp\|Q61753 | SERA_MOU | 57 kDa | 0.999441 | 3 | 100% | Q61753 | Phgdh | D-3-phosph | 18.26501 | 15.45136 | 18.07564 | 14.6353 | 17.04925 | 16.12053 | 2.394237 | 1.1893 | 3.599173 | 16.59952 | 4.741913 | -1.37011 | 0.002371 | 0.017008 | 1 | 1 |
| 944 | sp\|Q8K019 | BCLF1_MO | 106 kDa | 0.998011 | 3 | 100% | Q8K019 | Bclaf1 | Bcl-2-assoc | 21.51691 | 20.60907 | 21.94028 | 20.36035 | 22.17413 | 20.55908 | 1.367607 | 0.669924 | 2.065289 | 21.1933 | 4.665565 | -1.43814 | 0.0025 | 0.017844 | 1 | 1 |
| 83 | sp\|Q8VDN2 | AT1A1_MO | 113 kDa | 0.998975 | 17 | 59% | Q8VDN2 | Atp1a1 | Sodium/po | 23.47933 | 25.19346 | 24.01967 | 24.79375 | 23.26609 | 25.25398 | -1.49204 | -2.25466 | -0.72942 | 24.33438 | -4.65665 | -1.44903 | 0.002526 | 0.017932 | -1 | -1 |
| 163 | sp\|Q8BPB5 | FBLN3_MO | 55 kDa | 0.999504 | 11 | 100% | Q8BPB5 | Efemp1 | EGF-contai | 25.46593 | 23.13289 | 24.44394 | 23.06345 | 25.39581 | 23.70309 | 1.802079 | 0.881151 | 2.723007 | 24.20085 | 4.668441 | -1.45602 | 0.002566 | 0.017983 | 1 | 1 |
| 341 | sp\|P50516 | VATA_MOU | 68 kDa | 0.999405 | 7 | 100% | P50516 | Atp6v1a | V-type prot | 22.05077 | 24.05637 | 22.30866 | 23.72622 | 22.79099 | 23.69231 | -1.4415 | -2.18053 | -0.70246 | 23.10422 | -4.64249 | -1.46637 | 0.002567 | 0.017983 | -1 | -1 |
| 871 | sp\|P48758 | CBR1_MOU | 31 kDa | 0.998987 | 3 | 67% | P48758 | Cbr1 | Carbonyl re | 20.78965 | 22.03135 | 21.29824 | 22.7534 | 20.54205 | 22.71952 | -1.62478 | -2.45805 | -0.7915 | 21.68903 | -4.64116 | -1.46843 | 0.002573 | 0.017983 | -1 | -1 |
| 372 | sp\|P80560 | PTPR2_MO | 111 kDa | 0.998928 | 7 | 86% | P80560 | Ptprn2 | Receptor-ty | 21.65221 | 22.84576 | 21.98758 | 23.17468 | 21.52102 | 23.67379 | -1.51114 | -2.287 | -0.73528 | 22.47584 | -4.63579 | -1.47458 | 0.002587 | 0.017991 | -1 | -1 |
| 1046 | sp\|Q80TT2 | BAIP3_MO | 127 kDa | 0.999382 | 2 | 100% | Q80TT2 | Baiap3 | BAI1-associ | 18.3657 | 15.61973 | 18.83519 | 17.41713 | 19.38739 | 16.12937 | 2.474014 | 1.205602 | 3.742425 | 17.62575 | 4.654709 | -1.47529 | 0.002616 | 0.018057 | 1 | 1 |
| 790 | sp\|P03976 | KV2A5_MO | 12 kDa | 0.999638 | 3 | 33% | P03976 | NA | Ig kappa ch | 21.81688 | 19.43468 | 20.28002 | 18.3122 | 21.67842 | 17.72153 | 2.768972 | 1.348592 | 4.189351 | 19.87395 | 4.652269 | -1.47825 | 0.002623 | 0.018057 | 1 | 1 |
| 1191.1 | sp\|P36536 | SAR1A_MO | 22 kDa | 0.998052 | 2 | 100% | P36536 | Sar1a | GTP-bindin | 17.67067 | 20.11671 | 18.4915 | 19.82267 | 18.71094 | 19.83924 | -1.63517 | -2.47763 | -0.7927 | 19.10862 | -4.62303 | -1.49671 | 0.002649 | 0.018142 | -1 | -1 |
| 1015.1 | sp\|P47758 | SRPRB_MO | 30 kDa | 0.99986 | 2 | 100% | P47758 | Srprb | Signal reco | 21.12096 | 23.99881 | 21.22367 | 23.19838 | 22.45571 | 24.18518 | -2.19401 | -3.32386 | -1.06417 | 22.69712 | -4.63416 | -1.50024 | 0.002678 | 0.018163 | -1 | -1 |
| 1216 | sp\|Q3UPF5 | ZCCHV_MO | 107 kDa | 0.997739 | 2 | 100% | Q3UPF5 | Zc3hav1 | Zinc finger | 20.17213 | 21.26897 | 20.4747 | 22.14526 | 20.01053 | 21.8335 | -1.53012 | -2.32098 | -0.73927 | 20.98418 | -4.60499 | -1.5124 | 0.002681 | 0.018163 | -1 | -1 |
| 859.2 | sp\|Q6PDQ2 | CHD4_MOU | 218 kDa | 0.99907 | 3 | 100% | Q6PDQ2 | Chd4 | Chromodo | 22.31997 | 24.12919 | 22.92134 | 24.56679 | 22.56858 | 23.73819 | -1.54143 | -2.33953 | -0.74333 | 23.37401 | -4.59694 | -1.52232 | 0.002706 | 0.018163 | -1 | -1 |
| 860 | sp\|Q7TMF3 | NDUAC_M | 17 kDa | 0.999069 | 3 | 100% | Q7TMF3 | Ndufa12 | NADH dehy | 22.31997 | 24.12919 | 22.92134 | 24.56679 | 22.56858 | 23.73819 | -1.54143 | -2.33953 | -0.74333 | 23.37401 | -4.59694 | -1.52232 | 0.002706 | 0.018163 | -1 | -1 |
| 1153 | sp\|Q7TNG8 | LDHD_MOU | 52 kDa | 0.998398 | 2 | 100% | Q7TNG8 | Ldhd | Probable D | 19.67819 | 21.57733 | 19.17163 | 21.6382 | 20.60031 | 22.08775 | -1.95105 | -2.9622 | -0.9399 | 20.79223 | -4.60474 | -1.53608 | 0.002769 | 0.0185 | -1 | -1 |
| 310 | sp\|Q8VH51 | RBM39_M | 59 kDa | 0.999697 | 7 | 100% | Q8VH51 | Rbm39 | RNA-bindin | 23.28023 | 24.50378 | 23.38271 | 24.53827 | 23.06747 | 24.29109 | -1.20091 | -1.82752 | -0.5743 | 23.84393 | -4.56156 | -1.56599 | 0.002819 | 0.018737 | -1 | -1 |
| 1195 | sp\|Q9EQ06 | DHB11_M | 33 kDa | 0.998018 | 2 | 100% | Q9EQ06 | Hsd17b11 | Estradiol 17 | 17.55304 | 20.36387 | 17.31148 | 19.48534 | 18.65086 | 19.80308 | -2.04564 | -3.11407 | -0.97721 | 18.86128 | -4.56912 | -1.57965 | 0.002885 | 0.019082 | -1 | -1 |
| 1056.2 | tr\|Q63ZW6 | Q63ZW6_N | 162 kDa | 0.999285 | 2 | 100% | Q63ZW6 | Col4a5 | Col4a5 pro | 19.54683 | 18.24412 | 19.82987 | 18.05429 | 19.42494 | 16.95742 | 1.848605 | 0.881459 | 2.815752 | 18.67624 | 4.561446 | -1.58906 | 0.00291 | 0.019158 | 1 | 1 |
| 1022 | sp\|P00687 | AMY1_MOU | 58 kDa | 0.999718 | 2 | 100% | P00687 | Amy1 | Alpha-amyl | 22.9301 | 20.18507 | 21.64963 | 18.50636 | 23.00559 | 20.3898 | 2.834698 | 1.34859 | 4.320806 | 21.11109 | 4.55205 | -1.6006 | 0.002942 | 0.019244 | 1 | 1 |
| 1156 | sp\|P84309 | ADCY5_MO | 139 kDa | 0.998382 | 2 | 100% | P84309 | Adcy5 | Adenylate c | 18.79536 | 19.80088 | 18.23325 | 20.04702 | 17.82734 | 19.62704 | -1.53966 | -2.35002 | -0.72931 | 19.05515 | -4.52242 | -1.61494 | 0.002952 | 0.019244 | -1 | -1 |
| 983 | sp\|Q8BU30 | SYIC_MOUS | 144 kDa | 0.997151 | 3 | 100% | Q8BU30 | Iars | Isoleucine- | 17.57924 | 19.69756 | 15.54359 | 19.24074 | 16.4751 | 18.52327 | -2.62121 | -4.00471 | -1.23771 | 17.84325 | -4.5214 | -1.63832 | 0.003048 | 0.019775 | -1 | -1 |
| 1212 | sp\|P70280 | VAMP7_M | 25 kDa | 0.997775 | 2 | 100% | P70280 | Vamp7 | Vesicle-ass | 21.74346 | 23.0541 | 21.66027 | 23.37537 | 22.18273 | 23.04085 | -1.29462 | -1.98137 | -0.60788 | 22.50946 | -4.48695 | -1.65872 | 0.003075 | 0.019857 | -1 | -1 |
| 901 | sp\|P61255 | RL26_MOU | 17 kDa | 0.99868 | 3 | 100% | P61255 | Rpl26 | 60S ribosor | 24.22743 | 25.74522 | 24.67834 | 25.54254 | 24.42699 | 25.67184 | -1.20895 | -1.85119 | -0.56671 | 25.04873 | -4.48036 | -1.66695 | 0.003099 | 0.019916 | -1 | -1 |
| 824 | tr\|A0A140 | A0A140T8 | 13 kDa | 0.999377 | 3 | 100% | A0A140T8 | Igkv13-84 | Immunoglo | 23.25403 | 22.29525 | 23.27321 | 21.35865 | 23.14747 | 21.18339 | 1.61248 | 0.747207 | 2.477753 | 22.41867 | 4.442578 | -1.72742 | 0.0033 | 0.021108 | 1 | 1 |
| 436 | sp\|Q8VED5 | K2C79_MO | 58 kDa | 0.99927 | 6 | 17% | Q8VED5 | Krt79 | Keratin, typ | 24.02412 | 24.99836 | 23.48772 | 25.05226 | 23.6187 | 25.98452 | -1.63486 | -2.5154 | -0.75432 | 24.52761 | -4.42615 | -1.74796 | 0.003364 | 0.021255 | -1 | -1 |
| 958 | sp\|P99029 | PRDX5_M | 22 kDa | 0.997769 | 3 | 100% | P99029 | Prdx5 | Peroxiredo | 21.3084 | 22.36403 | 21.05864 | 22.15546 | 20.97149 | 22.29167 | -1.15754 | -1.78239 | -0.5327 | 21.69161 | -4.40927 | -1.75617 | 0.00337 | 0.021255 | -1 | -1 |
| 311 | tr\|A0A075 | A0A075B5 | 13 kDa | 0.999685 | 7 | 43% | A0A075B5 | Ighv4-1 | Immunoglo | 26.70387 | 28.2283 | 26.98323 | 28.15251 | 26.16557 | 27.77908 | -1.43574 | -2.21077 | -0.66071 | 27.33543 | -4.40917 | -1.75629 | 0.00337 | 0.021255 | -1 | -1 |
| 560 | sp\|P14106 | C1QB_MOU | 27 kDa | 0.998823 | 5 | 100% | P14106 | C1qb | Compleme | 26.98189 | 28.1934 | 27.18665 | 28.42181 | 26.79338 | 28.00411 | -1.21913 | -1.8792 | -0.55907 | 27.59687 | -4.39606 | -1.77284 | 0.003423 | 0.021488 | -1 | -1 |
| 894 | sp\|P21107 | TPM3_MOU | 33 kDa | 0.998754 | 3 | 33% | P21107 | Tpm3 | Tropomyos | 23.42961 | 22.32265 | 24.02872 | 22.19628 | 23.23099 | 22.03522 | 1.378388 | 0.630675 | 2.126101 | 22.87391 | 4.387702 | -1.78339 | 0.003457 | 0.021602 | 1 | 1 |
| 1019 | sp\|P07759 | SPA3K_MO | 47 kDa | 0.999735 | 2 | 100% | P07759 | Serpina3k | Serine prot | 18.43752 | 14.04793 | 19.3601 | NA | 18.39439 | 16.31902 | 3.547192 | 1.705851 | 5.388533 | 17.31179 | 4.775089 | -1.56869 | 0.003536 | 0.021945 | 1 | 1 |
| 917 | sp\|Q6IME9 | K2C72_MO | 57 kDa | 0.998428 | 3 | 33% | Q6IME9 | Krt72 | Keratin, typ | 18.93367 | 20.01606 | 18.19652 | 19.8107 | 18.74804 | 20.04097 | -1.32983 | -2.05467 | -0.60499 | 19.291 | -4.36672 | -1.80994 | 0.003544 | 0.021945 | -1 | -1 |
| 77 | sp\|Q91XU0 | WRIP1_M | 72 kDa | 0.999602 | 17 | 100% | Q91XU0 | Wrnip1 | ATPase WR | 25.96271 | 22.74806 | 24.0297 | 22.65917 | 26.33713 | 22.96971 | 2.650867 | 1.199294 | 4.10244 | 24.11775 | 4.358126 | -1.84173 | 0.003688 | 0.02264 | 1 | 1 |
| 990 | sp\|Q3THK7 | GUAA_MOU | 77 kDa | 0.99685 | 3 | 100% | Q3THK7 | Gmps | GMP synth | 19.43538 | 18.31149 | 19.37049 | 17.27847 | 19.24929 | 16.69458 | 1.92354 | 0.870142 | 2.976938 | 18.38995 | 4.357721 | -1.84224 | 0.00369 | 0.02264 | 1 | 1 |
| 456 | sp\|Q6ZQ58 | LARP1_MO | 121 kDa | 0.998957 | 6 | 100% | Q6ZQ58 | Larp1 | La-related p | 20.37757 | 22.14973 | 18.76882 | 21.93413 | 20.51588 | 22.09752 | -2.17303 | -3.36601 | -0.98005 | 20.97394 | -4.34694 | -1.85581 | 0.003737 | 0.022827 | -1 | -1 |
| 136 | sp\|P14148 | RL7_MOUS | 31 kDa | 0.99931 | 13 | 100% | P14148 | Rpl7 | 60S ribosor | 25.84312 | 24.28867 | 26.35916 | 24.82055 | 25.8448 | 24.87377 | 1.354696 | 0.607028 | 2.102364 | 25.33834 | 4.312547 | -1.87881 | 0.003781 | 0.022951 | 1 | 1 |
| 321 | sp\|O88665 | BRD7_MOU | 74 kDa | 0.999593 | 7 | 100% | O88665 | Brd7 | Bromodom | 22.8287 | 20.91494 | 21.83513 | 20.89656 | 23.0596 | 20.983 | 1.642973 | 0.737377 | 2.54857 | 21.75299 | 4.325039 | -1.87532 | 0.003791 | 0.022951 | 1 | 1 |
| 178 | sp\|Q68FH0 | PKP4_MOU | 132 kDa | 0.999035 | 11 | 100% | Q68FH0 | Pkp4 | Plakophilin | 22.72108 | 20.14036 | 21.18809 | 19.96626 | 21.97705 | 20.28417 | 1.83181 | 0.814718 | 2.848901 | 21.04617 | 4.298044 | -1.91761 | 0.00396 | 0.02377 | 1 | 1 |
| 28 | sp\|A2BH40 | ARI1A_MO | 242 kDa | 0.999848 | 25 | 92% | A2BH40 | Arid1a | AT-rich inte | 23.41653 | 22.00725 | 22.88232 | 21.53515 | 23.66679 | 22.13474 | 1.429503 | 0.633365 | 2.225641 | 22.60713 | 4.273634 | -1.92856 | 0.003962 | 0.02377 | 1 | 1 |
| 702 | sp\|Q9WV5 | VAPA_MOU | 28 kDa | 0.998865 | 4 | 75% | Q9WV55 | Vapa | Vesicle-ass | 21.03147 | 20.30019 | 21.34536 | 19.71504 | 21.1408 | 19.77819 | 1.241406 | 0.543529 | 1.939283 | 20.55184 | 4.233854 | -1.97965 | 0.004157 | 0.024829 | 1 | 1 |
| 654 | tr\|E9Q0F0 | E9Q0F0_M | 112 kDa | 0.999273 | 4 | 50% | E9Q0F0 | Krt78 | Keratin 78 | 25.82564 | 27.54229 | 26.53719 | 27.80519 | 26.68771 | 28.27118 | -1.5227 | -2.379 | -0.66641 | 27.11154 | -4.23894 | -1.98449 | 0.004199 | 0.024944 | -1 | -1 |
| 717 | sp\|Q9CYH2 | PXL2A_MO | 24 kDa | 0.998714 | 4 | 100% | Q9CYH2 | Prxl2a | Peroxiredo | 20.74462 | 19.6203 | 21.02182 | 19.35665 | 20.32793 | 19.31563 | 1.267262 | 0.552984 | 1.981541 | 20.06449 | 4.222793 | -1.9939 | 0.004213 | 0.024944 | 1 | 1 |
| 567 | sp\|P58269 | DPF3_MOU | 43 kDa | 0.99854 | 5 | 80% | P58269 | Dpf3 | Zinc finger | 17.7865 | 18.80425 | 17.87035 | 19.49178 | 17.77296 | 20.21977 | -1.69533 | -2.64963 | -0.74103 | 18.6576 | -4.23954 | -1.99203 | 0.004246 | 0.025026 | -1 | -1 |
| 24 | sp\|P28738 | KIF5C_MO | 109 kDa | 0.999682 | 27 | 74% | P28738 | Kif5c | Kinesin hea | 26.14704 | 27.13623 | 25.48745 | 27.13381 | 26.15793 | 27.62057 | -1.36606 | -2.13783 | -0.59429 | 26.61384 | -4.21291 | -2.00664 | 0.004263 | 0.025026 | -1 | -1 |
| 1027 | sp\|P04247 | MYG_MOU | 17 kDa | 0.999598 | 2 | 100% | P04247 | Mb | Myoglobin | 19.8928 | 18.99333 | 20.06902 | 18.93971 | 20.65781 | 18.32598 | 1.453536 | 0.631476 | 2.275597 | 19.47977 | 4.208661 | -2.01249 | 0.004288 | 0.02506 | 1 | 1 |
| 123 | sp\|Q6IFX2 | K1C42_MO | 50 kDa | 0.998174 | 14 | 29% | Q6IFX2 | Krt42 | Keratin, typ | 22.57409 | 24.7726 | 23.7388 | 25.0831 | 23.72628 | 25.27867 | -1.6984 | -2.65783 | -0.73897 | 24.19559 | -4.22453 | -2.01121 | 0.004324 | 0.025066 | -1 | -1 |
| 370 | sp\|Q64331 | MYO6_MO | 146 kDa | 0.998994 | 7 | 100% | Q64331 | Myo6 | Unconvent | 20.78207 | 22.9034 | 21.20147 | 23.22503 | 20.65398 | 21.85456 | -1.78182 | -2.78917 | -0.77447 | 21.77008 | -4.2212 | -2.01547 | 0.004341 | 0.025066 | -1 | -1 |
| 174 | sp\|P51881 | ADT2_MOU | 33 kDa | 0.999284 | 11 | 55% | P51881 | Slc25a5 | ADP/ATP tr | 24.52507 | 23.05248 | 23.82315 | 22.66325 | 24.18888 | 22.97536 | 1.282003 | 0.555055 | 2.008951 | 23.53803 | 4.197458 | -2.02661 | 0.004344 | 0.025066 | 1 | 1 |
| 483 | sp\|O08547 | SC22B_MO | 25 kDa | 0.999894 | 5 | 100% | O08547 | Sec22b | Vesicle-traf | 23.79215 | 22.57444 | 24.05767 | 22.73716 | 23.46862 | 22.24778 | 1.25302 | 0.538889 | 1.967152 | 23.1463 | 4.176194 | -2.05413 | 0.004458 | 0.025501 | 1 | 1 |
| 621 | sp\|Q9ER00 | STX12_MO | 31 kDa | 0.999496 | 4 | 100% | Q9ER00 | Stx12 | Syntaxin-12 | 23.70315 | 22.0989 | 23.50824 | 20.65139 | 23.50365 | 22.18932 | 1.925143 | 0.830354 | 3.019932 | 22.60911 | 4.196461 | -2.04716 | 0.004472 | 0.025501 | 1 | 1 |
| 720 | sp\|Q6ZPV2 | INO80_MO | 177 kDa | 0.998686 | 4 | 100% | Q6ZPV2 | Ino80 | Chromatin- | 18.01565 | 18.8961 | 17.12394 | 18.80979 | 17.46178 | 18.87278 | -1.32577 | -2.08196 | -0.56958 | 18.19667 | -4.17288 | -2.05842 | 0.004476 | 0.025501 | -1 | -1 |
| 394 | sp\|Q9CZT8 | RAB3B_MO | 25 kDa | 0.999744 | 6 | 33% | Q9CZT8 | Rab3b | Ras-related | 22.3992 | 21.13097 | 22.84088 | 20.58854 | 22.00084 | 21.22692 | 1.4315 | 0.611413 | 2.251587 | 21.69789 | 4.154831 | -2.08221 | 0.004578 | 0.025802 | 1 | 1 |
| 330 | sp\|Q9Z0H3 | SNF5_MOU | 44 kDa | 0.999526 | 7 | 100% | Q9Z0H3 | Smarcb1 | SWI/SNF-re | 24.42627 | 22.33792 | 23.4462 | 21.24587 | 24.44884 | 22.66562 | 2.023966 | 0.867448 | 3.180484 | 23.09512 | 4.176393 | -2.07294 | 0.004581 | 0.025802 | 1 | 1 |
| 543 | sp\|P21956 | MFGM_MO | 51 kDa | 0.999127 | 5 | 100% | P21956 | Mfge8 | Lactadherin | 20.43569 | 19.30064 | 21.21015 | 19.72779 | 20.78354 | 19.4737 | 1.309081 | 0.558832 | 2.059331 | 20.15525 | 4.152997 | -2.08424 | 0.004586 | 0.025802 | 1 | 1 |
| 918 | sp\|Q69ZQ2 | ISY1_MOUS | 33 kDa | 0.998423 | 3 | 100% | Q69ZQ2 | Isy1 | Pre-mRNA- | 22.5809 | 24.67487 | 22.99313 | 24.77975 | 21.84843 | 23.63831 | -1.89016 | -2.97268 | -0.80764 | 23.41923 | -4.1669 | -2.08516 | 0.004634 | 0.025929 | -1 | -1 |
| 596 | sp\|Q99K48 | NONO_MO | 55 kDa | 0.999738 | 4 | 75% | Q99K48 | Nono | Non-POU d | 20.89474 | 19.68457 | 21.03482 | 19.68744 | 20.48027 | 19.06889 | 1.322976 | 0.562791 | 2.083162 | 20.14179 | 4.142221 | -2.09826 | 0.004647 | 0.025929 | 1 | 1 |
| 675 | sp\|P05132 | KAPCA_MO | 41 kDa | 0.999091 | 4 | 50% | P05132 | Prkaca | cAMP-depe | 19.83985 | 21.77463 | 19.78772 | 21.18173 | 20.64757 | 22.04998 | -1.57706 | -2.4851 | -0.66903 | 20.88025 | -4.14048 | -2.11201 | 0.004735 | 0.026134 | -1 | -1 |
| 1277 | sp\|Q9D287 | SPF27_MO | 26 kDa | 0.99577 | 2 | 100% | Q9D287 | Bcas2 | Pre-mRNA- | 18.70835 | 13.9885 | 18.59317 | 17.27421 | 18.81194 | 15.04786 | 3.267629 | 1.385691 | 5.149567 | 17.07067 | 4.143602 | -2.1152 | 0.004767 | 0.026134 | 1 | 1 |
| 1278.1 | sp\|Q7TSF0 | DSG1C_MO | 101 kDa | 0.995714 | 2 | 100% | Q7TSF0 | Dsg1c | Desmoglein | 18.70835 | 13.9885 | 18.59317 | 17.27421 | 18.81194 | 15.04786 | 3.267629 | 1.385691 | 5.149567 | 17.07067 | 4.143602 | -2.1152 | 0.004767 | 0.026134 | 1 | 1 |
| 653 | sp\|P42225 | STAT1_MO | 87 kDa | 0.999283 | 4 | 100% | P42225 | Stat1 | Signal trans | 20.35865 | 19.5651 | 20.85073 | 19.40968 | 20.54185 | 19.40103 | 1.12514 | 0.475323 | 1.774957 | 20.02117 | 4.121132 | -2.12573 | 0.004768 | 0.026134 | 1 | 1 |
| 579 | sp\|P63087 | PP1G_MOU | 37 kDa | 0.997723 | 5 | 20% | P63087 | Ppp1cc | Serine/thre | 19.43867 | 21.97419 | 20.38385 | 21.23431 | 20.21514 | 22.78308 | -1.98464 | -3.12828 | -0.84101 | 21.00487 | -4.14139 | -2.11806 | 0.00478 | 0.026134 | -1 | -1 |
| 774 | tr\|A0A075 | A0A075B5 | 13 kDa | 0.995633 | 4 | 25% | A0A075B5 | Ighv7-3 | Immunoglo | 22.6625 | 21.39578 | 22.08317 | 21.19738 | 22.67557 | 21.21905 | 1.203013 | 0.50692 | 1.899106 | 21.87224 | 4.113428 | -2.13579 | 0.004813 | 0.026213 | 1 | 1 |
| 1152 | sp\|P01844 | LAC2_MOU | 11 kDa | 0.99843 | 2 | 50% | P01844 | Iglc2 | Ig lambda-2 | 20.71314 | 19.61536 | 20.98702 | 18.86239 | 20.24305 | 19.26072 | 1.401577 | 0.589401 | 2.213754 | 19.94695 | 4.107604 | -2.14373 | 0.00485 | 0.026309 | 1 | 1 |
| 744 | sp\|P54823 | DDX6_MOU | 54 kDa | 0.997909 | 4 | 100% | P54823 | Ddx6 | Probable AT | 22.85688 | 25.07803 | 24.25186 | 26.20248 | 24.08181 | 26.17252 | -2.08749 | -3.29646 | -0.87853 | 24.77393 | -4.12061 | -2.14492 | 0.004902 | 0.026481 | -1 | -1 |
| 628 | tr\|A0A140 | A0A140T8 | 13 kDa | 0.99946 | 4 | 50% | A0A140T8 | Igkv8-30 | Immunoglo | 21.70008 | 23.91505 | 22.5471 | 24.33119 | 21.3175 | 23.08488 | -1.92215 | -3.03787 | -0.80643 | 22.81597 | -4.11133 | -2.15694 | 0.004957 | 0.026675 | -1 | -1 |
| 507 | tr\|A0A075 | A0A075B5 | 13 kDa | 0.999496 | 5 | 20% | A0A075B5 | Ighv6-6 | Immunoglo | 23.10393 | 25.22683 | 23.57669 | 25.34276 | 22.26528 | 24.21038 | -1.94469 | -3.07589 | -0.81348 | 23.95431 | -4.1026 | -2.16827 | 0.00501 | 0.026853 | -1 | -1 |
| 468 | sp\|O70228 | ATP9A_MO | 119 kDa | 0.998178 | 6 | 67% | O70228 | Atp9a | Probable ph | 20.46668 | 19.03661 | 20.53157 | 19.15316 | 20.25761 | 19.58721 | 1.159627 | 0.481043 | 1.838212 | 19.83881 | 4.067386 | -2.19606 | 0.005094 | 0.027196 | 1 | 1 |
| 903.1 | sp\|Q6GSS7 | H2A2A_M | 14 kDa | 0.998649 | 3 | 33% | Q6GSS7 | Hist2h2aa2 | Histone H2 | 20.47444 | 22.10716 | 19.57708 | 21.36901 | 20.57363 | 21.82685 | -1.55929 | -2.47429 | -0.64429 | 20.98803 | -4.06359 | -2.21363 | 0.005214 | 0.027525 | -1 | -1 |
| 274 | sp\|P16014 | SCG1_MOU | 78 kDa | 0.999505 | 8 | 100% | P16014 | Chgb | Secretogra | 22.73764 | 23.72222 | 22.2271 | 23.6863 | 22.81412 | 23.85366 | -1.16111 | -1.84372 | -0.47849 | 23.17351 | -4.04853 | -2.22084 | 0.005214 | 0.027525 | -1 | -1 |
| 21 | sp\|O08810 | U5S1_MOU | 109 kDa | 0.999725 | 28 | 96% | O08810 | Eftud2 | 116 kDa U5 | 25.76133 | 24.08064 | 24.68752 | 22.86079 | 26.05939 | 23.60689 | 1.986638 | 0.821078 | 3.152199 | 24.50943 | 4.067567 | -2.21381 | 0.005229 | 0.027525 | 1 | 1 |
| 1234 | tr\|Q9JHF5 | Q9JHF5_M | 93 kDa | 0.997355 | 2 | 50% | Q9JHF5 | Tcirg1 | V-type prot | 23.00079 | 21.08417 | 23.83329 | 21.73598 | 22.65635 | 21.76223 | 1.636017 | 0.675878 | 2.596156 | 22.34547 | 4.066346 | -2.2154 | 0.005237 | 0.027525 | 1 | 1 |
| 196 | sp\|Q80WJ7 | LYRIC_MO | 64 kDa | 0.999538 | 10 | 100% | Q80WJ7 | Mtdh | Protein LYR | 24.06499 | 23.15208 | 24.28559 | 22.94461 | 24.34649 | 23.33512 | 1.088419 | 0.447462 | 1.729377 | 23.68815 | 4.041735 | -2.22978 | 0.005258 | 0.02753 | 1 | 1 |
| 776 | sp\|Q9D0M | DYL2_MOU | 10 kDa | 0.995494 | 4 | 50% | Q9D0M5 | Dynll2 | Dynein ligh | 22.52327 | 20.83074 | 22.79541 | 21.85832 | 22.67702 | 21.11447 | 1.39739 | 0.573796 | 2.220984 | 21.96654 | 4.038558 | -2.23429 | 0.005282 | 0.027546 | 1 | 1 |

| | | | | | | | | | | | | | | | | | | | | | | | | | |
|---|---|---|---|---|---|---|---|---|---|---|---|---|---|---|---|---|---|---|---|---|---|---|---|---|---|
| 612 | tr\|A0A140 | A0A140T8N | 13 kDa | 0.999605 | 4 | 25% | A0A140T8N | Igkv6-23 | Immunoglo | 20.91445 | 23.2108 | 22.11242 | 23.42372 | 20.73671 | 22.63175 | -1.83423 | -2.91343 | -0.75503 | 22.17164 | -4.05604 | -2.22884 | 0.005304 | 0.027554 | -1 | -1 |
| 963 | sp\|Q3UHK8 | TNR6A_MO | 203 kDa | 0.997696 | 3 | 100% | Q3UHK8 | Tnrc6a | Trinucleoti | 18.3557 | 20.51249 | 19.01337 | 20.02509 | NA | 20.34402 | -1.60933 | -2.52615 | -0.69251 | 19.65013 | -4.33477 | -2.02574 | 0.005344 | 0.027658 | -1 | -1 |
| 969 | sp\|Q08460 | KCMA1_MO | 134 kDa | 0.997601 | 3 | 100% | Q08460 | Kcnma1 | Calcium-ac | 17.678 | 15.88339 | 18.00248 | 16.26589 | 17.57318 | 16.89396 | 1.403468 | 0.57301 | 2.233927 | 17.04948 | 4.022597 | -2.25532 | 0.005387 | 0.027767 | 1 | 1 |
| 167 | sp\|Q91VR5 | DDX1_MOU | 82 kDa | 0.999436 | 11 | 100% | Q91VR5 | Ddx1 | ATP-depend | 23.30122 | 21.80544 | 22.70698 | 21.38053 | 22.84285 | 21.92862 | 1.245489 | 0.50796 | 1.983019 | 22.32761 | 4.019401 | -2.25922 | 0.005406 | 0.027767 | 1 | 1 |
| 826 | sp\|Q9CQN1 | TRAP1_MO | 80 kDa | 0.999366 | 3 | 100% | Q9CQN1 | Trap1 | Heat shock | 21.91922 | 18.94401 | 19.9797 | 19.23083 | 22.06275 | 18.69712 | 2.363236 | 0.965036 | 3.761437 | 20.13894 | 4.033558 | -2.2582 | 0.005452 | 0.027897 | 1 | 1 |
| 599 | tr\|A0A075 | A0A075B5I | 10 kDa | 0.999712 | 4 | 75% | A0A075B5I | Igkv9-124 | Immunoglo | 20.57192 | 22.66423 | 21.69615 | 23.13404 | 20.39761 | 22.24333 | -1.79197 | -2.85393 | -0.73002 | 21.78455 | -4.02696 | -2.26684 | 0.005496 | 0.028019 | -1 | -1 |
| 348 | sp\|P23249 | MOV10_MO | 114 kDa | 0.999342 | 7 | 100% | P23249 | Mov10 | Putative he | 21.27676 | 19.93984 | 21.55647 | 20.3593 | 20.98829 | 19.98992 | 1.177482 | 0.475892 | 1.879071 | 20.6851 | 3.994585 | -2.29202 | 0.005575 | 0.028171 | 1 | 1 |
| 1096 | sp\|O08807 | PRDX4_MO | 31 kDa | 0.999 | 2 | 50% | O08807 | Prdx4 | Peroxiredo | 17.9931 | 19.94334 | 18.22076 | 20.20226 | 18.03585 | 18.89354 | -1.59647 | -2.54548 | -0.64747 | 18.88147 | -4.01461 | -2.28301 | 0.00558 | 0.028171 | -1 | -1 |
| 116 | sp\|Q8CGC7 | SYEP_MOUS | 170 kDa | 0.999741 | 14 | 100% | Q8CGC7 | Eprs | Bifunctiona | 23.73363 | 25.04604 | 23.68682 | 25.21788 | 24.44025 | 25.48289 | -1.29537 | -2.06757 | -0.52317 | 24.60125 | -3.9927 | -2.29452 | 0.005589 | 0.028171 | -1 | -1 |
| 608 | sp\|Q99M28 | RNPS1_MO | 34 kDa | 0.999648 | 4 | 100% | Q99M28 | Rnps1 | RNA-bindin | 23.17282 | 21.85916 | 22.4934 | 21.54962 | 22.90669 | 21.61598 | 1.182714 | 0.472782 | 1.892646 | 22.26628 | 3.965189 | -2.33098 | 0.005784 | 0.029047 | 1 | 1 |
| 59 | sp\|Q3TKT4 | SMCA4_MO | 181 kDa | 0.999626 | 19 | 58% | Q3TKT4 | Smarca4 | Transcripti | 23.64992 | 21.92278 | 22.80156 | 22.20405 | 24.01612 | 21.35074 | 1.663339 | 0.66639 | 2.660288 | 22.65753 | 3.981607 | -2.32635 | 0.005812 | 0.029083 | 1 | 1 |
| 652 | sp\|Q9R226 | KHDR3_MO | 39 kDa | 0.99932 | 4 | 75% | Q9R226 | Khdrbs3 | KH domain | 23.03737 | 20.64425 | 21.33682 | 19.77296 | 22.90573 | 20.54607 | 2.105547 | 0.835703 | 3.375391 | 21.37387 | 3.956992 | -2.35877 | 0.005992 | 0.029873 | 1 | 1 |
| 919 | sp\|Q99KQ4 | NAMPT_MO | 55 kDa | 0.998411 | 3 | 100% | Q99KQ4 | Nampt | Nicotinami | 19.43594 | 21.40354 | 20.3241 | 21.20609 | 20.1842 | 21.21246 | -1.29262 | -2.07485 | -0.51038 | 20.62772 | -3.9331 | -2.37367 | 0.006021 | 0.029906 | -1 | -1 |
| 5 | sp\|Q8R0W | EPIPL_MOU | 723 kDa | 0.999889 | 53 | 96% | Q8R0W0 | Eppk1 | Epiplakin | 26.42704 | 23.51274 | 24.96372 | 24.08294 | 26.96435 | 24.25857 | 2.166955 | 0.852678 | 3.481232 | 25.03489 | 3.934718 | -2.3882 | 0.00616 | 0.030487 | 1 | 1 |
| 428 | sp\|P70333 | HNRH2_MO | 49 kDa | 0.999352 | 6 | 33% | P70333 | Hnrnph2 | Heterogene | 22.53427 | 20.68878 | 23.45805 | 20.71131 | 23.04984 | 22.05715 | 1.861641 | 0.730894 | 2.992388 | 22.08323 | 3.928993 | -2.39577 | 0.006204 | 0.030593 | 1 | 1 |
| 536 | sp\|Q922Q8 | LRC59_MO | 35 kDa | 0.999211 | 5 | 100% | Q922Q8 | Lrrc59 | Leucine-ric | 22.67913 | 21.96097 | 22.66218 | 21.11112 | 22.70258 | 21.02209 | 1.316565 | 0.514377 | 2.118754 | 22.02301 | 3.90631 | -2.40942 | 0.006227 | 0.030594 | 1 | 1 |
| 705 | sp\|P01786 | HVM17_MO | 12 kDa | 0.998848 | 4 | 25% | P01786 | NA | Ig heavy ch | 23.3802 | 25.11393 | 23.71698 | 25.62746 | 22.65071 | 24.32328 | -1.77226 | -2.8519 | -0.69261 | 24.13543 | -3.91739 | -2.41113 | 0.006295 | 0.030671 | -1 | -1 |
| 559 | sp\|Q80YR5 | SAFB2_MO | 112 kDa | 0.99883 | 5 | 100% | Q80YR5 | Safb2 | Scaffold att | 23.31818 | 21.69918 | 22.26191 | 21.08378 | 23.0486 | 21.47517 | 1.456853 | 0.568295 | 2.345411 | 22.1478 | 3.908624 | -2.41626 | 0.006302 | 0.030671 | 1 | 1 |
| 938 | sp\|Q8C0C7 | SYFA_MOUS | 58 kDa | 0.99806 | 3 | 100% | Q8C0C7 | Farsa | Phenylalan | 22.4362 | 23.05793 | 22.19186 | 23.36902 | 21.99061 | 23.60554 | -1.13794 | -1.83317 | -0.44271 | 22.77519 | -3.89574 | -2.42355 | 0.006311 | 0.030671 | -1 | -1 |
| 418 | sp\|P70372 | ELAV1_MO | 36 kDa | 0.999519 | 6 | 100% | P70372 | Elavl1 | ELAV-like p | 23.48833 | 22.39918 | 23.45239 | 22.14754 | 23.07889 | 22.30528 | 1.055871 | 0.408458 | 1.703284 | 22.81194 | 3.881775 | -2.44225 | 0.006423 | 0.031104 | 1 | 1 |
| 671 | sp\|Q922H4 | GMPPA_MO | 46 kDa | 0.999115 | 4 | 100% | Q922H4 | Gmppa | Mannose-1 | 21.88209 | 20.61914 | 21.71688 | 21.04773 | 21.85652 | 20.61797 | 1.056882 | 0.407984 | 1.70578 | 21.29006 | 3.876601 | -2.44919 | 0.006465 | 0.031134 | 1 | 1 |
| 400 | sp\|Q6ZWV3 | RL10_MOU | 25 kDa | 0.99967 | 6 | 100% | Q6ZWV3 | Rpl10 | 60S riboso | 23.70277 | 22.76863 | 24.73022 | 22.67266 | 23.89147 | 22.84036 | 1.347603 | 0.519987 | 2.175219 | 23.43435 | 3.875745 | -2.45063 | 0.006475 | 0.031134 | 1 | 1 |
| 148 | sp\|Q9WU2 | MTHR_MOU | 75 kDa | 0.999556 | 12 | 100% | Q9WU20 | Mthfr | Methylenet | 25.13436 | 23.70929 | 24.14165 | 22.34666 | 25.26627 | 21.77208 | 2.238083 | 0.863195 | 3.61297 | 23.72839 | 3.884718 | -2.45451 | 0.006557 | 0.031417 | 1 | 1 |
| 1090 | sp\|Q9CYR6 | AGM1_MO | 59 kDa | 0.999025 | 2 | 100% | Q9CYR6 | Pgm3 | Phosphoac | 21.62672 | 23.36316 | 21.61716 | 22.77109 | 21.69295 | 22.47944 | -1.22562 | -1.98167 | -0.46957 | 22.25842 | -3.85841 | -2.47361 | 0.006615 | 0.031586 | -1 | -1 |
| 338 | sp\|Q9CQV8 | 1433B_MO | 28 kDa | 0.999452 | 7 | 43% | Q9CQV8 | Ywhab | 14-3-3 prot | 20.69061 | 22.25122 | 20.94938 | 21.77248 | 20.85264 | 21.74035 | -1.09047 | -1.76435 | -0.4166 | 21.37611 | -3.85157 | -2.48279 | 0.006673 | 0.031749 | -1 | -1 |
| 632 | tr\|A0A075 | A0A075B5I | 13 kDa | 0.999431 | 4 | 25% | A0A075B5I | Ighv14-1 | Immunoglo | 20.65544 | 22.77383 | 22.0985 | 24.27004 | 21.14302 | 23.09677 | -2.08123 | -3.367 | -0.79545 | 22.3396 | -3.86282 | -2.48366 | 0.006739 | 0.031952 | -1 | -1 |
| 916.12 | sp\|Q8CGP6 | H2A1H_MO | 14 kDa | 0.998436 | 3 | 33% | Q8CGP6 | Hist1h2ah | Histone H2 | 20.6451 | 22.11734 | 20.2852 | 21.92949 | 21.20398 | 21.83807 | -1.25021 | -2.02814 | -0.47227 | 21.33653 | -3.82507 | -2.51847 | 0.006901 | 0.032606 | -1 | -1 |
| 186 | sp\|P62242 | RS8_MOUS | 24 kDa | 0.999717 | 10 | 100% | P62242 | Rps8 | 40S riboso | 26.12299 | 25.33812 | 26.36842 | 24.02069 | 25.86389 | 24.09249 | 1.634663 | 0.617928 | 2.651397 | 25.3011 | 3.836817 | -2.51838 | 0.006963 | 0.032784 | 1 | 1 |
| 1251.2 | sp\|Q9JKP5 | MBNL1_MO | 37 kDa | 0.996815 | 2 | 100% | Q9JKP5 | Mbnl1 | Muscleblin | 17.19217 | 18.18926 | 15.52598 | 18.93352 | 16.80841 | 18.13649 | -1.9109 | -3.10584 | -0.71596 | 17.46431 | -3.81631 | -2.54582 | 0.007145 | 0.033449 | -1 | -1 |
| 1017 | sp\|Q921T2 | TOIP1_MOU | 67 kDa | 0.999769 | 2 | 100% | Q921T2 | Tor1aip1 | Torsin-1A-i | 19.02487 | 20.18072 | 16.56918 | 20.30915 | 18.27851 | 20.41865 | -2.34532 | -3.81243 | -0.87821 | 19.13018 | -3.81497 | -2.54762 | 0.007157 | 0.033449 | -1 | -1 |
| 1100 | sp\|O88741 | GDAP1_MO | 41 kDa | 0.998964 | 2 | 100% | O88741 | Gdap1 | Ganglioside | 16.57023 | 17.09956 | 15.5506 | 17.64949 | 15.95305 | 17.3945 | -1.35656 | -2.20697 | -0.50614 | 16.7029 | -3.8019 | -2.55774 | 0.007195 | 0.033449 | -1 | -1 |
| 811 | sp\|Q62186 | SSRD_MOU | 19 kDa | 0.999444 | 3 | 100% | Q62186 | Ssr4 | Translocon | 23.63197 | 22.07744 | 23.67203 | 21.73563 | 23.51822 | 22.88704 | 1.374038 | 0.51262 | 2.235455 | 22.92039 | 3.802587 | -2.55813 | 0.007203 | 0.033449 | 1 | 1 |
| 638 | sp\|P01633 | KV5A1_MO | 16 kDa | 0.999406 | 4 | 50% | P01633 | Igk-V19-17 | Ig kappa ch | 22.52496 | 24.83582 | 24.19079 | 26.06789 | 22.61157 | 24.9458 | -2.17406 | -3.53917 | -0.80895 | 24.19614 | -3.80061 | -2.56687 | 0.007288 | 0.03373 | -1 | -1 |
| 637 | sp\|P01869 | IGH1M_MO | 43 kDa | 0.999409 | 4 | 100% | P01869 | Ighg1 | Ig gamma-1 | 25.16993 | 27.35553 | 25.73048 | 27.52137 | 24.17904 | 26.22533 | -2.00759 | -3.27374 | -0.74144 | 26.03028 | -3.7839 | -2.58932 | 0.007444 | 0.034334 | -1 | -1 |
| 378 | sp\|P62259 | 1433E_MO | 29 kDa | 0.998499 | 7 | 71% | P62259 | Ywhae | 14-3-3 prot | 21.92258 | 23.55076 | 22.10954 | 24.49489 | 21.71562 | 22.83284 | -1.71025 | -2.7925 | -0.62799 | 22.77104 | -3.77119 | -2.60643 | 0.007565 | 0.034773 | -1 | -1 |
| 1014 | sp\|P12961 | 7B2_MOUS | 24 kDa | 0.999876 | 2 | 100% | P12961 | Scg5 | Neuroendo | 18.97902 | 17.09026 | 18.34253 | 15.80426 | 19.94018 | 17.59953 | 2.255898 | 0.82642 | 3.685375 | 17.9593 | 3.766108 | -2.61326 | 0.007614 | 0.034782 | 1 | 1 |
| 740 | sp\|Q9Z108 | STAU1_MO | 54 kDa | 0.998308 | 4 | 100% | Q9Z108 | Stau1 | Double-stra | 20.90502 | 19.94529 | 21.08713 | 19.76023 | 20.53657 | 19.60812 | 1.071692 | 0.391062 | 1.752321 | 20.30706 | 3.747658 | -2.62327 | 0.007618 | 0.034782 | 1 | 1 |
| 88 | sp\|Q8CHC4 | SYNJ1_MOU | 173 kDa | 0.999829 | 16 | 100% | Q8CHC4 | Synj1 | Synaptojan | 23.53908 | 21.82992 | 22.44211 | 21.43725 | 23.00188 | 21.69697 | 1.339644 | 0.488383 | 2.190904 | 22.32454 | 3.750784 | -2.62676 | 0.007678 | 0.034939 | 1 | 1 |
| 159 | sp\|P62908 | RS3_MOUS | 27 kDa | 0.999714 | 11 | 100% | P62908 | Rps3 | 40S riboso | 26.7488 | 25.49396 | 26.55744 | 25.63845 | 26.84899 | 25.94945 | 1.024457 | 0.369736 | 1.679178 | 26.20618 | 3.724245 | -2.65514 | 0.007851 | 0.035516 | 1 | 1 |
| 1187 | sp\|Q8VHE0 | SEC63_MO | 88 kDa | 0.998093 | 2 | 100% | Q8VHE0 | Sec63 | Translocati | 19.45419 | 20.33196 | 19.3614 | 20.70331 | 19.32853 | 20.18215 | -1.02443 | -1.67926 | -0.36961 | 19.89359 | -3.72358 | -2.65604 | 0.007858 | 0.035516 | -1 | -1 |
| 56 | sp\|Q8CG16 | C1RA_MOU | 80 kDa | 0.999755 | 19 | 100% | Q8CG16 | C1ra | Compleme | 26.88471 | 27.83345 | 26.95398 | 28.23979 | 26.72597 | 27.65979 | -1.05612 | -1.73272 | -0.37952 | 27.38295 | -3.71519 | -2.66749 | 0.007943 | 0.035732 | -1 | -1 |
| 694 | tr\|A0A140 | A0A140T8N | 13 kDa | 0.998985 | 4 | 50% | A0A140T8N | Igkv6-15 | Immunoglo | 21.18154 | 23.5582 | 22.15962 | 23.61855 | 20.17522 | 22.6342 | -2.09819 | -3.44013 | -0.75626 | 22.22122 | -3.73134 | -2.66019 | 0.007958 | 0.035732 | -1 | -1 |
| 566 | sp\|Q4FK66 | PR38A_MO | 37 kDa | 0.998592 | 5 | 100% | Q4FK66 | Prpf38a | Pre-mRNA- | 21.39541 | 18.91557 | 20.33887 | 19.80857 | 21.57198 | 19.67731 | 1.634938 | 0.582709 | 2.687168 | 20.28462 | 3.708016 | -2.69176 | 0.008198 | 0.036481 | 1 | 1 |
| 406 | sp\|P14869 | RLA0_MOU | 34 kDa | 0.999608 | 6 | 100% | P14869 | Rplp0 | 60S acidic | 22.43835 | 21.56258 | 23.14436 | 21.87303 | 22.72625 | 21.48823 | 1.128374 | 0.39992 | 1.856827 | 22.20547 | 3.686818 | -2.70623 | 0.008239 | 0.036481 | 1 | 1 |
| 679.1 | sp\|P02301 | H3C_MOUS | 15 kDa | 0.999062 | 4 | 100% | P02301 | H3f3c | Histone H3 | 26.84963 | 25.79016 | 26.23383 | 25.38899 | 27.05354 | 24.9619 | 1.331984 | 0.472108 | 2.191861 | 26.04634 | 3.69281 | -2.70671 | 0.008286 | 0.036481 | 1 | 1 |
| 679.2 | sp\|P68433 | H31_MOUS | 15 kDa | 0.999062 | 4 | 100% | P68433 | Hist1h3i | Histone H3 | 26.84963 | 25.79016 | 26.23383 | 25.38899 | 27.05354 | 24.9619 | 1.331984 | 0.472108 | 2.191861 | 26.04634 | 3.69281 | -2.70671 | 0.008286 | 0.036481 | 1 | 1 |
| 679.3 | sp\|P84228 | H32_MOUS | 15 kDa | 0.999062 | 4 | 100% | P84228 | Hist2h3c2 | Histone H3 | 26.84963 | 25.79016 | 26.23383 | 25.38899 | 27.05354 | 24.9619 | 1.331984 | 0.472108 | 2.191861 | 26.04634 | 3.69281 | -2.70671 | 0.008286 | 0.036481 | 1 | 1 |
| 679.4 | sp\|P84244 | H33_MOUS | 15 kDa | 0.999062 | 4 | 100% | P84244 | H3f3b | Histone H3 | 26.84963 | 25.79016 | 26.23383 | 25.38899 | 27.05354 | 24.9619 | 1.331984 | 0.472108 | 2.191861 | 26.04634 | 3.69281 | -2.70671 | 0.008286 | 0.036481 | 1 | 1 |
| 208 | sp\|Q02257 | PLAK_MOU | 82 kDa | 0.999314 | 10 | 100% | Q02257 | Jup | Junction pl | 20.71432 | 22.87278 | 21.67919 | 22.86479 | 22.06726 | 23.68831 | -1.65504 | -2.7244 | -0.58568 | 22.31444 | -3.69347 | -2.71149 | 0.008352 | 0.036654 | -1 | -1 |
| 385 | sp\|P17427 | AP2A2_MO | 104 kDa | 0.997759 | 7 | 57% | P17427 | Ap2a2 | AP-2 compl | 21.497 | 19.59318 | 21.49084 | 19.78374 | 20.31035 | 19.56154 | 1.45324 | 0.512366 | 2.394115 | 20.37278 | 3.686009 | -2.72162 | 0.008433 | 0.03672 | 1 | 1 |
| 239 | sp\|Q9QWK | CD5L_MOU | 39 kDa | 0.999386 | 9 | 100% | Q9QWK4 | Cd5l | CD5 antige | 22.73384 | 24.67661 | 22.84612 | 24.54785 | 21.72721 | 23.40758 | -1.77496 | -2.92517 | -0.62474 | 23.3232 | -3.68264 | -2.7262 | 0.008469 | 0.03672 | -1 | -1 |
| 242 | sp\|Q9Z0H8 | CLIP2_MOU | 116 kDa | 0.999156 | 9 | 100% | Q9Z0H8 | Clip2 | CAP-Gly do | 21.51879 | 22.86324 | 20.35985 | 22.89537 | 22.03308 | 23.31907 | -1.72199 | -2.83807 | -0.60591 | 22.1649 | -3.68201 | -2.72705 | 0.008476 | 0.03672 | -1 | -1 |
| 304 | sp\|P19001 | K1C19_MO | 45 kDa | 0.994115 | 8 | 25% | P19001 | Krt19 | Keratin, typ | 21.29072 | 22.47524 | 22.15137 | 23.41253 | 21.4512 | 23.52047 | -1.50498 | -2.48047 | -0.5295 | 22.38359 | -3.68183 | -2.7273 | 0.008478 | 0.03672 | -1 | -1 |
| 793 | sp\|Q9CWZ3 | RBM8A_MO | 20 kDa | 0.999607 | 3 | 100% | Q9CWZ3 | Rbm8a | RNA-bindin | 24.88031 | 25.75978 | 24.53114 | 25.6741 | 24.92557 | 26.41514 | -1.17066 | -1.93145 | -0.40988 | 25.36434 | -3.66244 | -2.73961 | 0.008503 | 0.03672 | -1 | -1 |
| 677 | sp\|Q80WC | UBN2_MOU | 142 kDa | 0.999085 | 4 | 100% | Q80WC1 | Ubn2 | Ubinuclein | 19.17213 | 13.26706 | 17.38307 | 5.625675 | 21.08303 | 14.01973 | 8.241925 | 2.886949 | 13.5969 | 15.09178 | 3.673007 | -2.73929 | 0.008574 | 0.036911 | 1 | 1 |
| 195 | sp\|O54941 | SMCE1_MO | 47 kDa | 0.999567 | 10 | 100% | O54941 | Smarce1 | SWI/SNF-re | 24.90503 | 23.47545 | 23.76077 | 22.6846 | 25.01228 | 22.45604 | 1.687331 | 0.587594 | 2.787067 | 23.71569 | 3.661527 | -2.75491 | 0.008702 | 0.03734 | 1 | 1 |
| 486 | sp\|P61358 | RL27_MOU | 16 kDa | 0.999784 | 5 | 100% | P61358 | Rpl27 | 60S riboso | 24.21536 | 22.86925 | 24.38481 | 23.44313 | 24.006 | 23.12912 | 1.054886 | 0.36499 | 1.744781 | 23.67461 | 3.639342 | -2.77132 | 0.008762 | 0.037479 | 1 | 1 |
| 523 | sp\|O88712 | CTBP1_MO | 48 kDa | 0.99931 | 5 | 100% | O88712 | Ctbp1 | C-terminal- | 21.15973 | 22.87995 | 21.98129 | 22.82075 | 21.88324 | 22.78585 | -1.1541 | -1.90951 | -0.39869 | 22.2518 | -3.6363 | -2.7755 | 0.008796 | 0.037509 | -1 | -1 |
| 701 | sp\|Q60737 | CSK21_MO | 45 kDa | 0.998885 | 4 | 100% | Q60737 | Csnk2a1 | Casein kina | 21.30618 | 23.54571 | 20.38929 | 21.82659 | 21.0032 | 22.80043 | -1.82469 | -3.02029 | -0.62908 | 21.8119 | -3.6421 | -2.7814 | 0.008922 | 0.037925 | -1 | -1 |
| 439 | sp\|Q6NZF1 | ZC11A_MO | 86 kDa | 0.999213 | 6 | 100% | Q6NZF1 | Zc3h11a | Zinc finger | 24.71797 | 22.96025 | 23.5417 | 22.63279 | 25.27682 | 22.80947 | 1.711325 | 0.589156 | 2.833494 | 23.6565 | 3.639359 | -2.78514 | 0.008953 | 0.03794 | 1 | 1 |
| 1011 | sp\|Q9CQQ7 | AT5F1_MO | 29 kDa | 0.994424 | 3 | 100% | Q9CQQ7 | Atp5pb | ATP syntha | 22.55471 | 24.70787 | 22.84626 | 24.27014 | 21.49252 | 23.37996 | -1.82149 | -3.01905 | -0.62393 | 23.20858 | -3.62979 | -2.7982 | 0.009065 | 0.038024 | -1 | -1 |
| 391 | sp\|Q8R570 | SNP47_MO | 47 kDa | 0.999813 | 6 | 100% | Q8R570 | Snap47 | Synaptosor | 23.19089 | 24.21361 | 23.0016 | 24.28519 | 22.91097 | 23.72572 | -1.04035 | -1.7258 | -0.35491 | 23.55466 | -3.6125 | -2.80826 | 0.009073 | 0.038024 | -1 | -1 |
| 1047.1 | sp\|P01642 | KV5A9_MO | 13 kDa | 0.999377 | 2 | 100% | P01642 | Gm10881 | Ig kappa ch | 17.57796 | 19.35847 | 18.21446 | 19.68604 | NA | 19.06195 | -1.47261 | -2.41715 | -0.52807 | 18.77978 | -3.85033 | -2.59557 | 0.009085 | 0.038024 | -1 | -1 |
| 1047.2 | tr\|A0A140 | A0A140T8N | 13 kDa | 0.999377 | 2 | 100% | A0A140T8N | Igkv5-48 | Protein Igk | 17.57796 | 19.35847 | 18.21446 | 19.68604 | NA | 19.06195 | -1.47261 | -2.41715 | -0.52807 | 18.77978 | -3.85033 | -2.59557 | 0.009085 | 0.038024 | -1 | -1 |
| 508 | sp\|Q8K0E8 | FIBB_MOUS | 55 kDa | 0.999491 | 5 | 100% | Q8K0E8 | Fgb | Fibrinogen | 22.61871 | 20.39334 | 21.63383 | 21.20591 | 22.10401 | 20.67346 | 1.361281 | 0.463388 | 2.259174 | 21.43821 | 3.614243 | -2.81408 | 0.00917 | 0.038262 | 1 | 1 |
| 831 | sp\|Q0VBL3 | RBM15_MO | 106 kDa | 0.999348 | 3 | 100% | Q0VBL3 | Rbm15 | RNA-bindin | 22.4191 | 20.36229 | 20.53938 | 18.73121 | 21.96914 | 18.78609 | 2.34934 | 0.798737 | 3.899944 | 20.46787 | 3.615728 | -2.81743 | 0.009231 | 0.038395 | 1 | 1 |
| 1010 | tr\|A0A075 | A0A075B5Y | 13 kDa | 0.99458 | 3 | 33% | A0A075B5Y | Ighv1-37 | Immunoglo | 18.8997 | 22.01298 | 19.21375 | 19.69044 | 18.23639 | 21.6541 | -2.33589 | -3.88314 | -0.78865 | 19.95123 | -3.60283 | -2.83508 | 0.009386 | 0.038921 | -1 | -1 |
| 184 | sp\|Q8CFG8 | CS1B_MOU | 77 kDa | 0.999751 | 10 | 10% | Q8CFG8 | C1sb | Compleme | 21.15259 | 22.72476 | 21.61556 | 22.43485 | 20.79505 | 22.0608 | -1.21907 | -2.02888 | -0.40926 | 21.79727 | -3.58315 | -2.849 | 0.009431 | 0.038989 | -1 | -1 |
| 571 | tr\|A0A075 | A0A075B5T | 13 kDa | 0.998299 | 5 | 20% | A0A075B5T | Ighv6-3 | Immunoglo | 19.66521 | 21.26084 | 19.86692 | 21.61582 | 18.50265 | 20.43951 | -1.76046 | -2.92964 | -0.59129 | 20.22516 | -3.59333 | -2.8481 | 0.009502 | 0.039092 | -1 | -1 |
| 921 | sp\|Q9WVG | MS3L1_MO | 60 kDa | 0.998387 | 3 | 100% | Q9WVG9 | Msl3 | Male-specif | 18.51269 | 20.48919 | 19.83753 | 22.12256 | 19.09067 | 20.74367 | -1.97151 | -3.28119 | -0.66183 | 20.13272 | -3.59239 | -2.84938 | 0.009513 | 0.039092 | -1 | -1 |
| 1271 | sp\|P55937 | GOGA3_MO | 167 kDa | 0.996086 | 2 | 100% | P55937 | Golga3 | Golgin subf | 19.3429 | 20.73359 | 19.63075 | 20.40843 | 19.19605 | 20.16801 | -1.04677 | -1.7454 | -0.34815 | 19.91329 | -3.56625 | -2.87213 | 0.009638 | 0.039485 | -1 | -1 |
| 198 | sp\|P46096 | SYT1_MOU | 47 kDa | 0.999481 | 10 | 80% | P46096 | Syt1 | Synaptotag | 24.23333 | 23.41788 | 24.31347 | 23.2686 | 24.4028 | 23.53072 | 0.9108 | 0.301588 | 1.520012 | 23.86113 | 3.558406 | -2.88299 | 0.009738 | 0.039772 | 0 | 0 |
| 532 | sp\|Q6NVF9 | CPSF6_MO | 59 kDa | 0.999221 | 5 | 100% | Q6NVF9 | Cpsf6 | Cleavage ar | 23.60424 | 22.58169 | 23.01529 | 21.44957 | 23.17718 | 21.52069 | 1.414919 | 0.466095 | 2.363744 | 22.55811 | 3.558742 | -2.8956 | 0.009938 | 0.040469 | 1 | 1 |
| 112 | sp\|P06880 | SOMA_MOU | 25 kDa | 0.999893 | 14 | 100% | P06880 | Gh1 | Somatotro | 30.46798 | 29.19401 | 31.5654 | 30.12217 | 31.10104 | 29.50956 | 1.436224 | 0.468745 | 2.403703 | 30.32669 | 3.542676 | -2.91772 | 0.010148 | 0.040629 | 1 | 1 |
| 780 | sp\|P62830 | RL23_MOU | 15 kDa | 0.999855 | 3 | 100% | P62830 | Rpl23 | 60S riboso | 22.57372 | 24.72081 | 22.8465 | 24.40768 | 21.26138 | 23.37754 | -1.94148 | -3.25058 | -0.63238 | 23.19794 | -3.53924 | -2.92246 | 0.010194 | 0.040629 | -1 | -1 |
| 8 | sp\|Q6P4T2 | U520_MOU | 245 kDa | 0.999706 | 46 | 100% | Q6P4T2 | Snrnp200 | U5 small nu | 26.14711 | 24.35526 | 25.19469 | 24.83136 | 26.5794 | 24.24679 | 1.495931 | 0.486917 | 2.504946 | 25.22577 | 3.538059 | -2.92409 | 0.010209 | 0.040629 | 1 | 1 |
| 542.1 | sp\|P01787 | HVM18_MO | 14 kDa | 0.999129 | 5 | 60% | P01787 | NA | Ig heavy ch | 22.6187 | 24.57004 | 22.52038 | 24.12021 | 21.36085 | 23.18444 | -1.79158 | -3.00022 | -0.58295 | 23.06244 | -3.53747 | -2.92489 | 0.010217 | 0.040629 | -1 | -1 |
| 542.2 | sp\|P01788 | HVM19_MO | 14 kDa | 0.999129 | 5 | 60% | P01788 | NA | Ig heavy ch | 22.6187 | 24.57004 | 22.52038 | 24.12021 | 21.36085 | 23.18444 | -1.79158 | -3.00022 | -0.58295 | 23.06244 | -3.53747 | -2.92489 | 0.010217 | 0.040629 | -1 | -1 |
| 542.3 | sp\|P01790 | HVM21_MO | 14 kDa | 0.999129 | 5 | 60% | P01790 | NA | Ig heavy ch | 22.6187 | 24.57004 | 22.52038 | 24.12021 | 21.36085 | 23.18444 | -1.79158 | -3.00022 | -0.58295 | 23.06244 | -3.53747 | -2.92489 | 0.010217 | 0.040629 | -1 | -1 |
| 542.4 | sp\|P01791 | HVM22_MO | 14 kDa | 0.999129 | 5 | 60% | P01791 | NA | Ig heavy ch | 22.6187 | 24.57004 | 22.52038 | 24.12021 | 21.36085 | 23.18444 | -1.79158 | -3.00022 | -0.58295 | 23.06244 | -3.53747 | -2.92489 | 0.010217 | 0.040629 | -1 | -1 |
| 542.5 | sp\|P01794 | HVM25_MO | 14 kDa | 0.999129 | 5 | 60% | P01794 | NA | Ig heavy ch | 22.6187 | 24.57004 | 22.52038 | 24.12021 | 21.36085 | 23.18444 | -1.79158 | -3.00022 | -0.58295 | 23.06244 | -3.53747 | -2.92489 | 0.010217 | 0.040629 | -1 | -1 |
| 800 | sp\|Q9CQR5 | ZMAT5_MO | 20 kDa | 0.999518 | 3 | 100% | Q9CQR5 | Zmat5 | Zinc finger | 22.62015 | 24.90333 | 23.50593 | 25.25033 | 21.88105 | 23.76511 | -1.97054 | -3.30254 | -0.63855 | 23.65432 | -3.53048 | -2.93453 | 0.010311 | 0.040881 | -1 | -1 |
| 19 | sp\|Q8BTI8 | SRRM2_MO | 295 kDa | 0.999729 | 29 | 100% | Q8BTI8 | Srrm2 | Serine/argi | 25.50006 | 24.34192 | 25.24684 | 24.57461 | 26.15066 | 24.54117 | 1.146622 | 0.369694 | 1.923551 | 25.05921 | 3.512693 | -2.94644 | 0.010341 | 0.040881 | 1 | 1 |
| 1178 | sp\|Q93092 | TALDO_MO | 37 kDa | 0.998161 | 2 | 100% | Q93092 | Taldo1 | Transaldola | 22.04436 | 21.23042 | 22.13229 | 20.93387 | 21.7499 | 20.87381 | 0.962817 | 0.30772 | 1.617914 | 21.49411 | 3.498153 | -2.96668 | 0.010541 | 0.041552 | 0 | 0 |
| 343 | sp\|P68254 | 1433T_MO | 28 kDa | 0.99937 | 7 | 71% | P68254 | Ywhaq | 14-3-3 prot | 22.78802 | 21.55871 | 22.96714 | 21.86973 | 22.28267 | 21.25707 | 1.11744 | 0.350156 | 1.884724 | 22.12056 | 3.46632 | -3.01109 | 0.010994 | 0.043027 | 1 | 1 |
| 324 | tr\|A0A0B4 | A0A0B4J1H | 13 kDa | 0.999571 | 7 | 100% | A0A0B4J1H | Igkv1-135 | Immunoglo | 23.71496 | 25.78531 | 24.76285 | 26.71331 | 23.1627 | 25.08425 | -1.98079 | -3.33927 | -0.6223 | 24.87056 | -3.47964 | -3.00484 | 0.01102 | 0.043027 | -1 | -1 |
| 419 | sp\|P01801 | HVM32_MO | 13 kDa | 0.999484 | 6 | 17% | P01801 | NA | Ig heavy ch | 20.15813 | 21.63836 | 20.46698 | 22.09655 | 19.21493 | 20.92708 | -1.60732 | -2.70982 | -0.50482 | 20.75034 | -3.47917 | -3.00549 | 0.011027 | 0.043027 | -1 | -1 |
| 459 | sp\|Q569Z5 | DDX46_MO | 117 kDa | 0.998922 | 6 | 100% | Q569Z5 | Ddx46 | Probable A | 20.75107 | 21.78378 | 20.94048 | 21.49856 | 20.50994 | 21.75951 | -0.94679 | -1.59763 | -0.29595 | 21.20722 | -3.46243 | -3.01652 | 0.011051 | 0.043027 | 0 | 0 |
| 832.5 | tr\|A0A140 | A0A140T8C | 13 kDa | 0.999312 | 3 | 33% | A0A140T8C | Igkv3-4 | Immunoglo | 23.40676 | 22.58069 | 23.36312 | 21.70752 | 23.16474 | 22.31016 | 1.112086 | 0.347275 | 1.876898 | 22.7555 | 3.460868 | -3.0187 | 0.011074 | 0.043027 | 1 | 1 |
| 657 | sp\|Q9QUI0 | RHOA_MOU | 22 kDa | 0.999264 | 4 | 25% | Q9QUI0 | Rhoa | Transformi | 19.77692 | 17.76794 | 20.3614 | 18.64351 | 19.62169 | 19.0265 | 1.440683 | 0.448794 | 2.432572 | 19.19966 | 3.46622 | -3.02345 | 0.011216 | 0.043453 | 1 | 1 |
| 874 | sp\|Q91WJ8 | FUBP1_MO | 69 kDa | 0.998969 | 3 | 67% | Q91WJ8 | Fubp1 | Far upstrea | 19.29235 | 21.24632 | 20.78671 | 22.4224 | 19.608 | 21.36592 | -1.78252 | -3.01449 | -0.55055 | 20.78695 | -3.45291 | -3.04193 | 0.011414 | 0.044056 | -1 | -1 |
| 402 | sp\|G5E829 | AT2B1_MO | 135 kDa | 0.999659 | 6 | 100% | G5E829 | Atp2b1 | Plasma mer | 21.27672 | 24.26736 | 21.99073 | 23.04389 | 22.80853 | 24.06834 | -1.76787 | -2.99024 | -0.5455 | 22.90926 | -3.45142 | -3.04401 | 0.011436 | 0.044056 | -1 | -1 |
| 271 | sp\|Q9CXY6 | ILF2_MOUS | 43 kDa | 0.999564 | 8 | 100% | Q9CXY6 | Ilf2 | Interleukin | 22.558 | 21.56879 | 22.00128 | 21.26223 | 22.60948 | 20.60511 | 1.24421 | 0.382711 | 2.105708 | 21.76748 | 3.442967 | -3.05101 | 0.011475 | 0.04408 | 1 | 1 |
| 916.11 | sp\|Q8CGP5 | H2A1F_MO | 14 kDa | 0.998436 | 3 | 33% | Q8CGP5 | Hist1h2af | Histone H2 | 21.70109 | 19.53103 | 20.35261 | 19.26683 | 20.57721 | 19.59098 | 1.414022 | 0.431684 | 2.396359 | 20.16996 | 3.435153 | -3.06662 | 0.011684 | 0.044755 | 1 | 1 |
| 1290 | sp\|Q80XN0 | BDH_MOUS | 38 kDa | 0.995005 | 2 | 100% | Q80XN0 | Bdh1 | D-beta-hyd | 15.86316 | 18.82183 | 16.37755 | 17.56597 | 16.71688 | 17.37783 | -1.60269 | -2.71967 | -0.4857 | 17.12054 | -3.42415 | -3.08195 | 0.011855 | 0.045282 | -1 | -1 |
| 1009 | sp\|Q9EPN1 | NBEA_MOU | 327 kDa | 0.994717 | 3 | 33% | Q9EPN1 | Nbea | Neurobeac | 18.7965 | 22.44869 | 19.08408 | 20.13113 | 19.50881 | 23.80586 | -2.99877 | -5.09631 | -0.90123 | 20.62918 | -3.4118 | -3.09916 | 0.01205 | 0.045898 | -1 | -1 |
| 1062 | sp\|Q0P5V2 | SOBP_MOU | 92 kDa | 0.999243 | 2 | 100% | Q0P5V2 | Sobp | Sine oculis- | 18.26344 | 19.82591 | 18.13364 | 18.75869 | 16.92038 | 20.4005 | -1.88921 | -3.21155 | -0.56687 | 18.7171 | -3.40948 | -3.10239 | 0.012087 | 0.04591 | -1 | -1 |
| 1210 | sp\|Q8R326 | PSPC1_MO | 59 kDa | 0.99779 | 2 | 100% | Q8R326 | Pspc1 | Paraspeckle | 19.0285 | 17.98653 | 18.45406 | 16.52011 | 18.40393 | 16.43615 | 1.647899 | 0.491239 | 2.80456 | 17.80488 | 3.399973 | -3.11566 | 0.01224 | 0.046361 | 1 | 1 |
| 1286 | sp\|P42232 | STA5B_MO | 90 kDa | 0.995294 | 2 | 100% | P42232 | Stat5b | Signal trans | 18.58149 | 19.83637 | 17.8418 | 19.0329 | 18.54645 | 19.94087 | -1.28014 | -2.18298 | -0.37729 | 18.96331 | -3.38014 | -3.13885 | 0.012472 | 0.047066 | -1 | -1 |
| 786 | tr\|A0A075 | A0A075B67 | 11 kDa | 0.999674 | 3 | 33% | A0A075B67 | Ighv1-78 | Immunoglo | 21.42798 | 18.9608 | 20.20741 | 20.11767 | 21.60376 | 18.59672 | 1.854654 | 0.546871 | 3.162437 | 20.15239 | 3.384371 | -3.13746 | 0.012495 | 0.047066 | 1 | 1 |
| 74 | sp\|Q9Z1R2 | BAG6_MOU | 121 kDa | 0.999644 | 17 | 100% | Q9Z1R2 | Bag6 | Large prolin | 24.82119 | 26.31045 | 25.63808 | 25.99043 | 24.92573 | 26.42931 | -1.11506 | -1.90333 | -0.32679 | 25.68587 | -3.36687 | -3.15063 | 0.012552 | 0.047147 | -1 | -1 |
| 382 | sp\|Q99PU8 | DHX30_MO | 137 kDa | 0.99821 | 7 | 100% | Q99PU8 | Dhx30 | ATP-depend | 20.1292 | 18.1591 | 19.60263 | 17.52874 | 19.21206 | 18.85229 | 1.467917 | 0.430368 | 2.505465 | 18.914 | 3.376321 | -3.14872 | 0.01263 | 0.047309 | 1 | 1 |
| 1058 | sp\|Q5SUR0 | PUR4_MOU | 145 kDa | 0.999277 | 2 | 100% | Q5SUR0 | Pfas | Phosphorib | 18.74609 | 17.12656 | 18.66624 | 12.98109 | 19.0687 | 16.2118 | 3.387192 | 0.985407 | 5.788977 | 17.13341 | 3.365552 | -3.16379 | 0.012811 | 0.047575 | 1 | 1 |
| 319 | sp\|P63011 | RAB3A_MO | 25 kDa | 0.999637 | 7 | 71% | P63011 | Rab3a | Ras-related | 24.38117 | 23.10715 | 24.12305 | 23.33827 | 23.86707 | 23.14006 | 0.928602 | 0.269147 | 1.588057 | 23.65946 | 3.351545 | -3.17223 | 0.012812 | 0.047575 | 0 | 0 |
| 832.1 | sp\|P01665 | KV3AD_MO | 12 kDa | 0.999312 | 3 | 33% | P01665 | NA | Ig kappa ch | 24.16156 | 26.14511 | 25.44105 | 27.12187 | 23.31243 | 25.84295 | -2.06496 | -3.53051 | -0.59942 | 25.3375 | -3.36251 | -3.16805 | 0.012863 | 0.047575 | -1 | -1 |
| 832.2 | sp\|P01666 | KV3AE_MO | 12 kDa | 0.999312 | 3 | 33% | P01666 | NA | Ig kappa ch | 24.16156 | 26.14511 | 25.44105 | 27.12187 | 23.31243 | 25.84295 | -2.06496 | -3.53051 | -0.59942 | 25.3375 | -3.36251 | -3.16805 | 0.012863 | 0.047575 | -1 | -1 |
| 206 | tr\|E9Q1Z0 | E9Q1Z0_M | 58 kDa | 0.99933 | 10 | 10% | E9Q1Z0 | Krt90 | Keratin 90 | 25.17991 | 26.3187 | 25.04602 | 26.05911 | 25.64295 | 26.84973 | -1.11955 | -1.91582 | -0.32329 | 25.8494 | -3.34648 | -3.17938 | 0.012899 | 0.047575 | -1 | -1 |
| 785 | sp\|Q6PDM | MSL1_MOU | 67 kDa | 0.999675 | 3 | 100% | Q6PDM1 | Msl1 | Male-specif | 19.42529 | 20.31994 | 19.49852 | 20.25393 | 18.72494 | 21.48599 | -1.47037 | -2.51479 | -0.42595 | 19.95144 | -3.35972 | -3.17195 | 0.012911 | 0.047575 | -1 | -1 |
| 1230 | sp\|Q9Z1T1 | AP3B1_MO | 123 kDa | 0.997522 | 2 | 100% | Q9Z1T1 | Ap3b1 | AP-3 compl | 18.18484 | 18.58027 | 17.71707 | 20.40209 | 17.76675 | 21.07671 | -2.13014 | -3.6484 | -0.61188 | 18.95462 | -3.34822 | -3.18808 | 0.01311 | 0.047997 | -1 | -1 |
| 379 | sp\|Q9QZQ8 | H2AY_MOU | 40 kDa | 0.998402 | 7 | 100% | Q9QZQ8 | H2afy | Core histon | 23.14974 | 22.48891 | 22.8055 | 21.74509 | 23.35294 | 21.52854 | 1.18188 | 0.337856 | 2.025905 | 22.51178 | 3.335293 | -3.19822 | 0.013161 | 0.047997 | 1 | 1 |
| 410 | sp\|P14115 | RL27A_MO | 17 kDa | 0.999587 | 6 | 100% | P14115 | Rpl27a | 60S riboso | 25.0773 | 24.35798 | 25.51709 | 24.37646 | 25.06241 | 24.16433 | 0.91934 | 0.260875 | 1.577805 | 24.75926 | 3.323105 | -3.21241 | 0.013311 | 0.047997 | 0 | 0 |
| 627.1 | sp\|P10853 | H2B1F_MO | 14 kDa | 0.999468 | 4 | 100% | P10853 | Hist1h2bn | Histone H2 | 27.60808 | 26.65783 | 26.88069 | 25.35859 | 27.35263 | 25.46878 | 1.452069 | 0.411749 | 2.492388 | 26.55443 | 3.330972 | -3.21228 | 0.013415 | 0.047997 | 1 | 1 |
| 627.2 | sp\|P10854 | H2B1M_MO | 14 kDa | 0.999468 | 4 | 100% | P10854 | Hist1h2bm | Histone H2 | 27.60808 | 26.65783 | 26.88069 | 25.35859 | 27.35263 | 25.46878 | 1.452069 | 0.411749 | 2.492388 | 26.55443 | 3.330972 | -3.21228 | 0.013415 | 0.047997 | 1 | 1 |
| 627.3 | sp\|Q6ZWY9 | H2B1C_MO | 14 kDa | 0.999468 | 4 | 100% | Q6ZWY9 | Hist1h2bg | Histone H2 | 27.60808 | 26.65783 | 26.88069 | 25.35859 | 27.35263 | 25.46878 | 1.452069 | 0.411749 | 2.492388 | 26.55443 | 3.330972 | -3.21228 | 0.013415 | 0.047997 | 1 | 1 |
| 627.4 | sp\|Q8CGP1 | H2B1K_MO | 14 kDa | 0.999468 | 4 | 100% | Q8CGP1 | Hist1h2bk | Histone H2 | 27.60808 | 26.65783 | 26.88069 | 25.35859 | 27.35263 | 25.46878 | 1.452069 | 0.411749 | 2.492388 | 26.55443 | 3.330972 | -3.21228 | 0.013415 | 0.047997 | 1 | 1 |
| 627.5 | sp\|Q8CGP2 | H2B1P_MO | 14 kDa | 0.999468 | 4 | 100% | Q8CGP2 | Hist1h2bp | Histone H2 | 27.60808 | 26.65783 | 26.88069 | 25.35859 | 27.35263 | 25.46878 | 1.452069 | 0.411749 | 2.492388 | 26.55443 | 3.330972 | -3.21228 | 0.013415 | 0.047997 | 1 | 1 |
| 627.6 | sp\|Q64475 | H2B1B_MO | 14 kDa | 0.999468 | 4 | 100% | Q64475 | Hist1h2bb | Histone H2 | 27.60808 | 26.65783 | 26.88069 | 25.35859 | 27.35263 | 25.46878 | 1.452069 | 0.411749 | 2.492388 | 26.55443 | 3.330972 | -3.21228 | 0.013415 | 0.047997 | 1 | 1 |
| 627.7 | sp\|Q64478 | H2B1H_MO | 14 kDa | 0.999468 | 4 | 100% | Q64478 | Hist1h2bh | Histone H2 | 27.60808 | 26.65783 | 26.88069 | 25.35859 | 27.35263 | 25.46878 | 1.452069 | 0.411749 | 2.492388 | 26.55443 | 3.330972 | -3.21228 | 0.013415 | 0.047997 | 1 | 1 |
| 627.8 | sp\|Q64525 | H2B2B_MO | 14 kDa | 0.999468 | 4 | 100% | Q64525 | Hist2h2bb | Histone H2 | 27.60808 | 26.65783 | 26.88069 | 25.35859 | 27.35263 | 25.46878 | 1.452069 | 0.411749 | 2.492388 | 26.55443 | 3.330972 | -3.21228 | 0.013415 | 0.047997 | 1 | 1 |
| 782 | sp\|Q9JKK1 | STX6_MOU | 29 kDa | 0.999725 | 3 | 100% | Q9JKK1 | Stx6 | Syntaxin-6 | 21.04916 | 19.62749 | 19.84004 | 19.1177 | 21.38642 | 18.66022 | 1.623402 | 0.459569 | 2.787235 | 19.94684 | 3.328788 | -3.21535 | 0.013454 | 0.04801 | 1 | 1 |
| 1063.1 | tr\|A0A0A6 | A0A0A6YX | 42 kDa | 0.999242 | 2 | 100% | A0A0A6YX | Igha | Immunoglo | 21.34396 | 22.84424 | 22.27622 | 22.72817 | 20.9602 | 22.90266 | -1.29823 | -2.22974 | -0.36672 | 22.17591 | -3.32593 | -3.21936 | 0.013506 | 0.048067 | -1 | -1 |
| 66 | sp\|Q62261 | SPTB2_MO | 274 kDa | 0.999538 | 18 | 94% | Q62261 | Sptbn1 | Spectrin be | 23.05836 | 24.39588 | 23.34265 | 24.07382 | 23.25027 | 23.91527 | -0.91123 | -1.57009 | -0.25237 | 23.67271 | -3.29182 | -3.25671 | 0.013884 | 0.04917 | 0 | 0 |
| 1291 | sp\|Q3TDD9 | PPR21_MO | 88 kDa | 0.994795 | 2 | 100% | Q3TDD9 | Ppp1r21 | Protein pho | 19.72781 | 21.30293 | 20.62817 | 22.03197 | 20.77996 | 21.64606 | -1.28168 | -2.20723 | -0.35613 | 21.01948 | -3.30404 | -3.24936 | 0.013888 | 0.04917 | -1 | -1 |
| 220 | sp\|Q9R0H5 | K2C71_MO | 57 kDa | 0.99783 | 10 | 30% | Q9R0H5 | Krt71 | Keratin, typ | 20.71627 | 26.99334 | 22.32913 | 25.93382 | 25.47407 | 26.84644 | -3.75138 | -6.46264 | -1.04012 | 24.71551 | -3.30195 | -3.25309 | 0.013945 | 0.049244 | -1 | -1 |
| 982 | sp\|Q8BJM7 | TYW1_MOU | 82 kDa | 0.99719 | 3 | 100% | Q8BJM7 | Tyw1 | S-adenosyl- | 18.54003 | 20.42801 | 19.3526 | 19.63303 | 18.22847 | 19.99479 | -1.31158 | -2.26077 | -0.36239 | 19.36282 | -3.29757 | -3.25925 | 0.014027 | 0.049405 | -1 | -1 |
| 771 | sp\|P21126 | UBL4A_MO | 18 kDa | 0.996743 | 4 | 100% | P21126 | Ubl4a | Ubiquitin-l | 19.11274 | 20.47747 | 18.74828 | 20.26007 | 19.71726 | 20.14981 | -1.10302 | -1.90404 | -0.30201 | 19.74427 | -3.27752 | -3.27699 | 0.014155 | 0.049718 | -1 | -1 |
| 1064 | sp\|Q31125 | S39A7_MO | 51 kDa | 0.99924 | 2 | 100% | Q31125 | Slc39a7 | Zinc transp | 23.65166 | 23.03033 | 23.93482 | 22.70736 | 23.55237 | 22.70352 | 0.899217 | 0.245815 | 1.552619 | 23.26334 | 3.275553 | -3.27979 | 0.014193 | 0.049718 | 0 | 0 |
| 457 | sp\|P97762 | RP9_MOUS | 25 kDa | 0.998954 | 6 | 100% | P97762 | rp9 | Retinitis pi | 19.1707 | 20.75125 | 20.26302 | 21.88216 | 19.48914 | 20.76678 | -1.49244 | -2.57597 | -0.40891 | 20.38717 | -3.28706 | -3.27407 | 0.014226 | 0.049718 | -1 | -1 |
| 962 | sp\|P20357 | MTAP2_MO | 199 kDa | 0.997708 | 3 | 100% | P20357 | Map2 | Microtubul | 19.9232 | 19.08077 | 19.92721 | 17.82748 | 19.56382 | 18.68972 | 1.272088 | 0.347611 | 2.196565 | 19.1687 | 3.283124 | -3.27884 | 0.014283 | 0.049787 | 1 | 1 |
| 575 | sp\|P23116 | EIF3A_MOU | 162 kDa | 0.998125 | 5 | 100% | P23116 | Eif3a | Eukaryotic | 21.03761 | 22.24267 | 21.56231 | 23.9039 | 20.93486 | 22.34992 | -1.6539 | -2.85715 | -0.45065 | 22.00521 | -3.28022 | -3.28371 | 0.014357 | 0.049919 | -1 | -1 |
| 216 | sp\|Q91YR7 | PRP6_MOU | 107 kDa | 0.998848 | 10 | 100% | Q91YR7 | Prpf6 | Pre-mRNA- | 22.78952 | 23.57737 | 22.44745 | 23.3747 | 22.79357 | 23.79134 | -0.90429 | -1.56473 | -0.24385 | 23.12899 | -3.25891 | -3.30343 | 0.014516 | 0.050316 | 0 | 0 |
| 650 | sp\|Q9Z1Q9 | SYVC_MOU | 140 kDa | 0.999336 | 4 | 100% | Q9Z1Q9 | Vars | Valine--tRN | 21.27092 | 22.27429 | 21.92173 | 22.78289 | 21.34803 | 22.54732 | -1.02127 | -1.7675 | -0.27504 | 22.0242 | -3.2574 | -3.30558 | 0.014546 | 0.050316 | -1 | -1 |

| | | | | | | | | | | | | | | | | | | | | | | | | | |
|---|---|---|---|---|---|---|---|---|---|---|---|---|---|---|---|---|---|---|---|---|---|---|---|---|---|
| 1144 | sp\|Q61464 | ZN638_MO | 218 kDa | 0.998506 | 2 | 100% | Q61464 | Znf638 | Zinc finger | 18.73794 | 20.15227 | 17.33851 | 19.61122 | 19.09184 | 20.96889 | -1.8547 | -3.21253 | -0.49687 | 19.31678 | -3.25971 | -3.31265 | 0.014758 | 0.05092 | -1 | -1 |
| 1003 | sp\|Q8BX10 | PGAM5_M | 32 kDa | 0.995432 | 3 | 100% | Q8BX10 | Pgam5 | Serine/thre | 20.61479 | 19.25946 | 21.03657 | 20.16825 | 20.80334 | 19.797 | 1.076663 | 0.285382 | 1.867945 | 20.2799 | 3.238543 | -3.3324 | 0.014922 | 0.051357 | 1 | 1 |
| 1119.2 | sp\|C0HK80 | ARXS2_MO | 20 kDa | 0.998757 | 2 | 100% | C0HK80 | Arxes2 | Adipocyte- | 22.44177 | 21.28251 | 22.2516 | 21.36534 | 21.8199 | 20.95819 | 0.969081 | 0.255958 | 1.682204 | 21.68655 | 3.234421 | -3.33827 | 0.015006 | 0.051514 | 0 | 0 |
| 898 | sp\|P02468 | LAMC1_M | 177 kDa | 0.998706 | 3 | 100% | P02468 | Lamc1 | Laminin su | 17.36052 | 21.12654 | 18.73089 | 18.83599 | 18.10737 | 21.03991 | -2.26789 | -3.93636 | -0.59941 | 19.2002 | -3.24379 | -3.33516 | 0.015077 | 0.051539 | -1 | -1 |
| 218 | sp\|Q3UV17 | K22O_MOU | 63 kDa | 0.998045 | 10 | 10% | Q3UV17 | Krt76 | Keratin, typ | 24.17855 | 27.2646 | 26.69078 | 28.41119 | 26.21784 | 28.21609 | -2.26823 | -3.93761 | -0.59886 | 26.82984 | -3.24254 | -3.33692 | 0.015103 | 0.051539 | -1 | -1 |
| 613 | sp\|P62918 | RL8_MOUS | 28 kDa | 0.99957 | 4 | 100% | P62918 | Rpl8 | 60S riboson | 23.91971 | 22.96292 | 25.35297 | 23.62357 | 24.659 | 23.16656 | 1.392877 | 0.367367 | 2.418387 | 23.94745 | 3.241333 | -3.33863 | 0.015127 | 0.051539 | 1 | 1 |
| 730 | sp\|Q9CYN9 | RENR_MOU | 39 kDa | 0.998453 | 4 | 100% | Q9CYN9 | Atp6ap2 | Renin recep | 21.33995 | 20.01686 | 20.3396 | 19.81024 | 21.11157 | 19.49734 | 1.15556 | 0.302683 | 2.008438 | 20.35259 | 3.229248 | -3.35088 | 0.015248 | 0.05179 | 1 | 1 |
| 945 | sp\|Q62311 | TAF6_MOU | 73 kDa | 0.997951 | 3 | 100% | Q62311 | Taf6 | Transcripti | 21.53924 | 22.48376 | 20.59371 | 21.84033 | 21.20001 | 22.75409 | -1.2484 | -2.16977 | -0.32704 | 21.73519 | -3.23258 | -3.3499 | 0.015277 | 0.05179 | -1 | -1 |
| 484 | tr\|A0A0B4. | A0A0B4J1J | 11 kDa | 0.999849 | 5 | 20% | A0A0B4J1J | Ighv1-82 | Immunoglo | 22.72671 | 25.18137 | 24.90635 | 26.44601 | 23.22512 | 25.42486 | -2.06469 | -3.59068 | -0.53869 | 24.65174 | -3.22888 | -3.35625 | 0.015383 | 0.051893 | -1 | -1 |
| 1185 | sp\|O35963 | RB33B_MO | 26 kDa | 0.998107 | 2 | 100% | O35963 | Rab33b | Ras-related | 18.61365 | 21.92673 | 19.41745 | 24.03444 | 19.06121 | 20.10867 | -2.99251 | -5.20601 | -0.77901 | 20.52702 | -3.22632 | -3.35988 | 0.015436 | 0.051893 | -1 | -1 |
| 987 | sp\|P98195 | ATP9B_MO | 129 kDa | 0.996941 | 3 | 33% | P98195 | Atp9b | Probable p | 22.32723 | 21.06901 | 22.25345 | 21.34965 | 21.88627 | 21.36956 | 0.892905 | 0.231489 | 1.55432 | 21.70919 | 3.213155 | -3.36858 | 0.015446 | 0.051893 | 0 | 0 |
| 1075 | sp\|Q61033 | LAP2A_MO | 75 kDa | 0.999163 | 2 | 100% | Q61033 | Tmpo | Lamina-ass | 17.217 | 16.49091 | 18.70633 | 16.7869 | 17.75882 | 16.16876 | 1.411857 | 0.366835 | 2.456878 | 17.18812 | 3.224155 | -3.36294 | 0.015481 | 0.051893 | 1 | 1 |
| 617 | tr\|A0A0A6 | A0A0A6YV | 13 kDa | 0.999549 | 4 | 50% | A0A0A6YV | Ighv1-23 | Immunoglo | 22.47945 | 24.95764 | 24.66166 | 26.19997 | 23.08218 | 25.1787 | -2.03767 | -3.54674 | -0.5286 | 24.4266 | -3.22237 | -3.36548 | 0.015519 | 0.051893 | -1 | -1 |
| 408 | sp\|P25444 | RS2_MOUS | 31 kDa | 0.99959 | 6 | 100% | P25444 | Rps2 | 40S riboson | 24.90052 | 23.55319 | 25.20167 | 24.13509 | 24.62236 | 24.04582 | 0.996815 | 0.257387 | 1.736242 | 24.40978 | 3.208629 | -3.37504 | 0.015542 | 0.051893 | 0 | 0 |
| 641 | tr\|A0A075 | A0A075B5 | 11 kDa | 0.999388 | 4 | 25% | A0A075B5 | Ighv1-12 | Immunoglo | 22.45846 | 24.94402 | 24.64641 | 26.18745 | 23.05892 | 25.15689 | -2.04153 | -3.55472 | -0.52833 | 24.40869 | -3.21967 | -3.3693 | 0.015575 | 0.051893 | -1 | -1 |
| 1037 | sp\|Q922P9 | GLYR1_MO | 60 kDa | 0.999433 | 2 | 100% | Q922P9 | Glyr1 | Putative ox | 23.28803 | 25.61854 | 24.96253 | 26.65848 | 22.73394 | 25.13884 | -2.14379 | -3.74066 | -0.54692 | 24.73339 | -3.20378 | -3.39183 | 0.015913 | 0.052759 | -1 | -1 |
| 1038 | sp\|Q9R099 | TBL2_MOU | 50 kDa | 0.999431 | 2 | 100% | Q9R099 | Tbl2 | Transducin | 23.28803 | 25.61854 | 24.96253 | 26.65848 | 22.73394 | 25.13884 | -2.14379 | -3.74066 | -0.54692 | 24.73339 | -3.20378 | -3.39183 | 0.015913 | 0.052759 | -1 | -1 |
| 602 | tr\|A0A075 | A0A075B5 | 11 kDa | 0.999702 | 4 | 75% | A0A075B5 | Igkv8-28 | Immunoglo | 22.93043 | 24.69853 | 24.26432 | 25.97369 | 23.40683 | 24.76228 | -1.61097 | -2.81214 | -0.4098 | 24.33935 | -3.20061 | -3.39633 | 0.015982 | 0.052842 | -1 | -1 |
| 916.10 | sp\|Q8BFU2 | H2A3_MOU | 14 kDa | 0.998436 | 3 | 33% | Q8BFU2 | Hist3h2a | Histone H2 | 21.75737 | 23.81681 | 22.88746 | 23.58622 | 22.97043 | 24.67067 | -1.48615 | -2.59528 | -0.37702 | 23.28149 | -3.19765 | -3.40052 | 0.016046 | 0.052842 | -1 | -1 |
| 1109 | sp\|O35465 | FKBP8_MO | 44 kDa | 0.998852 | 2 | 100% | O35465 | Fkbp8 | Peptidyl-pr | 20.08854 | 22.03708 | 20.31717 | 21.40816 | 19.89263 | 20.68473 | -1.27721 | -2.23107 | -0.32335 | 20.73805 | -3.19542 | -3.40369 | 0.016094 | 0.052842 | -1 | -1 |
| 1110 | sp\|P45591 | COF2_MOU | 19 kDa | 0.998832 | 2 | 50% | P45591 | Cfl2 | Cofilin-2 | 20.08854 | 22.03708 | 20.31717 | 21.40816 | 19.89263 | 20.68473 | -1.27721 | -2.23107 | -0.32335 | 20.73805 | -3.19542 | -3.40369 | 0.016094 | 0.052842 | -1 | -1 |
| 258 | sp\|P04104 | K2C1_MOU | 66 kDa | 0.999759 | 8 | 75% | P04104 | Krt1 | Keratin, typ | 26.89041 | 28.2927 | 27.75588 | 28.70276 | 27.80706 | 28.99446 | -1.17885 | -2.06193 | -0.29577 | 28.07388 | -3.1824 | -3.41836 | 0.016272 | 0.053296 | -1 | -1 |
| 269 | sp\|Q3TTY5 | K22E_MOU | 71 kDa | 0.999598 | 8 | 75% | Q3TTY5 | Krt2 | Keratin, typ | 27.42303 | 27.91043 | 27.46561 | 28.48307 | 26.55497 | 28.61685 | -1.18891 | -2.0817 | -0.29613 | 27.74233 | -3.17466 | -3.42938 | 0.016444 | 0.053627 | -1 | -1 |
| 182 | sp\|Q8BTM8 | FLNA_MOU | 281 kDa | 0.999873 | 10 | 90% | Q8BTM8 | Flna | Filamin-A | 21.44473 | 22.20546 | 21.50788 | 22.2146 | 20.82194 | 22.18872 | -0.94474 | -1.65478 | -0.23471 | 21.73056 | -3.1669 | -3.43466 | 0.016452 | 0.053627 | 0 | 0 |
| 490 | sp\|P18527 | HVM56_M | 11 kDa | 0.999709 | 5 | 20% | P18527 | NA | Ig heavy ch | 22.83483 | 25.09691 | 23.69477 | 25.12349 | 22.16545 | 23.72507 | -1.75014 | -3.06584 | -0.43444 | 23.77342 | -3.17443 | -3.43351 | 0.016558 | 0.053715 | -1 | -1 |
| 500 | tr\|A0A075 | A0A075B5 | 13 kDa | 0.999544 | 5 | 40% | A0A075B5 | Ighv5-17 | Immunoglo | 22.83483 | 25.09691 | 23.69477 | 25.12349 | 22.16545 | 23.72507 | -1.75014 | -3.06584 | -0.43444 | 23.77342 | -3.17443 | -3.43351 | 0.016558 | 0.053715 | -1 | -1 |
| 1245 | sp\|Q8BVI5 | STX16_MO | 37 kDa | 0.997032 | 2 | 100% | Q8BVI5 | Stx16 | Syntaxin-16 | 20.14193 | 21.24357 | 20.01671 | 20.61781 | 19.15516 | 20.9595 | -1.16902 | -2.05031 | -0.28774 | 20.35578 | -3.1623 | -3.447 | 0.016723 | 0.054119 | -1 | -1 |
| 466 | sp\|P46638 | RB11B_MO | 24 kDa | 0.998356 | 6 | 100% | P46638 | Rab11b | Ras-related | 22.58047 | 26.01932 | 23.02968 | 23.33919 | 22.75785 | 25.93625 | -2.30892 | -4.04978 | -0.56806 | 23.94379 | -3.16516 | -3.44669 | 0.016768 | 0.054128 | -1 | -1 |
| 1136 | sp\|P80316 | TCPE_MOU | 60 kDa | 0.99857 | 2 | 100% | P80316 | Cct5 | T-complex | 22.13264 | 23.04299 | 22.06049 | 22.687 | 22.01365 | 22.94274 | -0.82198 | -1.44279 | -0.20116 | 22.47992 | -3.15137 | -3.45691 | 0.016805 | 0.054128 | 0 | 0 |
| 79 | sp\|Q8CAQ8 | MIC60_MO | 84 kDa | 0.999498 | 17 | 100% | Q8CAQ8 | Immt | MICOS com | 23.8663 | 24.79439 | 24.22846 | 24.78502 | 23.67939 | 24.78143 | -0.86223 | -1.51852 | -0.20593 | 24.35583 | -3.12696 | -3.4919 | 0.017377 | 0.055837 | 0 | 0 |
| 398 | sp\|P62264 | RS14_MOU | 16 kDa | 0.999699 | 6 | 100% | P62264 | Rps14 | 40S riboson | 24.61183 | 23.89033 | 25.18527 | 23.92356 | 24.67504 | 23.99588 | 0.887456 | 0.211267 | 1.563646 | 24.38032 | 3.123774 | -3.49647 | 0.017453 | 0.055949 | 0 | 0 |
| 1068 | sp\|Q9DCR2 | AP3S1_MO | 22 kDa | 0.999217 | 2 | 100% | Q9DCR2 | Ap3s1 | AP-3 comp | 21.64431 | 17.63543 | 19.06564 | 18.13848 | 21.87576 | 19.16112 | 2.550227 | 0.605097 | 4.495357 | 19.58679 | 3.128822 | -3.49845 | 0.017617 | 0.056254 | 1 | 0 |
| 213 | sp\|Q8BMK4 | CKAP4_MO | 64 kDa | 0.999127 | 10 | 100% | Q8BMK4 | Ckap4 | Cytoskeleto | 20.95514 | 22.1632 | 21.41182 | 23.17518 | 21.88914 | 22.60386 | -1.22871 | -2.16607 | -0.29136 | 22.03306 | -3.12822 | -3.49932 | 0.017631 | 0.056254 | -1 | 0 |
| 360 | sp\|B2RY56 | RBM25_M | 100 kDa | 0.999135 | 7 | 100% | B2RY56 | Rbm25 | RNA-bindin | 22.1694 | 21.06942 | 22.01795 | 21.60517 | 22.17716 | 20.71115 | 0.992924 | 0.232444 | 1.753404 | 21.62504 | 3.107628 | -3.51966 | 0.017845 | 0.05676 | 0 | 0 |
| 916.13 | sp\|Q8CGP7 | H2A1K_M | 14 kDa | 0.998436 | 3 | 33% | Q8CGP7 | Hist1h2ak | Histone H2 | 17.49933 | 19.37395 | 18.35155 | 18.9447 | 17.68718 | NA | -1.31331 | -2.30287 | -0.32374 | 18.37134 | -3.28478 | -3.31376 | 0.017878 | 0.05676 | -1 | 0 |
| 844 | sp\|Q9JII5 | DAZP1_MO | 43 kDa | 0.999212 | 3 | 100% | Q9JII5 | Dazap1 | DAZ-associ | 20.35194 | 21.99044 | 19.6933 | 21.21492 | 18.77912 | 20.58635 | -1.65578 | -2.9237 | -0.38787 | 20.43601 | -3.11648 | -3.51607 | 0.017916 | 0.05676 | -1 | 0 |
| 678 | sp\|Q9CZX8 | RS19_MOU | 16 kDa | 0.999069 | 4 | 100% | Q9CZX8 | Rps19 | 40S riboson | 21.1663 | 21.7784 | 21.68498 | 23.73654 | 20.76809 | 23.36253 | -1.7527 | -3.09593 | -0.40948 | 22.08281 | -3.11393 | -3.5197 | 0.017978 | 0.056825 | -1 | 0 |
| 161 | sp\|P97351 | RS3A_MOU | 30 kDa | 0.9996 | 11 | 100% | P97351 | Rps3a | 40S riboson | 23.82999 | 23.23401 | 24.11161 | 22.54046 | 23.52044 | 22.68903 | 0.999514 | 0.231542 | 1.767486 | 23.32092 | 3.097734 | -3.53388 | 0.018089 | 0.057033 | 0 | 0 |
| 806 | sp\|Q61598 | GDIB_MOU | 51 kDa | 0.999474 | 3 | 67% | Q61598 | Gdi2 | Rab GDP di | 22.89698 | 21.89981 | 22.61068 | 22.17156 | 22.89627 | 21.83406 | 0.832838 | 0.192612 | 1.473064 | 22.38489 | 3.096195 | -3.53609 | 0.018128 | 0.057033 | 0 | 0 |
| 1081 | sp\|Q3TJZ6 | FA98A_MO | 55 kDa | 0.999131 | 2 | 100% | Q3TJZ6 | Fam98a | Protein FAM | 26.18688 | 23.61623 | 25.0779 | 23.13685 | 26.61336 | 25.17665 | 1.982799 | 0.452619 | 3.51298 | 24.96798 | 3.092337 | -3.55056 | 0.018516 | 0.058038 | 1 | 0 |
| 849 | sp\|Q6ZPJ3 | UBE2O_MO | 141 kDa | 0.999172 | 3 | 100% | Q6ZPJ3 | Ube2o | (E3-indepe | 21.22238 | 20.05973 | 20.16797 | 19.47169 | 21.10626 | 18.95748 | 1.335902 | 0.304726 | 2.367077 | 20.16425 | 3.091666 | -3.55152 | 0.018533 | 0.058038 | 1 | 0 |
| 609 | sp\|P01639 | KV5A7_MO | 14 kDa | 0.999647 | 4 | 50% | P01639 | Gm5571 | Ig kappa ch | 20.90958 | 22.87286 | 21.92188 | 23.28731 | 20.92352 | 21.9382 | -1.4478 | -2.57159 | -0.324 | 21.97556 | -3.07447 | -3.57612 | 0.018974 | 0.059115 | -1 | 0 |
| 364 | sp\|Q9R0N5 | SYT5_MOU | 43 kDa | 0.999051 | 7 | 71% | Q9R0N5 | Syt5 | Synaptotag | 21.4152 | 20.62117 | 21.62429 | 21.05952 | 21.80493 | 20.41685 | 0.915628 | 0.204058 | 1.627197 | 21.15699 | 3.062684 | -3.58433 | 0.018985 | 0.059115 | 0 | 0 |
| 373 | sp\|P11983 | TCPA_MOU | 60 kDa | 0.998923 | 7 | 86% | P11983 | Tcp1 | T-complex | 22.03277 | 23.32591 | 23.27968 | 27.16754 | 22.5978 | 26.63184 | -3.07168 | -5.45696 | -0.6864 | 24.17259 | -3.07318 | -3.57797 | 0.019007 | 0.059115 | -1 | 0 |
| 1141 | sp\|Q8BTX9 | HSDL1_MO | 37 kDa | 0.998519 | 2 | 100% | Q8BTX9 | Hsdl1 | Inactive hy | 19.83488 | 24.78124 | 17.29591 | 24.24659 | 19.97972 | 20.23571 | -4.05101 | -7.20072 | -0.9013 | 21.06234 | -3.06933 | -3.58348 | 0.019108 | 0.059291 | -1 | 0 |
| 810 | sp\|Q9R112 | SQOR_MOU | 50 kDa | 0.99945 | 3 | 100% | Q9R112 | Sqor | Sulfide:qui | 20.86886 | 21.47694 | 20.97894 | 21.78368 | 20.24894 | 21.66869 | -0.94419 | -1.68116 | -0.20721 | 21.17101 | -3.04935 | -3.60355 | 0.019338 | 0.059868 | 0 | 0 |
| 530 | sp\|Q9WVR4 | FXR2_MOU | 74 kDa | 0.999225 | 5 | 60% | Q9WVR4 | Fxr2 | Fragile X m | 21.13431 | 22.50088 | 20.4116 | 21.58402 | 21.16765 | 22.02335 | -1.13157 | -2.01667 | -0.24646 | 21.4703 | -3.04774 | -3.61104 | 0.019562 | 0.060255 | -1 | 0 |
| 306.1 | sp\|P01863 | GCAA_MOU | 36 kDa | 0.999867 | 7 | 86% | P01863 | Ighg | Ig gamma-2 | 27.36892 | 28.74938 | 28.56541 | 29.53275 | 27.71947 | 29.09629 | -1.24154 | -2.21267 | -0.2704 | 28.50537 | -3.05092 | -3.60985 | 0.019596 | 0.060255 | -1 | 0 |
| 306.2 | sp\|P01865 | GCAM_MO | 44 kDa | 0.999867 | 7 | 86% | P01865 | Igh-1a | Ig gamma-2 | 27.36892 | 28.74938 | 28.56541 | 29.53275 | 27.71947 | 29.09629 | -1.24154 | -2.21267 | -0.2704 | 28.50537 | -3.05092 | -3.60985 | 0.019596 | 0.060255 | -1 | 0 |
| 157 | sp\|Q9D0E1 | HNRPM_M | 78 kDa | 0.999898 | 11 | 100% | Q9D0E1 | Hnrnpm | Heterogene | 23.81454 | 23.01013 | 23.27448 | 22.16757 | 23.58094 | 22.45551 | 1.012249 | 0.218618 | 1.80588 | 23.05053 | 3.035775 | -3.62314 | 0.019704 | 0.060451 | 1 | 0 |
| 90 | sp\|P07901 | HS90A_MO | 85 kDa | 0.99979 | 16 | 56% | P07901 | Hsp90aa1 | Heat shock | 23.48696 | 24.244 | 23.47314 | 24.19927 | 22.50916 | 24.20911 | -1.06104 | -1.89443 | -0.22765 | 23.68694 | -3.03044 | -3.63099 | 0.019856 | 0.060779 | -1 | 0 |
| 1205 | sp\|Q921F4 | HNRLL_MO | 64 kDa | 0.997883 | 2 | 100% | Q921F4 | Hnrnpll | Heterogene | 19.87285 | 20.43901 | 19.90515 | 20.61592 | 19.68001 | 20.79096 | -0.79596 | -1.42435 | -0.16757 | 20.21732 | -3.01482 | -3.65339 | 0.020285 | 0.061898 | 0 | 0 |
| 1097 | sp\|Q91YP0 | L2HDH_M | 51 kDa | 0.998996 | 2 | 100% | Q91YP0 | L2hgdh | L-2-hydrox | 19.68227 | 20.11841 | 19.74544 | 21.1738 | 19.49732 | 20.75919 | -1.04212 | -1.86511 | -0.21913 | 20.16274 | -3.01402 | -3.65471 | 0.020313 | 0.061898 | -1 | 0 |
| 323 | sp\|P35980 | RL18_MOU | 22 kDa | 0.999574 | 7 | 100% | P35980 | Rpl18 | 60S riboson | 25.86043 | 25.50647 | 26.16436 | 24.52851 | 25.80528 | 24.252 | 1.181033 | 0.245946 | 2.11612 | 25.35284 | 3.01412 | -3.66267 | 0.020613 | 0.062671 | 1 | 0 |
| 931 | sp\|Q3TXS7 | PSMD1_M | 106 kDa | 0.998211 | 3 | 100% | Q3TXS7 | Psmd1 | 26S protea | 21.5493 | 20.55774 | 22.5378 | 20.84932 | 21.24385 | 20.13167 | 1.264069 | 0.261182 | 2.266957 | 21.14495 | 3.00794 | -3.67155 | 0.020789 | 0.063065 | 1 | 0 |
| 905 | tr\|G3X8U3 | G3X8U3_M | 39 kDa | 0.998612 | 3 | 100% | G3X8U3 | 2210016F1 | Queuosine | 22.50088 | 21.59274 | 22.86089 | 22.04512 | 22.57885 | 21.81241 | 0.830117 | 0.166165 | 1.494069 | 22.23182 | 2.975797 | -3.70985 | 0.021415 | 0.064819 | 0 | 0 |
| 63 | sp\|Q9ESU6 | BRD4_MOU | 156 kDa | 0.999719 | 18 | 100% | Q9ESU6 | Brd4 | Bromodom | 25.06452 | 23.9049 | 23.7937 | 22.30319 | 25.51418 | 22.88037 | 1.761315 | 0.351272 | 3.171358 | 23.91014 | 2.980954 | -3.71037 | 0.021578 | 0.065165 | 1 | 0 |
| 322 | sp\|Q3TDQ1 | STT3B_MO | 93 kDa | 0.999583 | 7 | 86% | Q3TDQ1 | Stt3b | Dolichyl-di | 23.29999 | 22.48761 | 22.96804 | 22.32553 | 23.13935 | 22.27421 | 0.773343 | 0.150134 | 1.396552 | 22.74912 | 2.953514 | -3.74213 | 0.02209 | 0.066566 | 0 | 0 |
| 656 | sp\|O70404 | VAMP8_M | 11 kDa | 0.999267 | 4 | 100% | O70404 | Vamp8 | Vesicle-ass | 25.10347 | 23.49243 | 24.69223 | 23.33267 | 24.8207 | 24.45309 | 1.112737 | 0.215775 | 2.009699 | 24.31577 | 2.957416 | -3.74119 | 0.022164 | 0.066639 | 1 | 0 |
| 1137 | sp\|Q9CYG7 | TOM34_M | 34 kDa | 0.998561 | 2 | 100% | Q9CYG7 | Tomm34 | Mitochond | 19.79804 | 20.41211 | 19.44821 | 20.48668 | 19.50637 | 20.21103 | -0.78573 | -1.42265 | -0.14882 | 19.97708 | -2.93626 | -3.76716 | 0.022629 | 0.067396 | 0 | 0 |
| 353 | tr\|A0A2R8 | A0A2R8VH | 58 kDa | 0.999252 | 7 | 29% | A0A2R8VH | Gm5478 | Predicted p | 25.24538 | 25.88246 | 25.78906 | 26.98654 | 24.1742 | 26.94941 | -1.53659 | -2.78182 | -0.29136 | 25.83784 | -2.94483 | -3.76243 | 0.022683 | 0.067396 | -1 | 0 |
| 1088 | sp\|P27612 | PLAP_MOU | 87 kDa | 0.999063 | 2 | 100% | P27612 | Plaa | Phospholip | 18.74677 | 21.07718 | 18.88185 | 19.2794 | 18.94684 | 21.83052 | -1.87054 | -3.3868 | -0.35429 | 19.79376 | -2.94406 | -3.76353 | 0.022707 | 0.067396 | -1 | 0 |
| 929 | tr\|A0A075 | A0A075B5 | 13 kDa | 0.998258 | 3 | 33% | A0A075B5 | Ighv14-2 | Immunoglo | 19.51302 | 20.26388 | 19.08072 | 20.14489 | 19.59203 | 20.20793 | -0.81031 | -1.46773 | -0.15289 | 19.80041 | -2.93363 | -3.77097 | 0.022712 | 0.067396 | 0 | 0 |
| 101 | sp\|P01872 | IGHM_MOU | 50 kDa | 0.999648 | 15 | 100% | P01872 | Ighm | Immunoglo | 27.19116 | 28.49345 | 27.11103 | 28.4493 | 26.34852 | 27.41747 | -1.2365 | -2.23901 | -0.234 | 27.50182 | -2.94347 | -3.76439 | 0.022726 | 0.067396 | -1 | 0 |
| 267 | sp\|P62806 | H4_MOUSE | 11 kDa | 0.999599 | 8 | 100% | P62806 | Hist4h4 | Histone H4 | 28.31263 | 27.3346 | 27.27886 | 25.73448 | 28.04698 | 26.0879 | 1.49383 | 0.282503 | 2.705158 | 27.13258 | 2.943 | -3.76506 | 0.022741 | 0.067396 | 1 | 0 |
| 695 | sp\|Q8QZT1 | THIL_MOUS | 45 kDa | 0.99896 | 4 | 100% | Q8QZT1 | Acat1 | Acetyl-CoA | 20.63042 | 21.06243 | 20.81015 | 22.70028 | 20.11609 | 23.79761 | -2.00122 | -3.62438 | -0.37806 | 21.5195 | -2.94229 | -3.76609 | 0.022763 | 0.067396 | -1 | 0 |
| 1002.2 | sp\|Q80VQ0 | AL3B1_MO | 52 kDa | 0.995458 | 3 | 100% | Q80VQ0 | Aldh3b1 | Aldehyde d | 20.88722 | 19.67256 | 23.96654 | 17.32303 | 21.72996 | 20.42034 | 3.05593 | 0.571543 | 5.540317 | 20.66661 | 2.935451 | -3.77595 | 0.02298 | 0.067424 | 1 | 0 |
| 481 | tr\|A0A075 | A0A075B5 | 11 kDa | 0.999942 | 5 | 40% | A0A075B5 | Ighv1-43 | Immunoglo | 22.81384 | 25.18867 | 25.11803 | 26.68872 | 23.29034 | 25.45492 | -2.0367 | -3.69291 | -0.38049 | 24.75909 | -2.93469 | -3.77706 | 0.023005 | 0.067424 | -1 | 0 |
| 84 | sp\|Q922U2 | K2C5_MOU | 62 kDa | 0.998902 | 17 | 47% | Q922U2 | Krt5 | Keratin, typ | 23.34071 | 25.41956 | 25.3252 | 26.50245 | 24.71066 | 26.81444 | -1.78662 | -3.2417 | -0.33154 | 25.35217 | -2.9302 | -3.78353 | 0.023148 | 0.067424 | -1 | 0 |
| 729 | sp\|P62334 | PRS10_MO | 44 kDa | 0.998479 | 4 | 100% | P62334 | Psmc6 | 26S protea | 22.05284 | 23.49345 | 21.79907 | 27.08325 | 21.94051 | 23.56295 | -2.78241 | -5.04856 | -0.51626 | 23.32201 | -2.9301 | -3.78368 | 0.023151 | 0.067424 | -1 | 0 |
| 453 | sp\|Q9Z110 | P5CS_MOU | 87 kDa | 0.998979 | 6 | 100% | Q9Z110 | Aldh18a1 | Delta-1-pyr | 20.49007 | 21.0054 | 20.21388 | 21.84483 | 20.12265 | 20.96502 | -0.99621 | -1.80828 | -0.18414 | 20.77364 | -2.91998 | -3.79092 | 0.023155 | 0.067424 | 0 | 0 |
| 583 | sp\|P84091 | AP2M1_M | 50 kDa | 0.997211 | 5 | 100% | P84091 | Ap2m1 | AP-2 comp | 23.81037 | 22.32336 | 23.57002 | 22.19383 | 22.90525 | 22.73735 | 1.010369 | 0.186655 | 1.834082 | 22.92336 | 2.919616 | -3.79146 | 0.023167 | 0.067424 | 1 | 0 |
| 464 | sp\|Q6ZPR4 | KCNT1_MO | 138 kDa | 0.998579 | 6 | 100% | Q6ZPR4 | Kcnt1 | Potassium | 19.77841 | 17.82771 | 18.95494 | 16.91251 | 20.32247 | 18.98341 | 1.777396 | 0.328975 | 3.225818 | 18.79657 | 2.928464 | -3.78604 | 0.023204 | 0.067424 | 1 | 0 |
| 1163 | sp\|P00405 | COX2_MOU | 26 kDa | 0.998301 | 2 | 100% | P00405 | Mtco2 | Cytochrom | 16.45857 | 18.43037 | 18.35068 | 18.63351 | 16.46537 | 19.78843 | -1.85923 | -3.37446 | -0.344 | 18.02115 | -2.92823 | -3.78638 | 0.023212 | 0.067424 | -1 | 0 |
| 587 | sp\|P47857 | PFKAM_M | 85 kDa | 0.996794 | 5 | 60% | P47857 | Pfkm | ATP-depen | 20.18599 | 20.98899 | 19.55696 | 20.47379 | 19.05935 | 20.59214 | -1.08421 | -1.96856 | -0.19987 | 20.14287 | -2.92271 | -3.79137 | 0.023259 | 0.067424 | -1 | 0 |
| 751.2 | sp\|P01656 | KV3A3_MO | 12 kDa | 0.997764 | 4 | 25% | P01656 | NA | Ig kappa ch | 22.20678 | 21.61101 | 22.09707 | 21.46517 | 22.32532 | 21.23552 | 0.772497 | 0.141384 | 1.40361 | 21.82348 | 2.913337 | -3.80044 | 0.023366 | 0.067424 | 0 | 0 |
| 352 | sp\|P43274 | H14_MOUS | 22 kDa | 0.999273 | 7 | 29% | P43274 | Hist1h1e | Histone H1 | 24.95628 | 24.40787 | 25.13317 | 24.51737 | 25.59897 | 24.15171 | 0.870488 | 0.159246 | 1.58173 | 24.79423 | 2.91304 | -3.80087 | 0.023375 | 0.067424 | 0 | 0 |
| 1053 | sp\|Q9D1D4 | TMEDA_M | 25 kDa | 0.999297 | 2 | 100% | Q9D1D4 | Tmed10 | Transmemb | 21.72646 | 22.43562 | 21.72431 | 22.37141 | 21.81843 | 22.82564 | -0.78782 | -1.43156 | -0.14409 | 22.15031 | -2.91287 | -3.80112 | 0.023381 | 0.067424 | 0 | 0 |
| 974 | sp\|O88935 | SYN1_MOU | 74 kDa | 0.997544 | 3 | 67% | O88935 | Syn1 | Synapsin-1 | NA | 18.52299 | 16.46347 | 19.55697 | 18.02984 | 19.07375 | -1.80458 | -3.26115 | -0.34801 | 18.32941 | -3.07097 | -3.59725 | 0.02342 | 0.067424 | -1 | 0 |
| 636 | tr\|J3QK03 | J3QK03_M | 11 kDa | 0.99941 | 4 | 50% | J3QK03 | Ighv6-5 | Immunoglo | 19.24566 | 20.88704 | 19.94511 | 21.38176 | 18.5662 | 19.94772 | -1.48651 | -2.70306 | -0.26997 | 19.99558 | -2.91602 | -3.80401 | 0.023608 | 0.067824 | -1 | 0 |
| 880 | sp\|Q9CW0 | SMC3_MOU | 142 kDa | 0.998919 | 3 | 100% | Q9CW03 | Smc3 | Structural | 21.30276 | 20.49528 | 22.70691 | 21.01078 | 22.34286 | 21.09511 | 1.25045 | 0.220391 | 2.280509 | 21.49228 | 2.897043 | -3.83143 | 0.02424 | 0.06949 | 1 | 0 |
| 858 | sp\|Q61545 | EWS_MOUS | 68 kDa | 0.99907 | 3 | 100% | Q61545 | Ewsr1 | RNA-bindin | 18.34569 | 19.49539 | 18.86731 | 19.66019 | 17.73537 | 19.06338 | -1.0902 | -1.98993 | -0.19046 | 18.86122 | -2.88858 | -3.84081 | 0.024393 | 0.069783 | -1 | 0 |
| 770 | sp\|Q9WUA | SYFB_MOU | 66 kDa | 0.996919 | 4 | 100% | Q9WUA2 | Farsb | Phenylalan | 21.28569 | 19.8359 | 22.92422 | 20.56716 | 21.21848 | 20.46881 | 1.518838 | 0.263461 | 2.774214 | 21.05004 | 2.887274 | -3.84556 | 0.024571 | 0.070145 | 1 | 0 |
| 626 | sp\|Q9D1M | SEC13_MO | 36 kDa | 0.999481 | 4 | 100% | Q9D1M0 | Sec13 | Protein SEC | 22.36954 | 20.35718 | 21.84705 | 20.1058 | 21.82239 | 21.61808 | 1.319306 | 0.227413 | 2.4112 | 21.35334 | 2.883474 | -3.85105 | 0.024702 | 0.070369 | 1 | 0 |
| 600 | tr\|A0A140 | A0A140T8 | 13 kDa | 0.999708 | 4 | 25% | A0A140T8 | Igkv6-25 | Immunoglo | 20.74847 | 22.99187 | 22.60429 | 23.77756 | 20.8654 | 22.56049 | -1.70392 | -3.11668 | -0.29116 | 22.25801 | -2.87826 | -3.8586 | 0.024882 | 0.070733 | -1 | 0 |
| 590 | sp\|Q6P542 | ABCF1_MO | 95 kDa | 0.999828 | 4 | 100% | Q6P542 | Abcf1 | ATP-bindin | 20.71838 | 17.61124 | 18.74682 | 17.92869 | 18.71617 | 17.72178 | 1.639884 | 0.27567 | 3.004098 | 18.57385 | 2.868674 | -3.87247 | 0.025217 | 0.071535 | 1 | 0 |
| 127 | sp\|Q8BHS8 | SYBU_MOU | 73 kDa | 0.999745 | 13 | 85% | Q8BHS8 | Sybu | Syntabulin | 24.71025 | 23.33591 | 23.35876 | 20.98313 | 24.62368 | 22.5543 | 1.939781 | 0.322297 | 3.557264 | 23.26101 | 2.861959 | -3.88219 | 0.025454 | 0.072058 | 1 | 0 |
| 539 | sp\|P61620 | S61A1_MO | 52 kDa | 0.999172 | 5 | 100% | P61620 | Sec61a1 | Protein tra | 22.0348 | 21.37862 | 22.38587 | 21.40309 | 21.90202 | 21.20375 | 0.779075 | 0.125891 | 1.43226 | 21.71802 | 2.838864 | -3.90879 | 0.025941 | 0.073283 | 0 | 0 |
| 470 | sp\|P84084 | ARF5_MOU | 21 kDa | 0.998134 | 6 | 50% | P84084 | Arf5 | ADP-ribosy | 21.94885 | 23.92619 | 20.69065 | 21.74001 | 20.42162 | 22.96983 | -1.85831 | -3.42507 | -0.29154 | 21.94953 | -2.83051 | -3.92777 | 0.026598 | 0.07494 | -1 | 0 |
| 412 | sp\|Q9ER58 | TICN2_MO | 47 kDa | 0.999573 | 6 | 100% | Q9ER58 | Spock2 | Testican-2 | 21.94076 | 24.06247 | 21.67806 | 22.17376 | 21.94692 | 23.67778 | -1.44943 | -2.67238 | -0.22647 | 22.57996 | -2.82836 | -3.93087 | 0.026678 | 0.07494 | -1 | 0 |
| 862 | sp\|Q9CQC7 | NDUB4_M | 15 kDa | 0.999046 | 3 | 100% | Q9CQC7 | Ndufb4 | NADH dehy | 20.37357 | 21.85819 | 21.24276 | 21.69905 | 20.57247 | 21.43936 | -0.93594 | -1.72628 | -0.14559 | 21.19757 | -2.81858 | -3.93836 | 0.026693 | 0.07494 | 0 | 0 |
| 190 | sp\|P07356 | ANXA2_MO | 39 kDa | 0.999661 | 10 | 100% | P07356 | Anxa2 | Annexin A2 | 23.67308 | 24.04217 | 23.4087 | 24.61763 | 23.63475 | 24.476 | -0.80642 | -1.48961 | -0.12324 | 23.97539 | -2.80947 | -3.95165 | 0.027038 | 0.075752 | 0 | 0 |
| 1036 | sp\|Q9WUQ | PREB_MOU | 45 kDa | 0.999441 | 2 | 100% | Q9WUQ2 | Preb | Prolactin r | 15.84759 | 18.96055 | 17.54844 | 18.72038 | 16.36642 | 17.26072 | -1.7264 | -3.19625 | -0.25655 | 17.45068 | -2.80298 | -3.96771 | 0.027644 | 0.077288 | -1 | 0 |
| 1287 | sp\|P26638 | SYSC_MOU | 58 kDa | 0.995287 | 2 | 100% | P26638 | Sars | Serine--tRN | 17.36066 | 16.74451 | 17.39737 | 16.11356 | 16.99057 | 15.13956 | 1.250322 | 0.183512 | 2.317132 | 16.62437 | 2.796955 | -3.97645 | 0.027878 | 0.077783 | 1 | 0 |
| 225 | sp\|Q60930 | VDAC2_M | 32 kDa | 0.999813 | 9 | 100% | Q60930 | Vdac2 | Voltage-de | 23.99638 | 23.4321 | 24.1305 | 23.07336 | 23.76874 | 23.10344 | 0.762237 | 0.110134 | 1.41434 | 23.58409 | 2.782112 | -3.99158 | 0.028104 | 0.078142 | 0 | 0 |
| 1170 | sp\|P30275 | KCRU_MOU | 47 kDa | 0.998235 | 2 | 100% | P30275 | Ckmt1 | Creatine ki | 24.52315 | 23.90961 | 24.81755 | 23.8433 | 24.40754 | 23.7831 | 0.737407 | 0.106444 | 1.36837 | 24.21404 | 2.78166 | -3.99224 | 0.028122 | 0.078142 | 0 | 0 |
| 1147 | sp\|P63325 | RS10_MOU | 19 kDa | 0.998503 | 2 | 100% | P63325 | Rps10 | 40S riboson | 22.95405 | 21.94229 | 23.53032 | 22.16273 | 22.45879 | 21.94641 | 0.96391 | 0.138123 | 1.789697 | 22.4991 | 2.778373 | -3.99716 | 0.02826 | 0.078299 | 0 | 0 |
| 687 | tr\|A0A075 | A0A075B5 | 11 kDa | 0.999008 | 4 | 50% | A0A075B5 | Igkv8-27 | Immunoglo | 18.70096 | 21.10714 | 19.27954 | 21.14212 | 20.64572 | 24.14502 | -2.58935 | -4.80702 | -0.37168 | 20.83675 | -2.78641 | -3.99176 | 0.028294 | 0.078299 | -1 | 0 |
| 469 | sp\|Q68FL6 | SYMC_MOU | 101 kDa | 0.998165 | 6 | 100% | Q68FL6 | Mars | Methionine | 19.51202 | 21.3058 | 20.22308 | 20.2157 | 19.70669 | 21.35491 | -1.14487 | -2.12608 | -0.16366 | 20.38637 | -2.7845 | -3.99454 | 0.02837 | 0.07835 | -1 | 0 |
| 141 | sp\|Q9DBG3 | AP2B1_MO | 105 kDa | 0.997806 | 13 | 46% | Q9DBG3 | Ap2b1 | AP-2 comp | 23.15598 | 21.37204 | 23.25725 | 21.8094 | 21.75426 | 20.86725 | 1.372933 | 0.193299 | 2.552567 | 22.03603 | 2.77749 | -4.00472 | 0.028651 | 0.078964 | 1 | 0 |
| 114 | sp\|O70624 | MYOC_MOU | 55 kDa | 0.999859 | 14 | 100% | O70624 | Myoc | Myocilin | 24.06301 | 23.32106 | 23.72197 | 22.85188 | 23.66256 | 22.87739 | 0.799072 | 0.111022 | 1.487121 | 23.41631 | 2.764183 | -4.01776 | 0.028826 | 0.079287 | 0 | 0 |
| 951 | sp\|Q8K1M | DNM1L_M | 83 kDa | 0.997901 | 3 | 100% | Q8K1M6 | Dnm1l | Dynamin-1 | 20.22039 | 18.34894 | 20.94661 | 19.36239 | 20.20156 | 19.83991 | 1.27244 | 0.176408 | 2.368472 | 19.81997 | 2.770543 | -4.01481 | 0.028932 | 0.079416 | 1 | 0 |
| 14 | sp\|O88737 | BSN_MOUS | 419 kDa | 0.9995 | 35 | 100% | O88737 | Bsn | Protein bas | 23.04352 | 25.00809 | 21.45565 | 25.0599 | 24.43219 | 24.75986 | -1.96549 | -3.66361 | -0.26738 | 23.95987 | -2.76221 | -4.02692 | 0.029273 | 0.080049 | -1 | 0 |
| 392 | sp\|Q3UHK1 | MYCT_MOU | 69 kDa | 0.999778 | 6 | 100% | Q3UHK1 | Slc2a13 | Proton myo | 22.47252 | 23.45209 | 22.9104 | 23.64619 | 22.39145 | 23.13867 | -0.82086 | -1.53059 | -0.11112 | 23.00189 | -2.75279 | -4.03441 | 0.029295 | 0.080049 | 0 | 0 |
| 845 | tr\|Q80ZM5 | Q80ZM5_M | 20 kDa | 0.999209 | 3 | 100% | Q80ZM5 | H1fx | H1 histone | 21.13084 | 22.67627 | 21.70975 | 21.97843 | 21.35144 | 22.18377 | -0.88215 | -1.64517 | -0.11913 | 21.83842 | -2.75173 | -4.03597 | 0.02934 | 0.080049 | 0 | 0 |
| 299 | sp\|Q11011 | PSA_MOUS | 103 kDa | 0.998545 | 8 | 100% | Q11011 | Npepps | Puromycin | 21.21294 | 23.02534 | 22.30321 | 23.37755 | 21.85324 | 22.3792 | -1.13757 | -2.12269 | -0.15244 | 22.35858 | -2.75573 | -4.03633 | 0.029541 | 0.08039 | -1 | 0 |
| 526 | sp\|Q9ER69 | FL2D_MOU | 44 kDa | 0.999285 | 5 | 100% | Q9ER69 | Wtap | Pre-mRNA- | 18.5583 | 19.12888 | 17.4902 | 19.38718 | 18.68982 | 19.43559 | -1.07111 | -1.99903 | -0.14319 | 18.78166 | -2.75467 | -4.03784 | 0.029583 | 0.08039 | -1 | 0 |
| 1165 | sp\|P01027 | CO3_MOUS | 186 kDa | 0.998285 | 2 | 100% | P01027 | C3 | Compleme | 20.40933 | 19.88472 | 20.16579 | 19.19634 | 20.19402 | 18.81703 | 0.957015 | 0.126382 | 1.787647 | 19.77787 | 2.742406 | -4.0497 | 0.029737 | 0.080406 | 0 | 0 |
| 789 | sp\|Q9D384 | U1SBP_MO | 29 kDa | 0.999646 | 3 | 100% | Q9D384 | Snrnp35 | U11/U12 sn | 18.22159 | 19.78875 | 16.79628 | 19.43194 | 19.07134 | 19.60016 | -1.57722 | -2.94613 | -0.2083 | 18.81834 | -2.74957 | -4.04529 | 0.029798 | 0.080406 | -1 | 0 |
| 399 | sp\|Q3B7Z2 | OSBP1_MO | 89 kDa | 0.999671 | 6 | 100% | Q3B7Z2 | Osbp | Oxysterol-b | 22.60444 | 20.02101 | 21.7037 | 21.84946 | 22.58158 | 20.61257 | 1.468892 | 0.193964 | 2.743819 | 21.56212 | 2.749507 | -4.04539 | 0.029801 | 0.080406 | 1 | 0 |
| 414 | sp\|Q9Z1Z0 | USO1_MOU | 107 kDa | 0.999567 | 6 | 100% | Q9Z1Z0 | Uso1 | General ves | 22.22573 | 22.81753 | 22.1343 | 22.71304 | 21.71459 | 22.8099 | -0.75529 | -1.41134 | -0.09923 | 22.40251 | -2.74013 | -4.05291 | 0.029826 | 0.080406 | 0 | 0 |
| 662 | sp\|P63024 | VAMP3_M | 11 kDa | 0.999219 | 4 | 50% | P63024 | Vamp3 | Vesicle-ass | 22.11123 | 20.79044 | 20.86191 | 19.33916 | 21.63336 | 20.36416 | 1.370912 | 0.178921 | 2.562904 | 20.85004 | 2.744651 | -4.05245 | 0.030005 | 0.080729 | 1 | 0 |
| 463 | sp\|Q9DCD2 | SYF1_MOU | 100 kDa | 0.998711 | 6 | 100% | Q9DCD2 | Xab2 | Pre-mRNA- | 20.7791 | 20.10436 | 20.67987 | 20.2501 | 21.74623 | 19.77823 | 1.024172 | 0.131058 | 1.917286 | 20.55631 | 2.733759 | -4.06589 | 0.030322 | 0.081418 | 1 | 0 |
| 278 | sp\|Q8VDJ3 | VIGLN_MO | 142 kDa | 0.99943 | 8 | 100% | Q8VDJ3 | Hdlbp | Vigilin | 21.88987 | 21.0612 | 21.71129 | 20.23761 | 21.73413 | 21.2303 | 0.935393 | 0.117732 | 1.753054 | 21.31073 | 2.72297 | -4.07812 | 0.030569 | 0.081781 | 0 | 0 |
| 981 | sp\|Q6NSU3 | GL8D1_M | 42 kDa | 0.997254 | 3 | 100% | Q6NSU3 | Glt8d1 | Glycosyltra | 18.38342 | 16.25019 | 15.80608 | 13.96295 | 19.09967 | 15.22438 | 2.617219 | 0.330419 | 4.904018 | 16.45445 | 2.731257 | -4.07193 | 0.030577 | 0.081781 | 1 | 0 |
| 236 | sp\|Q00PI9 | HNRL2_MO | 85 kDa | 0.999487 | 9 | 100% | Q00PI9 | Hnrnpul2 | Heterogene | 23.46053 | 21.80072 | 23.27767 | 21.56276 | 22.95646 | 22.90901 | 1.140717 | 0.141444 | 2.13999 | 22.66119 | 2.724233 | -4.08215 | 0.030882 | 0.082417 | 1 | 0 |
| 836.1 | sp\|P01638 | KV5A6_MO | 13 kDa | 0.999277 | 3 | 100% | P01638 | NA | Ig kappa ch | 17.54744 | 14.73293 | 15.63094 | 16.03972 | 17.94353 | 13.2049 | 2.381449 | 0.29432 | 4.468578 | 15.84991 | 2.722969 | -4.08399 | 0.030937 | 0.082417 | 1 | 0 |
| 1102 | sp\|Q09163 | DLK1_MOU | 41 kDa | 0.998951 | 2 | 100% | Q09163 | Dlk1 | Protein del | 21.01135 | 19.91239 | 20.76942 | 20.2948 | 21.63649 | 20.45027 | 0.919935 | 0.112343 | 1.727526 | 20.67912 | 2.71136 | -4.0951 | 0.031077 | 0.082629 | 0 | 0 |
| 798.3 | tr\|A0A2I3 | A0A2I3BRL | 42 kDa | 0.999544 | 3 | 100% | A0A2I3BRL | Gm7324 | Predicted g | 21.97355 | 22.70747 | 21.68837 | 22.54516 | 22.18523 | 22.85399 | -0.75315 | -1.41625 | -0.09006 | 22.32563 | -2.70338 | -4.10667 | 0.031425 | 0.08339 | 0 | 0 |
| 1270 | sp\|Q9CQW | YKT6_MOU | 22 kDa | 0.996185 | 2 | 100% | Q9CQW1 | Ykt6 | Synaptobre | 15.1606 | 16.13552 | 15.10459 | 16.04965 | 15.83482 | 16.82286 | -0.96934 | -1.82324 | -0.11544 | 15.85134 | -2.7056 | -4.10645 | 0.031522 | 0.083484 | 0 | 0 |
| 294 | sp\|Q4G0F8 | UBN1_MOU | 122 kDa | 0.998983 | 8 | 100% | Q4G0F8 | Ubn1 | Ubinuclein | 22.94501 | 23.40816 | 20.99325 | 23.90496 | 22.96744 | 24.64749 | -1.68497 | -3.18058 | -0.18935 | 23.14439 | -2.68857 | -4.13407 | 0.032478 | 0.085848 | -1 | 0 |
| 462 | sp\|Q9Z1N5 | DX39B_MO | 49 kDa | 0.998744 | 6 | 100% | Q9Z1N5 | Ddx39b | Spliceosom | 21.87734 | 24.82251 | 21.67497 | 22.01079 | 21.87744 | 24.41092 | -1.93816 | -3.66133 | -0.21498 | 22.779 | -2.68417 | -4.14048 | 0.032681 | 0.086216 | -1 | 0 |
| 207 | sp\|P02535 | K1C10_MO | 58 kDa | 0.999318 | 10 | 80% | P02535 | Krt10 | Keratin, typ | 28.48804 | 28.91114 | 28.41481 | 29.1065 | 27.67688 | 30.3615 | -1.26647 | -2.39342 | -0.13953 | 28.82648 | -2.68191 | -4.14377 | 0.032785 | 0.086324 | -1 | 0 |
| 147 | sp\|P70704 | AT8A1_MO | 131 kDa | 0.999574 | 12 | 100% | P70704 | Atp8a1 | Phospholip | 22.34894 | 23.84426 | 23.58151 | 23.71198 | 22.43669 | 24.00702 | -1.06537 | -2.01426 | -0.11648 | 23.32173 | -2.67937 | -4.14746 | 0.032903 | 0.086466 | -1 | 0 |
| 80 | sp\|Q3TEA8 | HP1B3_MO | 61 kDa | 0.999402 | 17 | 100% | Q3TEA8 | Hp1bp3 | Heterochro | 24.89401 | 23.54595 | 25.31159 | 24.88947 | 25.29301 | 23.61559 | 1.149199 | 0.123926 | 2.174472 | 24.5916 | 2.674892 | -4.15399 | 0.033112 | 0.086848 | 1 | 0 |
| 813 | sp\|Q9CRB9 | MIC19_MO | 26 kDa | 0.999433 | 3 | 100% | Q9CRB9 | Chchd3 | MICOS com | 19.50863 | 21.93516 | 21.54366 | 21.97173 | 19.91339 | 21.48422 | -1.47514 | -2.79377 | -0.15652 | 21.05947 | -2.66971 | -4.16154 | 0.033356 | 0.087319 | -1 | 0 |
| 514 | sp\|P17751 | TPIS_MOUS | 32 kDa | 0.999415 | 5 | 100% | P17751 | Tpi1 | Triosephos | 21.1336 | 21.92856 | 20.82541 | 22.05441 | 21.35506 | 21.63712 | -0.76867 | -1.45694 | -0.0804 | 21.48902 | -2.65816 | -4.17286 | 0.033516 | 0.087525 | 0 | 0 |
| 992 | tr\|A0A140 | A0A140T8 | 13 kDa | 0.996713 | 3 | 67% | A0A140T8 | Igkv1-117 | Immunoglo | 20.8362 | 21.81629 | 21.39161 | 22.61264 | 21.5681 | 22.13768 | -0.92357 | -1.75132 | -0.09583 | 21.72709 | -2.65581 | -4.17641 | 0.033636 | 0.087525 | 0 | 0 |
| 993 | sp\|Q8BP47 | SYNC_MOU | 64 kDa | 0.99656 | 3 | 100% | Q8BP47 | Nars | Asparagine | 20.8362 | 21.81629 | 21.39161 | 22.61264 | 21.5681 | 22.13768 | -0.92357 | -1.75132 | -0.09583 | 21.72709 | -2.65581 | -4.17641 | 0.033636 | 0.087525 | 0 | 0 |
| 955 | sp\|Q9DBP5 | KCY_MOUS | 22 kDa | 0.99784 | 3 | 100% | Q9DBP5 | Cmpk1 | UMP-CMP k | 17.69648 | 16.82647 | 17.40337 | 14.87972 | 18.18634 | 11.29231 | 3.429234 | 0.354809 | 6.503659 | 16.04745 | 2.661851 | -4.17299 | 0.03373 | 0.087525 | 1 | 0 |
| 124 | sp\|Q7TSC1 | PRC2A_MO | 229 kDa | 0.999914 | 13 | 100% | Q7TSC1 | Prrc2a | Protein PRR | 21.8237 | 20.29848 | 21.32324 | 21.30351 | 21.97834 | 20.57564 | 0.982552 | 0.101322 | 1.863782 | 21.21715 | 2.658034 | -4.17637 | 0.033758 | 0.087525 | 0 | 0 |
| 1272 | sp\|Q8K0C9 | GMDS_MO | 42 kDa | 0.99604 | 2 | 100% | Q8K0C9 | Gmds | GDP-mann | 21.6135 | 20.78303 | 21.08911 | 20.67999 | 21.42575 | 19.87977 | 0.928523 | 0.094961 | 1.762084 | 20.91186 | 2.65141 | -4.18284 | 0.033848 | 0.087591 | 0 | 0 |

| | | | | | | | | | | | | | | | | | | | | | | | | | |
|---|---|---|---|---|---|---|---|---|---|---|---|---|---|---|---|---|---|---|---|---|---|---|---|---|---|
| 731 | sp\|Q8CGZ0 | CHERP_MO | 106 kDa | 0.998439 | 4 | 100% | Q8CGZ0 | Cherp | Calcium ho | 19.7326 | 18.91843 | 19.39718 | 18.98631 | 19.52252 | 18.5118 | 0.745254 | 0.075018 | 1.41549 | 19.17814 | 2.646533 | -4.18989 | 0.034077 | 0.088017 | 0 | 0 |
| 854 | sp\|P32921 | SYWC_MOU | 54 kDa | 0.999093 | 3 | 100% | P32921 | Wars | Tryptophar | 21.51485 | 22.31613 | 20.69197 | 22.03792 | 21.77243 | 22.54966 | -0.97482 | -1.8544 | -0.09524 | 21.81383 | -2.64207 | -4.19968 | 0.034533 | 0.089023 | 0 | 0 |
| 957 | sp\|Q99P58 | RB27B_MO | 25 kDa | 0.997785 | 3 | 33% | Q99P58 | Rab27b | Ras-related | 21.76208 | 21.26309 | 22.24039 | 20.23237 | 21.23287 | 20.50384 | 1.078678 | 0.103477 | 2.05388 | 21.20578 | 2.639661 | -4.20533 | 0.034808 | 0.089564 | 1 | 0 |
| 781 | tr\|Q3U4B1 | Q3U4B1_M | 37 kDa | 0.999788 | 3 | 67% | Q3U4B1 | Ssbp4 | Single stran | 20.36239 | 19.84935 | 21.6409 | 19.64033 | 20.25046 | 19.25902 | 1.168347 | 0.111184 | 2.225511 | 20.16707 | 2.637428 | -4.20858 | 0.034919 | 0.089678 | 1 | 0 |
| 980 | sp\|Q8BH04 | PCKGM_M | 71 kDa | 0.99726 | 3 | 100% | Q8BH04 | Pck2 | Phosphoen | 17.59617 | 18.48901 | 16.7916 | 19.04809 | 18.4791 | 19.05903 | -1.24308 | -2.36898 | -0.11719 | 18.24383 | -2.63483 | -4.21238 | 0.035048 | 0.08984 | -1 | 0 |
| 350 | sp\|Q61941 | NNTM_MO | 114 kDa | 0.999294 | 7 | 100% | Q61941 | Nnt | NAD(P) trar | 23.46375 | 24.9691 | 24.41784 | 26.04697 | 24.91549 | 25.53374 | -1.25091 | -2.38495 | -0.11687 | 24.89115 | -2.63238 | -4.21594 | 0.03517 | 0.089982 | -1 | 0 |
| 1093.1 | sp\|P61327 | MGN_MOU | 17 kDa | 0.999012 | 2 | 100% | P61327 | Magoh | Protein ma | 15.39886 | 19.18514 | 17.61647 | 19.42956 | 18.63677 | 19.17715 | -2.04658 | -3.90933 | -0.18384 | 18.24066 | -2.62196 | -4.23113 | 0.035694 | 0.091152 | -1 | 0 |
| 295 | sp\|Q9Z2K1 | K1C16_MO | 52 kDa | 0.998923 | 8 | 38% | Q9Z2K1 | Krt16 | Keratin, typ | 18.29305 | 20.6316 | 20.19063 | 20.98453 | 20.18094 | 22.0199 | -1.65714 | -3.16832 | -0.14596 | 20.38344 | -2.61694 | -4.23845 | 0.03595 | 0.091631 | -1 | 0 |
| 179 | sp\|Q04750 | TOP1_MOU | 91 kDa | 0.998937 | 11 | 100% | Q04750 | Top1 | DNA topois | 21.97346 | 23.4105 | 22.36539 | 22.74803 | 22.66678 | 23.59587 | -0.91625 | -1.75287 | -0.07964 | 22.79334 | -2.60727 | -4.24783 | 0.036078 | 0.091631 | 0 | 0 |
| 651 | sp\|Q91W50 | CSDE1_MO | 89 kDa | 0.999328 | 4 | 100% | Q91W50 | Csde1 | Cold shock | 20.14475 | 18.86037 | 19.51494 | 19.47688 | 20.01891 | 18.62934 | 0.904005 | 0.078517 | 1.729492 | 19.44086 | 2.606645 | -4.24842 | 0.036085 | 0.091631 | 0 | 0 |
| 301 | tr\|B2RT41 | B2RT41_M | 225 kDa | 0.998007 | 8 | 100% | B2RT41 | Zfc3h1 | Zinc finger, | 19.72281 | 20.72066 | 17.7719 | 20.22584 | 19.58492 | 20.47482 | -1.44723 | -2.77115 | -0.12331 | 19.75016 | -2.60872 | -4.25044 | 0.036373 | 0.09219 | -1 | 0 |
| 887 | sp\|Q9EPE9 | AT131_MO | 132 kDa | 0.998839 | 3 | 100% | Q9EPE9 | Atp13a1 | Manganese | 13.16176 | 18.05621 | 13.59603 | 14.18098 | NA | 17.3538 | -3.15144 | -6.02973 | -0.27314 | 15.26976 | -2.71397 | -4.0743 | 0.036789 | 0.092959 | -1 | 0 |
| 303 | sp\|Q9DCV7 | K2C7_MOU | 51 kDa | 0.995589 | 8 | 38% | Q9DCV7 | Krt7 | Keratin, typ | 20.06203 | 19.66068 | 20.01247 | 19.14906 | 20.61708 | 19.47183 | 0.803335 | 0.06582 | 1.54085 | 19.82886 | 2.592547 | -4.26898 | 0.036813 | 0.092959 | 0 | 0 |
| 577 | sp\|Q9R0N0 | GALK1_MO | 42 kDa | 0.99795 | 5 | 100% | Q9R0N0 | Galk1 | Galactokin | 21.59296 | 22.36245 | 21.54921 | 22.47905 | 21.12872 | 21.83593 | -0.80218 | -1.54233 | -0.06203 | 21.82472 | -2.57959 | -4.28797 | 0.037504 | 0.094439 | 0 | 0 |
| 827 | sp\|Q8BP67 | RL24_MOU | 18 kDa | 0.999366 | 3 | 100% | Q8BP67 | Rpl24 | 60S ribosor | 20.71275 | 19.88765 | 20.2485 | 19.1527 | 20.53617 | 19.91277 | 0.848101 | 0.065378 | 1.630823 | 20.07509 | 2.578935 | -4.28893 | 0.037539 | 0.094439 | 0 | 0 |
| 766 | sp\|Q8BX90 | FND3A_MO | 132 kDa | 0.997145 | 4 | 100% | Q8BX90 | Fndc3a | Fibronectir | 18.10043 | 17.85575 | 15.54876 | 18.62786 | 15.42358 | 18.30464 | -1.90516 | -3.66505 | -0.14527 | 17.31017 | -2.58344 | -4.28731 | 0.037706 | 0.094683 | -1 | 0 |
| 290 | sp\|P42932 | TCPQ_MOU | 60 kDa | 0.999068 | 8 | 100% | P42932 | Cct8 | T-complex | 22.82982 | 24.0613 | 22.47278 | 23.20813 | 22.5006 | 23.17574 | -0.88066 | -1.6984 | -0.06291 | 23.0414 | -2.56336 | -4.31185 | 0.038395 | 0.096212 | 0 | 0 |
| 1199 | sp\|Q8BML2 | OACYL_MO | 77 kDa | 0.997952 | 2 | 100% | Q8BML2 | Oacyl | O-acyltrans | 14.46419 | 15.15861 | 11.69788 | 15.35234 | 14.51149 | 16.6196 | -2.15233 | -4.15124 | -0.15342 | 14.63402 | -2.5696 | -4.30749 | 0.038456 | 0.096212 | -1 | 0 |
| 984 | sp\|Q8CIE2 | ZMIZ2_MO | 97 kDa | 0.997118 | 3 | 100% | Q8CIE2 | Zmiz2 | Zinc finger | 20.73819 | 20.30062 | 19.56023 | 22.01002 | 19.40958 | 21.97877 | -1.52714 | -2.9498 | -0.10447 | 20.66624 | -2.56169 | -4.31903 | 0.038893 | 0.097052 | -1 | 0 |
| 9 | sp\|Q99PV0 | PRP8_MOU | 274 kDa | 0.999619 | 45 | 100% | Q99PV0 | Prpf8 | Pre-mRNA- | 26.51989 | 24.63566 | 25.40605 | 25.26912 | 26.96499 | 25.38132 | 1.201608 | 0.081871 | 2.321345 | 25.69617 | 2.560927 | -4.32015 | 0.038935 | 0.097052 | 1 | 0 |
| 750.2 | tr\|A0A140 | A0A140T8 | 13 kDa | 0.997781 | 4 | 50% | A0A140T8 | Igkv12-41 | Protein Igk | 20.09583 | 22.51047 | 21.24994 | 22.65159 | 22.41439 | 24.81552 | -2.07247 | -4.01044 | -0.1345 | 22.28962 | -2.55207 | -4.33307 | 0.039431 | 0.097927 | -1 | 0 |
| 751.1 | sp\|P01654 | KV3A1_MO | 12 kDa | 0.997764 | 4 | 25% | P01654 | NA | Ig kappa ch | 20.09583 | 22.51047 | 21.24994 | 22.65159 | 22.41439 | 24.81552 | -2.07247 | -4.01044 | -0.1345 | 22.28962 | -2.55207 | -4.33307 | 0.039431 | 0.097927 | -1 | 0 |
| 144 | sp\|Q3UTJ2 | SRBS2_MO | 132 kDa | 0.999691 | 12 | 100% | Q3UTJ2 | Sorbs2 | Sorbin and | 23.3373 | 20.86913 | 21.87923 | 20.29638 | 23.36335 | 22.3805 | 1.677959 | 0.105736 | 3.250182 | 22.02098 | 2.546935 | -4.34056 | 0.039721 | 0.098467 | 1 | 0 |
| 169 | sp\|Q99JR8 | SMRD2_M | 59 kDa | 0.999372 | 11 | 91% | Q99JR8 | Smarcd2 | SWI/SNF-re | 23.43968 | 20.97937 | 22.3396 | 22.52246 | 23.50122 | 21.8694 | 1.303092 | 0.0767 | 2.529483 | 22.44195 | 2.535692 | -4.35696 | 0.040364 | 0.099878 | 1 | 0 |
| 1218 | sp\|O55022 | PGRC1_MO | 22 kDa | 0.997686 | 2 | 50% | O55022 | Pgrmc1 | Membrane | 18.78903 | 19.97469 | 15.9403 | 20.01927 | 18.39555 | 19.00039 | -1.95649 | -3.80505 | -0.10792 | 18.68654 | -2.52577 | -4.37143 | 0.04094 | 0.10112 | -1 | 0 |
| 187 | sp\|P10126 | EF1A1_MO | 50 kDa | 0.999707 | 10 | 30% | P10126 | Eef1a1 | Elongation | 24.32869 | 23.62159 | 24.18156 | 23.69618 | 24.27302 | 23.58216 | 0.627782 | 0.033005 | 1.222559 | 23.9472 | 2.512207 | -4.38672 | 0.041317 | 0.101864 | 0 | 0 |
| 41 | sp\|Q9R0I7 | YLPM1_MO | 155 kDa | 0.999814 | 22 | 100% | Q9R0I7 | Ylpm1 | YLP motif-c | 26.61066 | 24.99479 | 25.31427 | 25.07381 | 26.83904 | 25.33052 | 1.121616 | 0.053378 | 2.189854 | 25.69385 | 2.505688 | -4.40073 | 0.042133 | 0.103689 | 1 | 0 |
| 1191.2 | sp\|Q9CQC9 | SAR1B_MO | 22 kDa | 0.998052 | 2 | 100% | Q9CQC9 | Sar1b | GTP-bindin | 20.15253 | 21.06316 | 19.64633 | 20.38487 | 20.25306 | 21.82015 | -1.07209 | -2.09376 | -0.05041 | 20.55335 | -2.5042 | -4.4029 | 0.042223 | 0.103722 | -1 | 0 |
| 256 | sp\|Q99KI0 | ACON_MOU | 85 kDa | 0.99984 | 8 | 100% | Q99KI0 | Aco2 | Aconitate h | 21.84682 | 23.09235 | 22.84626 | 22.97281 | 21.96035 | 23.19137 | -0.8677 | -1.69881 | -0.03658 | 22.65166 | -2.48502 | -4.42663 | 0.042975 | 0.105378 | 0 | 0 |
| 1242 | sp\|P21278 | GNA11_MO | 42 kDa | 0.997167 | 2 | 100% | P21278 | Gna11 | Guanine nu | 16.86335 | 14.97846 | 18.33561 | 17.22075 | 17.6544 | 13.02014 | 2.544668 | 0.102554 | 4.986782 | 16.34545 | 2.486656 | -4.42848 | 0.043297 | 0.105893 | 1 | 0 |
| 794 | sp\|P01630 | KV2A6_MO | 12 kDa | 0.999602 | 3 | 33% | P01630 | NA | Ig kappa ch | 21.1021 | 20.68125 | 21.18696 | 20.40725 | 21.12151 | 20.41317 | 0.636303 | 0.025267 | 1.247339 | 20.81871 | 2.478552 | -4.43602 | 0.04337 | 0.105893 | 0 | 0 |
| 1280 | sp\|Q8K0L9 | ZBT20_MO | 81 kDa | 0.995702 | 2 | 100% | Q8K0L9 | Zbtb20 | Zinc finger | 20.26318 | 18.44949 | 22.24132 | 20.89245 | 21.43402 | 18.80873 | 1.929284 | 0.076287 | 3.782281 | 20.3482 | 2.48469 | -4.43135 | 0.043419 | 0.105893 | 1 | 0 |
| 422 | sp\|O08538 | ANGP1_MO | 58 kDa | 0.999433 | 6 | 100% | O08538 | Angpt1 | Angiopoiet | 21.66208 | 20.81513 | 22.21574 | 21.05985 | 21.15419 | 19.82284 | 1.111399 | 0.041083 | 2.181715 | 21.12164 | 2.478041 | -4.44105 | 0.043835 | 0.106715 | 1 | 0 |
| 33 | sp\|Q99104 | MYO5A_MO | 216 kDa | 0.999369 | 24 | 100% | Q99104 | Myo5a | Unconvent | 23.16301 | 23.89178 | 23.11372 | 23.74498 | 23.36102 | 23.89655 | -0.63186 | -1.24114 | -0.02257 | 23.52851 | -2.46829 | -4.45105 | 0.044016 | 0.106964 | 0 | 0 |
| 475 | sp\|Q80SY5 | PR38B_MO | 64 kDa | 0.997437 | 6 | 100% | Q80SY5 | Prpf38b | Pre-mRNA- | 19.65407 | 21.97681 | 19.6132 | 20.7489 | 20.77621 | 20.90881 | -1.19701 | -2.35397 | -0.04005 | 20.613 | -2.46905 | -4.45416 | 0.044404 | 0.107713 | -1 | 0 |
| 624 | sp\|Q9QYH6 | MAGD1_M | 86 kDa | 0.999491 | 4 | 100% | Q9QYH6 | Maged1 | Melanoma- | 20.44559 | 18.32565 | 20.25479 | 19.83698 | 20.48935 | 19.70369 | 1.107803 | 0.035008 | 2.180599 | 19.84268 | 2.464316 | -4.46106 | 0.044706 | 0.108018 | 1 | 0 |
| 668.1 | sp\|P19253 | RL13A_MO | 23 kDa | 0.99914 | 4 | 100% | P19253 | Rpl13a | 60S ribosor | 23.91459 | 23.18001 | 23.71477 | 23.09677 | 23.68246 | 23.16763 | 0.622468 | 0.019363 | 1.225572 | 23.45937 | 2.456548 | -4.46825 | 0.044769 | 0.108018 | 0 | 0 |
| 668.2 | tr\|A0A1B0 | A0A1B0GS | 51 kDa | 0.99914 | 4 | 100% | A0A1B0GS | Gm45713 | Predicted g | 23.91459 | 23.18001 | 23.71477 | 23.09677 | 23.68246 | 23.16763 | 0.622468 | 0.019363 | 1.225572 | 23.45937 | 2.456548 | -4.46825 | 0.044769 | 0.108018 | 0 | 0 |
| 1142.2 | tr\|W4VSN7 | W4VSN7_N | 18 kDa | 0.998518 | 2 | 100% | W4VSN7 | Gm3550 | 60S ribosor | 21.53802 | 19.7664 | 22.0227 | 20.84075 | 21.50734 | 21.23563 | 1.075095 | 0.031224 | 2.118966 | 21.15181 | 2.457824 | -4.47053 | 0.045125 | 0.108491 | 1 | 0 |
| 1143 | sp\|P41105 | RL28_MOU | 16 kDa | 0.998509 | 2 | 100% | P41105 | Rpl28 | 60S ribosor | 21.53802 | 19.7664 | 22.0227 | 20.84075 | 21.50734 | 21.23563 | 1.075095 | 0.031224 | 2.118966 | 21.15181 | 2.457824 | -4.47053 | 0.045125 | 0.108491 | 1 | 0 |
| 300 | sp\|Q6NV83 | SR140_MO | 118 kDa | 0.998301 | 8 | 100% | Q6NV83 | U2surp | U2 snRNP-a | 23.30277 | 23.86703 | 22.72884 | 23.66706 | 22.96029 | 23.55779 | -0.69999 | -1.38126 | -0.01872 | 23.3473 | -2.44553 | -4.48438 | 0.045487 | 0.109035 | 0 | 0 |
| 1148 | sp\|Q9QX66 | DPF1_MOU | 44 kDa | 0.998488 | 2 | 50% | Q9QX66 | Dpf1 | Zinc finger | 19.46792 | 19.66093 | 19.06247 | 19.91646 | 19.11227 | 20.19687 | -0.71053 | -1.40218 | -0.01889 | 19.56949 | -2.44515 | -4.48495 | 0.045512 | 0.109035 | 0 | 0 |
| 238 | sp\|O08553 | DPYL2_MO | 62 kDa | 0.999407 | 9 | 67% | O08553 | Dpysl2 | Dihydropyr | 22.47915 | 22.79795 | 21.95698 | 22.93316 | 22.20652 | 22.92139 | -0.66995 | -1.32271 | -0.01719 | 22.54919 | -2.4428 | -4.48839 | 0.045667 | 0.109213 | 0 | 0 |
| 891 | sp\|Q8BG32 | PSD11_MO | 47 kDa | 0.998773 | 3 | 100% | Q8BG32 | Psmd11 | 26S protea | 22.34552 | 19.62844 | 19.25061 | 18.66524 | 22.13427 | 19.29053 | 2.048736 | 0.048762 | 4.04871 | 20.2191 | 2.444625 | -4.48977 | 0.045988 | 0.109594 | 1 | 0 |
| 892 | sp\|Q80WC | AGFG2_MO | 49 kDa | 0.99877 | 3 | 100% | Q80WC7 | Agfg2 | Arf-GAP do | 22.34552 | 19.62844 | 19.25061 | 18.66524 | 22.13427 | 19.29053 | 2.048736 | 0.048762 | 4.04871 | 20.2191 | 2.444625 | -4.48977 | 0.045988 | 0.109594 | 1 | 0 |
| 92 | sp\|P14685 | PSMD3_M | 61 kDa | 0.999619 | 16 | 100% | P14685 | Psmd3 | 26S protea | 23.61728 | 24.04203 | 23.26095 | 23.82918 | 23.25207 | 24.38788 | -0.7096 | -1.40298 | -0.01621 | 23.73157 | -2.43579 | -4.49865 | 0.046131 | 0.109744 | 0 | 0 |
| 961 | sp\|Q64378 | FKBP5_MO | 51 kDa | 0.997747 | 3 | 100% | Q64378 | Fkbp5 | Peptidyl-pr | 21.81719 | 19.52478 | 21.06636 | 20.02997 | 21.8661 | 21.34804 | 1.282288 | 0.028515 | 2.536062 | 20.94207 | 2.440716 | -4.49547 | 0.046246 | 0.109825 | 1 | 0 |
| 110 | sp\|Q7TQG1 | PKHA6_MO | 131 kDa | 0.999124 | 15 | 100% | Q7TQG1 | Plekha6 | Pleckstrin h | 23.65664 | 21.53064 | 22.01761 | 22.06882 | 23.09271 | 21.79914 | 1.122787 | 0.023842 | 2.221731 | 22.36093 | 2.438216 | -4.49911 | 0.046413 | 0.109911 | 1 | 0 |
| 881 | sp\|Q91V41 | RAB14_MO | 24 kDa | 0.998909 | 3 | 67% | Q91V41 | Rab14 | Ras-related | 20.1225 | 20.80785 | 20.42186 | 21.48924 | 20.29558 | 20.80112 | -0.75275 | -1.48972 | -0.01578 | 20.65636 | -2.43111 | -4.5055 | 0.046445 | 0.109911 | 0 | 0 |
| 1128 | sp\|P35486 | ODPA_MOU | 43 kDa | 0.998636 | 2 | 100% | P35486 | Pdha1 | Pyruvate de | 21.51543 | 20.80337 | 21.10743 | 20.53453 | 21.93836 | 20.8999 | 0.774473 | 0.015479 | 1.533466 | 21.13317 | 2.428672 | -4.50906 | 0.046608 | 0.110037 | 0 | 0 |
| 1162 | tr\|A0A075 | A0A075B5 | 13 kDa | 0.99832 | 2 | 50% | A0A075B5 | Igkv3-1 | Immunoglo | 19.78814 | 19.06371 | 20.4904 | 19.56496 | 19.70587 | 18.55576 | 0.933328 | 0.018131 | 1.848526 | 19.52814 | 2.431786 | -4.50729 | 0.04671 | 0.110037 | 0 | 0 |
| 544 | sp\|O08576 | RUN3A_MO | 50 kDa | 0.999109 | 5 | 100% | O08576 | Rundc3a | RUN domai | 19.70988 | 23.13447 | 19.16494 | 21.78753 | 22.66871 | 23.89923 | -2.4259 | -4.80614 | -0.04565 | 21.72746 | -2.43221 | -4.50786 | 0.046815 | 0.110037 | -1 | 0 |
| 518 | sp\|P09103 | PDIA1_MO | 57 kDa | 0.999354 | 5 | 100% | P09103 | P4hb | Protein dis | 20.44397 | 21.02852 | 20.36413 | 21.43987 | 20.71194 | 21.07258 | -0.67364 | -1.3352 | -0.01209 | 20.8435 | -2.42361 | -4.51647 | 0.046951 | 0.110037 | 0 | 0 |
| 688 | sp\|Q9WU4 | NCOR2_M | 270 kDa | 0.999006 | 4 | 100% | Q9WU42 | Ncor2 | Nuclear rec | 18.56896 | 19.38305 | 18.10729 | 20.49671 | 19.62972 | 20.01871 | -1.1975 | -2.37403 | -0.02098 | 19.36741 | -2.42899 | -4.51256 | 0.047032 | 0.110037 | -1 | 0 |
| 900 | tr\|E9QAP7 | E9QAP7_M | 105 kDa | 0.998684 | 3 | 100% | E9QAP7 | Taf4 | TATA-box-b | 19.83317 | 20.60674 | 19.70936 | 21.62335 | 20.98159 | 22.32683 | -1.34427 | -2.66536 | -0.02317 | 20.84684 | -2.4283 | -4.51357 | 0.047079 | 0.110037 | -1 | 0 |
| 681 | sp\|Q9D0W | PPIL1_MOU | 18 kDa | 0.999051 | 4 | 100% | Q9D0W5 | Ppil1 | Peptidyl-pr | 21.37567 | 20.23097 | 20.71761 | 20.84909 | 21.91929 | 19.55584 | 1.125558 | 0.019056 | 2.232059 | 20.77474 | 2.427541 | -4.51467 | 0.04713 | 0.110037 | 1 | 0 |
| 297 | sp\|Q01853 | TERA_MOU | 89 kDa | 0.998647 | 8 | 100% | Q01853 | Vcp | Transitiona | 22.64674 | 23.74505 | 23.27683 | 23.41815 | 22.3216 | 25.62771 | -1.51524 | -3.00499 | -0.0255 | 23.50601 | -2.42729 | -4.51504 | 0.047147 | 0.110037 | -1 | 0 |
| 275 | sp\|Q64012 | RALY_MOU | 33 kDa | 0.999479 | 8 | 100% | Q64012 | Raly | RNA-bindir | 24.25149 | 23.26493 | 23.75608 | 23.47938 | 24.43591 | 23.54171 | 0.719155 | 0.008507 | 1.429802 | 23.78825 | 2.408623 | -4.5384 | 0.04798 | 0.111635 | 0 | 0 |
| 594.1 | sp\|P01661 | KV3A9_MO | 14 kDa | 0.999744 | 4 | 50% | P01661 | NA | Ig kappa ch | 17.65179 | 21.03122 | 20.27979 | 20.51485 | 19.21963 | 20.36423 | -1.58636 | -3.15483 | -0.0179 | 19.84358 | -2.41367 | -4.53487 | 0.048079 | 0.111635 | -1 | 0 |
| 594.2 | sp\|P01662 | KV3AA_MO | 12 kDa | 0.999744 | 4 | 50% | P01662 | NA | Ig kappa ch | 17.65179 | 21.03122 | 20.27979 | 20.51485 | 19.21963 | 20.36423 | -1.58636 | -3.15483 | -0.0179 | 19.84358 | -2.41367 | -4.53487 | 0.048079 | 0.111635 | -1 | 0 |
| 98 | sp\|Q80U23 | SNPH_MOU | 54 kDa | 0.999775 | 15 | 87% | Q80U23 | Snph | Syntaphilin | 24.56258 | 20.65571 | 23.42978 | 21.67479 | 24.41434 | 23.97825 | 2.032649 | 0.019173 | 4.046125 | 23.11924 | 2.409166 | -4.54144 | 0.048392 | 0.112169 | 1 | 0 |
| 635 | sp\|P61027 | RAB10_MO | 23 kDa | 0.99942 | 4 | 50% | P61027 | Rab10 | Ras-related | 20.99505 | 20.28275 | 20.83494 | 19.72103 | 20.90705 | 20.52256 | 0.7369 | 0.002363 | 1.471436 | 20.5439 | 2.387788 | -4.56887 | 0.049449 | 0.114084 | 0 | 0 |
| 907 | sp\|Q68FF6 | GIT1_MOU | 85 kDa | 0.998593 | 3 | 100% | Q68FF6 | Git1 | ARF GTPase | 20.14337 | 19.68871 | 20.58869 | 18.81858 | 21.31187 | 20.16075 | 1.125295 | 0.003098 | 2.247492 | 20.11866 | 2.39303 | -4.56494 | 0.049528 | 0.114084 | 1 | 0 |
| 1288 | sp\|Q99KY4 | GAK_MOUS | 144 kDa | 0.99518 | 2 | 100% | Q99KY4 | Gak | Cyclin-G-as | 19.7152 | 20.58733 | 20.4211 | 22.09889 | 20.70414 | 23.73377 | -1.85985 | -3.71486 | -0.00484 | 21.21007 | -2.39266 | -4.56548 | 0.049554 | 0.114084 | -1 | 0 |
| 1289 | tr\|A0A140 | A0A140T8 | 13 kDa | 0.995165 | 2 | 50% | A0A140T8 | Igkv1-122 | Immunoglo | 19.7152 | 20.58733 | 20.4211 | 22.09889 | 20.70414 | 23.73377 | -1.85985 | -3.71486 | -0.00484 | 21.21007 | -2.39266 | -4.56548 | 0.049554 | 0.114084 | -1 | 0 |
| 1179 | sp\|O89079 | COPE_MOU | 35 kDa | 0.998156 | 2 | 100% | O89079 | Cope | Coatomer s | 19.07455 | 21.38768 | 20.87227 | 21.07544 | 19.69889 | 20.73699 | -1.1848 | -2.36972 | 0.000114 | 20.4743 | -2.38621 | -4.57487 | 0.050016 | 0.114541 | -1 | 0 |
| 1202 | sp\|Q8BTV2 | CPSF7_MO | 52 kDa | 0.997928 | 2 | 100% | Q8BTV2 | Cpsf7 | Cleavage ar | 20.75181 | 20.37373 | 21.80442 | 20.05874 | 20.7041 | 20.13299 | 0.898292 | -0.00017 | 1.796752 | 20.63763 | 2.383485 | -4.57738 | 0.050032 | 0.114541 | 0 | 0 |
| 259 | sp\|Q921F2 | TADBP_MO | 45 kDa | 0.999752 | 8 | 100% | Q921F2 | Tardbp | TAR DNA-bi | 24.50671 | 23.36841 | 23.37408 | 22.91112 | 24.68604 | 23.31289 | 0.99147 | -0.00039 | 1.983329 | 23.69321 | 2.385503 | -4.5759 | 0.050068 | 0.114541 | 0 | 0 |
| 36 | sp\|E1U8D0 | SOGA1_MO | 159 kDa | 0.99968 | 23 | 91% | E1U8D0 | Soga1 | Protein SO | 23.38885 | 22.81849 | 22.73312 | 22.3781 | 23.91569 | 21.72737 | 1.037901 | -0.00087 | 2.076668 | 22.82694 | 2.384452 | -4.57743 | 0.050143 | 0.114541 | 1 | 0 |
| 85 | sp\|Q9CWF | TBB2B_MO | 50 kDa | 0.998684 | 17 | 6% | Q9CWF2 | Tubb2b | Tubulin bet | 20.83364 | 19.967 | 21.31004 | 20.26951 | 20.2755 | 19.1449 | 1.012591 | -0.00103 | 2.026213 | 20.3001 | 2.384014 | -4.57807 | 0.050175 | 0.114541 | 1 | 0 |
| 193 | sp\|O55201 | SPT5H_MO | 121 kDa | 0.999629 | 10 | 100% | O55201 | Supt5h | Transcripti | 24.24779 | 24.5721 | 22.28038 | 25.1202 | 24.24871 | 25.2634 | -1.39294 | -2.79075 | 0.004878 | 24.28876 | -2.37811 | -4.58666 | 0.050603 | 0.115324 | -1 | 0 |
| 1066 | sp\|Q9DAW | PRP4_MOU | 58 kDa | 0.999236 | 2 | 100% | Q9DAW6 | Prpf4 | U4/U6 sma | 18.80277 | 18.89969 | 19.31853 | 22.16425 | 18.67556 | 21.70541 | -1.99083 | -3.99383 | 0.012164 | 19.9277 | -2.37195 | -4.59563 | 0.051054 | 0.116157 | -1 | 0 |
| 997 | sp\|Q9CQY5 | MAGT1_M | 38 kDa | 0.996064 | 3 | 100% | Q9CQY5 | Magt1 | Magnesium | 19.52609 | 19.13462 | 19.692 | 18.33757 | 19.02055 | 18.0906 | 0.891949 | -0.01032 | 1.794217 | 18.9669 | 2.356669 | -4.61647 | 0.052008 | 0.11774 | 0 | 0 |
| 356.1 | sp\|P15864 | H12_MOUS | 21 kDa | 0.999195 | 7 | 29% | P15864 | Hist1h1c | Histone H1 | 23.14631 | 22.63602 | 23.42859 | 22.65585 | 23.57128 | 22.92237 | 0.643983 | -0.00752 | 1.295483 | 23.06007 | 2.352665 | -4.62019 | 0.05203 | 0.11774 | 0 | 0 |
| 356.2 | sp\|P43277 | H13_MOUS | 22 kDa | 0.999195 | 7 | 29% | P43277 | Hist1h1d | Histone H1 | 23.14631 | 22.63602 | 23.42859 | 22.65585 | 23.57128 | 22.92237 | 0.643983 | -0.00752 | 1.295483 | 23.06007 | 2.352665 | -4.62019 | 0.05203 | 0.11774 | 0 | 0 |
| 722 | sp\|Q99KK2 | NEUA_MOU | 48 kDa | 0.998657 | 4 | 100% | Q99KK2 | Cmas | N-acylneur | 21.61629 | 20.1071 | 21.84499 | 21.25798 | 21.27918 | 20.71487 | 0.886838 | -0.01071 | 1.78439 | 21.13673 | 2.355475 | -4.61821 | 0.052097 | 0.11774 | 0 | 0 |
| 971 | sp\|Q9D394 | RUFY3_MO | 53 kDa | 0.997581 | 3 | 100% | Q9D394 | Rufy3 | Protein RU | 18.15332 | 21.67238 | 16.86989 | 18.88303 | NA | 19.03884 | -2.35315 | -4.73929 | 0.032991 | 18.92349 | -2.44447 | -4.44132 | 0.052314 | 0.118033 | -1 | 0 |
| 289 | sp\|Q91VI7 | RINI_MOUS | 50 kDa | 0.999122 | 8 | 100% | Q91VI7 | Rnh1 | Ribonuclea | 22.87007 | 23.23037 | 22.4386 | 23.09264 | 21.80077 | 23.20573 | -0.80643 | -1.62633 | 0.013471 | 22.77303 | -2.34114 | -4.63708 | 0.052916 | 0.119193 | 0 | 0 |
| 754 | sp\|Q9R233 | TPSN_MOU | 50 kDa | 0.997633 | 4 | 100% | Q9R233 | Tapbp | Tapasin | 19.3476 | 20.46715 | 15.40894 | 19.93171 | 18.25698 | 19.42601 | -2.27045 | -4.58034 | 0.039446 | 18.8064 | -2.34569 | -4.63383 | 0.053024 | 0.119237 | -1 | 0 |
| 309 | sp\|Q60932 | VDAC1_MO | 32 kDa | 0.99972 | 7 | 100% | Q60932 | Vdac1 | Voltage-dep | 23.46704 | 22.68136 | 24.19251 | 22.91082 | 23.37237 | 23.11738 | 0.774119 | -0.01397 | 1.562205 | 23.29025 | 2.337947 | -4.64168 | 0.053153 | 0.119331 | 0 | 0 |
| 433 | sp\|Q9D051 | ODPB_MOU | 39 kDa | 0.999287 | 6 | 100% | Q9D051 | Pdhb | Pyruvate de | 22.47638 | 23.53241 | 22.59785 | 23.1129 | 22.4291 | 22.88218 | -0.67472 | -1.36272 | 0.01329 | 22.83847 | -2.33415 | -4.64722 | 0.053447 | 0.119791 | 0 | 0 |
| 374 | sp\|Q9Z2I9 | SUCB1_MO | 50 kDa | 0.998806 | 7 | 100% | Q9Z2I9 | Sucla2 | Succinate-- | 22.6028 | 24.62411 | 22.48925 | 23.09585 | 22.63647 | 23.18563 | -1.05902 | -2.14532 | 0.02728 | 23.10569 | -2.32651 | -4.6617 | 0.054511 | 0.121975 | -1 | 0 |
| 387 | sp\|B2RQC6 | PYR1_MOU | 243 kDa | 0.99735 | 7 | 100% | B2RQC6 | Cad | CAD protei | 20.18287 | 21.52596 | 19.64392 | 20.26748 | 20.4574 | 21.93899 | -1.14941 | -2.3297 | 0.030872 | 20.66944 | -2.32402 | -4.66532 | 0.054707 | 0.122213 | -1 | 0 |
| 676 | sp\|Q9EPU4 | CPSF1_MO | 161 kDa | 0.999086 | 4 | 100% | Q9EPU4 | Cpsf1 | Cleavage ar | 20.60048 | 19.8573 | 19.9053 | 19.49584 | 20.3059 | 18.54551 | 0.971007 | -0.02799 | 1.970003 | 19.78506 | 2.319579 | -4.67177 | 0.055059 | 0.122724 | 0 | 0 |
| 1275 | sp\|Q7TNI2 | TM59L_MO | 38 kDa | 0.995772 | 2 | 100% | Q7TNI2 | Tmem59l | Transmemb | 17.12223 | 15.45169 | 17.60215 | 16.91581 | 17.53475 | NA | 1.235958 | -0.03787 | 2.509787 | 16.92533 | 2.405047 | -4.49528 | 0.055117 | 0.122724 | 0 | 0 |
| 759 | tr\|A0A075 | A0A075B5 | 13 kDa | 0.997521 | 4 | 50% | A0A075B5 | Ighv7-2 | Immunoglo | 20.51741 | 18.7803 | 19.95402 | 20.36438 | 21.16758 | 18.33081 | 1.387845 | -0.04441 | 2.8201 | 19.85242 | 2.312445 | -4.68213 | 0.055629 | 0.123515 | 0 | 0 |
| 792 | sp\|Q3V3Q7 | PACS2_MO | 95 kDa | 0.999629 | 3 | 67% | Q3V3Q7 | Pacs2 | Phosphofu | 19.66859 | 19.39897 | 19.95115 | 18.58753 | 19.50722 | 18.91582 | 0.741546 | -0.02375 | 1.506841 | 19.33821 | 2.306268 | -4.68789 | 0.055654 | 0.123515 | 0 | 0 |
| 409.1 | sp\|P61205 | ARF3_MOU | 21 kDa | 0.999587 | 6 | 50% | P61205 | Arf3 | ADP-ribosy | 22.93053 | 22.17259 | 23.22057 | 22.59268 | 22.81393 | 22.2623 | 0.645819 | -0.02293 | 1.314569 | 22.66544 | 2.298519 | -4.69918 | 0.056284 | 0.124505 | 0 | 0 |
| 409.2 | sp\|P84078 | ARF1_MOU | 21 kDa | 0.999587 | 6 | 50% | P84078 | Arf1 | ADP-ribosy | 22.93053 | 22.17259 | 23.22057 | 22.59268 | 22.81393 | 22.2623 | 0.645819 | -0.02293 | 1.314569 | 22.66544 | 2.298519 | -4.69918 | 0.056284 | 0.124505 | 0 | 0 |
| 1255 | tr\|G3X972 | G3X972_M | 119 kDa | 0.996704 | 2 | 100% | G3X972 | Sec24c | SEC24 relat | 16.35178 | 18.29013 | 17.49884 | 19.74665 | 18.02601 | 18.05301 | -1.40439 | -2.8617 | 0.052929 | 17.9944 | -2.29977 | -4.70054 | 0.056657 | 0.125125 | 0 | 0 |
| 642 | sp\|P61372 | ISL1_MOUS | 39 kDa | 0.999383 | 4 | 100% | P61372 | Isl1 | Insulin gen | 21.44796 | 19.90153 | 20.70495 | 18.87481 | 20.57345 | 20.54201 | 1.136004 | -0.04445 | 2.316459 | 20.34079 | 2.29658 | -4.70516 | 0.056918 | 0.125498 | 0 | 0 |
| 947 | sp\|Q80VI1 | TRI56_MOU | 80 kDa | 0.997927 | 3 | 100% | Q80VI1 | Trim56 | E3 ubiquiti | 19.90851 | 19.84905 | 19.02738 | 21.15116 | 19.64108 | 20.53434 | -0.98586 | -2.01146 | 0.039744 | 20.01859 | -2.29396 | -4.70896 | 0.057134 | 0.125769 | 0 | 0 |
| 248 | sp\|Q6ZWN | RS9_MOUS | 23 kDa | 0.998829 | 9 | 100% | Q6ZWN5 | Rps9 | 40S ribosor | 25.33911 | 24.4305 | 25.31446 | 24.9012 | 25.10763 | 24.54685 | 0.627551 | -0.0273 | 1.282399 | 24.93996 | 2.280916 | -4.72482 | 0.057743 | 0.126438 | 0 | 0 |
| 643 | sp\|O70305 | ATX2_MOU | 136 kDa | 0.999382 | 4 | 100% | O70305 | Atxn2 | Ataxin-2 | 23.61795 | 24.8761 | 24.78795 | 25.92741 | 23.7155 | 30.00502 | -2.89571 | -5.91873 | 0.127315 | 25.48832 | -2.28594 | -4.72061 | 0.0578 | 0.126438 | 0 | 0 |
| 578 | sp\|P35441 | TSP1_MOU | 130 kDa | 0.997925 | 5 | 100% | P35441 | Thbs1 | Thrombosp | 19.30392 | 18.31727 | 19.15528 | 18.17946 | 19.74027 | 19.17603 | 0.842234 | -0.0373 | 1.721768 | 18.97871 | 2.282833 | -4.72389 | 0.05787 | 0.126438 | 0 | 0 |
| 861 | sp\|Q8BK64 | AHSA1_MO | 38 kDa | 0.999048 | 3 | 100% | Q8BK64 | Ahsa1 | Activator o | 22.09504 | 21.6814 | 21.98867 | 21.1492 | 22.26285 | 21.58001 | 0.645314 | -0.02855 | 1.319181 | 21.79286 | 2.279277 | -4.7272 | 0.05788 | 0.126438 | 0 | 0 |
| 748 | sp\|Q6DID3 | SCAF8_MO | 140 kDa | 0.997845 | 4 | 25% | Q6DID3 | Scaf8 | SR-related | 20.36778 | 22.42936 | 22.17318 | 23.78716 | 19.41568 | 21.51597 | -1.92528 | -3.9363 | 0.085743 | 21.61486 | -2.28469 | -4.72241 | 0.057904 | 0.126438 | 0 | 0 |
| 117 | sp\|Q03265 | ATPA_MOU | 60 kDa | 0.999674 | 14 | 100% | Q03265 | Atp5f1a | ATP syntha | 27.64126 | 27.00649 | 27.78058 | 26.94099 | 27.46944 | 27.16288 | 0.593641 | -0.02717 | 1.214454 | 27.33361 | 2.275954 | -4.73204 | 0.058161 | 0.126794 | 0 | 0 |
| 804 | sp\|Q4QQM | P5I11_MO | 21 kDa | 0.999496 | 3 | 100% | Q4QQM4 | Trp53i11 | Tumor prot | 13.88303 | 15.97338 | 14.62379 | 14.68579 | NA | 15.76436 | -1.2211 | -2.50283 | 0.060635 | 14.98607 | -2.36148 | -4.55494 | 0.058401 | 0.127114 | 0 | 0 |
| 783 | tr\|A0A075 | A0A075B5 | 10 kDa | 0.999719 | 3 | 100% | A0A075B5 | Igkv4-63 | Immunoglo | 19.42727 | 21.06022 | 21.05137 | 22.2924 | 21.26137 | 22.81953 | -1.47738 | -3.02855 | 0.073783 | 21.31869 | -2.27293 | -4.73947 | 0.058897 | 0.127808 | 0 | 0 |
| 698 | sp\|Q91V12 | BACH_MOU | 43 kDa | 0.998903 | 4 | 100% | Q91V12 | Acot7 | Cytosolic a | 22.18701 | 21.56343 | 21.69473 | 21.30658 | 22.27314 | 21.31347 | 0.657133 | -0.03294 | 1.347204 | 21.72306 | 2.266522 | -4.74576 | 0.058964 | 0.127808 | 0 | 0 |
| 610 | sp\|Q5SUF2 | LC7L3_MO | 51 kDa | 0.999627 | 4 | 100% | Q5SUF2 | Luc7l3 | Luc7-like p | 21.09798 | 21.80842 | 20.7883 | 21.59458 | 21.53058 | 22.33529 | -0.77381 | -1.58657 | 0.038954 | 21.52586 | -2.26617 | -4.74634 | 0.059003 | 0.127808 | 0 | 0 |
| 326 | sp\|P26039 | TLN1_MOU | 270 kDa | 0.999568 | 7 | 100% | P26039 | Tln1 | Talin-1 | 20.72182 | 21.70198 | 21.38482 | 21.37281 | 20.78427 | 22.10382 | -0.76257 | -1.56408 | 0.03895 | 21.34492 | -2.26447 | -4.74875 | 0.05914 | 0.127901 | 0 | 0 |
| 48 | sp\|Q61656 | DDX5_MOU | 69 kDa | 0.999587 | 20 | 60% | Q61656 | Ddx5 | Probable AT | 24.37106 | 23.50322 | 24.3068 | 23.23213 | 24.07786 | 23.96358 | 0.685599 | -0.03733 | 1.408529 | 23.90911 | 2.257227 | -4.75928 | 0.059766 | 0.129049 | 0 | 0 |
| 758 | sp\|P62196 | PRS8_MOU | 46 kDa | 0.997543 | 4 | 100% | P62196 | Psmc5 | 26S protea | 19.3257 | 20.27089 | 17.8908 | 18.97265 | 16.0805 | 19.59572 | -1.84742 | -3.80081 | 0.105969 | 18.68938 | -2.25698 | -4.76258 | 0.060271 | 0.129801 | 0 | 0 |
| 1121 | sp\|P27546 | MAP4_MOU | 117 kDa | 0.998723 | 2 | 100% | P27546 | Map4 | Microtubu | 19.63936 | 19.57015 | 19.57633 | 17.9499 | 19.49861 | 17.39109 | 1.267717 | -0.07295 | 2.608388 | 18.93757 | 2.256583 | -4.76315 | 0.060306 | 0.129801 | 0 | 0 |
| 1161 | sp\|Q8VEK0 | CC50A_MO | 41 kDa | 0.998326 | 2 | 100% | Q8VEK0 | Tmem30a | Cell cycle c | 18.2963 | 17.94929 | 18.50579 | 17.21228 | 18.18613 | 17.75457 | 0.690692 | -0.04123 | 1.422612 | 17.98406 | 2.246065 | -4.7755 | 0.060744 | 0.130537 | 0 | 0 |
| 49 | sp\|P07724 | ALBU_MOU | 69 kDa | 0.999586 | 20 | 100% | P07724 | Alb | Serum albu | 25.14534 | 25.54977 | 24.56097 | 26.08565 | 25.69402 | 26.99156 | -1.07555 | -2.21818 | 0.067077 | 25.67122 | -2.24635 | -4.77797 | 0.061205 | 0.13132 | 0 | 0 |
| 105 | sp\|P20152 | VIME_MOU | 54 kDa | 0.999598 | 15 | 93% | P20152 | Vim | Vimentin | 23.75672 | 23.50756 | 23.68261 | 26.28432 | 23.55944 | 26.29925 | -1.69745 | -3.50863 | 0.113727 | 24.51498 | -2.23659 | -4.79209 | 0.062075 | 0.132976 | 0 | 0 |
| 896 | sp\|P62245 | RS15A_MO | 15 kDa | 0.998735 | 3 | 100% | P62245 | Rps15a | 40S ribosor | 21.14847 | 20.39999 | 20.74654 | 19.2188 | 19.97439 | 18.54891 | 1.233898 | -0.08482 | 2.552619 | 20.00618 | 2.232939 | -4.79737 | 0.062404 | 0.133259 | 0 | 0 |
| 897 | sp\|Q8R1A4 | DOCK7_MO | 241 kDa | 0.998714 | 3 | 100% | Q8R1A4 | Dock7 | Dedicator c | 21.14847 | 20.39999 | 20.74654 | 19.2188 | 19.97439 | 18.54891 | 1.233898 | -0.08482 | 2.552619 | 20.00618 | 2.232939 | -4.79737 | 0.062404 | 0.133259 | 0 | 0 |
| 125 | sp\|P16858 | G3P_MOUS | 36 kDa | 0.999792 | 13 | 85% | P16858 | Gapdh | Glyceraldel | 28.16674 | 27.56843 | 28.26585 | 27.57199 | 28.00266 | 27.61463 | 0.560067 | -0.0435 | 1.163636 | 27.86505 | 2.208585 | -4.8299 | 0.064149 | 0.136665 | 0 | 0 |
| 424 | sp\|Q9ERI2 | RB27A_MO | 25 kDa | 0.999413 | 6 | 67% | Q9ERI2 | Rab27a | Ras-related | 23.33434 | 22.84304 | 23.4829 | 22.62545 | 22.95118 | 22.32735 | 0.657527 | -0.05125 | 1.366301 | 22.92738 | 2.208042 | -4.83069 | 0.0642 | 0.136665 | 0 | 0 |
| 1282 | sp\|Q8VI75 | IPO4_MOU | 119 kDa | 0.995585 | 2 | 100% | Q8VI75 | Ipo4 | Importin-4 | 16.87967 | 18.7431 | 15.36351 | 24.20768 | 16.46692 | 17.10444 | -3.78171 | -7.87076 | 0.307342 | 18.12755 | -2.20707 | -4.83475 | 0.064785 | 0.137693 | 0 | 0 |
| 1268 | sp\|Q9ERS2 | NDUAD_M | 17 kDa | 0.996241 | 2 | 100% | Q9ERS2 | Ndufa13 | NADH dehy | 16.35494 | 18.75895 | 17.40005 | 19.32248 | 18.3782 | 17.97596 | -1.30807 | -2.72885 | 0.112713 | 18.03177 | -2.19712 | -4.84912 | 0.065725 | 0.139458 | 0 | 0 |
| 837 | sp\|Q9D819 | IPYR_MOUS | 33 kDa | 0.999266 | 3 | 100% | Q9D819 | Ppa1 | Inorganic p | 21.30183 | 18.63942 | 20.36794 | 20.15095 | 21.9805 | 20.65114 | 1.402921 | -0.12232 | 2.928164 | 20.5153 | 2.195051 | -4.8521 | 0.065922 | 0.139458 | 0 | 0 |
| 926 | sp\|P01795 | HVM26_M | 16 kDa | 0.998317 | 3 | 67% | P01795 | NA | Ig heavy ch | 19.84225 | 22.13909 | 20.03015 | 21.77417 | 19.08465 | 19.6113 | -1.5225 | -3.1778 | 0.132798 | 20.4136 | -2.19499 | -4.8522 | 0.065928 | 0.139458 | 0 | 0 |
| 659 | sp\|Q8K2C9 | HACD3_MO | 43 kDa | 0.999233 | 4 | 100% | Q8K2C9 | Hacd3 | Very-long-c | 21.67203 | 19.79713 | 20.94981 | 21.38432 | 21.63156 | 19.32649 | 1.248487 | -0.10953 | 2.606505 | 20.79356 | 2.193961 | -4.85368 | 0.066026 | 0.139458 | 0 | 0 |
| 737 | sp\|B1AZI6 | THOC2_MO | 183 kDa | 0.998344 | 4 | 100% | B1AZI6 | Thoc2 | THO compl | 14.35192 | 16.63941 | 13.08975 | 15.25741 | 15.14647 | 14.98837 | -1.43235 | -2.9972 | 0.132508 | 14.91222 | -2.18436 | -4.86752 | 0.066951 | 0.141117 | 0 | 0 |
| 1029 | sp\|P61969 | LMO4_MOU | 18 kDa | 0.999551 | 2 | 100% | P61969 | Lmo4 | LIM domai | 20.35958 | 19.44375 | 19.82501 | 19.74339 | 20.07819 | 18.65762 | 0.806004 | -0.07475 | 1.686757 | 19.68459 | 2.181612 | -4.87049 | 0.06702 | 0.141117 | 0 | 0 |
| 674 | sp\|Q6PHN9 | RAB35_MO | 23 kDa | 0.999093 | 4 | 75% | Q6PHN9 | Rab35 | Ras-related | 22.62656 | 21.8727 | 22.81318 | 22.32889 | 22.57005 | 22.02688 | 0.593767 | -0.05576 | 1.243294 | 22.37304 | 2.175805 | -4.87736 | 0.067286 | 0.141457 | 0 | 0 |
| 690 | tr\|Q8C6B0 | Q8C6B0_M | 28 kDa | 0.998998 | 4 | 100% | Q8C6B0 | Mettl7a1 | MCG20149 | 22.2672 | 23.67224 | 22.61654 | 22.55147 | 22.44558 | 23.56109 | -0.81849 | -1.71562 | 0.078633 | 22.85236 | -2.17498 | -4.88006 | 0.067668 | 0.141622 | 0 | 0 |
| 903.2 | sp\|Q64523 | H2A2C_MO | 14 kDa | 0.998649 | 3 | 33% | Q64523 | Hist2h2ac | Histone H2 | 18.29777 | 20.58504 | 16.49685 | 18.40752 | 18.35316 | 18.94464 | -1.59647 | -3.34661 | 0.153667 | 18.51416 | -2.17691 | -4.87827 | 0.067678 | 0.141622 | 0 | 0 |
| 904 | sp\|P61021 | RAB5B_MO | 24 kDa | 0.998636 | 3 | 33% | P61021 | Rab5b | Ras-related | 18.29777 | 20.58504 | 16.49685 | 18.40752 | 18.35316 | 18.94464 | -1.59647 | -3.34661 | 0.153667 | 18.51416 | -2.17691 | -4.87827 | 0.067678 | 0.141622 | 0 | 0 |
| 2 | sp\|Q61879 | MYH10_M | 229 kDa | 0.999667 | 65 | 80% | Q61879 | Myh10 | Myosin-10 | 25.34386 | 25.94549 | 25.2567 | 25.98808 | 25.65435 | 26.03852 | -0.5724 | -1.20085 | 0.056055 | 25.7045 | -2.16783 | -4.88889 | 0.068072 | 0.142227 | 0 | 0 |
| 413 | sp\|Q8C9B9 | DIDO1_MO | 247 kDa | 0.999571 | 6 | 100% | Q8C9B9 | Dido1 | Death-indu | 19.48619 | 19.17092 | 19.42985 | 28.65219 | 19.07651 | 24.02508 | -4.61855 | -9.70578 | 0.468683 | 21.64012 | -2.16658 | -4.89314 | 0.068698 | 0.142911 | 0 | 0 |
| 623.1 | sp\|P51410 | RL9_MOUS | 22 kDa | 0.999492 | 4 | 100% | P51410 | Rpl9 | 60S ribosor | 22.37216 | 21.71602 | 22.55843 | 21.55681 | 22.40655 | 22.18749 | 0.625609 | -0.06332 | 1.314537 | 22.13291 | 2.161373 | -4.89822 | 0.068715 | 0.142911 | 0 | 0 |
| 623.2 | tr\|A0A140 | A0A140T8 | 22 kDa | 0.999492 | 4 | 100% | A0A140T8 | Rpl9-ps6 | Ribosomal | 22.37216 | 21.71602 | 22.55843 | 21.55681 | 22.40655 | 22.18749 | 0.625609 | -0.06332 | 1.314537 | 22.13291 | 2.161373 | -4.89822 | 0.068715 | 0.142911 | 0 | 0 |
| 229 | sp\|Q9CSN1 | SNW1_MO | 61 kDa | 0.999739 | 9 | 100% | Q9CSN1 | Snw1 | SNW doma | 22.63537 | 23.28721 | 21.94737 | 24.45001 | 23.21464 | 28.76926 | -2.90303 | -6.10542 | 0.29935 | 24.05064 | -2.16336 | -4.89777 | 0.069019 | 0.143092 | 0 | 0 |
| 847 | sp\|P21550 | ENOB_MOU | 47 kDa | 0.999183 | 3 | 33% | P21550 | Eno3 | Beta-enolas | 21.59499 | 19.76458 | 21.30936 | 20.68132 | 21.82217 | 21.3675 | 0.971038 | -0.10022 | 2.042298 | 21.08999 | 2.163179 | -4.89804 | 0.069037 | 0.143092 | 0 | 0 |
| 972 | tr\|V9GX06 | V9GX06_M | 20 kDa | 0.997556 | 3 | 33% | V9GX06 | Gm11214 | Predicted g | 18.50721 | 19.78297 | 18.03757 | 18.59997 | 18.84494 | 19.82273 | -0.93865 | -1.97457 | 0.097268 | 18.93257 | -2.16236 | -4.89921 | 0.069119 | 0.143092 | 0 | 0 |
| 20 | sp\|Q99NB9 | SF3B1_MO | 146 kDa | 0.999583 | 29 | 100% | Q99NB9 | Sf3b1 | Splicing fac | 25.87324 | 24.79496 | 25.11067 | 24.78684 | 26.0706 | 25.20829 | 0.754806 | -0.07876 | 1.588375 | 25.30743 | 2.15534 | -4.90698 | 0.069331 | 0.143313 | 0 | 0 |

| | | | | | | | | | | | | | | | | | | | | | | | | | |
|---|---|---|---|---|---|---|---|---|---|---|---|---|---|---|---|---|---|---|---|---|---|---|---|---|---|
| 927 | sp\|Q99JI4 | PSMD6_MO | 46 kDa | 0.998291 | 3 | 100% | Q99JI4 | Psmd6 | 26S protea | 20.53188 | 22.94741 | 20.64792 | 24.1763 | 20.45174 | 20.35352 | -1.94856 | -4.10926 | 0.212132 | 21.51813 | -2.15215 | -4.91392 | 0.07015 | 0.144407 | 0 | 0 |
| 1188 | sp\|P53026 | RL10A_MO | 25 kDa | 0.998088 | 2 | 100% | P53026 | Rpl10a | 60S ribosor | 16.47305 | 15.93451 | 15.75128 | 18.88918 | 16.27828 | 19.73769 | -2.01959 | -4.25931 | 0.220138 | 17.17733 | -2.15188 | -4.91429 | 0.070177 | 0.144407 | 0 | 0 |
| 1206 | sp\|Q91YI0 | ARLY_MOU | 52 kDa | 0.997863 | 2 | 100% | Q91YI0 | Asl | Argininosu | 19.60739 | 19.38953 | 19.69633 | 18.52972 | 19.49555 | 18.84432 | 0.678569 | -0.07372 | 1.430856 | 19.26047 | 2.146896 | -4.91912 | 0.07018 | 0.144407 | 0 | 0 |
| 749 | tr\|A0A140 | A0A140T8F | 13 kDa | 0.997834 | 4 | 25% | A0A140T8F | Igkv6-14 | Immunogl | 21.24037 | 23.49166 | 24.40437 | 26.1255 | 22.75383 | 24.90559 | -2.0414 | -4.31043 | 0.227639 | 23.82022 | -2.14702 | -4.92128 | 0.070673 | 0.144981 | 0 | 0 |
| 750.1 | sp\|P01635 | KV5A3_MO | 13 kDa | 0.997781 | 4 | 50% | P01635 | NA | Ig kappa ch | 21.24037 | 23.49166 | 24.40437 | 26.1255 | 22.75383 | 24.90559 | -2.0414 | -4.31043 | 0.227639 | 23.82022 | -2.14702 | -4.92128 | 0.070673 | 0.144981 | 0 | 0 |
| 1273 | sp\|Q99L04 | DHRS1_MO | 34 kDa | 0.996 | 2 | 100% | Q99L04 | Dhrs1 | Dehydroge | 18.49905 | 16.14366 | 17.89996 | 17.81497 | 17.73644 | 17.08379 | 1.031013 | -0.11706 | 2.179087 | 17.52964 | 2.143111 | -4.92691 | 0.071075 | 0.145585 | 0 | 0 |
| 1139 | sp\|Q8VCW | ACSF2_MO | 68 kDa | 0.998538 | 2 | 100% | Q8VCW8 | Acsf2 | Acyl-CoA sy | 18.50746 | 18.33504 | 19.98435 | 17.84882 | 18.7316 | 18.18022 | 0.953114 | -0.10877 | 2.015002 | 18.59791 | 2.141986 | -4.92852 | 0.071191 | 0.145602 | 0 | 0 |
| 1176 | sp\|Q61335 | BAP31_MO | 28 kDa | 0.998178 | 2 | 100% | Q61335 | Bcap31 | B-cell recep | 20.72006 | 21.74586 | 21.5107 | 21.60528 | 21.16145 | 23.15 | -1.03631 | -2.19588 | 0.123269 | 21.64889 | -2.13275 | -4.94179 | 0.07215 | 0.147121 | 0 | 0 |
| 1177.1 | sp\|P01750 | HVM06_MO | 13 kDa | 0.998168 | 2 | 50% | P01750 | NA | Ig heavy ch | 20.72006 | 21.74586 | 21.5107 | 21.60528 | 21.16145 | 23.15 | -1.03631 | -2.19588 | 0.123269 | 21.64889 | -2.13275 | -4.94179 | 0.07215 | 0.147121 | 0 | 0 |
| 316 | sp\|P53994 | RAB2A_MO | 24 kDa | 0.999651 | 7 | 29% | P53994 | Rab2a | Ras-related | 22.90826 | 22.36791 | 22.9107 | 22.24305 | 22.60549 | 22.13464 | 0.559615 | -0.06757 | 1.186804 | 22.52834 | 2.123693 | -4.95256 | 0.072594 | 0.147803 | 0 | 0 |
| 791 | tr\|A0A140 | A0A140T8F | 13 kDa | 0.999633 | 3 | 67% | A0A140T8F | Igkv12-46 | Immunogl | 22.37028 | 24.61678 | 25.16711 | 26.85868 | 22.747 | 24.67052 | -1.95387 | -4.14615 | 0.238421 | 24.40506 | -2.1269 | -4.95018 | 0.072764 | 0.147928 | 0 | 0 |
| 38 | sp\|Q8BSQ9 | PB1_MOUS | 187 kDa | 0.999426 | 23 | 100% | Q8BSQ9 | Pbrm1 | Protein pol | 24.00882 | 24.38111 | 22.83492 | 24.53214 | 24.1884 | 24.79239 | -0.89117 | -1.89175 | 0.109415 | 24.12296 | -2.12548 | -4.95223 | 0.072914 | 0.148012 | 0 | 0 |
| 589 | sp\|P57776 | EF1D_MOU | 31 kDa | 0.99988 | 4 | 100% | P57776 | Eef1d | Elongation | 20.68779 | 21.34694 | 21.48152 | 21.72065 | 20.77347 | 22.20864 | -0.77782 | -1.65251 | 0.096874 | 21.36983 | -2.11991 | -4.95935 | 0.073303 | 0.148527 | 0 | 0 |
| 177 | sp\|Q3TLH4 | PRC2C_MO | 311 kDa | 0.999166 | 11 | 100% | Q3TLH4 | Prrc2c | Protein PRI | 22.51569 | 21.66818 | 21.91524 | 21.97381 | 23.06239 | 21.50143 | 0.783296 | -0.09788 | 1.664476 | 22.10612 | 2.119119 | -4.96048 | 0.073387 | 0.148527 | 0 | 0 |
| 995 | sp\|A0A0B4 | FCGR4_MO | 28 kDa | 0.996145 | 3 | 100% | A0A0B4J1C | Fcgr4 | Low affinity | 19.48471 | 19.5456 | 19.12275 | 21.14874 | 19.69717 | 20.48786 | -0.95919 | -2.04133 | 0.12295 | 19.91447 | -2.1153 | -4.96683 | 0.073999 | 0.149541 | 0 | 0 |
| 1060.1 | sp\|P84104 | SRSF3_MO | 19 kDa | 0.999248 | 2 | 100% | P84104 | Srsf3 | Serine/argi | 18.04716 | 17.88871 | 18.42212 | 16.55398 | 17.99105 | 17.46613 | 0.850503 | -0.11351 | 1.814518 | 17.72819 | 2.105441 | -4.98096 | 0.075064 | 0.151377 | 0 | 0 |
| 669 | sp\|Q61466 | SMRD1_MO | 58 kDa | 0.999133 | 4 | 100% | Q61466 | Smarcd1 | SWI/SNF-re | 20.89988 | 19.41932 | 19.69688 | 19.84245 | 20.92136 | 19.68011 | 0.858749 | -0.11489 | 1.832393 | 20.07667 | 2.10483 | -4.98184 | 0.07513 | 0.151377 | 0 | 0 |
| 726 | sp\|P62852 | RS25_MOU | 14 kDa | 0.998563 | 4 | 100% | P62852 | Rps25 | 40S ribosor | 23.24155 | 23.15851 | 23.94058 | 22.4012 | 23.2139 | 22.62173 | 0.738199 | -0.09965 | 1.576043 | 23.09625 | 2.097779 | -4.99011 | 0.075454 | 0.151805 | 0 | 0 |
| 1080 | sp\|P63321 | RALA_MOU | 24 kDa | 0.999136 | 2 | 100% | P63321 | Rala | Ras-related | 17.08238 | 18.81028 | 19.13867 | 19.00508 | 17.31019 | 19.30791 | -1.19734 | -2.55797 | 0.163281 | 18.44242 | -2.10006 | -4.98867 | 0.075652 | 0.151976 | 0 | 0 |
| 1196 | sp\|Q91YW | DNJC3_MO | 57 kDa | 0.998015 | 2 | 100% | Q91YW3 | Dnajc3 | DnaJ homo | 19.86516 | 19.15825 | 19.3162 | 16.8684 | 19.68905 | 18.91287 | 1.3103 | -0.17984 | 2.800441 | 18.96832 | 2.098428 | -4.99101 | 0.075831 | 0.152111 | 0 | 0 |
| 1115 | sp\|Q9Z2Z6 | MCAT_MOU | 33 kDa | 0.998798 | 2 | 100% | Q9Z2Z6 | Slc25a20 | Mitochond | 18.16067 | 19.06355 | 18.15772 | 18.1919 | 17.55955 | 18.83327 | -0.73693 | -1.57661 | 0.102748 | 18.32778 | -2.09008 | -5.00134 | 0.07635 | 0.152926 | 0 | 0 |
| 832.3 | sp\|P01667 | KV3AF_MO | 12 kDa | 0.999312 | 3 | 33% | P01667 | NA | Ig kappa ch | 21.28451 | 22.07846 | 21.4009 | 22.0484 | 21.74552 | 21.97898 | -0.5583 | -1.19923 | 0.082625 | 21.75613 | -2.0733 | -5.02497 | 0.078126 | 0.156253 | 0 | 0 |
| 832.4 | sp\|P01669 | KV3AH_MO | 12 kDa | 0.999312 | 3 | 33% | P01669 | NA | Ig kappa ch | 21.11795 | 21.2825 | 20.81303 | 21.2857 | 20.13261 | 21.63708 | -0.7139 | -1.53462 | 0.106829 | 21.04481 | -2.07042 | -5.02912 | 0.078464 | 0.156697 | 0 | 0 |
| 876 | sp\|Q8BH43 | WASF2_MO | 54 kDa | 0.998953 | 3 | 100% | Q8BH43 | Wasf2 | Wiskott-Al | 19.41742 | 20.28957 | 19.92755 | 20.26697 | 19.30488 | 19.94935 | -0.61868 | -1.33059 | 0.093226 | 19.85929 | -2.06845 | -5.03191 | 0.078681 | 0.156898 | 0 | 0 |
| 604 | sp\|P35979 | RL12_MOU | 18 kDa | 0.99968 | 4 | 100% | P35979 | Rpl12 | 60S ribosor | 21.17717 | 20.86809 | 22.20451 | 20.85737 | 21.13721 | 20.43754 | 0.785296 | -0.12088 | 1.691476 | 21.11365 | 2.065917 | -5.03671 | 0.079285 | 0.157871 | 0 | 0 |
| 747 | sp\|P54775 | PRS6B_MO | 47 kDa | 0.997862 | 4 | 100% | P54775 | Psmc4 | 26S protea | 17.61825 | 19.01362 | 18.03245 | 18.91802 | 18.36808 | 18.24818 | -0.72035 | -1.55389 | 0.113188 | 18.36643 | -2.05703 | -5.0483 | 0.080011 | 0.159083 | 0 | 0 |
| 848 | sp\|Q8BI72 | CARF_MOU | 60 kDa | 0.999175 | 3 | 100% | Q8BI72 | Cdkn2aip | CDKN2A-in | 18.8391 | 22.32455 | 19.24099 | 18.67581 | 19.26623 | 23.77319 | -2.47574 | -5.34395 | 0.392464 | 20.35331 | -2.0599 | -5.04607 | 0.080188 | 0.159202 | 0 | 0 |
| 686 | sp\|Q9D1H7 | GET4_MOU | 37 kDa | 0.999009 | 4 | 100% | Q9D1H7 | Get4 | Golgi to ER | 20.27562 | 21.25011 | 20.78394 | 21.18487 | 20.79132 | 21.13495 | -0.57302 | -1.24005 | 0.094015 | 20.90347 | -2.04466 | -5.06594 | 0.081456 | 0.161483 | 0 | 0 |
| 429 | sp\|Q91YT0 | NDUV1_MO | 51 kDa | 0.999344 | 6 | 100% | Q91YT0 | Ndufv1 | NADH dehy | 22.94854 | 21.28818 | 21.62113 | 22.16856 | 22.89369 | 20.90478 | 1.033948 | -0.17229 | 2.240188 | 21.97081 | 2.045575 | -5.06649 | 0.081871 | 0.162069 | 0 | 0 |
| 1105 | sp\|Q64471 | GSTT1_MO | 27 kDa | 0.998915 | 2 | 100% | Q64471 | Gstt1 | Glutathion | 20.76699 | 20.15229 | 21.15158 | 20.10874 | 20.58863 | 20.46888 | 0.59243 | -0.09974 | 1.284605 | 20.53952 | 2.037147 | -5.07667 | 0.082353 | 0.162785 | 0 | 0 |
| 1007 | sp\|P63168 | DYL1_MOU | 10 kDa | 0.995001 | 3 | 33% | P63168 | Dynll1 | Dynein ligh | 20.14808 | 20.3688 | 20.73643 | 22.17295 | 20.19844 | 21.59913 | -1.01931 | -2.21235 | 0.173729 | 20.87064 | -2.03893 | -5.07595 | 0.082664 | 0.163162 | 0 | 0 |
| 670 | sp\|P19783 | COX41_MO | 20 kDa | 0.999124 | 4 | 100% | P19783 | Cox4i1 | Cytochrom | 22.38329 | 24.04041 | 23.76296 | 23.92295 | 22.54927 | 23.41614 | -0.89466 | -1.94702 | 0.157702 | 23.34584 | -2.02882 | -5.09033 | 0.083884 | 0.16533 | 0 | 0 |
| 194.1 | sp\|P03987 | IGHG3_MO | 44 kDa | 0.999623 | 10 | 100% | P03987 | NA | Ig gamma-3 | 31.06637 | 31.41955 | 31.23238 | 31.84414 | 30.74192 | 31.51004 | -0.57769 | -1.26244 | 0.107071 | 31.3024 | -2.00797 | -5.11826 | 0.08593 | 0.168627 | 0 | 0 |
| 194.2 | tr\|A0A1Y7 | A0A1Y7VJN | 44 kDa | 0.999623 | 10 | 100% | A0A1Y7VJN | Ighg3 | Immunogl | 31.06637 | 31.41955 | 31.23238 | 31.84414 | 30.74192 | 31.51004 | -0.57769 | -1.26244 | 0.107071 | 31.3024 | -2.00797 | -5.11826 | 0.08593 | 0.168627 | 0 | 0 |
| 194.3 | tr\|A0A075 | A0A075B5I | 36 kDa | 0.999623 | 10 | 100% | A0A075B5I | Ighg3 | Immunogl | 31.06637 | 31.41955 | 31.23238 | 31.84414 | 30.74192 | 31.51004 | -0.57769 | -1.26244 | 0.107071 | 31.3024 | -2.00797 | -5.11826 | 0.08593 | 0.168627 | 0 | 0 |
| 73 | sp\|Q9QWL | K1C17_MO | 48 kDa | 0.999655 | 17 | 41% | Q9QWL7 | Krt17 | Keratin, typ | 21.78826 | 23.94931 | 24.59178 | 25.21657 | 23.7104 | 25.73277 | -1.60274 | -3.50653 | 0.301047 | 24.16485 | -2.00907 | -5.11837 | 0.086321 | 0.169149 | 0 | 0 |
| 1203 | sp\|Q91Z49 | UIF_MOUSI | 36 kDa | 0.997924 | 2 | 100% | Q91Z49 | Fyttd1 | UAP56-inte | 17.54395 | 20.67548 | 21.10342 | 22.47061 | 19.29936 | 20.56734 | -1.92223 | -4.20837 | 0.363908 | 20.27669 | -2.00657 | -5.12192 | 0.086635 | 0.169519 | 0 | 0 |
| 1266 | sp\|Q80WM | HPLN4_MO | 43 kDa | 0.996345 | 2 | 100% | Q80WM4 | Hapln4 | Hyaluronar | 17.31378 | 17.18517 | 16.70941 | 18.95847 | 16.57449 | 17.54914 | -1.0317 | -2.26042 | 0.197016 | 17.38174 | -2.00379 | -5.12586 | 0.086984 | 0.169755 | 0 | 0 |
| 11 | sp\|Q3UHU | MTCL1_MO | 214 kDa | 0.99993 | 39 | 92% | Q3UHU5 | Mtcl1 | Microtubu | 25.32065 | 23.95463 | 24.22275 | 24.3119 | 25.82497 | 24.38785 | 0.904665 | -0.17285 | 1.982179 | 24.67046 | 2.003621 | -5.1261 | 0.087006 | 0.169755 | 0 | 0 |
| 989 | sp\|Q6NZA9 | TAF9B_MO | 27 kDa | 0.996854 | 3 | 100% | Q6NZA9 | Taf9b | Transcripti | 19.22655 | 21.38171 | 18.28191 | 19.95401 | 19.96765 | 19.74334 | -1.20098 | -2.63262 | 0.230657 | 19.7592 | -2.00195 | -5.12847 | 0.087217 | 0.169922 | 0 | 0 |
| 263 | sp\|P18872 | GNAO_MOU | 40 kDa | 0.999709 | 8 | 75% | P18872 | Gnao1 | Guanine nu | 23.37746 | 22.92768 | 23.83198 | 23.16522 | 23.26878 | 22.4688 | 0.638839 | -0.12464 | 1.402315 | 23.17332 | 1.991576 | -5.14155 | 0.088006 | 0.171215 | 0 | 0 |
| 973 | sp\|P61211 | ARL1_MOU | 20 kDa | 0.997547 | 3 | 100% | P61211 | Arl1 | ADP-ribosy | 21.06796 | 20.32074 | 21.05429 | 20.73157 | 20.71281 | 19.47779 | 0.768323 | -0.15257 | 1.689217 | 20.56086 | 1.990481 | -5.14453 | 0.08862 | 0.172162 | 0 | 0 |
| 872 | sp\|Q8VDM | HNRL1_MO | 96 kDa | 0.998975 | 3 | 100% | Q8VDM6 | Hnrnpul1 | Heterogene | 17.87002 | 20.53286 | 17.82245 | 18.14826 | 18.4014 | 19.03686 | -1.20803 | -2.65826 | 0.242192 | 18.63531 | -1.9879 | -5.14837 | 0.089011 | 0.172674 | 0 | 0 |
| 809 | sp\|P21460 | CYTC_MOU | 16 kDa | 0.999461 | 3 | 100% | P21460 | Cst3 | Cystatin-C | 20.25379 | 20.61908 | 20.32381 | 21.16063 | 19.97906 | 20.55024 | -0.5911 | -1.30197 | 0.11978 | 20.4811 | -1.97909 | -5.15927 | 0.089622 | 0.173514 | 0 | 0 |
| 1125 | sp\|Q8VBT0 | TMX1_MOU | 31 kDa | 0.998676 | 2 | 100% | Q8VBT0 | Tmx1 | Thioredoxi | 20.1296 | 20.37123 | 21.13759 | 18.59105 | 21.40044 | 20.22167 | 1.161229 | -0.23704 | 2.559497 | 20.30859 | 1.981884 | -5.15687 | 0.089791 | 0.173514 | 0 | 0 |
| 692 | sp\|P01671 | KV3AJ_MO | 12 kDa | 0.998994 | 4 | 50% | P01671 | NA | Ig kappa ch | 21.86401 | 22.76124 | 21.36318 | 27.61294 | 22.02664 | 22.61566 | -2.57867 | -5.68444 | 0.527106 | 23.04061 | -1.98142 | -5.15753 | 0.089851 | 0.173514 | 0 | 0 |
| 1224 | sp\|Q9JIK9 | RT34_MOU | 26 kDa | 0.997577 | 2 | 100% | Q9JIK9 | Mrps34 | 28S ribosor | 17.41549 | 17.08561 | 17.37655 | 16.21332 | 17.34728 | 16.93214 | 0.636085 | -0.12989 | 1.402057 | 17.06173 | 1.976531 | -5.16289 | 0.089956 | 0.173514 | 0 | 0 |
| 344 | sp\|Q05920 | PYC_MOUS | 130 kDa | 0.999363 | 7 | 100% | Q05920 | Pc | Pyruvate ca | 21.57737 | 22.62811 | 21.58681 | 22.18397 | 22.1268 | 22.26508 | -0.5954 | -1.31398 | 0.123184 | 22.06136 | -1.97211 | -5.16915 | 0.090537 | 0.174169 | 0 | 0 |
| 1016 | sp\|Q9CRB2 | NHP2_MOU | 17 kDa | 0.999784 | 2 | 100% | Q9CRB2 | Nhp2 | H/ACA ribo | 20.50024 | 20.11229 | 18.97951 | 21.27965 | 19.62813 | 20.64152 | -0.97519 | -2.15292 | 0.202529 | 20.19022 | -1.97605 | -5.16511 | 0.090553 | 0.174169 | 0 | 0 |
| 165 | sp\|Q99LF4 | RTCB_MOU | 55 kDa | 0.999492 | 11 | 100% | Q99LF4 | Rtcb | tRNA-splici | 23.6508 | 24.02761 | 23.06105 | 23.62805 | 23.19807 | 24.02312 | -0.58962 | -1.30277 | 0.12353 | 23.59812 | -1.96785 | -5.17519 | 0.0911 | 0.174975 | 0 | 0 |
| 351 | sp\|Q9EQP2 | EHD4_MOU | 61 kDa | 0.999293 | 7 | 100% | Q9EQP2 | Ehd4 | EH domain- | 21.90479 | 23.40897 | 22.56605 | 22.56219 | 21.64188 | 22.47898 | -0.77914 | -1.72611 | 0.167832 | 22.42714 | -1.96349 | -5.18284 | 0.092216 | 0.176867 | 0 | 0 |
| 515 | sp\|P35550 | FBRL_MOU | 34 kDa | 0.999391 | 5 | 100% | P35550 | Fbl | rRNA 2'-O-r | 21.53485 | 20.90654 | 20.80488 | 20.82081 | 21.67181 | 19.73344 | 0.850251 | -0.18442 | 1.884925 | 20.91206 | 1.961076 | -5.18625 | 0.092539 | 0.176918 | 0 | 0 |
| 26 | tr\|E9Q6R7 | E9Q6R7_M | 393 kDa | 0.999404 | 26 | 100% | E9Q6R7 | Utrn | Utrophin | 23.74854 | 24.11449 | 22.95712 | 24.01407 | 23.29688 | 23.72689 | -0.61764 | -1.36898 | 0.133705 | 23.643 | -1.95657 | -5.19114 | 0.092608 | 0.176918 | 0 | 0 |
| 1254 | sp\|Q61838 | PZP_MOUS | 166 kDa | 0.996716 | 2 | 100% | Q61838 | Pzp | Pregnancy | 18.06137 | 17.17247 | 17.91895 | 17.92188 | 18.16303 | 15.57041 | 1.159525 | -0.25201 | 2.571064 | 17.46802 | 1.960368 | -5.18725 | 0.092634 | 0.176918 | 0 | 0 |
| 814 | sp\|P62889 | RL30_MOU | 13 kDa | 0.999429 | 3 | 100% | P62889 | Rpl30 | 60S ribosor | 18.76563 | 20.81154 | 22.01423 | 23.84083 | 20.20538 | 22.31093 | -1.99269 | -4.43086 | 0.445487 | 21.32476 | -1.95041 | -5.20128 | 0.093981 | 0.179237 | 0 | 0 |
| 272 | sp\|Q9CY64 | BIEA_MOUS | 34 kDa | 0.999545 | 8 | 100% | Q9CY64 | Blvra | Biliverdin r | 22.22623 | 21.43275 | 22.26378 | 21.66638 | 21.86617 | 21.6715 | 0.52852 | -0.11822 | 1.175259 | 21.85447 | 1.94506 | -5.2074 | 0.094173 | 0.179353 | 0 | 0 |
| 582 | sp\|Q80XI3 | IF4G3_MO | 175 kDa | 0.997392 | 5 | 60% | Q80XI3 | Eif4g3 | Eukaryotic | 18.57404 | 17.95481 | 18.62386 | 17.98449 | 18.65492 | 18.36391 | 0.516538 | -0.11883 | 1.151901 | 18.35934 | 1.934999 | -5.22158 | 0.095562 | 0.181742 | 0 | 0 |
| 129 | sp\|Q8C2Q3 | RBM14_MO | 69 kDa | 0.999661 | 13 | 100% | Q8C2Q3 | Rbm14 | RNA-bindir | 23.65097 | 23.01881 | 24.15315 | 23.52274 | 23.68625 | 23.22118 | 0.575876 | -0.13336 | 1.285108 | 23.54218 | 1.932598 | -5.22497 | 0.095897 | 0.182123 | 0 | 0 |
| 355 | sp\|Q00612 | G6PD1_MO | 59 kDa | 0.99922 | 7 | 100% | Q00612 | G6pdx | Glucose-6- | 21.28601 | 21.79778 | 21.25247 | 22.04192 | 21.36447 | 21.60195 | -0.5129 | -1.14668 | 0.120881 | 21.55743 | -1.92617 | -5.23402 | 0.096798 | 0.183578 | 0 | 0 |
| 733 | sp\|P80314 | TCPB_MOU | 57 kDa | 0.998386 | 4 | 100% | P80314 | Cct2 | T-complex | 21.83593 | 22.83208 | 19.75246 | 21.64363 | 21.73009 | 22.54547 | -1.23423 | -2.76294 | 0.294472 | 21.72328 | -1.92674 | -5.23454 | 0.097257 | 0.18419 | 0 | 0 |
| 368 | sp\|Q9D554 | SF3A3_MO | 59 kDa | 0.999004 | 7 | 100% | Q9D554 | Sf3a3 | Splicing fac | 23.46252 | 22.1916 | 22.37233 | 22.83474 | 23.45323 | 21.57347 | 0.896091 | -0.21557 | 2.007748 | 22.64798 | 1.923677 | -5.23884 | 0.09769 | 0.184635 | 0 | 0 |
| 712 | sp\|P59708 | SF3B6_MO | 15 kDa | 0.998754 | 4 | 100% | P59708 | Sf3b6 | Splicing fac | 23.31885 | 24.30827 | 22.60936 | 23.4416 | 23.45743 | 23.78346 | -0.7159 | -1.60402 | 0.172225 | 23.48649 | -1.92164 | -5.24118 | 0.097764 | 0.184635 | 0 | 0 |
| 633 | sp\|Q8R1Q8 | DC1L1_MO | 57 kDa | 0.999422 | 4 | 100% | Q8R1Q8 | Dync1li1 | Cytoplasmi | 20.62262 | 20.38908 | 21.32239 | 19.68148 | 21.79843 | 20.99653 | 0.89212 | -0.21566 | 1.999895 | 20.80175 | 1.921863 | -5.24139 | 0.097947 | 0.184723 | 0 | 0 |
| 1239 | sp\|Q920R6 | VPP4_MOU | 96 kDa | 0.997235 | 2 | 50% | Q920R6 | Atp6v0a4 | V-type prot | 19.78293 | 20.66062 | 19.81136 | 20.35308 | 20.27215 | 23.29947 | -1.48224 | -3.3259 | 0.361409 | 20.6966 | -1.91863 | -5.24592 | 0.098406 | 0.185331 | 0 | 0 |
| 337 | sp\|P04919 | B3AT_MOU | 103 kDa | 0.999457 | 7 | 100% | P04919 | Slc4a1 | Band 3 anic | 22.61503 | 21.81561 | 22.10359 | 20.26714 | 21.51081 | 21.30427 | 0.947472 | -0.23211 | 2.127058 | 21.60274 | 1.916848 | -5.24842 | 0.09866 | 0.185552 | 0 | 0 |
| 1048 | sp\|O08529 | CAN2_MOU | 80 kDa | 0.999364 | 2 | 100% | O08529 | Capn2 | Calpain-2 c | 19.61498 | 18.73694 | 21.28691 | 19.80762 | 20.13886 | 19.53021 | 0.988664 | -0.24366 | 2.220993 | 19.85259 | 1.914578 | -5.2516 | 0.098985 | 0.185905 | 0 | 0 |
| 192 | sp\|Q61584 | FXR1_MOU | 76 kDa | 0.999635 | 10 | 90% | Q61584 | Fxr1 | Fragile X m | 22.1959 | 20.3458 | 21.5801 | 20.38229 | 21.96489 | 22.04761 | 0.988393 | -0.24478 | 2.221564 | 21.41943 | 1.912746 | -5.25416 | 0.099248 | 0.186141 | 0 | 0 |
| 244 | sp\|P67778 | PHB_MOUS | 30 kDa | 0.999094 | 9 | 100% | P67778 | Phb | Prohibitin | 22.88709 | 23.70663 | 23.35839 | 23.59607 | 22.54393 | 23.30905 | -0.60745 | -1.36706 | 0.152164 | 23.23353 | -1.90335 | -5.26608 | 0.100064 | 0.187413 | 0 | 0 |
| 920 | sp\|P70452 | STX4_MOU | 34 kDa | 0.998396 | 3 | 100% | P70452 | Stx4 | Syntaxin-4 | 21.55224 | 21.87244 | 22.14579 | 22.66145 | 21.24589 | 22.50962 | -0.69986 | -1.57974 | 0.180017 | 21.9979 | -1.8962 | -5.27683 | 0.101437 | 0.189723 | 0 | 0 |
| 1055 | sp\|A2AI08 | TPRN_MOU | 80 kDa | 0.999289 | 2 | 100% | A2AI08 | Tprn | Taperin | 17.34622 | 19.36158 | 15.93158 | 17.5636 | 15.94543 | 16.6191 | -1.44035 | -3.25393 | 0.373222 | 17.12792 | -1.89533 | -5.27853 | 0.101781 | 0.189842 | 0 | 0 |
| 1056.1 | sp\|P02463 | CO4A1_MO | 161 kDa | 0.999285 | 2 | 100% | P02463 | Col4a1 | Collagen al | 17.34622 | 19.36158 | 15.93158 | 17.5636 | 15.94543 | 16.6191 | -1.44035 | -3.25393 | 0.373222 | 17.12792 | -1.89533 | -5.27853 | 0.101781 | 0.189842 | 0 | 0 |
| 1175 | sp\|Q8VCM | NCLN_MOU | 63 kDa | 0.998178 | 2 | 100% | Q8VCM8 | Ncln | Nicalin | 22.78731 | 23.43039 | 23.23011 | 23.26334 | 22.39577 | 23.48805 | -0.58953 | -1.33325 | 0.154194 | 23.09916 | -1.88667 | -5.28944 | 0.10252 | 0.190797 | 0 | 0 |
| 870 | sp\|P97461 | RS5_MOUS | 23 kDa | 0.998991 | 3 | 100% | P97461 | Rps5 | 40S ribosor | 20.62235 | 21.00697 | 21.06139 | 21.68757 | 20.61873 | 21.3633 | -0.58513 | -1.32344 | 0.153187 | 21.06005 | -1.8863 | -5.28996 | 0.102574 | 0.190797 | 0 | 0 |
| 1049 | sp\|P62855 | RS26_MOU | 13 kDa | 0.999332 | 2 | 100% | P62855 | Rps26 | 40S ribosor | 18.8746 | 20.18957 | 19.34226 | 20.1773 | 20.08921 | 20.0271 | -0.69597 | -1.57515 | 0.183218 | 19.78334 | -1.88713 | -5.2895 | 0.102778 | 0.190915 | 0 | 0 |
| 798.1 | sp\|Q9WV0 | RBMX_MO | 42 kDa | 0.999544 | 3 | 100% | Q9WV02 | Rbmx | RNA-bindir | 22.08296 | 20.84523 | 20.38172 | 19.78061 | 21.85413 | 20.63443 | 1.019515 | -0.2713 | 2.310336 | 20.92984 | 1.884858 | -5.29314 | 0.103334 | 0.191684 | 0 | 0 |
| 237 | sp\|Q61666 | HIRA_MOU | 112 kDa | 0.999458 | 9 | 100% | Q61666 | Hira | Protein HIR | 22.95569 | 21.48586 | 21.55902 | 21.54089 | 24.35422 | 22.00045 | 1.280577 | -0.34562 | 2.906779 | 22.31602 | 1.879239 | -5.30097 | 0.104177 | 0.192984 | 0 | 0 |
| 819 | sp\|Q5FWK | RHG01_MO | 50 kDa | 0.999402 | 3 | 100% | Q5FWK3 | Arhgap1 | Rho GTPase | 22.7233 | 22.13758 | 22.16103 | 20.45174 | 21.2809 | 19.67692 | 1.299665 | -0.35361 | 2.952937 | 21.40525 | 1.876022 | -5.30545 | 0.104663 | 0.193579 | 0 | 0 |
| 1180 | sp\|Q8VHG | AMOT_MO | 121 kDa | 0.99815 | 2 | 100% | Q8VHG2 | Amot | Angiomotir | 19.98924 | 19.4095 | 20.58648 | 19.8315 | 19.79393 | 19.24965 | 0.626333 | -0.17017 | 1.422835 | 19.81005 | 1.871625 | -5.31045 | 0.104784 | 0.193579 | 0 | 0 |
| 1078 | sp\|Q9CR78 | MSD3_MO | 32 kDa | 0.999141 | 2 | 100% | Q9CR78 | Msantd3 | Myb/SANT- | 17.06162 | 19.77194 | 19.20468 | 18.48108 | 18.38776 | 20.1465 | -1.24849 | -2.83945 | 0.342482 | 18.84226 | -1.87272 | -5.31005 | 0.105163 | 0.194016 | 0 | 0 |
| 58 | sp\|O55143 | AT2A2_MO | 115 kDa | 0.999641 | 19 | 100% | O55143 | Atp2a2 | Sarcoplasm | 24.40454 | 25.50284 | 24.36559 | 25.76005 | 24.10731 | 24.17567 | -0.85371 | -1.94399 | 0.236566 | 24.71933 | -1.86864 | -5.31573 | 0.105786 | 0.1949 | 0 | 0 |
| 423 | sp\|E9PV24 | FIBA_MOUS | 87 kDa | 0.999433 | 6 | 100% | E9PV24 | Fga | Fibrinogen | 24.08215 | 25.11773 | 22.61217 | 24.17888 | 24.01419 | 24.22623 | -0.93811 | -2.13696 | 0.260739 | 24.03856 | -1.86741 | -5.31743 | 0.105974 | 0.19498 | 0 | 0 |
| 68 | sp\|P50446 | K2C6A_MO | 59 kDa | 0.999228 | 18 | 22% | P50446 | Krt6a | Keratin, typ | 19.02408 | 21.00874 | 21.5277 | 22.14506 | 21.24919 | 23.36931 | -1.57405 | -3.58756 | 0.439465 | 21.38735 | -1.86558 | -5.31998 | 0.106254 | 0.195232 | 0 | 0 |
| 829 | sp\|Q9R0U0 | SRS10_MO | 31 kDa | 0.999358 | 3 | 100% | Q9R0U0 | Srsf10 | Serine/argi | 22.44571 | 21.7926 | 22.13316 | 22.21904 | 22.70403 | 21.46182 | 0.603146 | -0.16847 | 1.374765 | 22.12606 | 1.860458 | -5.32602 | 0.106497 | 0.195412 | 0 | 0 |
| 743 | sp\|Q61187 | TS101_MO | 44 kDa | 0.997956 | 4 | 100% | Q61187 | Tsg101 | Tumor susc | 20.45914 | 22.6445 | 19.68307 | 20.46677 | 20.51144 | 20.91239 | -1.12334 | -2.56695 | 0.320278 | 20.77955 | -1.85699 | -5.33191 | 0.107582 | 0.197136 | 0 | 0 |
| 200 | sp\|Q9WTI7 | MYO1C_MO | 122 kDa | 0.999458 | 10 | 100% | Q9WTI7 | Myo1c | Unconvent | 22.21048 | 21.46957 | 21.66281 | 20.82646 | 21.94149 | 21.65874 | 0.620004 | -0.17962 | 1.41963 | 21.62826 | 1.845474 | -5.34686 | 0.108838 | 0.199169 | 0 | 0 |
| 584 | sp\|P17742 | PPIA_MOU | 18 kDa | 0.997028 | 5 | 100% | P17742 | Ppia | Peptidyl-pr | 23.37981 | 22.72229 | 22.4707 | 23.99901 | 22.3828 | 24.41963 | -0.9692 | -2.2225 | 0.284091 | 23.22904 | -1.84549 | -5.34786 | 0.109383 | 0.199897 | 0 | 0 |
| 1257 | sp\|Q8BH24 | TM9S4_MO | 75 kDa | 0.996566 | 2 | 100% | Q8BH24 | Tm9sf4 | Transmemb | 17.17198 | 14.69957 | 18.04486 | 16.30367 | 17.3162 | 17.56562 | 1.321395 | -0.38945 | 3.032236 | 16.85032 | 1.843205 | -5.35103 | 0.109745 | 0.200289 | 0 | 0 |
| 586 | sp\|Q9R1C7 | PR40A_MO | 108 kDa | 0.996922 | 5 | 100% | Q9R1C7 | Prpf40a | Pre-mRNA- | 20.39896 | 20.29174 | 20.55306 | 19.57126 | 20.89738 | 20.21221 | 0.591395 | -0.17685 | 1.359639 | 20.32077 | 1.832226 | -5.36525 | 0.110949 | 0.202009 | 0 | 0 |
| 403.1 | sp\|C0HKD8 | MFA1A_MO | 52 kDa | 0.999627 | 6 | 100% | C0HKD8 | Mfap1a | Microfibril | 22.26411 | 21.18953 | 21.34933 | 21.40389 | 22.88534 | 21.54029 | 0.788361 | -0.2372 | 1.813919 | 21.77208 | 1.83449 | -5.36309 | 0.111135 | 0.202009 | 0 | 0 |
| 403.2 | sp\|C0HKD9 | MFA1B_MO | 52 kDa | 0.999627 | 6 | 100% | C0HKD9 | Mfap1b | Microfibril | 22.26411 | 21.18953 | 21.34933 | 21.40389 | 22.88534 | 21.54029 | 0.788361 | -0.2372 | 1.813919 | 21.77208 | 1.83449 | -5.36309 | 0.111135 | 0.202009 | 0 | 0 |
| 332 | sp\|Q61990 | PCBP2_MO | 38 kDa | 0.999483 | 7 | 57% | Q61990 | Pcbp2 | Poly(rC)-bi | 23.73992 | 23.27985 | 23.70337 | 23.15774 | 23.41051 | 22.93915 | 0.492353 | -0.15059 | 1.135295 | 23.37176 | 1.822656 | -5.3785 | 0.112499 | 0.204215 | 0 | 0 |
| 756 | sp\|Q9JKY5 | HIP1R_MO | 119 kDa | 0.997556 | 4 | 100% | Q9JKY5 | Hip1r | Huntingtin | 22.81746 | 22.1869 | 22.6072 | 22.54227 | 22.95581 | 22.14697 | 0.501441 | -0.15502 | 1.157903 | 22.54277 | 1.818074 | -5.38483 | 0.113249 | 0.205088 | 0 | 0 |
| 565 | sp\|Q6ZWV | RL35_MOU | 15 kDa | 0.998603 | 5 | 100% | Q6ZWV7 | Rpl35 | 60S ribosor | 22.7477 | 23.54159 | 23.15752 | 23.56501 | 22.9289 | 23.22948 | -0.50066 | -1.15635 | 0.15504 | 23.19503 | -1.81735 | -5.38584 | 0.113368 | 0.205088 | 0 | 0 |
| 135 | sp\|P21619 | LMNB2_MO | 67 kDa | 0.99944 | 13 | 92% | P21619 | Lmnb2 | Lamin-B2 | 23.36068 | 24.06857 | 23.58733 | 23.98207 | 23.68842 | 23.99327 | -0.46916 | -1.08375 | 0.145422 | 23.78006 | -1.81695 | -5.38639 | 0.113434 | 0.205088 | 0 | 0 |
| 416 | sp\|Q02053 | UBA1_MOU | 118 kDa | 0.99956 | 6 | 100% | Q02053 | Uba1 | Ubiquitin-l | 21.0104 | 22.86488 | 22.57702 | 22.02513 | 21.44343 | 22.76394 | -0.87436 | -2.02318 | 0.274455 | 22.11413 | -1.81631 | -5.38818 | 0.114088 | 0.205997 | 0 | 0 |
| 1030.2 | tr\|A0A075 | A0A075B5I | 13 kDa | 0.999544 | 2 | 100% | A0A075B5I | Igkv17-121 | Immunogl | 21.3192 | 21.76438 | 21.22886 | 21.73695 | 21.26646 | 21.65953 | -0.44878 | -1.03994 | 0.142378 | 21.4959 | -1.80689 | -5.40028 | 0.115099 | 0.207546 | 0 | 0 |
| 866 | sp\|Q9CQ62 | DECR_MOU | 36 kDa | 0.999025 | 3 | 100% | Q9CQ62 | Decr1 | 2,4-dienoy | 20.71805 | 19.86758 | 21.12629 | 20.55935 | 20.62873 | 20.34668 | 0.566492 | -0.18323 | 1.316215 | 20.54112 | 1.798429 | -5.41194 | 0.116517 | 0.209823 | 0 | 0 |
| 1094 | sp\|Q52KI8 | SRRM1_MO | 107 kDa | 0.99901 | 2 | 100% | Q52KI8 | Srrm1 | Serine/argi | 17.7902 | 19.55372 | 19.21977 | 19.57931 | 19.4818 | 19.97399 | -0.87175 | -2.0303 | 0.286805 | 19.26646 | -1.79567 | -5.41659 | 0.117536 | 0.211148 | 0 | 0 |
| 358 | sp\|O54734 | OST48_MO | 49 kDa | 0.999154 | 7 | 100% | O54734 | Ddost | Dolichyl-di | 23.30466 | 23.23761 | 23.62532 | 22.90064 | 23.32284 | 22.39744 | 0.572374 | -0.18863 | 1.333376 | 23.13142 | 1.790175 | -5.4233 | 0.117917 | 0.211148 | 0 | 0 |
| 1225 | sp\|O35326 | SRSF5_MO | 31 kDa | 0.997565 | 2 | 50% | O35326 | Srsf5 | Serine/argi | 20.6504 | 20.17127 | 19.96271 | 19.11824 | 20.30169 | 19.62527 | 0.666673 | -0.22021 | 1.553556 | 19.9716 | 1.792008 | -5.42128 | 0.117938 | 0.211148 | 0 | 0 |
| 1226 | sp\|Q925I1 | ATAD3_MO | 67 kDa | 0.997556 | 2 | 100% | Q925I1 | Atad3 | ATPase fam | 20.6504 | 20.17127 | 19.96271 | 19.11824 | 20.30169 | 19.62527 | 0.666673 | -0.22021 | 1.553556 | 19.9716 | 1.792008 | -5.42128 | 0.117938 | 0.211148 | 0 | 0 |
| 467 | sp\|P63154 | CRNL1_MO | 83 kDa | 0.998287 | 6 | 100% | P63154 | Crnkl1 | Crooked ne | 21.55484 | 19.96839 | 20.45266 | 20.71832 | 21.73213 | 20.69854 | 0.784789 | -0.25989 | 1.829471 | 20.85415 | 1.792749 | -5.42059 | 0.118031 | 0.211148 | 0 | 0 |
| 968 | sp\|Q99MR | ACL6B_MO | 47 kDa | 0.997607 | 3 | 67% | Q99MR0 | Actl6b | Actin-like p | 20.41374 | 19.47328 | 20.52814 | 19.83619 | 19.86589 | 19.85308 | 0.548409 | -0.18283 | 1.279649 | 19.99505 | 1.785029 | -5.43037 | 0.118798 | 0.212241 | 0 | 0 |
| 250 | sp\|P11276 | FINC_MOU | 273 kDa | 0.998514 | 9 | 100% | P11276 | Fn1 | Fibronectir | 21.63618 | 22.45669 | 21.69182 | 22.90146 | 21.75908 | 21.68862 | -0.65323 | -1.53032 | 0.223865 | 22.02231 | -1.77547 | -5.44396 | 0.120786 | 0.215508 | 0 | 0 |
| 1174 | sp\|Q9D832 | DNJB4_MO | 38 kDa | 0.998188 | 2 | 100% | Q9D832 | Dnajb4 | DnaJ homo | 17.64287 | 17.58509 | 19.29275 | 18.16608 | 18.72357 | 17.32004 | 0.862661 | -0.29839 | 2.02371 | 18.12173 | 1.773129 | -5.44747 | 0.121414 | 0.216343 | 0 | 0 |
| 902 | sp\|Q8BRF7 | SCFD1_MO | 72 kDa | 0.998675 | 3 | 100% | Q8BRF7 | Scfd1 | Sec1 family | 20.57609 | 20.29953 | 21.49423 | 20.33285 | 20.81199 | 20.5077 | 0.580742 | -0.20093 | 1.362417 | 20.6704 | 1.76831 | -5.4533 | 0.121705 | 0.216386 | 0 | 0 |
| 634 | sp\|Q6DFW | NOP58_MO | 60 kDa | 0.999422 | 4 | 100% | Q6DFW4 | Nop58 | Nucleolar p | 19.71475 | 18.06997 | 19.89668 | 18.17703 | 20.23419 | 20.29432 | 1.10143 | -0.38262 | 2.585477 | 19.39782 | 1.771169 | -5.45015 | 0.121757 | 0.216386 | 0 | 0 |
| 893 | sp\|Q8BKT7 | THOC5_MO | 79 kDa | 0.998758 | 3 | 100% | Q8BKT7 | Thoc5 | THO compl | 18.8306 | 17.41741 | 17.07995 | 15.33539 | 19.42075 | 17.99951 | 1.526329 | -0.53378 | 3.586435 | 17.6806 | 1.76811 | -5.45433 | 0.122294 | 0.21699 | 0 | 0 |
| 937 | sp\|Q9QZD9 | EIF3I_MOU | 36 kDa | 0.998119 | 3 | 100% | Q9QZD9 | Eif3i | Eukaryotic | 18.84203 | 18.23063 | 19.08623 | 18.46689 | 18.65219 | 18.44887 | 0.478018 | -0.16686 | 1.122898 | 18.62114 | 1.764277 | -5.45882 | 0.122417 | 0.21699 | 0 | 0 |
| 506 | sp\|Q60931 | VDAC3_MO | 31 kDa | 0.999502 | 5 | 100% | Q60931 | Vdac3 | Voltage-de | 22.41538 | 21.37195 | 22.1468 | 21.35602 | 22.03295 | 22.13765 | 0.576504 | -0.20395 | 1.356958 | 21.91013 | 1.758149 | -5.4672 | 0.123505 | 0.218634 | 0 | 0 |
| 335 | sp\|Q9R0Q1 | SYTL4_MO | 76 kDa | 0.999466 | 7 | 100% | Q9R0Q1 | Sytl4 | Synaptotag | 21.71318 | 22.30569 | 21.82421 | 22.07671 | 21.96402 | 22.56567 | -0.48222 | -1.13654 | 0.1721 | 22.07491 | -1.7541 | -5.47272 | 0.124229 | 0.219628 | 0 | 0 |
| 930.2 | sp\|P63001 | RAC1_MOU | 21 kDa | 0.998224 | 3 | 100% | P63001 | Rac1 | Ras-related | 19.77637 | 20.1645 | 19.79768 | 20.15404 | 19.59646 | 20.1729 | -0.44031 | -1.0383 | 0.157675 | 19.94366 | -1.75255 | -5.47485 | 0.124509 | 0.219836 | 0 | 0 |
| 1281 | sp\|Q8C5N3 | CWC22_MO | 105 kDa | 0.995614 | 2 | 100% | Q8C5N3 | Cwc22 | Pre-mRNA- | 21.6343 | 22.10314 | 20.99153 | 21.64782 | 21.26518 | 21.73397 | -0.5313 | -1.25582 | 0.193215 | 21.56266 | -1.7454 | -5.4846 | 0.125801 | 0.221768 | 0 | 0 |
| 988 | sp\|Q9EQZ6 | RPGF4_MO | 115 kDa | 0.996858 | 3 | 100% | Q9EQZ6 | Rapgef4 | Rap guanin | 18.6788 | 18.85052 | 18.92413 | 21.09611 | 19.87571 | 21.20583 | -1.22461 | -2.89674 | 0.447525 | 19.77185 | -1.74774 | -5.48209 | 0.12593 | 0.221768 | 0 | 0 |
| 380 | sp\|Q5XJY5 | COPD_MOU | 57 kDa | 0.998264 | 7 | 100% | Q5XJY5 | Arcn1 | Coatomer s | 21.91961 | 22.57273 | 21.47775 | 22.50547 | 21.56178 | 21.68752 | -0.6022 | -1.42422 | 0.219825 | 21.95414 | -1.74372 | -5.48689 | 0.126115 | 0.221806 | 0 | 0 |
| 1002.1 | sp\|J3QMK6 | AL3B3_MO | 53 kDa | 0.995458 | 3 | 100% | J3QMK6 | Aldh3b3 | Aldehyde d | 22.21939 | 22.81456 | 21.61647 | 22.09614 | 22.11875 | 22.81778 | -0.59129 | -1.39911 | 0.216523 | 22.28052 | -1.74226 | -5.48889 | 0.126382 | 0.221988 | 0 | 0 |
| 709 | sp\|P09405 | NUCL_MOU | 77 kDa | 0.998826 | 4 | 100% | P09405 | Ncl | Nucleolin | 21.14682 | 21.73915 | 20.82734 | 21.32898 | 21.34954 | 21.79336 | -0.5126 | -1.21465 | 0.189454 | 21.3642 | -1.73783 | -5.49489 | 0.127181 | 0.223101 | 0 | 0 |
| 1077 | sp\|Q6NZR2 | MSD2_MOU | 61 kDa | 0.999153 | 2 | 100% | Q6NZR2 | Msantd2 | Myb/SANT- | 20.39327 | 21.18038 | 20.69604 | 21.30296 | 20.7497 | 20.82728 | -0.49054 | -1.16355 | 0.182477 | 20.85827 | -1.7348 | -5.49902 | 0.127739 | 0.223685 | 0 | 0 |
| 558 | sp\|P11352 | GPX1_MOU | 22 kDa | 0.99885 | 5 | 100% | P11352 | Gpx1 | Glutathion | 23.09901 | 21.90279 | 21.44827 | 21.09699 | 22.18951 | 21.22025 | 0.838921 | -0.3135 | 1.99134 | 21.82614 | 1.737246 | -5.49634 | 0.127843 | 0.223685 | 0 | 0 |
| 708 | sp\|P06745 | G6PI_MOU | 63 kDa | 0.998826 | 4 | 100% | P06745 | Gpi | Glucose-6- | 20.75272 | 20.49471 | 21.15124 | 20.06202 | 21.24761 | 20.89434 | 0.566832 | -0.21646 | 1.35012 | 20.76711 | 1.722398 | -5.51586 | 0.130043 | 0.227241 | 0 | 0 |
| 94 | sp\|P99024 | TBB5_MOU | 50 kDa | 0.999508 | 16 | 12% | P99024 | Tubb5 | Tubulin bet | 24.32673 | 23.33734 | 23.97248 | 23.32587 | 23.15299 | 22.78236 | 0.668879 | -0.26087 | 1.598628 | 23.48296 | 1.716852 | -5.52396 | 0.131642 | 0.229738 | 0 | 0 |
| 1028 | sp\|Q04447 | KCRB_MOU | 43 kDa | 0.999575 | 2 | 100% | Q04447 | Ckb | Creatine ki | 20.49903 | 20.64227 | 21.28593 | 19.82857 | 20.6927 | 20.18824 | 0.606192 | -0.24072 | 1.4531 | 20.52279 | 1.705553 | -5.53889 | 0.133474 | 0.232389 | 0 | 0 |
| 658 | sp\|Q9JJL8 | SYSM_MOU | 58 kDa | 0.999254 | 4 | 100% | Q9JJL8 | Sars2 | Serine--tRN | 21.71237 | 19.6845 | 19.78574 | 19.84282 | 21.36138 | 20.49805 | 0.9447 | -0.37597 | 2.265374 | 20.48081 | 1.707062 | -5.53717 | 0.133504 | 0.232389 | 0 | 0 |
| 538 | sp\|P62827 | RAN_MOUS | 24 kDa | 0.99919 | 5 | 100% | P62827 | Ran | GTP-bindin | 24.67969 | 24.23354 | 24.92683 | 24.34232 | 24.72777 | 24.44456 | 0.437956 | -0.17475 | 1.050659 | 24.55912 | 1.701299 | -5.54439 | 0.134056 | 0.233051 | 0 | 0 |
| 581 | sp\|Q9Z1Q2 | ABHGA_MO | 63 kDa | 0.997559 | 5 | 100% | Q9Z1Q2 | Abhd16a | Phosphatic | 20.92941 | 19.01781 | 21.03051 | 19.78409 | 20.58343 | 21.04451 | 0.898982 | -0.3616 | 2.159564 | 20.3983 | 1.701888 | -5.54414 | 0.134498 | 0.233519 | 0 | 0 |
| 784 | sp\|Q91WS | CISD1_MO | 12 kDa | 0.999709 | 3 | 100% | Q91WS0 | Cisd1 | CDGSH iror | 19.06946 | 20.72622 | 20.99448 | 20.75499 | 20.37419 | 22.02083 | -1.0213 | -2.45437 | 0.41177 | 20.65669 | -1.70074 | -5.54569 | 0.13472 | 0.233606 | 0 | 0 |
| 103 | sp\|Q9ESE1 | LRBA_MOU | 317 kDa | 0.999616 | 15 | 87% | Q9ESE1 | Lrba | Lipopolysa | 22.55533 | 22.18078 | 22.66058 | 21.9909 | 22.33142 | 22.05555 | 0.440037 | -0.18104 | 1.061111 | 22.29576 | 1.686345 | -5.56454 | 0.13697 | 0.236913 | 0 | 0 |
| 939 | sp\|Q60854 | SPB6_MOU | 43 kDa | 0.998046 | 3 | 100% | Q60854 | Serpinb6 | Serpin B6 | 20.36318 | 19.72682 | 20.19545 | 20.32601 | 20.44886 | 18.83408 | 0.706861 | -0.2918 | 1.705522 | 19.9824 | 1.689142 | -5.56128 | 0.136976 | 0.236913 | 0 | 0 |
| 455 | sp\|P27046 | MA2A1_MO | 132 kDa | 0.99896 | 6 | 100% | P27046 | Man2a1 | Alpha-man | 21.29452 | 20.97972 | 21.1154 | 22.428 | 21.24594 | 22.5988 | -0.78356 | -1.89997 | 0.332854 | 21.61039 | -1.67493 | -5.58033 | 0.139791 | 0.241473 | 0 | 0 |
| 591.1 | sp\|P01644 | KV5AB_MO | 12 kDa | 0.999808 | 4 | 50% | P01644 | NA | Ig kappa ch | 30.56848 | 30.99447 | 29.93028 | 30.48349 | 30.59641 | 31.45695 | -0.61324 | -1.48958 | 0.263096 | 30.67168 | -1.66823 | -5.58912 | 0.140917 | 0.24264 | 0 | 0 |
| 591.2 | sp\|P01645 | KV5AC_MO | 12 kDa | 0.999808 | 4 | 50% | P01645 | NA | Ig kappa ch | 30.56848 | 30.99447 | 29.93028 | 30.48349 | 30.59641 | 31.45695 | -0.61324 | -1.48958 | 0.263096 | 30.67168 | -1.66823 | -5.58912 | 0.140917 | 0.24264 | 0 | 0 |
| 1193 | sp\|Q9R0P6 | SC11A_MO | 21 kDa | 0.998027 | 2 | 100% | Q9R0P6 | Sec11a | Signal pept | 19.80913 | 19.89768 | 19.16466 | 19.66622 | 19.19866 | 20.19005 | -0.52717 | -1.28024 | 0.225906 | 19.6544 | -1.66614 | -5.59163 | 0.141003 | 0.24264 | 0 | 0 |

| | | | | | | | | | | | | | | | | | | | | | | | | | |
|---|---|---|---|---|---|---|---|---|---|---|---|---|---|---|---|---|---|---|---|---|---|---|---|---|---|
| 245 | sp\|Q9QZE5 | COPG1_MO | 98 kDa | 0.999019 | 9 | 78% | Q9QZE5 | Copg1 | Coatomer s | 21.82022 | 21.22272 | 22.36363 | 21.18759 | 21.36307 | 21.45236 | 0.561419 | -0.24255 | 1.365389 | 21.56827 | 1.662133 | -5.597 | 0.141826 | 0.243474 | 0 | 0 |
| 998 | sp\|P01636 | KV5A4_MO | 12 kDa | 0.995948 | 3 | 33% | P01636 | NA | Ig kappa ch | 15.55487 | 16.57059 | 11.80174 | 15.69161 | 14.5461 | 14.63078 | -1.66342 | -4.04801 | 0.721159 | 14.79928 | -1.66472 | -5.59397 | 0.141847 | 0.243474 | 0 | 0 |
| 334 | sp\|E9Q286 | ICE1_MOUS | 242 kDa | 0.999467 | 7 | 100% | E9Q286 | Ice1 | Little elong | 21.68771 | 22.01328 | 20.78232 | 22.21347 | 22.00889 | 22.13213 | -0.62666 | -1.52559 | 0.272277 | 21.8063 | -1.66187 | -5.59761 | 0.142206 | 0.243782 | 0 | 0 |
| 648 | sp\|Q8BZL8 | LMO3_MO | 17 kDa | 0.999349 | 4 | 25% | Q8BZL8 | Lmo3 | LIM domai | 20.00626 | 19.19275 | 20.54784 | 19.28175 | 19.32354 | 19.53644 | 0.622229 | -0.27263 | 1.517089 | 19.6481 | 1.657637 | -5.60325 | 0.14307 | 0.24488 | 0 | 0 |
| 713 | sp\|Q9QY76 | VAPB_MOU | 27 kDa | 0.998753 | 4 | 75% | Q9QY76 | Vapb | Vesicle-asso | 22.38498 | 21.55169 | 20.56986 | 20.79709 | 22.67565 | 19.97025 | 1.103821 | -0.48528 | 2.692922 | 21.32492 | 1.65767 | -5.60338 | 0.143284 | 0.24488 | 0 | 0 |
| 320 | sp\|Q8CBW | ABI1_MOU | 52 kDa | 0.999605 | 7 | 100% | Q8CBW3 | Abi1 | Abl interact | 22.78899 | 22.29972 | 21.21939 | 19.54282 | 22.51208 | 20.89948 | 1.259481 | -0.55427 | 3.073235 | 21.54375 | 1.657157 | -5.60406 | 0.143389 | 0.24488 | 0 | 0 |
| 1030.1 | tr\|A0A075 | A0A075B5 | 12 kDa | 0.999544 | 2 | 100% | A0A075B5 | Igkv17-127 | Immunoglo | 19.53636 | 19.24193 | 20.70039 | 19.70731 | 19.47109 | 17.9754 | 0.927737 | -0.41117 | 2.266641 | 19.43875 | 1.653582 | -5.60882 | 0.144123 | 0.245824 | 0 | 0 |
| 836.2 | tr\|A0A140 | A0A140T8 | 13 kDa | 0.999277 | 3 | 100% | A0A140T8 | Igkv14-111 | Immunoglo | 24.1277 | 24.30344 | 23.1277 | 25.28147 | 23.97254 | 23.92815 | -0.7617 | -1.86681 | 0.343406 | 24.1235 | -1.64487 | -5.6204 | 0.145927 | 0.248589 | 0 | 0 |
| 359 | sp\|P19096 | FAS_MOUS | 272 kDa | 0.999148 | 7 | 100% | P19096 | Fasn | Fatty acid s | 22.23164 | 21.60814 | 20.94065 | 22.22972 | 21.35763 | 23.94744 | -1.08512 | -2.66505 | 0.494807 | 22.05254 | -1.63905 | -5.62813 | 0.147144 | 0.250347 | 0 | 0 |
| 1069 | sp\|Q9JJK2 | LANC2_MO | 51 kDa | 0.999217 | 2 | 100% | Q9JJK2 | Lancl2 | LanC-like p | 19.56307 | 18.28328 | 20.40465 | 17.77966 | 19.43289 | 20.14441 | 1.064422 | -0.48929 | 2.618137 | 19.26799 | 1.634906 | -5.63362 | 0.148016 | 0.251515 | 0 | 0 |
| 732 | sp\|P97427 | DPYL1_MO | 62 kDa | 0.9984 | 4 | 25% | P97427 | Crmp1 | Dihydropyr | 21.39214 | 21.71855 | 22.07068 | 22.69001 | 21.04639 | 22.07159 | -0.65698 | -1.61689 | 0.302937 | 21.83156 | -1.63331 | -5.63573 | 0.148353 | 0.251773 | 0 | 0 |
| 954 | sp\|Q9Z2D6 | MECP2_MO | 52 kDa | 0.997848 | 3 | 100% | Q9Z2D6 | Mecp2 | Methyl-CpG | 21.28345 | 22.56401 | 21.75678 | 21.30964 | 20.97674 | 22.17584 | -0.6775 | -1.66852 | 0.313511 | 21.67774 | -1.63148 | -5.63816 | 0.14874 | 0.252115 | 0 | 0 |
| 682 | sp\|P31324 | KAP3_MOU | 46 kDa | 0.99905 | 4 | 75% | P31324 | Prkar2b | cAMP-depe | 19.3267 | 19.26679 | 19.49492 | 18.43125 | 19.51251 | 19.09077 | 0.515104 | -0.24099 | 1.2712 | 19.18716 | 1.621507 | -5.65102 | 0.150315 | 0.254466 | 0 | 0 |
| 1126 | sp\|Q8BG51 | MIRO1_MO | 72 kDa | 0.998666 | 2 | 100% | Q8BG51 | Rhot1 | Mitochond | 20.20886 | 20.71566 | 20.85144 | 21.28954 | 20.24017 | 20.86126 | -0.522 | -1.28958 | 0.245585 | 20.69449 | -1.61862 | -5.65484 | 0.150938 | 0.255195 | 0 | 0 |
| 714 | sp\|P80317 | TCPZ_MOU | 58 kDa | 0.998736 | 4 | 100% | P80317 | Cct6a | T-complex | 19.29978 | 19.64555 | 19.76247 | 20.61356 | 19.23408 | 19.79159 | -0.58479 | -1.4457 | 0.276129 | 19.72451 | -1.61931 | -5.65411 | 0.151122 | 0.255195 | 0 | 0 |
| 1221 | sp\|Q9D6J6 | NDUV2_MO | 27 kDa | 0.997654 | 2 | 100% | Q9D6J6 | Ndufv2 | NADH dehy | 22.13019 | 20.19584 | 20.13594 | 20.94936 | 22.87734 | 20.62026 | 1.126002 | -0.54083 | 2.79283 | 21.15149 | 1.612126 | -5.66371 | 0.152899 | 0.257322 | 0 | 0 |
| 280 | sp\|Q922B2 | SYDC_MOU | 57 kDa | 0.999384 | 8 | 100% | Q922B2 | Dars | Aspartate-- | 21.61051 | 21.51561 | 22.17834 | 21.29702 | 21.6656 | 21.24156 | 0.466756 | -0.22352 | 1.157031 | 21.58477 | 1.609416 | -5.66699 | 0.152935 | 0.257322 | 0 | 0 |
| 597 | sp\|Q8VIJ6 | SFPQ_MOU | 75 kDa | 0.999723 | 4 | 75% | Q8VIJ6 | Sfpq | Splicing fac | 22.94767 | 22.14397 | 22.44565 | 22.7139 | 22.78353 | 21.33934 | 0.659882 | -0.31709 | 1.636855 | 22.39568 | 1.611886 | -5.66403 | 0.152951 | 0.257322 | 0 | 0 |
| 81 | sp\|Q62167 | DDX3X_MO | 73 kDa | 0.999216 | 17 | 24% | Q62167 | Ddx3x | ATP-depen | 22.3851 | 20.92064 | 22.03355 | 21.79823 | 22.2409 | 22.12249 | 0.606064 | -0.29485 | 1.506975 | 21.91682 | 1.603726 | -5.67465 | 0.154517 | 0.259635 | 0 | 0 |
| 308 | sp\|Q3URS9 | CCD51_MO | 45 kDa | 0.999785 | 7 | 100% | Q3URS9 | Ccdc51 | Coiled-coil | 21.81858 | 19.80296 | 20.33623 | 20.88405 | 22.13494 | 20.88448 | 0.906089 | -0.4455 | 2.257682 | 20.97688 | 1.599837 | -5.67987 | 0.155595 | 0.2608 | 0 | 0 |
| 769 | sp\|P47962 | RL5_MOUS | 34 kDa | 0.997008 | 4 | 100% | P47962 | Rpl5 | 60S riboson | 20.29553 | 20.55558 | 20.1062 | 20.45196 | 19.40144 | 20.35779 | -0.52072 | -1.29662 | 0.255184 | 20.19475 | -1.59734 | -5.68288 | 0.155595 | 0.2608 | 0 | 0 |
| 1089 | sp\|Q9D6M | GHC1_MOU | 35 kDa | 0.999038 | 2 | 100% | Q9D6M3 | Slc25a22 | Mitochond | 20.1199 | 19.48487 | 19.73123 | 19.403 | 19.61848 | 19.25781 | 0.441311 | -0.21875 | 1.101371 | 19.60255 | 1.591337 | -5.69076 | 0.156933 | 0.262717 | 0 | 0 |
| 1246 | sp\|P31938 | MP2K1_MO | 43 kDa | 0.996991 | 2 | 100% | P31938 | Map2k1 | Dual specif | 22.18774 | 23.36572 | 20.20119 | 22.89809 | 22.16394 | 21.60415 | -1.10503 | -2.76553 | 0.555474 | 22.07014 | -1.58813 | -5.69521 | 0.158206 | 0.264522 | 0 | 0 |
| 850 | tr\|A0A140 | A0A140T8 | 13 kDa | 0.999137 | 3 | 100% | A0A140T8 | Igkv6-32 | Immunoglo | 21.53047 | 22.3488 | 21.3377 | 22.36287 | 22.07352 | 21.80139 | -0.52379 | -1.31132 | 0.263743 | 21.90912 | -1.58303 | -5.70165 | 0.158802 | 0.265191 | 0 | 0 |
| 661 | sp\|P10922 | H10_MOUS | 21 kDa | 0.999226 | 4 | 100% | P10922 | H1f0 | Histone H1 | 24.15853 | 23.79194 | 23.69151 | 23.53611 | 24.62709 | 23.59527 | 0.517935 | -0.26154 | 1.297408 | 23.90007 | 1.581521 | -5.70362 | 0.159144 | 0.265436 | 0 | 0 |
| 71 | sp\|P05784 | K1C18_MO | 48 kDa | 0.999757 | 17 | 82% | P05784 | Krt18 | Keratin, typ | 24.42293 | 23.90281 | 24.39502 | 24.4042 | 25.01887 | 24.04625 | 0.494518 | -0.25144 | 1.24048 | 24.36502 | 1.577855 | -5.70841 | 0.159977 | 0.266497 | 0 | 0 |
| 1219 | tr\|A0A075 | A0A075B5 | 11 kDa | 0.997683 | 2 | 50% | A0A075B5 | Ighv1-67 | Immunoglo | 18.28485 | 18.9101 | 19.63178 | 20.75041 | 19.16561 | 20.13085 | -0.90304 | -2.26833 | 0.462241 | 19.47893 | -1.57847 | -5.70783 | 0.16039 | 0.266857 | 0 | 0 |
| 1026 | sp\|P63242 | IF5A1_MOU | 17 kDa | 0.999606 | 2 | 100% | P63242 | Eif5a | Eukaryotic | 19.69219 | 22.40331 | 21.48355 | 23.10251 | 23.93995 | 25.12685 | -1.839 | -4.62536 | 0.947373 | 22.62473 | -1.57504 | -5.71229 | 0.161171 | 0.267828 | 0 | 0 |
| 296 | sp\|P33175 | KIF5A_MOU | 117 kDa | 0.998745 | 8 | 12% | P33175 | Kif5a | Kinesin hea | 20.31659 | 18.92504 | 18.20802 | 18.88076 | 20.677 | 18.26149 | 1.044776 | -0.54016 | 2.629718 | 19.21148 | 1.573116 | -5.71481 | 0.161612 | 0.268233 | 0 | 0 |
| 549 | sp\|Q9DBE8 | ALG2_MOU | 47 kDa | 0.998978 | 5 | 100% | Q9DBE8 | Alg2 | Alpha-1,3/1 | 22.52353 | 22.30899 | 22.37436 | 21.4002 | 22.23581 | 21.89774 | 0.508925 | -0.26412 | 1.281966 | 22.12344 | 1.566937 | -5.72266 | 0.162482 | 0.269307 | 0 | 0 |
| 1211 | sp\|Q3UMB | WASC4_MO | 136 kDa | 0.997787 | 2 | 100% | Q3UMB9 | Washc4 | WASH com | 14.9721 | 17.51921 | 12.97503 | 18.14232 | 17.28625 | 15.24019 | -1.88945 | -4.76407 | 0.985176 | 16.02252 | -1.56857 | -5.72072 | 0.162657 | 0.269307 | 0 | 0 |
| 1119.1 | sp\|C0HK79 | ARXS1_MO | 20 kDa | 0.998757 | 2 | 100% | C0HK79 | Arxes1 | Adipocyte- | 20.79483 | 20.16016 | 21.48443 | 20.81437 | 20.45198 | 19.88434 | 0.624119 | -0.33068 | 1.578914 | 20.59835 | 1.559939 | -5.73194 | 0.164659 | 0.272289 | 0 | 0 |
| 1168 | sp\|Q8VE97 | SRSF4_MO | 56 kDa | 0.998262 | 2 | 50% | Q8VE97 | Srsf4 | Serine/argi | 19.61238 | 19.2232 | 18.76235 | 19.98669 | 19.16938 | 20.07829 | -0.58136 | -1.47177 | 0.309057 | 19.47205 | -1.55649 | -5.73635 | 0.165247 | 0.272413 | 0 | 0 |
| 82 | sp\|Q7TMM | TBB2A_MO | 50 kDa | 0.999055 | 17 | 6% | Q7TMM9 | Tubb2a | Tubulin bet | 22.94849 | 22.60223 | 23.258 | 22.37028 | 22.49043 | 22.2841 | 0.480102 | -0.25483 | 1.215036 | 22.65892 | 1.554842 | -5.73839 | 0.1653 | 0.272413 | 0 | 0 |
| 807 | sp\|P35922 | FMR1_MOU | 69 kDa | 0.999463 | 3 | 33% | P35922 | Fmr1 | Synaptic fu | 22.07779 | 21.50725 | 22.6493 | 21.71026 | 21.96177 | 22.00088 | 0.490158 | -0.26038 | 1.240697 | 21.98454 | 1.554401 | -5.73896 | 0.165404 | 0.272413 | 0 | 0 |
| 573 | sp\|Q6PHZ2 | KCC2D_MO | 56 kDa | 0.998209 | 5 | 60% | Q6PHZ2 | Camk2d | Calcium/ca | 21.98621 | 23.10317 | 21.25374 | 23.33645 | 22.40286 | 21.69289 | -0.8299 | -2.10257 | 0.442771 | 22.29589 | -1.55618 | -5.73681 | 0.165537 | 0.272413 | 0 | 0 |
| 680.1 | sp\|P0DP26 | CALM1_MO | 17 kDa | 0.99906 | 4 | 100% | P0DP26 | Calm1 | Calmodulin | 21.50927 | 20.84945 | 21.26034 | 21.24079 | 21.23826 | 20.51891 | 0.466241 | -0.25108 | 1.183559 | 21.10284 | 1.547029 | -5.74852 | 0.167145 | 0.273785 | 0 | 0 |
| 680.2 | sp\|P0DP27 | CALM2_MO | 17 kDa | 0.99906 | 4 | 100% | P0DP27 | Calm2 | Calmodulin | 21.50927 | 20.84945 | 21.26034 | 21.24079 | 21.23826 | 20.51891 | 0.466241 | -0.25108 | 1.183559 | 21.10284 | 1.547029 | -5.74852 | 0.167145 | 0.273785 | 0 | 0 |
| 680.3 | sp\|P0DP28 | CALM3_MO | 17 kDa | 0.99906 | 4 | 100% | P0DP28 | Calm3 | Calmodulin | 21.50927 | 20.84945 | 21.26034 | 21.24079 | 21.23826 | 20.51891 | 0.466241 | -0.25108 | 1.183559 | 21.10284 | 1.547029 | -5.74852 | 0.167145 | 0.273785 | 0 | 0 |
| 521 | sp\|P50481 | LHX3_MOU | 44 kDa | 0.999329 | 5 | 100% | P50481 | Lhx3 | LIM/homeo | 20.32543 | 20.08119 | 20.6918 | 17.99755 | 19.63035 | 19.75225 | 0.938862 | -0.50738 | 2.385107 | 19.74643 | 1.549211 | -5.74584 | 0.167178 | 0.273785 | 0 | 0 |
| 948 | sp\|Q922U1 | PRPF3_MO | 77 kDa | 0.997918 | 3 | 100% | Q922U1 | Prpf3 | U4/U6 sma | 19.899 | 21.03168 | 17.8648 | 20.67317 | 21.16724 | 20.87095 | -1.21492 | -3.0918 | 0.661955 | 20.25114 | -1.54477 | -5.75158 | 0.168232 | 0.275143 | 0 | 0 |
| 99 | sp\|Q8BMS1 | ECHA_MOU | 83 kDa | 0.999761 | 15 | 100% | Q8BMS1 | Hadha | Trifunction | 23.52318 | 23.5922 | 24.0243 | 23.32312 | 23.7086 | 22.89657 | 0.481398 | -0.26185 | 1.224646 | 23.51133 | 1.541598 | -5.75554 | 0.168438 | 0.275143 | 0 | 0 |
| 55 | sp\|P14733 | LMNB1_MO | 67 kDa | 0.999761 | 19 | 95% | P14733 | Lmnb1 | Lamin-B1 | 24.72842 | 25.12719 | 24.94358 | 25.52291 | 24.66242 | 24.99112 | -0.4356 | -1.10862 | 0.237419 | 24.99594 | -1.5405 | -5.75696 | 0.168701 | 0.275143 | 0 | 0 |
| 181 | sp\|E9Q557 | DESP_MOU | 333 kDa | 0.998772 | 11 | 91% | E9Q557 | Dsp | Desmoplak | 26.4519 | 27.72025 | 25.8171 | 26.80747 | 27.65173 | 27.94761 | -0.85153 | -2.16928 | 0.466223 | 27.06601 | -1.54211 | -5.75501 | 0.168864 | 0.275143 | 0 | 0 |
| 991 | sp\|Q6ZWY3 | RS27L_MO | 9 kDa | 0.996755 | 3 | 33% | Q6ZWY3 | Rps27l | 40S riboson | 18.43975 | 19.48885 | 19.44739 | 19.30052 | 19.01831 | 19.71013 | -0.53135 | -1.35324 | 0.290544 | 19.23416 | -1.53882 | -5.75914 | 0.169114 | 0.275143 | 0 | 0 |
| 339 | sp\|P97467 | AMD_MOU | 109 kDa | 0.999426 | 7 | 100% | P97467 | Pam | Peptidyl-gl | 22.78346 | 22.61932 | 22.60715 | 22.17065 | 22.95593 | 22.27116 | 0.42847 | -0.23447 | 1.091414 | 22.56794 | 1.538311 | -5.75979 | 0.169225 | 0.275143 | 0 | 0 |
| 228 | sp\|P63239 | NEC1_MOU | 84 kDa | 0.999749 | 9 | 89% | P63239 | Pcsk1 | Neuroendo | 23.54351 | 23.21976 | 23.43057 | 22.859 | 23.50683 | 23.18907 | 0.40436 | -0.22185 | 1.030567 | 23.29146 | 1.536919 | -5.76159 | 0.16956 | 0.275357 | 0 | 0 |
| 1171 | sp\|Q9JLI8 | SART3_MO | 110 kDa | 0.998231 | 2 | 100% | Q9JLI8 | Sart3 | Squamous c | 24.53754 | 24.33501 | 24.85107 | 24.11704 | 24.42377 | 24.11128 | 0.41635 | -0.22905 | 1.061747 | 24.39595 | 1.53544 | -5.7635 | 0.169915 | 0.275604 | 0 | 0 |
| 761 | sp\|P48453 | PP2BB_MO | 59 kDa | 0.997338 | 4 | 100% | P48453 | Ppp3cb | Serine/thre | 18.54963 | 18.63115 | 18.91097 | 20.08064 | 18.65731 | 19.31212 | -0.63533 | -1.62261 | 0.35194 | 19.02364 | -1.53573 | -5.76324 | 0.170395 | 0.276052 | 0 | 0 |
| 1117 | sp\|P47757 | CAPZB_MO | 31 kDa | 0.998764 | 2 | 100% | P47757 | Capzb | F-actin-cap | 20.26947 | 20.08153 | 20.29621 | 19.23993 | 20.47194 | 20.16228 | 0.517961 | -0.28701 | 1.322931 | 20.08689 | 1.531576 | -5.76848 | 0.170859 | 0.276278 | 0 | 0 |
| 835 | tr\|V9GX38 | V9GX38_M | 38 kDa | 0.999287 | 3 | 33% | V9GX38 | Gm17190 | Predicted g | 19.21437 | 20.68553 | 19.61596 | 19.31395 | 19.52128 | 20.22245 | -0.62344 | -1.59369 | 0.34681 | 19.76226 | -1.53342 | -5.76621 | 0.170951 | 0.276278 | 0 | 0 |
| 1095 | tr\|A0A075 | A0A075B5 | 10 kDa | 0.999003 | 2 | 50% | A0A075B5 | Igkv5-39 | Immunoglo | 19.70284 | 19.95872 | 20.11754 | 19.06697 | 19.98818 | 19.23474 | 0.516042 | -0.28655 | 1.318634 | 19.67816 | 1.530351 | -5.77006 | 0.171145 | 0.276278 | 0 | 0 |
| 231 | sp\|Q9Z315 | SNUT1_MO | 91 kDa | 0.999674 | 9 | 100% | Q9Z315 | Sart1 | U4/U6.U5 t | 20.94027 | 21.37044 | 20.48983 | 21.14614 | 21.17071 | 21.45395 | -0.45657 | -1.16768 | 0.254536 | 21.09522 | -1.52818 | -5.77285 | 0.171672 | 0.276799 | 0 | 0 |
| 666 | sp\|P62849 | RS24_MOU | 15 kDa | 0.999157 | 4 | 100% | P62849 | Rps24 | 40S riboson | 22.15371 | 21.20951 | 21.4961 | 20.34672 | 21.88396 | 21.91178 | 0.688588 | -0.38846 | 1.76564 | 21.5003 | 1.525715 | -5.77612 | 0.172822 | 0.278112 | 0 | 0 |
| 1235 | sp\|Q02248 | CTNB1_MO | 85 kDa | 0.997288 | 2 | 100% | Q02248 | Ctnnb1 | Catenin bet | 18.0649 | 16.35402 | 18.39154 | 18.53567 | 17.51481 | 15.79664 | 1.09497 | -0.61902 | 2.808956 | 17.44293 | 1.524564 | -5.7776 | 0.173102 | 0.278112 | 0 | 0 |
| 1236 | sp\|Q7TMY8 | HUWE1_M | 483 kDa | 0.997286 | 2 | 100% | Q7TMY8 | Huwe1 | E3 ubiquiti | 18.0649 | 16.35402 | 18.39154 | 18.53567 | 17.51481 | 15.79664 | 1.09497 | -0.61902 | 2.808956 | 17.44293 | 1.524564 | -5.7776 | 0.173102 | 0.278112 | 0 | 0 |
| 742 | sp\|Q8BGC4 | PTGR3_MO | 41 kDa | 0.997995 | 4 | 100% | Q8BGC4 | Zadh2 | Prostagland | 20.78401 | 22.5345 | 21.10698 | 20.55711 | 20.97627 | 22.24272 | -0.82236 | -2.11124 | 0.466515 | 21.36693 | -1.52266 | -5.78005 | 0.173569 | 0.278266 | 0 | 0 |
| 134 | tr\|F8VQC1 | F8VQC1_M | 75 kDa | 0.999512 | 13 | 100% | F8VQC1 | Srp72 | Signal recog | 23.30278 | 28.82273 | 23.30846 | 22.81869 | 23.55645 | 24.85482 | -2.10952 | -5.41609 | 1.197052 | 24.44399 | -1.5225 | -5.78026 | 0.173608 | 0.278266 | 0 | 0 |
| 533 | sp\|Q3UEB3 | PUF60_MO | 60 kDa | 0.99922 | 5 | 100% | Q3UEB3 | Puf60 | Poly(U)-bin | 21.10803 | 22.10785 | 20.97972 | 20.94478 | 21.21398 | 21.97337 | -0.57476 | -1.47611 | 0.326596 | 21.38795 | -1.52014 | -5.78324 | 0.173969 | 0.278515 | 0 | 0 |
| 1131 | sp\|Q9JKX6 | NUDT5_MO | 24 kDa | 0.99861 | 2 | 100% | Q9JKX6 | Nudt5 | ADP-sugar p | 20.77212 | 19.03802 | 21.93863 | 21.74438 | 21.59847 | 20.32745 | 1.066458 | -0.61433 | 2.747243 | 20.90318 | 1.514197 | -5.79089 | 0.175652 | 0.280546 | 0 | 0 |
| 1132 | sp\|P67871 | CSK2B_MO | 25 kDa | 0.998588 | 2 | 100% | P67871 | Csnk2b | Casein kina | 20.77212 | 19.03802 | 21.93863 | 21.74438 | 21.59847 | 20.32745 | 1.066458 | -0.61433 | 2.747243 | 20.90318 | 1.514197 | -5.79089 | 0.175652 | 0.280546 | 0 | 0 |
| 145 | sp\|P52480 | KPYM_MOU | 58 kDa | 0.99964 | 12 | 100% | P52480 | Pkm | Pyruvate ki | 24.438 | 24.67056 | 24.85356 | 25.20332 | 24.62281 | 25.48808 | -0.48253 | -1.24671 | 0.281638 | 24.87939 | -1.50293 | -5.80523 | 0.177916 | 0.283828 | 0 | 0 |
| 1189 | sp\|O89051 | ITM2B_MO | 30 kDa | 0.998086 | 2 | 100% | O89051 | Itm2b | Integral me | 17.42239 | 17.98042 | 18.1624 | 12.28564 | 17.16919 | 16.38427 | 2.03455 | -1.19426 | 5.263359 | 16.56738 | 1.503754 | -5.80423 | 0.178255 | 0.283854 | 0 | 0 |
| 799 | sp\|Q9CX86 | ROA0_MOU | 31 kDa | 0.999526 | 3 | 100% | Q9CX86 | Hnrnpa0 | Heterogene | 20.31193 | 22.76135 | 23.80267 | 25.02931 | 19.66552 | 21.72003 | -1.91019 | -4.94357 | 1.123187 | 22.21513 | -1.5028 | -5.80545 | 0.178494 | 0.283854 | 0 | 0 |
| 445 | sp\|P57780 | ACTN4_MO | 105 kDa | 0.999129 | 6 | 100% | P57780 | Actn4 | Alpha-actin | 21.12255 | 20.96159 | 20.70941 | 21.58629 | 20.60384 | 21.27276 | -0.46161 | -1.19389 | 0.270672 | 21.04274 | -1.50037 | -5.8085 | 0.17856 | 0.283854 | 0 | 0 |
| 6 | sp\|Q9JHU4 | DYHC1_MO | 532 kDa | 0.999666 | 50 | 100% | Q9JHU4 | Dync1h1 | Cytoplasmi | 25.49044 | 26.32785 | 25.53898 | 25.23382 | 25.2618 | 26.5393 | -0.60325 | -1.56291 | 0.356414 | 25.73203 | -1.50013 | -5.80885 | 0.179166 | 0.283972 | 0 | 0 |
| 1051 | sp\|P58281 | OPA1_MOU | 111 kDa | 0.999313 | 2 | 100% | P58281 | Opa1 | Dynamin-li | 23.23577 | 22.40208 | 22.09365 | 22.37312 | 23.05695 | 21.84288 | 0.589432 | -0.34849 | 1.527354 | 22.50074 | 1.499748 | -5.80933 | 0.179262 | 0.283972 | 0 | 0 |
| 1052 | sp\|Q9R0P9 | UCHL1_MO | 25 kDa | 0.99931 | 2 | 100% | Q9R0P9 | Uchl1 | Ubiquitin c | 23.23577 | 22.40208 | 22.09365 | 22.37312 | 23.05695 | 21.84288 | 0.589432 | -0.34849 | 1.527354 | 22.50074 | 1.499748 | -5.80933 | 0.179262 | 0.283972 | 0 | 0 |
| 1173 | sp\|P55249 | LX12E_MO | 75 kDa | 0.998211 | 2 | 100% | P55249 | Alox12e | Arachidona | 18.10271 | 17.45414 | 19.11794 | 18.08479 | 17.63845 | 17.2586 | 0.68719 | -0.40822 | 1.782596 | 17.94277 | 1.497106 | -5.8127 | 0.17993 | 0.284697 | 0 | 0 |
| 768 | sp\|P28660 | NCKP1_MO | 129 kDa | 0.997017 | 4 | 100% | P28660 | Nckap1 | Nck-associa | 21.67127 | 20.4981 | 21.94754 | 21.20502 | 21.66573 | 21.82739 | 0.584678 | -0.34827 | 1.517631 | 21.46918 | 1.495574 | -5.81465 | 0.180318 | 0.284954 | 0 | 0 |
| 525 | sp\|P35455 | NEU2_MOU | 18 kDa | 0.999285 | 5 | 80% | P35455 | Avp | Vasopressin | 19.70739 | 25.82576 | 22.65475 | 23.37696 | 23.98801 | 22.72488 | -1.85915 | -4.82726 | 1.10896 | 23.04629 | -1.49481 | -5.81562 | 0.180513 | 0.284954 | 0 | 0 |
| 942 | sp\|P01668 | KV3AG_MO | 12 kDa | 0.998019 | 3 | 33% | P01668 | NA | Ig kappa ch | 16.2335 | 16.01487 | 16.56614 | 19.27991 | 17.00409 | 18.07093 | -1.18732 | -3.08422 | 0.709571 | 17.19491 | -1.49374 | -5.81697 | 0.180783 | 0.285048 | 0 | 0 |
| 752 | sp\|Q8BW4 | PMGT2_M | 69 kDa | 0.997653 | 4 | 100% | Q8BW41 | Pomgnt2 | Protein O-l | 19.13876 | 19.83357 | 19.25925 | 21.29196 | 18.27615 | 18.62164 | -1.02434 | -2.66515 | 0.616473 | 19.40355 | -1.48983 | -5.82195 | 0.181781 | 0.28629 | 0 | 0 |
| 683 | sp\|P35293 | RAB18_MO | 23 kDa | 0.999044 | 4 | 75% | P35293 | Rab18 | Ras-related | 20.96792 | 21.09972 | 20.77681 | 21.18074 | 20.53618 | 21.18275 | -0.3941 | -1.02538 | 0.237174 | 20.95735 | -1.4859 | -5.82691 | 0.182242 | 0.286396 | 0 | 0 |
| 188 | sp\|Q7TPV4 | MBB1A_M | 152 kDa | 0.999675 | 10 | 100% | Q7TPV4 | Mybbp1a | Myb-bindin | 22.38563 | 22.14493 | 22.77721 | 22.26044 | 22.49628 | 22.07004 | 0.394568 | -0.2375 | 1.026638 | 22.35576 | 1.485788 | -5.82705 | 0.182271 | 0.286396 | 0 | 0 |
| 1063.2 | tr\|A0A075 | A0A075B6 | 37 kDa | 0.999242 | 2 | 100% | A0A075B6 | Igha | Immunoglo | 22.16911 | 21.41782 | 22.21547 | 21.74372 | 21.77535 | 21.74489 | 0.417832 | -0.25194 | 1.087601 | 21.84439 | 1.484834 | -5.82827 | 0.182516 | 0.286449 | 0 | 0 |
| 493 | tr\|A0A075 | A0A075B5 | 10 kDa | 0.999661 | 5 | 40% | A0A075B5 | Igkv6-13 | Immunoglo | 23.9316 | 26.1743 | 21.65218 | 23.36178 | 21.12807 | 22.27183 | -1.69869 | -4.42912 | 1.031741 | 23.08663 | -1.48468 | -5.82847 | 0.183099 | 0.287032 | 0 | 0 |
| 454 | sp\|Q9JKF1 | IQGA1_MO | 189 kDa | 0.998976 | 6 | 100% | Q9JKF1 | Iqgap1 | Ras GTPase | 20.51589 | 21.37397 | 20.64323 | 20.4434 | 19.80473 | 20.8257 | -0.55974 | -1.46383 | 0.34435 | 20.60115 | -1.47594 | -5.83954 | 0.185145 | 0.289903 | 0 | 0 |
| 868 | sp\|O09106 | HDAC1_MO | 55 kDa | 0.998995 | 3 | 100% | O09106 | Hdac1 | Histone dea | 20.94364 | 19.98263 | 19.54545 | 19.88414 | 20.646 | 19.32672 | 0.647203 | -0.39984 | 1.69425 | 20.05476 | 1.475114 | -5.84058 | 0.185576 | 0.290243 | 0 | 0 |
| 1023 | sp\|Q922V4 | PLRG1_MO | 57 kDa | 0.999706 | 2 | 100% | Q922V4 | Plrg1 | Pleiotropic | 20.29109 | 20.47224 | 19.97076 | 21.5589 | 19.66128 | 19.93901 | -0.68234 | -1.78896 | 0.424269 | 20.31554 | -1.47149 | -5.84515 | 0.18652 | 0.291384 | 0 | 0 |
| 665 | sp\|Q8R4X3 | RBM12_MO | 103 kDa | 0.999174 | 4 | 100% | Q8R4X3 | Rbm12 | RNA-bindin | 21.42093 | 22.19443 | 20.42327 | 21.16665 | 21.4883 | 21.87072 | -0.6331 | -1.66062 | 0.394418 | 21.42738 | -1.47039 | -5.84654 | 0.186808 | 0.291499 | 0 | 0 |
| 404 | sp\|Q9JIF7 | COPB_MOU | 107 kDa | 0.999621 | 6 | 100% | Q9JIF7 | Copb1 | Coatomer s | 21.62258 | 21.66188 | 22.07055 | 20.78293 | 21.48591 | 21.22945 | 0.501593 | -0.31278 | 1.315963 | 21.47555 | 1.466061 | -5.85202 | 0.187414 | 0.292018 | 0 | 0 |
| 78 | sp\|P29341 | PABP1_MO | 71 kDa | 0.999579 | 17 | 88% | P29341 | Pabpc1 | Polyadenyl | 25.00281 | 21.99944 | 23.58065 | 21.92393 | 24.69661 | 25.22812 | 1.37619 | -0.86179 | 3.61417 | 23.73859 | 1.467483 | -5.85021 | 0.187572 | 0.292018 | 0 | 0 |
| 1253 | sp\|Q99K01 | PDXD1_MO | 87 kDa | 0.996729 | 2 | 100% | Q99K01 | Pdxdc1 | Pyridoxal-d | 19.63986 | 19.35901 | 19.33071 | 20.2351 | 19.73239 | 23.0122 | -1.30112 | -3.42569 | 0.823451 | 20.21821 | -1.46149 | -5.85775 | 0.189153 | 0.294141 | 0 | 0 |
| 1181 | sp\|P15508 | SPTB1_MO | 245 kDa | 0.998147 | 2 | 100% | P15508 | Sptb | Spectrin be | 20.9722 | 20.28426 | 21.14096 | 20.10071 | 20.66647 | 20.94354 | 0.483705 | -0.30819 | 1.275604 | 20.68469 | 1.453824 | -5.86742 | 0.190654 | 0.296136 | 0 | 0 |
| 531 | sp\|Q7TN33 | CELF6_MO | 48 kDa | 0.999224 | 5 | 100% | Q7TN33 | Celf6 | CUGBP Elav | 22.46618 | 22.44068 | 21.56423 | 22.07976 | 21.39844 | 22.51871 | -0.53677 | -1.41692 | 0.343383 | 22.078 | -1.45386 | -5.86734 | 0.19097 | 0.296287 | 0 | 0 |
| 383 | sp\|Q6PH08 | ERC2_MOU | 111 kDa | 0.998165 | 7 | 100% | Q6PH08 | Erc2 | ERC protein | 19.38581 | 19.82916 | 18.82525 | 21.31985 | 21.08043 | 21.08828 | -0.98193 | -2.60431 | 0.640441 | 20.2548 | -1.44438 | -5.8792 | 0.193736 | 0.299975 | 0 | 0 |
| 427 | sp\|P08752 | GNAI2_MO | 40 kDa | 0.999401 | 6 | 50% | P08752 | Gnai2 | Guanine nu | 20.31076 | 20.35625 | 21.4175 | 20.17157 | 20.58204 | 20.23514 | 0.515778 | -0.336 | 1.367553 | 20.51221 | 1.443223 | -5.88067 | 0.193789 | 0.299975 | 0 | 0 |
| 262 | sp\|Q91ZR2 | SNX18_MO | 68 kDa | 0.999726 | 8 | 100% | Q91ZR2 | Snx18 | Sorting nex | 25.12914 | 22.76566 | 23.16367 | 22.64179 | 25.67968 | 24.90054 | 1.221498 | -0.81592 | 3.258911 | 24.04675 | 1.430751 | -5.89619 | 0.197457 | 0.305305 | 0 | 0 |
| 822 | sp\|Q9D358 | PPAC_MOU | 18 kDa | 0.999395 | 3 | 100% | Q9D358 | Acp1 | Low molec | 20.3845 | 21.89027 | 19.23984 | 20.2441 | 17.99841 | 19.13001 | -1.21388 | -3.24866 | 0.820911 | 19.81452 | -1.42366 | -5.90499 | 0.199419 | 0.307987 | 0 | 0 |
| 598 | sp\|Q9DB77 | QCR2_MOU | 48 kDa | 0.999719 | 4 | 100% | Q9DB77 | Uqcrc2 | Cytochrom | 20.6501 | 21.16469 | 21.97187 | 22.2635 | 20.61804 | 21.90472 | -0.69763 | -1.86898 | 0.473716 | 21.42882 | -1.42132 | -5.9079 | 0.200071 | 0.308642 | 0 | 0 |
| 818 | sp\|Q9R1Q9 | VAS1_MOU | 51 kDa | 0.999414 | 3 | 100% | Q9R1Q9 | Atp6ap1 | V-type prot | 20.97313 | 21.84734 | 21.39982 | 22.21377 | 20.63937 | 20.82034 | -0.62304 | -1.66982 | 0.423733 | 21.31563 | -1.42041 | -5.90902 | 0.200323 | 0.30868 | 0 | 0 |
| 1103 | sp\|P28658 | ATX10_MO | 54 kDa | 0.998947 | 2 | 100% | P28658 | Atxn10 | Ataxin-10 | 20.16387 | 21.32252 | 21.62698 | 20.57814 | 19.67945 | 22.24518 | -0.89184 | -2.407 | 0.623311 | 20.93602 | -1.4047 | -5.92843 | 0.204752 | 0.315146 | 0 | 0 |
| 943 | sp\|P35283 | RAB12_MO | 27 kDa | 0.998013 | 3 | 67% | P35283 | Rab12 | Ras-related | 19.41794 | 19.38581 | 19.9271 | 18.94686 | 19.13382 | 18.7218 | 0.474797 | -0.33195 | 1.281539 | 19.25555 | 1.40086 | -5.93327 | 0.205324 | 0.315578 | 0 | 0 |
| 1070 | sp\|Q921K9 | BCL7B_MO | 22 kDa | 0.999205 | 2 | 50% | Q921K9 | Bcl7b | B-cell CLL/l | 17.47632 | 18.15401 | 18.98205 | 18.70716 | 17.57501 | 19.38307 | -0.73696 | -1.99131 | 0.517395 | 18.3796 | -1.40208 | -5.93165 | 0.205498 | 0.315578 | 0 | 0 |
| 516 | sp\|O88569 | ROA2_MOU | 37 kDa | 0.999369 | 5 | 100% | O88569 | Hnrnpa2b1 | Heterogene | 22.61404 | 22.24579 | 22.23639 | 22.64133 | 23.21183 | 21.26329 | 0.637284 | -0.45107 | 1.725638 | 22.36878 | 1.397378 | -5.93743 | 0.206844 | 0.317286 | 0 | 0 |
| 568 | sp\|P62317 | SMD2_MOU | 14 kDa | 0.998349 | 5 | 100% | P62317 | Snrpd2 | Small nucle | 24.09942 | 24.53585 | 23.38422 | 24.60029 | 24.5672 | 24.43173 | -0.50568 | -1.37111 | 0.359758 | 24.26979 | -1.39294 | -5.94293 | 0.207911 | 0.318563 | 0 | 0 |
| 362 | sp\|P13020 | GELS_MOU | 86 kDa | 0.999091 | 7 | 100% | P13020 | Gsn | Gelsolin | 21.74806 | 22.1444 | 22.07291 | 22.40759 | 21.85665 | 22.22665 | -0.36701 | -0.99512 | 0.261108 | 22.07605 | -1.39071 | -5.94575 | 0.208237 | 0.318701 | 0 | 0 |
| 1209 | sp\|Q7TT50 | MRCKB_M | 195 kDa | 0.997802 | 2 | 100% | Q7TT50 | Cdc42bpb | Serine/thre | 15.95387 | 16.43879 | 17.26305 | 16.34319 | 16.7925 | 15.13228 | 0.698387 | -0.50136 | 1.898139 | 16.32062 | 1.389171 | -5.94749 | 0.209213 | 0.319834 | 0 | 0 |
| 684 | sp\|O70152 | DPM1_MO | 29 kDa | 0.999033 | 4 | 100% | O70152 | Dpm1 | Dolichol-ph | 21.71079 | 21.51958 | 21.43645 | 22.1934 | 21.49215 | 22.20897 | -0.42752 | -1.16226 | 0.307224 | 21.76022 | -1.38491 | -5.95285 | 0.209924 | 0.32056 | 0 | 0 |
| 704 | sp\|Q9JIS5 | SV2A_MOU | 83 kDa | 0.998858 | 4 | 100% | Q9JIS5 | Sv2a | Synaptic ve | 22.04964 | 21.73632 | 22.53426 | 22.16459 | 22.27226 | 21.75317 | 0.400694 | -0.28846 | 1.089852 | 22.08504 | 1.383869 | -5.95412 | 0.210227 | 0.320662 | 0 | 0 |
| 1274 | sp\|E9Q3L2 | PI4KA_MO | 237 kDa | 0.995953 | 2 | 100% | E9Q3L2 | Pi4ka | Phosphatid | 19.80407 | 19.23859 | 18.72606 | 17.14825 | 19.66021 | 19.16098 | 0.880835 | -0.64237 | 2.404038 | 18.95636 | 1.380027 | -5.95866 | 0.211881 | 0.322467 | 0 | 0 |
| 1054 | sp\|Q80UU9 | PGRC2_MO | 23 kDa | 0.99929 | 2 | 50% | Q80UU9 | Pgrmc2 | Membrane | 18.21978 | 17.55232 | 19.34481 | 14.31366 | 17.49705 | 18.60673 | 1.529647 | -1.11556 | 4.174858 | 17.58906 | 1.380008 | -5.95868 | 0.211887 | 0.322467 | 0 | 0 |
| 933 | sp\|P01592 | IGJ_MOUSE | 18 kDa | 0.998193 | 3 | 100% | P01592 | Jchain | Immunoglo | 22.27519 | 22.79066 | 22.75113 | 22.5666 | 21.48053 | 22.70681 | -0.51908 | -1.42437 | 0.386217 | 22.42849 | -1.3669 | -5.9747 | 0.215556 | 0.327613 | 0 | 0 |
| 173 | sp\|Q8BKI2 | TNR6B_MO | 192 kDa | 0.999298 | 11 | 100% | Q8BKI2 | Tnrc6b | Trinucleoti | 22.3601 | 21.52462 | 21.34639 | 20.98332 | 22.02357 | 21.68924 | 0.51096 | -0.3806 | 1.402523 | 21.65454 | 1.366245 | -5.97549 | 0.215751 | 0.327613 | 0 | 0 |
| 707 | sp\|Q6NZL0 | SOGA3_MO | 103 kDa | 0.998834 | 4 | 50% | Q6NZL0 | Soga3 | Protein SO | 18.26011 | 14.90411 | 17.94117 | 17.57236 | 18.01283 | 18.30415 | 1.144496 | -0.85604 | 3.145028 | 17.49912 | 1.365272 | -5.9766 | 0.21625 | 0.328003 | 0 | 0 |
| 444 | sp\|Q60605 | MYL6_MOU | 17 kDa | 0.999136 | 6 | 83% | Q60605 | Myl6 | Myosin ligh | 23.07272 | 23.44481 | 22.62557 | 22.88893 | 22.75131 | 23.32471 | -0.40295 | -1.10892 | 0.303017 | 23.01801 | -1.35853 | -5.98495 | 0.217749 | 0.329908 | 0 | 0 |
| 570 | sp\|O35474 | EDIL3_MOU | 54 kDa | 0.998321 | 5 | 100% | O35474 | Edil3 | EGF-like rep | 21.17285 | 22.25191 | 22.40025 | 23.03696 | 22.31698 | 22.36921 | -0.58933 | -1.62668 | 0.448015 | 22.25803 | -1.35577 | -5.98809 | 0.219105 | 0.331591 | 0 | 0 |
| 952 | sp\|Q9CY66 | GAR1_MOU | 23 kDa | 0.997891 | 3 | 100% | Q9CY66 | Gar1 | H/ACA ribo | 19.33262 | 20.28371 | 19.13859 | 19.31163 | 19.57088 | 19.8274 | -0.46021 | -1.27234 | 0.351917 | 19.57747 | -1.34882 | -5.99667 | 0.220702 | 0.333555 | 0 | 0 |
| 106 | sp\|Q9D6F9 | TBB4A_MO | 50 kDa | 0.999582 | 15 | 13% | Q9D6F9 | Tubb4a | Tubulin bet | 21.58144 | 21.19593 | 21.18172 | 20.21757 | 20.72466 | 20.52791 | 0.51547 | -0.39534 | 1.42628 | 20.90487 | 1.349211 | -5.99608 | 0.220894 | 0.333555 | 0 | 0 |
| 775 | tr\|E9PUQ3 | E9PUQ3_M | 34 kDa | 0.995543 | 4 | 100% | E9PUQ3 | Nkapd1 | NKAP doma | 15.11682 | 17.07619 | 15.6913 | NA | 14.82018 | 15.12651 | -0.89191 | -2.50394 | 0.72012 | 15.5662 | -1.37145 | -5.85894 | 0.22179 | 0.334374 | 0 | 0 |
| 890 | sp\|Q9DCC4 | P5CR3_MO | 29 kDa | 0.998784 | 3 | 100% | Q9DCC4 | Pycr3 | Pyrroline-5 | 17.19474 | 15.69695 | 15.76588 | 18.78786 | 16.91062 | 19.25783 | -1.29047 | -3.57763 | 0.996696 | 17.26898 | -1.34648 | -5.99929 | 0.22193 | 0.334374 | 0 | 0 |
| 1050 | tr\|A0A075 | A0A075B6 | 13 kDa | 0.999313 | 2 | 100% | A0A075B6 | Igkv19-93 | Immunoglo | 14.4726 | 14.52237 | 15.42281 | 16.51854 | 12.29857 | NA | -1.45579 | -4.08993 | 1.178343 | 14.64698 | -1.36991 | -5.86077 | 0.222241 | 0.334472 | 0 | 0 |
| 1146 | sp\|Q62425 | NDUA4_MO | 9 kDa | 0.998505 | 2 | 100% | Q62425 | Ndufa4 | Cytochrom | 19.29007 | 20.39892 | 20.14955 | 20.29324 | 19.23964 | 19.56314 | -0.52535 | -1.45851 | 0.407815 | 19.82243 | -1.34351 | -6.00287 | 0.222841 | 0.335003 | 0 | 0 |
| 447 | sp\|P14131 | RS16_MOU | 16 kDa | 0.999093 | 6 | 100% | P14131 | Rps16 | 40S riboson | 24.59531 | 25.3173 | 24.80137 | 22.79347 | 24.28328 | 22.67738 | 0.963933 | -0.75061 | 2.678475 | 24.07802 | 1.341681 | -6.00506 | 0.223402 | 0.335474 | 0 | 0 |
| 664 | sp\|Q9D6Z1 | NOP56_MO | 64 kDa | 0.999188 | 4 | 100% | Q9D6Z1 | Nop56 | Nucleolar p | 20.08395 | 19.86843 | 20.11259 | 19.96428 | 20.09317 | 19.26276 | 0.398075 | -0.31275 | 1.108898 | 19.89753 | 1.332922 | -6.01577 | 0.225591 | 0.338386 | 0 | 0 |
| 249 | tr\|O35452 | O35452_M | 435 kDa | 0.998622 | 9 | 100% | O35452 | Tnxb | Tenascin X | 20.15781 | 19.76681 | 19.0807 | 19.38652 | 20.4381 | 18.59189 | 0.643793 | -0.50911 | 1.796694 | 19.57031 | 1.332617 | -6.01592 | 0.226204 | 0.338932 | 0 | 0 |
| 763 | sp\|Q3UL36 | ARGL1_MO | 33 kDa | 0.99725 | 4 | 100% | Q3UL36 | Arglu1 | Arginine an | 20.71566 | 20.34174 | 20.5412 | 19.3493 | 20.2237 | 20.31957 | 0.489988 | -0.3891 | 1.369076 | 20.24853 | 1.328763 | -6.02061 | 0.2272 | 0.340048 | 0 | 0 |
| 54 | sp\|Q6NZJ6 | IF4G1_MO | 176 kDa | 0.999849 | 19 | 89% | Q6NZJ6 | Eif4g1 | Eukaryotic | 24.22671 | 25.37446 | 23.71419 | 23.7381 | 24.68741 | 25.90715 | -0.79714 | -2.23128 | 0.637009 | 24.608 | -1.32645 | -6.02329 | 0.22813 | 0.340699 | 0 | 0 |
| 1283 | sp\|Q61011 | GBB3_MOU | 37 kDa | 0.995547 | 2 | 50% | Q61011 | Gnb3 | Guanine nu | NA | 15.81033 | 16.70831 | 17.15926 | 18.45037 | 16.99706 | 0.923794 | -0.77224 | 2.619823 | 17.02507 | 1.350121 | -5.88427 | 0.228137 | 0.340699 | 0 | 0 |
| 458 | sp\|P21661 | NEC2_MOU | 71 kDa | 0.998953 | 6 | 83% | P21661 | Pcsk2 | Neuroendo | 20.94724 | 20.62929 | 20.81091 | 20.11026 | 21.40032 | 21.03659 | 0.460778 | -0.36991 | 1.291465 | 20.82244 | 1.320311 | -6.03082 | 0.229553 | 0.342435 | 0 | 0 |
| 1237 | tr\|Q9Z1A1 | Q9Z1A1_M | 43 kDa | 0.997261 | 2 | 100% | Q9Z1A1 | Tfg | TFG protein | 18.08519 | 17.72852 | 17.96482 | 17.15726 | 18.02598 | 17.97065 | 0.40652 | -0.32981 | 1.142852 | 17.82207 | 1.31404 | -6.03828 | 0.231531 | 0.345007 | 0 | 0 |
| 1099 | sp\|P19157 | GSTP1_MO | 24 kDa | 0.998964 | 2 | 100% | P19157 | Gstp1 | Glutathion | 20.60479 | 20.60859 | 20.79604 | 20.62818 | 20.1613 | 22.90224 | -0.85896 | -2.41813 | 0.700207 | 20.95019 | -1.31471 | -6.03724 | 0.231832 | 0.345076 | 0 | 0 |
| 922 | sp\|Q99MJ9 | DDX50_MO | 82 kDa | 0.998358 | 3 | 100% | Q99MJ9 | Ddx50 | ATP-depen | 22.13603 | 22.16664 | 23.14478 | 21.84751 | 22.00701 | 21.77412 | 0.499853 | -0.40967 | 1.409377 | 22.17935 | 1.310186 | -6.04271 | 0.233075 | 0.346032 | 0 | 0 |
| 923 | sp\|P63276 | RS17_MOU | 16 kDa | 0.998349 | 3 | 100% | P63276 | Rps17 | 40S riboson | 22.13603 | 22.16664 | 23.14478 | 21.84751 | 22.00701 | 21.77412 | 0.499853 | -0.40967 | 1.409377 | 22.17935 | 1.310186 | -6.04271 | 0.233075 | 0.346032 | 0 | 0 |
| 839 | sp\|Q60668 | HNRPD_MO | 38 kDa | 0.999248 | 3 | 67% | Q60668 | Hnrnpd | Heterogene | 19.12274 | 19.24007 | 18.46981 | 19.39219 | 19.46803 | 19.84987 | -0.47385 | -1.33638 | 0.388686 | 19.25712 | -1.30966 | -6.04333 | 0.23324 | 0.346032 | 0 | 0 |
| 1204 | sp\|Q99KL7 | RAB28_MO | 25 kDa | 0.997906 | 2 | 100% | Q99KL7 | Rab28 | Ras-related | 16.11008 | 18.09839 | 16.67935 | 16.22784 | 15.26952 | 16.19803 | -0.82177 | -2.32294 | 0.679402 | 16.43053 | -1.30638 | -6.04711 | 0.234492 | 0.34751 | 0 | 0 |
| 728 | sp\|Q8BJ71 | NUP93_MO | 93 kDa | 0.998481 | 4 | 100% | Q8BJ71 | Nup93 | Nuclear po | 19.45614 | 20.75315 | 19.3343 | 19.10635 | 19.85234 | 20.75413 | -0.65695 | -1.86034 | 0.546433 | 19.87607 | -1.30281 | -6.05133 | 0.235642 | 0.348833 | 0 | 0 |
| 1085 | sp\|Q8K021 | SCAM1_MO | 38 kDa | 0.999085 | 2 | 100% | Q8K021 | Scamp1 | Secretory c | 19.68145 | 19.55658 | 18.13059 | 19.58311 | 19.35914 | 19.75749 | -0.57533 | -1.62998 | 0.479309 | 19.34473 | -1.30186 | -6.05245 | 0.235946 | 0.348902 | 0 | 0 |
| 736 | sp\|P54869 | HMCS2_MO | 57 kDa | 0.998356 | 4 | 100% | P54869 | Hmgcs2 | Hydroxyme | 16.87567 | 15.86102 | 16.68381 | 13.96195 | 17.54384 | 17.75657 | 1.174594 | -0.98221 | 3.331401 | 16.44714 | 1.299652 | -6.05505 | 0.236661 | 0.349577 | 0 | 0 |
| 703 | sp\|Q9CPR4 | RL17_MOU | 21 kDa | 0.998861 | 4 | 100% | Q9CPR4 | Rpl17 | 60S riboson | 23.9601 | 23.76732 | 23.98207 | 22.82783 | 23.66429 | 23.66947 | 0.447283 | -0.37497 | 1.269532 | 23.64518 | 1.294795 | -6.06103 | 0.237736 | 0.350783 | 0 | 0 |
| 1217 | sp\|Q64511 | TOP2B_MO | 182 kDa | 0.997738 | 2 | 100% | Q64511 | Top2b | DNA topois | 17.75285 | 17.04339 | 18.51347 | 17.761 | 18.60066 | 18.37083 | 0.563923 | -0.47847 | 1.606314 | 18.00703 | 1.291041 | -6.06518 | 0.23946 | 0.352944 | 0 | 0 |

| | | | | | | | | | | | | | | | | | | | | | | | | | |
|---|---|---|---|---|---|---|---|---|---|---|---|---|---|---|---|---|---|---|---|---|---|---|---|---|---|
| 851 | sp\|P01193 | COLI_MOU | 27 kDa | 0.999134 | 3 | 100% | P01193 | Pomc | Pro-opiom | 20.26525 | 20.81112 | 20.26936 | 20.07104 | 19.80083 | 20.70913 | -0.41861 | -1.19569 | 0.358466 | 20.32112 | -1.28218 | -6.07583 | 0.241865 | 0.355715 | 0 | 0 |
| 852 | sp\|Q8R3Q0 | SARAF_MO | 36 kDa | 0.99911 | 3 | 100% | Q8R3Q0 | Saraf | Store-opera | 20.26525 | 20.81112 | 20.26936 | 20.07104 | 19.80083 | 20.70913 | -0.41861 | -1.19569 | 0.358466 | 20.32112 | -1.28218 | -6.07583 | 0.241865 | 0.355715 | 0 | 0 |
| 60 | sp\|Q3UPL0 | SC31A_MO | 134 kDa | 0.999301 | 19 | 100% | Q3UPL0 | Sec31a | Protein tra | 24.52068 | 24.10077 | 23.12058 | 24.35929 | 24.19547 | 25.25077 | -0.6247 | -1.79127 | 0.541869 | 24.25793 | -1.27794 | -6.0805 | 0.243775 | 0.358135 | 0 | 0 |
| 772 | sp\|Q61554 | FBN1_MOU | 312 kDa | 0.996719 | 4 | 100% | Q61554 | Fbn1 | Fibrillin-1 | 15.93234 | 15.79776 | 15.85361 | 18.33627 | NA | 16.53124 | -0.99545 | -2.89501 | 0.904119 | 16.49024 | -1.29896 | -5.94421 | 0.244045 | 0.358143 | 0 | 0 |
| 53.1 | sp\|P60710 | ACTB_MOU | 42 kDa | 0.999884 | 19 | 32% | P60710 | Actb | Actin, cyto | 27.26109 | 26.84488 | 27.26374 | 27.07091 | 27.37189 | 27.01326 | 0.322558 | -0.28051 | 0.925628 | 27.13763 | 1.273038 | -6.08651 | 0.244903 | 0.358627 | 0 | 0 |
| 53.2 | sp\|P63260 | ACTG_MOU | 42 kDa | 0.999884 | 19 | 32% | P63260 | Actg1 | Actin, cyto | 27.26109 | 26.84488 | 27.26374 | 27.07091 | 27.37189 | 27.01326 | 0.322558 | -0.28051 | 0.925628 | 27.13763 | 1.273038 | -6.08651 | 0.244903 | 0.358627 | 0 | 0 |
| 1035 | sp\|Q4VA53 | PDS5B_MO | 164 kDa | 0.999478 | 2 | 100% | Q4VA53 | Pds5b | Sister chro | 21.34795 | 22.48577 | 21.53638 | 21.21233 | 21.59444 | 22.27492 | -0.49808 | -1.4396 | 0.443435 | 21.74197 | -1.26248 | -6.09848 | 0.248955 | 0.364167 | 0 | 0 |
| 1138 | sp\|P42125 | ECI1_MOUS | 32 kDa | 0.998555 | 2 | 100% | P42125 | Eci1 | Enoyl-CoA | 14.26245 | 15.81581 | 15.69376 | 14.09489 | 14.49537 | 19.20698 | -1.55536 | -4.50138 | 1.390656 | 15.59488 | -1.25993 | -6.10143 | 0.249817 | 0.364725 | 0 | 0 |
| 501 | sp\|Q9CZ13 | QCR1_MOU | 53 kDa | 0.999537 | 5 | 100% | Q9CZ13 | Uqcrc1 | Cytochrom | 22.12501 | 23.54609 | 22.59612 | 22.46173 | 22.52523 | 22.64883 | -0.4701 | -1.36011 | 0.419912 | 22.6505 | -1.25918 | -6.10242 | 0.249874 | 0.364725 | 0 | 0 |
| 716 | sp\|P62900 | RL31_MOU | 14 kDa | 0.998717 | 4 | 100% | P62900 | Rpl31 | 60S riboso | 23.02013 | 22.98067 | 23.86283 | 23.02079 | 23.06881 | 22.69608 | 0.418077 | -0.37444 | 1.210591 | 23.10822 | 1.255595 | -6.10674 | 0.250791 | 0.36567 | 0 | 0 |
| 512 | sp\|A2AR02 | PPIG_MOU | 88 kDa | 0.999448 | 5 | 100% | A2AR02 | Ppig | Peptidyl-pr | 23.55085 | 24.8093 | 21.79787 | 22.3406 | 23.17949 | 24.19385 | -0.93851 | -2.7215 | 0.844478 | 23.31199 | -1.25615 | -6.1058 | 0.251102 | 0.36573 | 0 | 0 |
| 108 | sp\|Q9JIX8 | ACINU_MO | 151 kDa | 0.999387 | 15 | 100% | Q9JIX8 | Acin1 | Apoptotic | 23.83579 | 23.0013 | 22.59459 | 22.90041 | 23.74044 | 22.78002 | 0.496363 | -0.44931 | 1.442033 | 23.14209 | 1.252595 | -6.1099 | 0.252315 | 0.366731 | 0 | 0 |
| 183 | sp\|P60843 | IF4A1_MOU | 46 kDa | 0.999862 | 10 | 40% | P60843 | Eif4a1 | Eukaryotic | 21.07199 | 19.89601 | 19.96077 | 20.14829 | 20.41824 | 20.07783 | 0.442961 | -0.40009 | 1.28601 | 20.26219 | 1.251492 | -6.11139 | 0.252331 | 0.366731 | 0 | 0 |
| 629 | sp\|P17182 | ENOA_MOU | 47 kDa | 0.999435 | 4 | 50% | P17182 | Eno1 | Alpha-enol | 20.38999 | 20.77463 | 19.48551 | 20.29795 | 20.581 | 20.80718 | -0.47442 | -1.37898 | 0.430142 | 20.38938 | -1.25031 | -6.11265 | 0.252899 | 0.367164 | 0 | 0 |
| 361 | sp\|Q6IFZ6 | K2C1B_MO | 61 kDa | 0.999128 | 7 | 43% | Q6IFZ6 | Krt77 | Keratin, typ | 20.03582 | 19.79831 | 20.56672 | 21.47475 | 19.98644 | 21.14134 | -0.60848 | -1.77024 | 0.553288 | 20.50056 | -1.2499 | -6.113 | 0.253238 | 0.367262 | 0 | 0 |
| 994 | sp\|Q99L88 | SNTB1_MO | 58 kDa | 0.996484 | 3 | 67% | Q99L88 | Sntb1 | Beta-1-synt | 19.19151 | 17.90519 | 18.17066 | 18.85729 | 19.5099 | 18.42548 | 0.561371 | -0.512 | 1.634739 | 18.67667 | 1.248109 | -6.11506 | 0.253854 | 0.367762 | 0 | 0 |
| 548 | sp\|Q922D8 | C1TC_MOU | 101 kDa | 0.999011 | 5 | 100% | Q922D8 | Mthfd1 | C-1-tetrahy | 20.59735 | 21.76774 | 21.13135 | 21.43041 | 21.11702 | 20.91412 | -0.42218 | -1.23018 | 0.385817 | 21.15966 | -1.24368 | -6.12045 | 0.254891 | 0.368871 | 0 | 0 |
| 802 | tr\|A0A075 | A0A075B5Y | 13 kDa | 0.999506 | 3 | 33% | A0A075B5Y | Ighv1-80 | Immunoglo | 19.71265 | 19.61132 | 18.62336 | 20.98819 | 19.85256 | 19.55977 | -0.65691 | -1.92068 | 0.606873 | 19.72464 | -1.24046 | -6.12384 | 0.256495 | 0.370797 | 0 | 0 |
| 906 | sp\|Q69ZU6 | THS7A_MO | 184 kDa | 0.998597 | 3 | 100% | Q69ZU6 | Thsd7a | Thrombosp | 20.64849 | 20.98868 | 20.97098 | 24.70158 | 20.98408 | 20.72049 | -1.26907 | -3.71729 | 1.179148 | 21.50238 | -1.23705 | -6.12774 | 0.257681 | 0.372114 | 0 | 0 |
| 1265 | sp\|Q8C092 | TAF5_MOU | 87 kDa | 0.996358 | 2 | 100% | Q8C092 | Taf5 | Transcripti | 22.4533 | 22.08904 | 22.9978 | 22.66527 | 22.49715 | 22.00323 | 0.396904 | -0.37131 | 1.16512 | 22.45097 | 1.229712 | -6.13644 | 0.259753 | 0.374666 | 0 | 0 |
| 118 | sp\|P20029 | BIP_MOUSI | 72 kDa | 0.999596 | 14 | 86% | P20029 | Hspa5 | Endoplasm | 24.07169 | 23.69779 | 24.68037 | 24.43839 | 24.2952 | 23.64415 | 0.422311 | -0.39557 | 1.240194 | 24.13793 | 1.229032 | -6.13721 | 0.260001 | 0.374666 | 0 | 0 |
| 449 | tr\|D3Z2H9 | D3Z2H9_M | 29 kDa | 0.999056 | 6 | 67% | D3Z2H9 | Tpm3-rs7 | Tropomyos | 21.91828 | 21.58947 | 21.8854 | 22.19357 | 23.30543 | 21.6336 | 0.564156 | -0.53714 | 1.665453 | 22.08763 | 1.222489 | -6.14433 | 0.262792 | 0.377613 | 0 | 0 |
| 767 | sp\|Q61292 | LAMB2_MC | 197 kDa | 0.997106 | 4 | 100% | Q61292 | Lamb2 | Laminin su | 19.84692 | 20.85124 | 18.91676 | 19.40409 | 20.62722 | 21.61181 | -0.82542 | -2.43689 | 0.78606 | 20.20967 | -1.22236 | -6.14448 | 0.262838 | 0.377613 | 0 | 0 |
| 1214 | sp\|Q8CG76 | ARK72_MO | 41 kDa | 0.997748 | 2 | 100% | Q8CG76 | Akr7a2 | Aflatoxin B | 18.67074 | 14.65707 | 19.2279 | 18.10806 | 18.62914 | 19.53054 | 1.410706 | -1.34372 | 4.16513 | 18.13724 | 1.222239 | -6.14461 | 0.262881 | 0.377613 | 0 | 0 |
| 715 | sp\|Q8CJ67 | STAU2_MO | 63 kDa | 0.998727 | 4 | 100% | Q8CJ67 | Stau2 | Double-stra | 18.31204 | 17.39822 | 18.52715 | 17.42763 | 18.25092 | 18.72317 | 0.513697 | -0.49246 | 1.519854 | 18.10652 | 1.218406 | -6.14896 | 0.264242 | 0.379166 | 0 | 0 |
| 367 | sp\|P61264 | STX1B_MO | 33 kDa | 0.999014 | 7 | 100% | P61264 | Stx1b | Syntaxin-1I | 23.69454 | 23.33416 | 22.91814 | 22.26714 | 23.53783 | 23.1099 | 0.47977 | -0.46332 | 1.422856 | 23.14362 | 1.214038 | -6.1539 | 0.2658 | 0.380308 | 0 | 0 |
| 95 | sp\|E9Q7E2 | ARID2_MO | 196 kDa | 0.999408 | 16 | 100% | E9Q7E2 | Arid2 | AT-rich inte | 22.45908 | 22.99555 | 21.34756 | 20.54714 | 22.72861 | 20.01082 | 0.993916 | -0.95993 | 2.947763 | 21.68146 | 1.213974 | -6.15397 | 0.265822 | 0.380308 | 0 | 0 |
| 212 | sp\|Q9WU7 | PDC6I_MO | 96 kDa | 0.999134 | 10 | 100% | Q9WU78 | Pdcd6ip | Programme | 21.80837 | 22.37338 | 22.09878 | 22.73573 | 21.50208 | 21.65206 | -0.45065 | -1.33611 | 0.434819 | 22.0284 | -1.21327 | -6.1549 | 0.265879 | 0.380308 | 0 | 0 |
| 1220 | sp\|P61022 | CHP1_MOU | 22 kDa | 0.997671 | 2 | 100% | P61022 | Chp1 | Calcineurin | 16.26696 | 17.42891 | 16.9435 | 16.1818 | 16.31512 | 17.54921 | -0.54478 | -1.61759 | 0.528036 | 16.78092 | -1.21184 | -6.15638 | 0.266586 | 0.380917 | 0 | 0 |
| 691 | sp\|P51150 | RAB7A_MO | 23 kDa | 0.998994 | 4 | 100% | P51150 | Rab7a | Ras-related | 21.92977 | 22.84273 | 23.07417 | 21.85513 | 23.66143 | 22.09604 | 0.623826 | -0.60879 | 1.856442 | 22.57654 | 1.207775 | -6.16097 | 0.268047 | 0.382601 | 0 | 0 |
| 801 | tr\|E9Q6E5 | E9Q6E5_M | 57 kDa | 0.99951 | 3 | 100% | E9Q6E5 | Srsf11 | Serine/argi | 20.82789 | 20.43009 | 21.63236 | 20.80723 | 20.65197 | 20.61885 | 0.418684 | -0.41066 | 1.248028 | 20.82807 | 1.201639 | -6.16821 | 0.269788 | 0.384682 | 0 | 0 |
| 396 | sp\|P05064 | ALDOA_MC | 39 kDa | 0.999727 | 6 | 100% | P05064 | Aldoa | Fructose-bi | 22.58627 | 23.47733 | 23.39034 | 23.49036 | 22.6982 | 22.93938 | -0.41075 | -1.22639 | 0.40489 | 23.09698 | -1.19867 | -6.17153 | 0.270866 | 0.385813 | 0 | 0 |
| 999 | sp\|Q3U0S6 | RAIN_MOU | 104 kDa | 0.99588 | 3 | 100% | Q3U0S6 | Rasip1 | Ras-interac | 19.04704 | 19.79216 | 18.94951 | 19.88899 | 19.72588 | 19.26755 | -0.40876 | -1.22583 | 0.408315 | 19.44519 | -1.19077 | -6.18038 | 0.273757 | 0.389522 | 0 | 0 |
| 1118 | sp\|Q9CZX7 | PP4P2_MO | 28 kDa | 0.998761 | 2 | 100% | Q9CZX7 | Pip4p2 | Type 2 pho | 18.33188 | 19.50028 | 18.0082 | 18.10931 | 18.87474 | 19.21301 | -0.53593 | -1.61676 | 0.544901 | 18.6729 | -1.18332 | -6.18834 | 0.276978 | 0.393691 | 0 | 0 |
| 30 | sp\|Q6DIC0 | SMCA2_MC | 180 kDa | 0.999766 | 25 | 68% | Q6DIC0 | Smarca2 | Probable gl | 24.44188 | 24.3204 | 23.4863 | 23.00031 | 24.71935 | 23.62814 | 0.566228 | -0.57753 | 1.709982 | 23.93273 | 1.181435 | -6.19043 | 0.277675 | 0.39427 | 0 | 0 |
| 232 | sp\|O08599 | STXB1_MO | 68 kDa | 0.999644 | 9 | 100% | O08599 | Stxbp1 | Syntaxin-bi | 23.33014 | 24.06757 | 23.7112 | 23.70792 | 23.53412 | 23.76879 | -0.32294 | -0.97536 | 0.329477 | 23.68662 | -1.17814 | -6.19444 | 0.278422 | 0.394916 | 0 | 0 |
| 1124 | sp\|Q8BRT1 | CLAP2_MO | 141 kDa | 0.998677 | 2 | 100% | Q8BRT1 | Clasp2 | CLIP-associ | 20.42563 | 20.1827 | 20.86269 | 20.08948 | 20.35398 | 20.37979 | 0.330107 | -0.3428 | 1.003011 | 20.38238 | 1.167624 | -6.20607 | 0.282364 | 0.40009 | 0 | 0 |
| 753 | sp\|P97789 | XRN1_MOU | 194 kDa | 0.997634 | 4 | 100% | P97789 | Xrn1 | 5'-3' exorib | 20.61666 | 19.6441 | 20.93442 | 20.87949 | 20.85339 | 20.54193 | 0.446318 | -0.46527 | 1.357907 | 20.57833 | 1.167248 | -6.20626 | 0.282802 | 0.400292 | 0 | 0 |
| 211.1 | sp\|Q91VC3 | IF4A3_MOU | 47 kDa | 0.99918 | 10 | 80% | Q91VC3 | Eif4a3 | Eukaryotic | 23.05602 | 22.71626 | 22.55292 | 24.13776 | 23.62744 | 24.12589 | -0.58118 | -1.7747 | 0.612336 | 23.36938 | -1.16207 | -6.21181 | 0.284937 | 0.402054 | 0 | 0 |
| 211.2 | tr\|A0A0N4 | A0A0N4SVI | 47 kDa | 0.99918 | 10 | 80% | A0A0N4SVI | Gm5580 | Predicted p | 23.05602 | 22.71626 | 22.55292 | 24.13776 | 23.62744 | 24.12589 | -0.58118 | -1.7747 | 0.612336 | 23.36938 | -1.16207 | -6.21181 | 0.284937 | 0.402054 | 0 | 0 |
| 211.3 | tr\|E9PV04 | E9PV04_M | 47 kDa | 0.99918 | 10 | 80% | E9PV04 | Gm8994 | Predicted g | 23.05602 | 22.71626 | 22.55292 | 24.13776 | 23.62744 | 24.12589 | -0.58118 | -1.7747 | 0.612336 | 23.36938 | -1.16207 | -6.21181 | 0.284937 | 0.402054 | 0 | 0 |
| 1159 | sp\|O88685 | PRS6A_MO | 50 kDa | 0.998334 | 2 | 100% | O88685 | Psmc3 | 26S protea | 18.05916 | 18.96047 | 18.32641 | 18.41132 | 17.6215 | 17.92422 | -0.42965 | -1.31643 | 0.457131 | 18.21718 | -1.15502 | -6.21969 | 0.287433 | 0.405155 | 0 | 0 |
| 576 | sp\|Q3UIL6 | PKHA7_MC | 127 kDa | 0.99801 | 5 | 100% | Q3UIL6 | Plekha7 | Pleckstrin h | 20.29183 | 19.15625 | 19.08372 | 20.08616 | 19.99944 | 18.16854 | 0.654681 | -0.699 | 2.008361 | 19.46432 | 1.154157 | -6.22049 | 0.287951 | 0.405464 | 0 | 0 |
| 221 | sp\|Q9ESX5 | DKC1_MOU | 57 kDa | 0.9999 | 9 | 100% | Q9ESX5 | Dkc1 | H/ACA ribo | 22.24927 | 22.80967 | 22.01281 | 21.40163 | 22.69208 | 20.97133 | 0.59051 | -0.64337 | 1.824388 | 22.0228 | 1.142106 | -6.23362 | 0.292591 | 0.41157 | 0 | 0 |
| 996 | sp\|Q61699 | HS105_MO | 96 kDa | 0.996089 | 3 | 67% | Q61699 | Hsph1 | Heat shock | 22.72279 | 22.25554 | 23.556 | 22.73277 | 22.00289 | 21.60967 | 0.561236 | -0.61544 | 1.737916 | 22.47994 | 1.138249 | -6.23781 | 0.294089 | 0.413248 | 0 | 0 |
| 93 | sp\|Q8BH59 | CMC1_MOI | 75 kDa | 0.999516 | 16 | 88% | Q8BH59 | Slc25a12 | Calcium-bi | 24.71949 | 24.52611 | 24.85689 | 23.9849 | 24.46802 | 24.52528 | 0.336037 | -0.36807 | 1.040146 | 24.51345 | 1.135921 | -6.24071 | 0.294532 | 0.413442 | 0 | 0 |
| 696 | sp\|P18760 | COF1_MOU | 19 kDa | 0.998935 | 4 | 75% | P18760 | Cfl1 | Cofilin-1 | 22.48757 | 21.98246 | 21.6139 | 22.46261 | 21.58576 | 22.44953 | -0.40246 | -1.25169 | 0.446778 | 22.09697 | -1.1295 | -6.24744 | 0.297287 | 0.416878 | 0 | 0 |
| 1067 | sp\|Q9QYJ0 | DNJA2_MO | 46 kDa | 0.999232 | 2 | 100% | Q9QYJ0 | Dnaja2 | DnaJ homo | 19.01908 | 18.39065 | 17.81519 | 20.52241 | 19.60757 | 19.81672 | -0.76264 | -2.38218 | 0.856895 | 19.19527 | -1.12378 | -6.25342 | 0.299765 | 0.419307 | 0 | 0 |
| 1262 | sp\|Q9QZ06 | TOLIP_MOU | 30 kDa | 0.996417 | 2 | 100% | Q9QZ06 | Tollip | Toll-interac | 18.56015 | 17.36388 | 16.65478 | 17.4022 | 15.91283 | 18.96257 | -0.86697 | -2.71008 | 0.976144 | 17.47607 | -1.12254 | -6.25475 | 0.300256 | 0.419307 | 0 | 0 |
| 1263 | sp\|Q8C050 | KS6A5_MO | 97 kDa | 0.996414 | 2 | 100% | Q8C050 | Rps6ka5 | Ribosomal | 18.56015 | 17.36388 | 16.65478 | 17.4022 | 15.91283 | 18.96257 | -0.86697 | -2.71008 | 0.976144 | 17.47607 | -1.12254 | -6.25475 | 0.300256 | 0.419307 | 0 | 0 |
| 1264 | sp\|Q9JME5 | AP3B2_MO | 119 kDa | 0.99637 | 2 | 100% | Q9JME5 | Ap3b2 | AP-3 comp | 18.56015 | 17.36388 | 16.65478 | 17.4022 | 15.91283 | 18.96257 | -0.86697 | -2.71008 | 0.976144 | 17.47607 | -1.12254 | -6.25475 | 0.300256 | 0.419307 | 0 | 0 |
| 425 | sp\|Q60673 | PTPRN_MO | 106 kDa | 0.999409 | 6 | 83% | Q60673 | Ptprn | Receptor-ty | 21.69706 | 22.63807 | 22.76399 | 22.60891 | 21.73337 | 22.21977 | -0.42411 | -1.3273 | 0.479078 | 22.27686 | -1.11943 | -6.25824 | 0.301308 | 0.420344 | 0 | 0 |
| 40 | sp\|P17156 | HSP72_MO | 70 kDa | 0.999892 | 22 | 55% | P17156 | Hspa2 | Heat shock | 25.21283 | 24.77557 | 24.97384 | 24.75298 | 25.17259 | 24.9733 | 0.285805 | -0.3294 | 0.90101 | 24.97685 | 1.105734 | -6.27309 | 0.306517 | 0.427171 | 0 | 0 |
| 975 | sp\|O08664 | BCL7C_MO | 23 kDa | 0.997539 | 3 | 67% | O08664 | Bcl7c | B-cell CLL/l | 18.52368 | 19.65756 | 18.48339 | 20.09007 | 18.47839 | 17.74715 | -0.66977 | -2.11973 | 0.780188 | 18.83004 | -1.10236 | -6.27628 | 0.308336 | 0.429264 | 0 | 0 |
| 411 | sp\|Q9Z2X1 | HNRPF_MC | 46 kDa | 0.999584 | 6 | 83% | Q9Z2X1 | Hnrnpf | Heterogene | 21.42801 | 21.04381 | 20.36141 | 20.96755 | 21.14544 | 22.90569 | -0.66073 | -2.09308 | 0.771612 | 21.30865 | -1.10085 | -6.27787 | 0.308945 | 0.42953 | 0 | 0 |
| 91 | sp\|P56480 | ATPB_MOU | 56 kDa | 0.999697 | 16 | 100% | P56480 | Atp5f1b | ATP syntha | 25.16068 | 25.45215 | 25.34254 | 25.33352 | 25.15319 | 25.75801 | -0.29575 | -0.93615 | 0.344647 | 25.36668 | -1.09921 | -6.28001 | 0.30916 | 0.42953 | 0 | 0 |
| 856 | sp\|P12367 | KAP2_MOU | 45 kDa | 0.999076 | 3 | 67% | P12367 | Prkar2a | cAMP-depe | 20.75131 | 21.2365 | 21.2305 | 21.37016 | 21.18599 | 21.46858 | -0.30248 | -0.96358 | 0.358626 | 21.20717 | -1.08899 | -6.29079 | 0.313336 | 0.434564 | 0 | 0 |
| 625 | sp\|P01897 | HA1L_MOU | 41 kDa | 0.999491 | 4 | 100% | P01897 | H2-L | H-2 class I h | 21.84149 | 21.48162 | 22.86674 | 22.06264 | 21.88092 | 21.80673 | 0.41272 | -0.49091 | 1.316352 | 21.99002 | 1.088827 | -6.29072 | 0.313672 | 0.434564 | 0 | 0 |
| 606 | sp\|Q99MR | MCCA_MOI | 79 kDa | 0.999677 | 4 | 100% | Q99MR8 | Mccc1 | Methylcrot | 20.12466 | 20.95707 | 20.72556 | 20.63492 | 20.0551 | 20.36848 | -0.35171 | -1.12112 | 0.417703 | 20.47763 | -1.08799 | -6.29183 | 0.313745 | 0.434564 | 0 | 0 |
| 595 | sp\|P84099 | RL19_MOU | 23 kDa | 0.99974 | 4 | 100% | P84099 | Rpl19 | 60S riboso | 24.42087 | 24.4852 | 26.29186 | 24.81166 | 24.85375 | 24.6224 | 0.549073 | -0.65824 | 1.756381 | 24.91429 | 1.085331 | -6.29423 | 0.315288 | 0.436255 | 0 | 0 |
| 302 | sp\|P70398 | USP9X_MC | 291 kDa | 0.997909 | 8 | 100% | P70398 | Usp9x | Probable u | 21.59574 | 21.49756 | 21.545 | 23.19385 | 21.72931 | 21.73635 | -0.51924 | -1.66188 | 0.623403 | 21.88297 | -1.08445 | -6.29516 | 0.315653 | 0.436316 | 0 | 0 |
| 935 | sp\|Q8K212 | PACS1_MO | 105 kDa | 0.998128 | 3 | 67% | Q8K212 | Pacs1 | Phosphofu | 17.17446 | 16.60528 | 17.72118 | 18.03294 | 17.3244 | 20.77297 | -1.06371 | -3.41231 | 1.284883 | 17.93854 | -1.08085 | -6.29892 | 0.317137 | 0.436854 | 0 | 0 |
| 1079 | sp\|Q8BVE3 | VATH_MOU | 56 kDa | 0.999136 | 2 | 100% | Q8BVE3 | Atp6v1h | V-type prot | 19.61019 | 20.00108 | 20.18677 | 20.03633 | 20.07403 | 21.13036 | -0.43226 | -1.38761 | 0.523094 | 20.17313 | -1.07977 | -6.30005 | 0.317587 | 0.436854 | 0 | 0 |
| 795 | sp\|Q9JLM8 | DCLK1_MO | 84 kDa | 0.999593 | 3 | 100% | Q9JLM8 | Dclk1 | Serine/thre | 21.98741 | 21.39565 | 21.15474 | 21.41427 | 22.03333 | 21.29436 | 0.357063 | -0.43091 | 1.145036 | 21.54663 | 1.078536 | -6.30174 | 0.317654 | 0.436854 | 0 | 0 |
| 796 | sp\|P70296 | PEBP1_MO | 21 kDa | 0.99957 | 3 | 100% | P70296 | Pebp1 | Phosphatid | 21.98741 | 21.39565 | 21.15474 | 21.41427 | 22.03333 | 21.29436 | 0.357063 | -0.43091 | 1.145036 | 21.54663 | 1.078536 | -6.30174 | 0.317654 | 0.436854 | 0 | 0 |
| 797 | sp\|P35282 | RAB21_MO | 24 kDa | 0.999555 | 3 | 67% | P35282 | Rab21 | Ras-related | 21.98741 | 21.39565 | 21.15474 | 21.41427 | 22.03333 | 21.29436 | 0.357063 | -0.43091 | 1.145036 | 21.54663 | 1.078536 | -6.30174 | 0.317654 | 0.436854 | 0 | 0 |
| 22 | sp\|Q99PL5 | RRBP1_MO | 173 kDa | 0.999797 | 27 | 100% | Q99PL5 | Rrbp1 | Ribosome-l | 25.41022 | 25.37928 | 25.6512 | 24.91899 | 25.72088 | 25.53594 | 0.316031 | -0.38255 | 1.014609 | 25.43609 | 1.076752 | -6.3036 | 0.318396 | 0.437431 | 0 | 0 |
| 29 | sp\|Q6A0A9 | F120A_MO | 122 kDa | 0.999791 | 25 | 92% | Q6A0A9 | FAM120A | Constitutiv | 24.79018 | 24.02983 | 24.82944 | 23.89092 | 24.68235 | 25.12409 | 0.419041 | -0.51009 | 1.348176 | 24.5578 | 1.076289 | -6.30368 | 0.319032 | 0.437862 | 0 | 0 |
| 706 | sp\|Q76MZ3 | 2AAA_MOU | 65 kDa | 0.998838 | 4 | 100% | Q76MZ3 | Ppp2r1a | Serine/thre | 21.04498 | 20.67588 | 21.5613 | 20.78137 | 21.19417 | 21.33114 | 0.337356 | -0.41124 | 1.085948 | 21.09814 | 1.072616 | -6.30791 | 0.320121 | 0.438912 | 0 | 0 |
| 1108 | sp\|P54116 | STOM_MOU | 31 kDa | 0.998855 | 2 | 100% | P54116 | Stom | Erythrocyte | 17.72699 | 17.93445 | 18.19323 | NA | 16.65377 | 15.12948 | 0.992699 | -1.2862 | 3.271602 | 17.12758 | 1.079749 | -6.18572 | 0.323827 | 0.443545 | 0 | 0 |
| 1155 | sp\|Q9EPK6 | SIL1_MOUS | 52 kDa | 0.998388 | 2 | 100% | Q9EPK6 | Sil1 | Nucleotide | 18.82831 | 20.58365 | 19.21105 | 19.52668 | 20.09733 | 19.51783 | -0.49715 | -1.61323 | 0.618926 | 19.62748 | -1.06303 | -6.31744 | 0.324586 | 0.444136 | 0 | 0 |
| 878 | sp\|P05202 | AATM_MOU | 47 kDa | 0.998944 | 3 | 100% | P05202 | Got2 | Aspartate a | 19.09702 | 19.5882 | 19.82708 | 20.85778 | 19.37733 | 19.33916 | -0.49457 | -1.6105 | 0.621365 | 19.68109 | -1.05764 | -6.323 | 0.326866 | 0.446316 | 0 | 0 |
| 1083 | sp\|Q9D666 | SUN1_MOU | 102 kDa | 0.999095 | 2 | 100% | Q9D666 | Sun1 | SUN domai | 18.52893 | 17.95948 | 17.8518 | 18.07803 | 18.55362 | 17.93081 | 0.322011 | -0.40416 | 1.048177 | 18.15044 | 1.055443 | -6.32567 | 0.327364 | 0.446316 | 0 | 0 |
| 1084 | sp\|Q8BH58 | TIPRL_MOU | 31 kDa | 0.999091 | 2 | 100% | Q8BH58 | Tiprl | TIP41-like p | 18.52893 | 17.95948 | 17.8518 | 18.07803 | 18.55362 | 17.93081 | 0.322011 | -0.40416 | 1.048177 | 18.15044 | 1.055443 | -6.32567 | 0.327364 | 0.446316 | 0 | 0 |
| 97 | sp\|P68368 | TBA4A_MO | 50 kDa | 0.999782 | 15 | 33% | P68368 | Tuba4a | Tubulin alp | 24.50154 | 24.53722 | 24.01647 | 24.29535 | 23.90417 | 24.50605 | -0.30548 | -0.99498 | 0.384026 | 24.29347 | -1.0545 | -6.32665 | 0.327768 | 0.446316 | 0 | 0 |
| 1031 | sp\|P62315 | SMD1_MOI | 13 kDa | 0.999534 | 2 | 100% | P62315 | Snrpd1 | Small nucle | 22.91578 | 20.40756 | 23.37379 | 23.58675 | 22.82967 | 22.65532 | 0.823206 | -1.03823 | 2.684642 | 22.62814 | 1.055385 | -6.32533 | 0.327825 | 0.446316 | 0 | 0 |
| 46 | sp\|P39054 | DYN2_MOU | 98 kDa | 0.999773 | 20 | 45% | P39054 | Dnm2 | Dynamin-2 | 24.10148 | 23.12313 | 22.72644 | 25.41301 | 24.45549 | 25.04125 | -0.76466 | -2.49598 | 0.96666 | 24.14347 | -1.054 | -6.32675 | 0.328413 | 0.446578 | 0 | 0 |
| 76 | sp\|P68369 | TBA1A_MO | 50 kDa | 0.999608 | 17 | 41% | P68369 | Tuba1a | Tubulin alp | 26.08717 | 25.4928 | 25.21534 | 25.23483 | 25.58089 | 25.13188 | 0.341299 | -0.43062 | 1.113215 | 25.45715 | 1.052365 | -6.32884 | 0.328676 | 0.446578 | 0 | 0 |
| 823 | sp\|P62996 | TRA2B_MO | 34 kDa | 0.999382 | 3 | 100% | P62996 | Tra2b | Transforme | 22.90713 | 22.94679 | 24.26385 | 23.21417 | 22.91069 | 22.47936 | 0.480447 | -0.61046 | 1.571358 | 23.12033 | 1.051009 | -6.32982 | 0.32969 | 0.447507 | 0 | 0 |
| 498 | sp\|P28652 | KCC2B_MO | 60 kDa | 0.999561 | 5 | 60% | P28652 | Camk2b | Calcium/ca | 21.79719 | 21.57455 | 21.06306 | 20.87704 | 21.43291 | 20.71653 | 0.375013 | -0.4776 | 1.227627 | 21.24355 | 1.04831 | -6.33278 | 0.330635 | 0.448342 | 0 | 0 |
| 314 | sp\|P49312 | ROA1_MOU | 34 kDa | 0.99966 | 7 | 71% | P49312 | Hnrnpa1 | Heterogene | 22.99258 | 22.15958 | 21.18278 | 22.92846 | 21.91006 | 22.57914 | -0.52725 | -1.73049 | 0.675984 | 22.2921 | -1.04573 | -6.33522 | 0.331952 | 0.449479 | 0 | 0 |
| 838 | sp\|Q9D7B6 | ACAD8_MC | 45 kDa | 0.999265 | 3 | 100% | Q9D7B6 | Acad8 | Isobutyryl- | 21.34062 | 19.95169 | 19.84672 | 21.7007 | 20.51211 | 22.10983 | -0.6876 | -2.2574 | 0.882206 | 20.91028 | -1.0453 | -6.33566 | 0.332137 | 0.449479 | 0 | 0 |
| 1222 | sp\|Q9ES74 | NEK7_MOU | 35 kDa | 0.997627 | 2 | 100% | Q9ES74 | Nek7 | Serine/thre | 17.30336 | 17.51005 | 16.46635 | 17.802 | 17.17102 | 16.83538 | -0.40223 | -1.32263 | 0.518171 | 17.18136 | -1.04261 | -6.33845 | 0.333246 | 0.450371 | 0 | 0 |
| 978 | sp\|Q8BJY1 | PSMD5_MC | 56 kDa | 0.997369 | 3 | 100% | Q8BJY1 | Psmd5 | 26S protea | 17.99708 | 18.99322 | 17.13591 | 16.03473 | 17.32847 | 13.41864 | 1.338288 | -1.72607 | 4.402648 | 16.81801 | 1.042223 | -6.3388 | 0.33346 | 0.450371 | 0 | 0 |
| 697 | sp\|Q5U4D9 | THOC6_MC | 37 kDa | 0.998918 | 4 | 100% | Q5U4D9 | Thoc6 | THO compl | 17.57242 | 18.87889 | 17.98383 | 17.10629 | 18.95096 | 22.66261 | -1.38019 | -4.54878 | 1.788396 | 18.85917 | -1.0395 | -6.34157 | 0.334636 | 0.451508 | 0 | 0 |
| 371 | sp\|Q8BMA | SRP68_MO | 71 kDa | 0.998976 | 7 | 100% | Q8BMA6 | Srp68 | Signal reco | 22.36527 | 22.85536 | 22.53385 | 22.82614 | 22.44061 | 22.48292 | -0.2749 | -0.90703 | 0.35724 | 22.58403 | -1.03505 | -6.34651 | 0.336137 | 0.453084 | 0 | 0 |
| 307 | sp\|P06151 | LDHA_MOU | 36 kDa | 0.999832 | 7 | 86% | P06151 | Ldha | L-lactate de | 21.97781 | 22.52739 | 23.03656 | 23.13185 | 21.88315 | 22.49528 | -0.419 | -1.39376 | 0.555759 | 22.50867 | -1.02581 | -6.35543 | 0.340596 | 0.458492 | 0 | 0 |
| 649 | sp\|Q9CQD1 | RAB5A_MO | 24 kDa | 0.999339 | 4 | 50% | Q9CQD1 | Rab5a | Ras-related | 20.33762 | 20.37961 | 20.13024 | 20.02577 | 20.43603 | 19.51389 | 0.328206 | -0.4347 | 1.091108 | 20.13719 | 1.02395 | -6.35772 | 0.340988 | 0.458492 | 0 | 0 |
| 640 | sp\|Q9DCD0 | 6PGD_MOU | 53 kDa | 0.9994 | 4 | 100% | Q9DCD0 | Pgd | 6-phospho | 23.19676 | 23.46002 | 21.70685 | 21.87411 | 22.37085 | 23.85979 | -0.63982 | -2.13033 | 0.850696 | 22.74473 | -1.0244 | -6.35685 | 0.341213 | 0.458492 | 0 | 0 |
| 727 | sp\|B2RSH2 | GNAI1_MO | 40 kDa | 0.998489 | 4 | 25% | B2RSH2 | Gnai1 | Guanine nu | 19.58567 | 20.01871 | 20.06553 | 20.19325 | 19.85572 | 20.12743 | -0.27749 | -0.92323 | 0.368256 | 19.97438 | -1.02278 | -6.35889 | 0.341502 | 0.458492 | 0 | 0 |
| 495 | sp\|Q922R8 | PDIA6_MO | 48 kDa | 0.999609 | 5 | 100% | Q922R8 | Pdia6 | Protein dis | 23.45302 | 24.10854 | 23.73881 | 23.15559 | 23.21565 | 24.3297 | -0.39545 | -1.3219 | 0.531007 | 23.66688 | -1.01848 | -6.36282 | 0.3438 | 0.46112 | 0 | 0 |
| 644 | sp\|P50396 | GDIA_MOU | 51 kDa | 0.999371 | 4 | 75% | P50396 | Gdi1 | Rab GDP di | 21.27701 | 22.72952 | 22.70496 | 21.70335 | 21.18493 | 22.2775 | -0.51449 | -1.72397 | 0.694981 | 21.97954 | -1.01516 | -6.36612 | 0.345291 | 0.462663 | 0 | 0 |
| 630 | sp\|Q9QXV0 | PCSK1_MO | 27 kDa | 0.999433 | 4 | 100% | Q9QXV0 | Pcsk1n | ProSAAS | 20.71248 | 20.08099 | 20.83897 | 21.9361 | 22.41646 | 19.77664 | 0.724726 | -0.98124 | 2.430693 | 20.96027 | 1.013804 | -6.36747 | 0.345891 | 0.463009 | 0 | 0 |
| 936 | sp\|Q9D5V5 | CUL5_MOU | 91 kDa | 0.998121 | 3 | 100% | Q9D5V5 | Cul5 | Cullin-5 | 19.62982 | 20.14108 | 20.61344 | 20.37752 | 19.29841 | 20.21986 | -0.39893 | -1.34032 | 0.542464 | 20.04669 | -1.01129 | -6.36998 | 0.347008 | 0.464046 | 0 | 0 |
| 619 | sp\|P62880 | GBB2_MOU | 37 kDa | 0.99954 | 4 | 50% | P62880 | Gnb2 | Guanine nu | 18.66145 | 19.46895 | 20.7353 | 18.50855 | 19.12446 | 18.76646 | 0.592415 | -0.80868 | 1.993514 | 19.21086 | 1.009039 | -6.37223 | 0.348011 | 0.464929 | 0 | 0 |
| 1241.1 | sp\|P0C0S6 | H2AZ_MOU | 14 kDa | 0.997171 | 2 | 50% | P0C0S6 | H2afz | Histone H2 | 20.56766 | 16.89908 | 20.61792 | 21.18222 | 20.36502 | 20.32369 | 1.048535 | -1.43777 | 3.534836 | 19.9926 | 1.006422 | -6.37483 | 0.34918 | 0.466031 | 0 | 0 |
| 432 | sp\|Q8BJ05 | ZC3HE_MO | 82 kDa | 0.999309 | 6 | 100% | Q8BJ05 | Zc3h14 | Zinc finger | 20.97267 | 22.06546 | 22.57092 | 25.0766 | 22.87565 | 22.0813 | -0.93471 | -3.15566 | 1.286244 | 22.6071 | -1.00435 | -6.37688 | 0.350105 | 0.466806 | 0 | 0 |
| 1065 | sp\|P51174 | ACADL_MO | 48 kDa | 0.99924 | 2 | 100% | P51174 | Acadl | Long-chain | 21.87831 | 21.67857 | 22.95876 | 22.51112 | 21.83541 | 20.93909 | 0.51457 | -0.71022 | 1.739363 | 21.96688 | 1.002612 | -6.37861 | 0.350886 | 0.466933 | 0 | 0 |
| 233 | sp\|P48036 | ANXA5_MC | 36 kDa | 0.999564 | 9 | 100% | P48036 | Anxa5 | Annexin A5 | 22.32253 | 22.97193 | 22.33126 | 24.03029 | 23.66282 | 22.85298 | -0.51286 | -1.73359 | 0.70787 | 23.02863 | -1.00261 | -6.37861 | 0.350889 | 0.466933 | 0 | 0 |
| 520 | sp\|Q9DB20 | ATPO_MOU | 23 kDa | 0.99933 | 5 | 100% | Q9DB20 | Atp5po | ATP syntha | 22.19405 | 22.36927 | 22.79422 | 22.20827 | 22.96171 | 22.44518 | 0.309088 | -0.42688 | 1.045053 | 22.49545 | 0.999601 | -6.382 | 0.351825 | 0.46772 | 0 | 0 |
| 916.14 | sp\|Q8R1M | H2AJ_MOU | 14 kDa | 0.998436 | 3 | 33% | Q8R1M2 | H2afj | Histone H2 | 20.55017 | 20.91647 | 20.73682 | 20.46705 | 19.89855 | 20.78899 | -0.32899 | -1.11598 | 0.457994 | 20.55968 | -0.995 | -6.38655 | 0.353905 | 0.469451 | 0 | 0 |
| 166 | sp\|Q7TMQ | WDR91_M | 83 kDa | 0.999462 | 11 | 100% | Q7TMQ7 | Wdr91 | WD repeat- | 22.68742 | 21.59898 | 20.86125 | 21.73298 | 23.6267 | 21.82641 | 0.672337 | -0.93895 | 2.283628 | 22.05562 | 0.995781 | -6.38536 | 0.353962 | 0.469451 | 0 | 0 |
| 1018 | sp\|P35454 | NEU1_MOU | 13 kDa | 0.999755 | 2 | 50% | P35454 | Oxt | Oxytocin-n | 20.47943 | 20.51376 | 20.70255 | 20.34504 | 20.75425 | 20.30329 | 0.258046 | -0.35958 | 0.875675 | 20.51639 | 0.99442 | -6.38711 | 0.354165 | 0.469451 | 0 | 0 |
| 440 | sp\|P62192 | PRS4_MOU | 49 kDa | 0.999207 | 6 | 100% | P62192 | Psmc1 | 26S protea | 21.47137 | 21.79434 | 21.7285 | 21.83467 | 21.39242 | 21.7357 | -0.25748 | -0.87476 | 0.359804 | 21.6595 | -0.99279 | -6.38872 | 0.354906 | 0.469589 | 0 | 0 |
| 43 | sp\|Q501J6 | DDX17_MC | 72 kDa | 0.999633 | 22 | 64% | Q501J6 | Ddx17 | Probable A | 23.96127 | 23.33629 | 23.52084 | 23.37517 | 23.65664 | 23.61145 | 0.271949 | -0.38015 | 0.924048 | 23.57694 | 0.9926 | -6.3889 | 0.354991 | 0.469589 | 0 | 0 |
| 151 | sp\|Q8K400 | STXB5_MO | 128 kDa | 0.99924 | 12 | 92% | Q8K400 | Stxbp5 | Syntaxin-bi | 24.20699 | 23.99182 | 23.40208 | 22.24267 | 24.0704 | 23.85512 | 0.529957 | -0.74392 | 1.803832 | 23.62818 | 0.992807 | -6.38829 | 0.355308 | 0.469589 | 0 | 0 |
| 376 | sp\|P46978 | STT3A_MO | 81 kDa | 0.998661 | 7 | 86% | P46978 | Stt3a | Dolichyl-di | 23.98039 | 22.85556 | 23.86563 | 23.63066 | 23.53746 | 23.84965 | 0.349206 | -0.49171 | 1.190125 | 23.61989 | 0.989083 | -6.39225 | 0.356697 | 0.470965 | 0 | 0 |
| 738 | sp\|P01803 | HVM34_M | 13 kDa | 0.998328 | 4 | 25% | P01803 | NA | Ig heavy ch | 21.00019 | 20.56115 | 21.76202 | 20.72628 | 20.83225 | 21.22736 | 0.359887 | -0.50794 | 1.22771 | 21.01821 | 0.988618 | -6.39256 | 0.357048 | 0.47097 | 0 | 0 |
| 1112 | sp\|Q6PDM | SRSF1_MOI | 28 kDa | 0.998826 | 2 | 100% | Q6PDM2 | Srsf1 | Serine/argi | 18.06532 | 17.94049 | 18.01049 | 17.1134 | 18.4661 | 18.34514 | 0.380959 | -0.54432 | 1.306237 | 17.99016 | 0.982364 | -6.39855 | 0.360036 | 0.474449 | 0 | 0 |
| 899.2 | sp\|P63163 | RSMN_MOI | 25 kDa | 0.998705 | 3 | 100% | P63163 | Snrpn | Small nucle | 19.21848 | 16.87568 | 19.38276 | 19.58512 | 19.55483 | 19.54426 | 0.717003 | -1.045 | 2.479002 | 19.02686 | 0.971104 | -6.40946 | 0.36525 | 0.480854 | 0 | 0 |
| 284 | sp\|Q8BFZ9 | ERLN2_MO | 38 kDa | 0.999317 | 8 | 100% | Q8BFZ9 | Erlin2 | Erlin-2 | 23.85399 | 23.55766 | 23.98913 | 23.68708 | 23.47934 | 23.22943 | 0.282766 | -0.41261 | 0.978147 | 23.63277 | 0.967844 | -6.41302 | 0.366361 | 0.481848 | 0 | 0 |
| 1032 | sp\|Q3UNZ8 | QORL2_MC | 38 kDa | 0.999523 | 2 | 100% | Q3UNZ8 | Cryzl2 | Quinone ox | 17.66629 | 16.0421 | 16.95147 | 17.82363 | 17.80533 | 17.07797 | 0.49313 | -0.72547 | 1.711726 | 17.2278 | 0.965723 | -6.41465 | 0.367749 | 0.483205 | 0 | 0 |
| 172 | sp\|O35286 | DHX15_MC | 91 kDa | 0.999312 | 11 | 100% | O35286 | Dhx15 | Pre-mRNA- | 23.27429 | 22.46308 | 22.84967 | 22.13879 | 23.18034 | 23.51594 | 0.395499 | -0.59624 | 1.387244 | 22.90368 | 0.951693 | -6.4281 | 0.374324 | 0.490904 | 0 | 0 |
| 1145 | tr\|A0A075 | A0A075B5S | 11 kDa | 0.998505 | 2 | 100% | A0A075B5S | Ighv9-4 | Immunoglo | 20.1762 | 18.50193 | 18.55013 | 19.61981 | 19.24259 | 18.30321 | 0.514654 | -0.7759 | 1.805211 | 19.06564 | 0.951675 | -6.42811 | 0.374332 | 0.490904 | 0 | 0 |
| 143 | sp\|O88532 | ZFR_MOUS | 117 kDa | 0.999727 | 12 | 100% | O88532 | Zfr | Zinc finger | 22.00516 | 21.81905 | 21.97103 | 22.60968 | 22.07328 | 22.50433 | -0.29454 | -1.03761 | 0.44854 | 22.16375 | -0.94342 | -6.43636 | 0.377851 | 0.495039 | 0 | 0 |
| 605 | sp\|Q8BWT | THIM_MOU | 42 kDa | 0.999678 | 4 | 100% | Q8BWT1 | Acaa2 | 3-ketoacyl- | 22.02005 | 22.32851 | 22.10197 | 21.93375 | 21.3777 | 22.1269 | -0.29648 | -1.04647 | 0.453512 | 21.98148 | -0.94089 | -6.43875 | 0.379055 | 0.495846 | 0 | 0 |
| 1233 | sp\|Q9CZJ2 | HS12B_MO | 76 kDa | 0.997383 | 2 | 100% | Q9CZJ2 | Hspa12b | Heat shock | 19.82611 | 19.5062 | 19.87865 | 19.79408 | 19.70304 | 19.34939 | 0.252714 | -0.38727 | 0.892701 | 19.67625 | 0.93985 | -6.43974 | 0.379553 | 0.495846 | 0 | 0 |
| 718 | sp\|P61089 | UBE2N_MC | 17 kDa | 0.998699 | 4 | 100% | P61089 | Ube2n | Ubiquitin-c | 21.65186 | 22.14184 | 22.4304 | 22.11026 | 21.35633 | 22.16883 | -0.32745 | -1.15675 | 0.501849 | 21.97659 | -0.93984 | -6.43974 | 0.379564 | 0.495846 | 0 | 0 |
| 504 | sp\|Q9D8N0 | EF1G_MOU | 50 kDa | 0.99953 | 5 | 100% | Q9D8N0 | Eef1g | Elongation | 22.32356 | 22.52574 | 23.27353 | 22.48682 | 22.06277 | 24.34953 | -0.56741 | -2.01111 | 0.876291 | 22.83699 | -0.93793 | -6.44114 | 0.38086 | 0.49706 | 0 | 0 |
| 979 | tr\|Q80ZX0 | Q80ZX0_M | 136 kDa | 0.997282 | 3 | 100% | Q80ZX0 | Sec24b | Sec24-relat | 13.61362 | 17.84742 | 18.20925 | 17.65841 | 18.26751 | 18.00711 | -1.14085 | -4.04913 | 1.767419 | 17.26722 | -0.93615 | -6.44282 | 0.381712 | 0.497694 | 0 | 0 |
| 287 | sp\|Q03517 | SCG2_MOU | 71 kDa | 0.999259 | 8 | 100% | Q03517 | Scg2 | Secretogra | 23.56611 | 23.04486 | 23.21411 | 24.68829 | 23.72483 | 24.05759 | -0.42856 | -1.52229 | 0.665162 | 23.71597 | -0.9351 | -6.44381 | 0.382216 | 0.497872 | 0 | 0 |
| 588 | sp\|Q99JP0 | M4K3_MOI | 101 kDa | 0.996695 | 5 | 100% | Q99JP0 | Map4k3 | Mitogen-ac | 20.31399 | 21.87734 | 19.24963 | 19.43373 | 20.36377 | 20.46836 | -0.61734 | -2.19667 | 0.961978 | 20.28447 | -0.93284 | -6.44593 | 0.383299 | 0.498804 | 0 | 0 |
| 1166 | sp\|Q99J99 | THTM_MOI | 33 kDa | 0.998285 | 2 | 100% | Q99J99 | Mpst | 3-mercapto | 20.49583 | 19.26034 | 20.56021 | 19.61356 | 20.28542 | 21.08983 | 0.459249 | -0.73687 | 1.655373 | 20.21753 | 0.916269 | -6.46137 | 0.391323 | 0.508757 | 0 | 0 |
| 569 | sp\|Q8JZX4 | SPF45_MO | 45 kDa | 0.998333 | 5 | 100% | Q8JZX4 | Rbm17 | Splicing fac | 20.76884 | 21.00199 | 19.31749 | 20.67929 | 20.19204 | 19.90931 | -0.43741 | -1.57854 | 0.703726 | 20.31149 | -0.91475 | -6.46278 | 0.392066 | 0.509235 | 0 | 0 |
| 397 | sp\|P62983 | RS27A_MO | 18 kDa | 0.999709 | 6 | 100% | P62983 | Rps27a | Ubiquitin-4 | 25.49872 | 25.82533 | 25.87552 | 25.85119 | 25.4093 | 25.84478 | -0.24592 | -0.888 | 0.396163 | 25.71747 | -0.9116 | -6.46609 | 0.393229 | 0.510257 | 0 | 0 |
| 985 | sp\|Q99JI6 | RAP1B_MO | 21 kDa | 0.997076 | 3 | 33% | Q99JI6 | Rap1b | Ras-related | 18.51436 | 22.86944 | 17.55084 | 18.56424 | 18.54881 | 17.00714 | -1.27561 | -4.61591 | 2.064704 | 18.84247 | -0.91134 | -6.46592 | 0.393734 | 0.510424 | 0 | 0 |
| 357 | sp\|Q99ME | WDR6_MO | 122 kDa | 0.999167 | 7 | 100% | Q99ME2 | Wdr6 | WD repeat- | 23.62871 | 22.82514 | 23.9283 | 25.85653 | 23.42609 | 24.2862 | -0.66159 | -2.39871 | 1.075531 | 23.99183 | -0.90889 | -6.46818 | 0.394938 | 0.511495 | 0 | 0 |
| 494 | sp\|Q64332 | SYN2_MOU | 63 kDa | 0.99965 | 5 | 80% | Q64332 | Syn2 | Synapsin-2 | 21.17829 | 21.07763 | 21.27296 | 20.62693 | 21.7488 | 21.5495 | 0.315333 | -0.51235 | 1.143012 | 21.24235 | 0.906837 | -6.47046 | 0.395577 | 0.511834 | 0 | 0 |
| 667 | sp\|P01867 | IGG2B_MO | 44 kDa | 0.999146 | 4 | 100% | P01867 | Igh-3 | Ig gamma-2 | 27.36625 | 27.68074 | 27.58628 | 28.37094 | 28.37658 | 28.31208 | -0.34488 | -1.25256 | 0.562794 | 27.94881 | -0.90583 | -6.47115 | 0.396297 | 0.512277 | 0 | 0 |
| 1043 | sp\|Q791V5 | MTCH2_MC | 33 kDa | 0.999396 | 2 | 100% | Q791V5 | Mtch2 | Mitochond | 19.64914 | 21.03393 | 21.41856 | 21.57474 | 20.86456 | 20.70817 | -0.46153 | -1.6803 | 0.757246 | 20.87485 | -0.9037 | -6.47294 | 0.397491 | 0.512476 | 0 | 0 |
| 551 | tr\|Q8C5R8 | Q8C5R8_M | 35 kDa | 0.998927 | 5 | 20% | Q8C5R8 | Prps1l1 | Phosphorib | 15.3737 | 15.25413 | 16.09713 | 16.77661 | 14.93804 | NA | -0.54575 | -2.02753 | 0.936037 | 15.68792 | -0.91293 | -6.3496 | 0.398263 | 0.512476 | 0 | 0 |
| 552.1 | sp\|Q9D7G0 | PRPS1_MO | 35 kDa | 0.998927 | 5 | 20% | Q9D7G0 | Prps1 | Ribose-pho | 15.3737 | 15.25413 | 16.09713 | 16.77661 | 14.93804 | NA | -0.54575 | -2.02753 | 0.936037 | 15.68792 | -0.91293 | -6.3496 | 0.398263 | 0.512476 | 0 | 0 |

| | | | | | | | | | | | | | | | | | | | | | | | | | |
|---|---|---|---|---|---|---|---|---|---|---|---|---|---|---|---|---|---|---|---|---|---|---|---|---|---|
| 552.2 | tr\|G3UXL2 | G3UXL2_M | 35 kDa | 0.998927 | 5 | 20% | G3UXL2 | Prps1l3 | Phosphorib | 15.3737 | 15.25413 | 16.09713 | 16.77661 | 14.93804 | NA | -0.54575 | -2.02753 | 0.936037 | 15.68792 | -0.91293 | -6.3496 | 0.398263 | 0.512476 | 0 | 0 |
| 234 | sp\|P68040 | RACK1_MO | 35 kDa | 0.999549 | 9 | 100% | P68040 | Rack1 | Receptor o | 23.12812 | 23.29674 | 23.12434 | 22.45088 | 23.56158 | 23.15558 | 0.303612 | -0.49822 | 1.105448 | 23.11954 | 0.901229 | -6.4756 | 0.39834 | 0.512476 | 0 | 0 |
| 210 | sp\|Q922F4 | TBB6_MOU | 50 kDa | 0.999274 | 10 | 10% | Q922F4 | Tubb6 | Tubulin bet | 18.24358 | 17.75739 | 18.99348 | 18.88422 | 19.11044 | 18.58991 | 0.371989 | -0.61673 | 1.360706 | 18.5965 | 0.897861 | -6.47827 | 0.400383 | 0.514615 | 0 | 0 |
| 156 | sp\|P46735 | MYO1B_MO | 129 kDa | 0.997764 | 12 | 100% | P46735 | Myo1b | Unconvent | 22.29839 | 22.89147 | 22.598 | 22.30505 | 22.39406 | 22.88695 | -0.26434 | -0.96908 | 0.440398 | 22.56232 | -0.89276 | -6.48331 | 0.40255 | 0.516911 | 0 | 0 |
| 327 | sp\|P38647 | GRP75_MO | 73 kDa | 0.999554 | 7 | 100% | P38647 | Hspa9 | Stress-70 p | 22.60589 | 23.28218 | 23.21156 | 22.99888 | 25.86785 | 23.22692 | 0.72577 | -1.218 | 2.669537 | 23.53221 | 0.891056 | -6.48445 | 0.403771 | 0.517989 | 0 | 0 |
| 1004 | sp\|Q9EP69 | SAC1_MOU | 67 kDa | 0.995413 | 3 | 100% | Q9EP69 | Sacm1l | Phosphatid | 19.02959 | 18.83593 | 20.23165 | 18.8804 | 18.97748 | 19.39891 | 0.3745 | -0.63609 | 1.385088 | 19.22566 | 0.884357 | -6.4905 | 0.407128 | 0.52085 | 0 | 0 |
| 1005 | sp\|Q2YDW | MSTO1_MO | 61 kDa | 0.995317 | 3 | 100% | Q2YDW2 | Msto1 | Protein mis | 19.02959 | 18.83593 | 20.23165 | 18.8804 | 18.97748 | 19.39891 | 0.3745 | -0.63609 | 1.385088 | 19.22566 | 0.884357 | -6.4905 | 0.407128 | 0.52085 | 0 | 0 |
| 724 | sp\|Q8R001 | MARE2_MO | 37 kDa | 0.998599 | 4 | 100% | Q8R001 | Mapre2 | Microtubul | 20.94129 | 18.07386 | 20.39233 | 22.55165 | 20.97621 | 18.72213 | 0.987397 | -1.67726 | 3.65205 | 20.27624 | 0.884305 | -6.49055 | 0.407154 | 0.52085 | 0 | 0 |
| 1229 | sp\|Q91WQ | SYYC_MOU | 59 kDa | 0.997532 | 2 | 100% | Q91WQ3 | Yars | Tyrosine--tl | 18.69858 | 18.45269 | 19.22397 | 17.54956 | 18.75455 | 19.37554 | 0.433104 | -0.74118 | 1.607384 | 18.67582 | 0.880179 | -6.49426 | 0.409232 | 0.523014 | 0 | 0 |
| 563 | sp\|Q3UGY8 | BIG3_MOU | 240 kDa | 0.998646 | 5 | 100% | Q3UGY8 | Arfgef3 | Brefeldin A | 20.78891 | 22.74401 | 21.24238 | 20.29615 | 21.76054 | 22.57253 | -0.60696 | -2.25753 | 1.043624 | 21.56742 | -0.87755 | -6.49661 | 0.41056 | 0.524218 | 0 | 0 |
| 450 | sp\|Q99LC3 | NDUAA_MO | 41 kDa | 0.99905 | 6 | 100% | Q99LC3 | Ndufa10 | NADH dehy | 22.84203 | 23.20579 | 22.62237 | 23.09226 | 23.16437 | 23.05759 | -0.24229 | -0.90058 | 0.416001 | 22.9974 | -0.87603 | -6.49837 | 0.410967 | 0.524244 | 0 | 0 |
| 138 | sp\|Q91W3 | NCOA5_MO | 65 kDa | 0.999193 | 13 | 100% | Q91W39 | Ncoa5 | Nuclear rec | 23.69294 | 22.71697 | 23.73591 | 24.60557 | 23.25543 | 24.93077 | -0.52301 | -1.95117 | 0.905146 | 23.82293 | -0.87395 | -6.49983 | 0.412385 | 0.525558 | 0 | 0 |
| 977 | sp\|Q9QXX4 | CMC2_MOU | 74 kDa | 0.997372 | 3 | 33% | Q9QXX4 | Slc25a13 | Calcium-bi | 14.71319 | 16.20681 | 17.39441 | 17.07012 | 16.14617 | 16.74452 | -0.58923 | -2.20038 | 1.021922 | 16.3792 | -0.87277 | -6.50088 | 0.412982 | 0.525825 | 0 | 0 |
| 421 | sp\|Q8VEM8 | MPCP_MOU | 40 kDa | 0.99946 | 6 | 100% | Q8VEM8 | Slc25a3 | Phosphate | 23.74263 | 23.55037 | 25.03682 | 24.06591 | 23.70909 | 23.74246 | 0.376601 | -0.65598 | 1.409185 | 23.97455 | 0.870376 | -6.50301 | 0.414199 | 0.526879 | 0 | 0 |
| 537 | sp\|Q9ERK4 | XPO2_MOU | 110 kDa | 0.999209 | 5 | 100% | Q9ERK4 | Cse1l | Exportin-2 | 20.79164 | 20.64448 | 21.54968 | 20.55356 | 20.57623 | 20.83262 | 0.29563 | -0.51532 | 1.106583 | 20.8247 | 0.86771 | -6.50576 | 0.415205 | 0.527159 | 0 | 0 |
| 580 | sp\|Q9QUP5 | HPLN1_MO | 40 kDa | 0.997599 | 5 | 100% | Q9QUP5 | Hapln1 | Hyaluronar | 22.02736 | 19.64185 | 18.98758 | 19.97603 | 19.70585 | 19.08239 | 0.673508 | -1.17919 | 2.52621 | 19.90351 | 0.867536 | -6.50553 | 0.415646 | 0.527159 | 0 | 0 |
| 846 | sp\|Q9QXL2 | KI21A_MO | 187 kDa | 0.999184 | 3 | 100% | Q9QXL2 | Kif21a | Kinesin-like | 21.47721 | 21.9467 | 21.17533 | 21.08011 | 21.61225 | 22.20781 | -0.32328 | -1.21203 | 0.565467 | 21.58323 | -0.86715 | -6.50603 | 0.415699 | 0.527159 | 0 | 0 |
| 615 | sp\|P14152 | MDHC_MO | 37 kDa | 0.999556 | 4 | 100% | P14152 | Mdh1 | Malate deh | 22.40132 | 23.1796 | 23.65911 | 24.09262 | 23.45055 | 23.38866 | -0.3833 | -1.43879 | 0.672189 | 23.36198 | -0.86663 | -6.50633 | 0.416107 | 0.527159 | 0 | 0 |
| 377 | sp\|Q9WTM | RUVB2_MO | 51 kDa | 0.998589 | 7 | 100% | Q9WTM5 | Ruvbl2 | RuvB-like 2 | 24.87667 | 25.25968 | 24.24096 | 24.34858 | 25.02871 | 25.64906 | -0.37033 | -1.39137 | 0.650717 | 24.90061 | -0.86555 | -6.50729 | 0.416661 | 0.527159 | 0 | 0 |
| 312 | sp\|Q06890 | CLUS_MOU | 52 kDa | 0.999684 | 7 | 100% | Q06890 | Clu | Clusterin | 23.54192 | 23.0837 | 23.17588 | 22.67431 | 23.29486 | 23.44701 | 0.269214 | -0.47183 | 1.010263 | 23.20295 | 0.864673 | -6.50846 | 0.416751 | 0.527159 | 0 | 0 |
| 283 | sp\|Q9QYX7 | PCLO_MOU | 551 kDa | 0.999324 | 8 | 100% | Q9QYX7 | Pclo | Protein pic | 20.14311 | 20.43764 | 18.60328 | 20.35687 | 21.07699 | 20.60297 | -0.5247 | -1.98292 | 0.933525 | 20.20348 | -0.85869 | -6.51334 | 0.420177 | 0.530997 | 0 | 0 |
| 279 | sp\|O55029 | COPB2_MO | 102 kDa | 0.999399 | 8 | 100% | O55029 | Copb2 | Coatomer s | 23.47247 | 23.11115 | 23.40375 | 25.75206 | 23.46798 | 23.28906 | -0.60269 | -2.27938 | 1.074 | 23.74941 | -0.85781 | -6.51412 | 0.42063 | 0.531074 | 0 | 0 |
| 561 | sp\|Q99JR1 | SFXN1_MO | 36 kDa | 0.99881 | 5 | 100% | Q99JR1 | Sfxn1 | Sideroflexir | 27.07349 | 24.86069 | 23.05498 | 27.04255 | 22.95534 | 24.37574 | -1.06506 | -4.04586 | 1.915749 | 24.8938 | -0.85269 | -6.51861 | 0.423274 | 0.533915 | 0 | 0 |
| 448 | sp\|Q6P9Z1 | SMRD3_MO | 55 kDa | 0.999065 | 6 | 83% | Q6P9Z1 | Smarcd3 | SWI/SNF-re | 20.44882 | 18.04715 | 19.53106 | 21.11497 | 20.76712 | 19.59368 | 0.663739 | -1.20016 | 2.527636 | 19.91713 | 0.849817 | -6.52111 | 0.424759 | 0.535291 | 0 | 0 |
| 384 | sp\|O08788 | DCTN1_MO | 142 kDa | 0.997846 | 7 | 100% | O08788 | Dctn1 | Dynactin su | 20.77595 | 21.78121 | 21.37536 | 21.65293 | 22.37911 | 22.26562 | -0.38978 | -1.48603 | 0.706473 | 21.70503 | -0.84851 | -6.52225 | 0.425436 | 0.535646 | 0 | 0 |
| 859.1 | sp\|A2A8L1 | CHD5_MOU | 223 kDa | 0.99907 | 3 | 100% | A2A8L1 | Chd5 | Chromodo | 20.5247 | 20.64026 | 20.70125 | 21.02256 | 20.74777 | 20.98767 | -0.22559 | -0.86517 | 0.413994 | 20.7707 | -0.8395 | -6.53045 | 0.429786 | 0.540621 | 0 | 0 |
| 282 | sp\|O35129 | PHB2_MOU | 33 kDa | 0.999352 | 8 | 100% | O35129 | Phb2 | Prohibitin- | 23.17873 | 23.53199 | 23.42839 | 23.62756 | 23.45151 | 23.54396 | -0.21496 | -0.82519 | 0.39527 | 23.46036 | -0.83843 | -6.53138 | 0.430347 | 0.540825 | 0 | 0 |
| 115 | sp\|Q8K4Z5 | SF3A1_MO | 89 kDa | 0.99977 | 14 | 100% | Q8K4Z5 | Sf3a1 | Splicing fac | 24.38031 | 23.60007 | 23.05652 | 23.54372 | 24.49486 | 23.7132 | 0.358233 | -0.67735 | 1.393819 | 23.79811 | 0.825525 | -6.54206 | 0.437486 | 0.549287 | 0 | 0 |
| 509 | sp\|Q8CH25 | SLTM_MOU | 117 kDa | 0.999485 | 5 | 100% | Q8CH25 | Sltm | SAFB-like tr | 20.54435 | 20.50325 | 20.07133 | 20.34102 | 21.04583 | 22.2248 | -0.46918 | -1.82722 | 0.888849 | 20.78843 | -0.82449 | -6.54294 | 0.438036 | 0.549469 | 0 | 0 |
| 1008 | sp\|O54781 | SRPK2_MO | 77 kDa | 0.994925 | 3 | 100% | O54781 | Srpk2 | SRSF protei | 17.84747 | 17.02365 | 19.49351 | 20.82396 | 22.13089 | 18.39678 | 1.075826 | -2.05966 | 4.211316 | 19.28604 | 0.818817 | -6.54775 | 0.441047 | 0.552735 | 0 | 0 |
| 42 | sp\|Q9QXS1 | PLEC_MOU | 534 kDa | 0.999798 | 22 | 91% | Q9QXS1 | Plec | Plectin | 21.27534 | 21.50789 | 19.03068 | 20.93863 | 21.87201 | 21.50565 | -0.59138 | -2.32525 | 1.142484 | 21.0217 | -0.81396 | -6.55185 | 0.443638 | 0.55547 | 0 | 0 |
| 1104 | sp\|P32067 | LA_MOUSE | 48 kDa | 0.998931 | 2 | 100% | P32067 | Ssb | Lupus La pr | 20.01918 | 20.60156 | 21.11963 | 20.78247 | 19.76958 | 20.51648 | -0.3307 | -1.30301 | 0.641602 | 20.46815 | -0.81168 | -6.55377 | 0.444857 | 0.556482 | 0 | 0 |
| 64 | sp\|P27773 | PDIA3_MO | 57 kDa | 0.999569 | 18 | 100% | P27773 | Pdia3 | Protein dis | 24.38726 | 25.09534 | 25.66653 | 25.41644 | 24.77461 | 25.25201 | -0.3118 | -1.23713 | 0.613539 | 25.0987 | -0.80397 | -6.56025 | 0.448979 | 0.561121 | 0 | 0 |
| 197 | sp\|Q8K3I4 | MYRIP_MO | 95 kDa | 0.999504 | 10 | 100% | Q8K3I4 | Myrip | Rab effecto | 22.9535 | 23.70629 | 22.69515 | 21.50695 | 23.04048 | 22.11664 | 0.45308 | -0.90257 | 1.808728 | 22.66984 | 0.797588 | -6.56553 | 0.452454 | 0.564878 | 0 | 0 |
| 395 | sp\|Q9DBG6 | RPN2_MOU | 69 kDa | 0.999742 | 6 | 100% | Q9DBG6 | Rpn2 | Dolichyl-di | 23.176 | 23.30928 | 23.77754 | 22.80779 | 23.16465 | 23.24731 | 0.251272 | -0.49976 | 1.002308 | 23.24709 | 0.796313 | -6.56695 | 0.452819 | 0.564878 | 0 | 0 |
| 152 | sp\|P02088 | HBB1_MOU | 16 kDa | 0.999235 | 12 | 17% | P02088 | Hbb-b1 | Hemoglobi | 27.20821 | 27.32291 | 27.35415 | 26.57818 | 26.98872 | 26.91582 | 0.244722 | -0.4877 | 0.977145 | 27.06133 | 0.795263 | -6.56782 | 0.453389 | 0.56507 | 0 | 0 |
| 1042 | sp\|P61514 | RL37A_MO | 10 kDa | 0.9994 | 2 | 100% | P61514 | Rpl37a | 60S ribosor | 18.52674 | 18.99861 | 18.9645 | 18.35634 | 18.12218 | 19.10495 | -0.28216 | -1.13391 | 0.569586 | 18.67889 | -0.78956 | -6.57232 | 0.456664 | 0.568147 | 0 | 0 |
| 1000 | sp\|Q9Z130 | HNRDL_MO | 34 kDa | 0.995659 | 3 | 33% | Q9Z130 | Hnrnpdl | Heterogene | 15.64052 | 16.48313 | 17.55449 | 18.10705 | 15.99414 | 16.22825 | -0.54309 | -2.18411 | 1.097925 | 16.66793 | -0.78979 | -6.57196 | 0.456696 | 0.568147 | 0 | 0 |
| 241 | sp\|Q99JY0 | ECHB_MOU | 51 kDa | 0.999181 | 9 | 100% | Q99JY0 | Hadhb | Trifunction | 24.82575 | 24.57784 | 24.83541 | 24.6437 | 24.59079 | 24.41817 | 0.204082 | -0.41255 | 0.820711 | 24.64861 | 0.78774 | -6.574 | 0.457492 | 0.568615 | 0 | 0 |
| 721 | sp\|Q07076 | ANXA7_MO | 50 kDa | 0.998675 | 4 | 100% | Q07076 | Anxa7 | Annexin A7 | 19.86612 | 20.26402 | 20.35462 | 19.49889 | 20.14255 | 19.84985 | 0.250173 | -0.50751 | 1.007856 | 19.99601 | 0.785874 | -6.57553 | 0.458513 | 0.569363 | 0 | 0 |
| 451 | sp\|Q99MR | SRRT_MOU | 100 kDa | 0.999027 | 6 | 100% | Q99MR6 | Srrt | Serrate RNA | 22.35835 | 20.59488 | 21.76943 | 23.15277 | 21.79816 | 20.44457 | 0.577903 | -1.1802 | 2.336009 | 21.68636 | 0.784441 | -6.57634 | 0.459621 | 0.570216 | 0 | 0 |
| 723 | sp\|P62874 | GBB1_MOU | 37 kDa | 0.998625 | 4 | 50% | P62874 | Gnb1 | Guanine nu | 20.48363 | 20.24224 | 22.07345 | 20.94028 | 20.21039 | 20.33389 | 0.41702 | -0.86757 | 1.701614 | 20.71398 | 0.774714 | -6.58425 | 0.464974 | 0.576329 | 0 | 0 |
| 473 | sp\|O88738 | BIRC6_MO | 532 kDa | 0.997622 | 6 | 100% | O88738 | Birc6 | Baculoviral | 19.78433 | 19.10187 | 19.46946 | 20.917 | 20.65113 | 21.40261 | -0.50552 | -2.06481 | 1.053767 | 20.22107 | -0.77368 | -6.58508 | 0.465544 | 0.57651 | 0 | 0 |
| 663 | sp\|P63044 | VAMP2_MO | 13 kDa | 0.999194 | 4 | 50% | P63044 | Vamp2 | Vesicle-ass | 22.39879 | 20.19269 | 20.82648 | 22.3521 | 22.06751 | 21.2371 | 0.503633 | -1.05858 | 2.065851 | 21.51245 | 0.769349 | -6.58857 | 0.467944 | 0.578953 | 0 | 0 |
| 67 | sp\|Q6URW | MYH14_MO | 229 kDa | 0.999296 | 18 | 61% | Q6URW6 | Myh14 | Myosin-14 | 20.49185 | 21.29343 | 20.83259 | 20.83399 | 20.99145 | 20.86522 | -0.22558 | -0.92547 | 0.474314 | 20.88476 | -0.76713 | -6.59071 | 0.468863 | 0.579129 | 0 | 0 |
| 1285 | sp\|Q8BK72 | RT27_MOU | 48 kDa | 0.995366 | 2 | 100% | Q8BK72 | Mrps27 | 28S ribosor | 18.15662 | 18.08956 | 18.97539 | 17.58982 | 19.35127 | 19.48795 | 0.438648 | -0.92517 | 1.802468 | 18.60844 | 0.767555 | -6.59001 | 0.46894 | 0.579129 | 0 | 0 |
| 17 | sp\|E9QAT4 | SC16A_MO | 254 kDa | 0.999775 | 33 | 100% | E9QAT4 | Sec16a | Protein tra | 25.65975 | 25.0515 | 25.36399 | 25.07134 | 25.17908 | 25.42681 | 0.217724 | -0.45979 | 0.895238 | 25.29208 | 0.764874 | -6.59251 | 0.470119 | 0.580056 | 0 | 0 |
| 886 | sp\|Q61595 | KTN1_MOU | 153 kDa | 0.998844 | 3 | 100% | Q61595 | Ktn1 | Kinectin | 20.08275 | 19.8958 | 19.93524 | 20.56563 | 19.58447 | 19.87592 | -0.24496 | -1.0121 | 0.522181 | 19.98997 | -0.76001 | -6.59639 | 0.472834 | 0.582876 | 0 | 0 |
| 647 | sp\|P14094 | AT1B1_MO | 35 kDa | 0.99935 | 4 | 100% | P14094 | Atp1b1 | Sodium/po | 22.14003 | 22.47536 | 23.12338 | 22.05102 | 22.28595 | 22.21878 | 0.268067 | -0.57393 | 1.110062 | 22.38242 | 0.758314 | -6.59764 | 0.47387 | 0.583622 | 0 | 0 |
| 1076 | sp\|Q5I012 | S38AA_MO | 117 kDa | 0.999159 | 2 | 100% | Q5I012 | Slc38a10 | Putative so | 19.62342 | 20.02419 | 20.14219 | 18.95157 | 19.26409 | 19.13336 | 0.306861 | -0.65958 | 1.273298 | 19.52314 | 0.757738 | -6.59785 | 0.474418 | 0.583766 | 0 | 0 |
| 556 | sp\|Q9CQE8 | RTRAF_MO | 28 kDa | 0.998864 | 5 | 100% | Q9CQE8 | RTRAF | RNA transc | 22.03878 | 21.55377 | 21.01344 | 21.64723 | 21.6659 | 22.40987 | -0.29759 | -1.24356 | 0.648388 | 21.7215 | -0.75073 | -6.60339 | 0.478355 | 0.588078 | 0 | 0 |
| 1106 | sp\|A2RTL5 | RSRC2_MO | 44 kDa | 0.99891 | 2 | 100% | A2RTL5 | Rsrc2 | Arginine/se | 17.73595 | 19.10717 | 19.39336 | 18.56739 | 19.2903 | 20.06157 | -0.43884 | -1.84101 | 0.963343 | 19.02596 | -0.74688 | -6.60641 | 0.480528 | 0.589968 | 0 | 0 |
| 293 | sp\|O35643 | AP1B1_MO | 104 kDa | 0.998995 | 8 | 12% | O35643 | Ap1b1 | AP-1 comp | 17.59438 | 19.14216 | 18.71509 | 18.10126 | 17.95741 | 18.02964 | -0.3354 | -1.40766 | 0.736867 | 18.25666 | -0.74646 | -6.60674 | 0.480763 | 0.589968 | 0 | 0 |
| 417 | sp\|Q91V61 | SFXN3_MO | 35 kDa | 0.999544 | 6 | 100% | Q91V61 | Sfxn3 | Sideroflexir | 23.16663 | 23.23275 | 23.11899 | 22.09398 | 22.64576 | 22.77427 | 0.276794 | -0.61996 | 1.173546 | 22.83873 | 0.735832 | -6.61515 | 0.486677 | 0.596368 | 0 | 0 |
| 154 | tr\|A0A1W2 | A0A1W2P7 | 92 kDa | 0.998967 | 12 | 8% | A0A1W2P7 | Gm32717 | Predicted g | 17.95369 | 18.27901 | 15.44668 | 18.25918 | 18.91957 | 17.6597 | -0.62598 | -2.65647 | 1.404498 | 17.75297 | -0.73572 | -6.6151 | 0.486858 | 0.596368 | 0 | 0 |
| 840 | sp\|P62911 | RL32_MOU | 16 kDa | 0.999242 | 3 | 100% | P62911 | Rpl32 | 60S ribosor | 18.28103 | 20.69263 | 18.42216 | 14.75673 | 17.89942 | 16.03677 | 1.038826 | -2.35028 | 4.427932 | 17.68146 | 0.731489 | -6.61837 | 0.489275 | 0.598788 | 0 | 0 |
| 37 | sp\|Q921M | SF3B3_MO | 136 kDa | 0.999613 | 23 | 100% | Q921M3 | Sf3b3 | Splicing fac | 25.64794 | 25.23526 | 24.34345 | 24.97936 | 25.86222 | 24.62941 | 0.336525 | -0.76367 | 1.436715 | 25.11627 | 0.729962 | -6.61955 | 0.49015 | 0.599317 | 0 | 0 |
| 1087 | sp\|Q64737 | PUR2_MOU | 108 kDa | 0.999065 | 2 | 100% | Q64737 | Gart | Trifunction | 22.93929 | 22.68533 | 22.87242 | 22.97413 | 23.01155 | 22.59121 | 0.190864 | -0.43652 | 0.818253 | 22.84565 | 0.724084 | -6.62438 | 0.493229 | 0.602539 | 0 | 0 |
| 1244 | sp\|Q9CQT1 | MTNA_MOU | 39 kDa | 0.997045 | 2 | 100% | Q9CQT1 | Mri1 | Methylthio | 18.89307 | 18.24818 | 17.94123 | 17.54076 | 18.2714 | 18.46826 | 0.282833 | -0.6502 | 1.215861 | 18.22715 | 0.723411 | -6.62457 | 0.49391 | 0.602828 | 0 | 0 |
| 345 | sp\|P62823 | RAB3C_MO | 26 kDa | 0.99936 | 7 | 43% | P62823 | Rab3c | Ras-related | 21.05358 | 21.422 | 21.33234 | 21.5557 | 20.83664 | 20.91379 | -0.22298 | -0.9581 | 0.512142 | 21.18568 | -0.72194 | -6.62602 | 0.494461 | 0.602958 | 0 | 0 |
| 1044 | sp\|P62307 | RUXF_MOU | 10 kDa | 0.999395 | 2 | 100% | P62307 | Snrpf | Small nucle | 19.43933 | 18.93213 | 20.63188 | 19.08913 | 18.99318 | 19.95017 | 0.364319 | -0.8579 | 1.586534 | 19.50597 | 0.711352 | -6.6337 | 0.500882 | 0.610159 | 0 | 0 |
| 1252 | sp\|Q6NSW | SPKAP_MO | 185 kDa | 0.996787 | 2 | 100% | Q6NSW3 | Sphkap | A-kinase an | 16.86386 | 16.99018 | 17.16571 | 15.96874 | 16.41088 | 16.67591 | 0.268538 | -0.63249 | 1.169567 | 16.67921 | 0.710494 | -6.63448 | 0.501266 | 0.610159 | 0 | 0 |
| 1284 | sp\|Q80T69 | RSBN1_MO | 89 kDa | 0.995403 | 2 | 100% | Q80T69 | Rsbn1 | Lysine-spec | 20.7875 | 21.33942 | 20.92294 | 21.14441 | 21.03247 | 20.8425 | -0.19447 | -0.85057 | 0.461629 | 21.01154 | -0.70548 | -6.63842 | 0.504013 | 0.612953 | 0 | 0 |
| 393 | sp\|P01942 | HBA_MOUS | 15 kDa | 0.999751 | 6 | 100% | P01942 | Hba | Hemoglobi | 26.46401 | 26.63959 | 26.74094 | 26.81191 | 26.31126 | 26.62693 | -0.18741 | -0.82663 | 0.451822 | 26.59911 | -0.6978 | -6.64413 | 0.508514 | 0.617872 | 0 | 0 |
| 349 | sp\|Q8K3H0 | DP13A_MO | 79 kDa | 0.999321 | 7 | 100% | Q8K3H0 | Appl1 | DCC-intera | 22.19632 | 22.04202 | 22.30079 | 21.03712 | 21.46042 | 22.02368 | 0.284905 | -0.69085 | 1.260656 | 21.84339 | 0.696805 | -6.64455 | 0.509378 | 0.618368 | 0 | 0 |
| 547 | sp\|Q62158 | TRI27_MO | 59 kDa | 0.99902 | 5 | 100% | Q62158 | Trim27 | Zinc finger | 19.25237 | 18.23601 | 17.57046 | 19.03589 | 19.73267 | 18.01041 | 0.424394 | -1.03513 | 1.883919 | 18.63963 | 0.693918 | -6.64668 | 0.511075 | 0.619873 | 0 | 0 |
| 808 | sp\|Q8K2B3 | SDHA_MOU | 73 kDa | 0.999462 | 3 | 100% | Q8K2B3 | Sdha | Succinate d | 21.86345 | 22.38283 | 21.19742 | 21.06094 | 21.97075 | 22.58293 | -0.3317 | -1.47796 | 0.814565 | 21.84305 | -0.69057 | -6.64914 | 0.513046 | 0.621529 | 0 | 0 |
| 517 | sp\|Q9R1A8 | COP1_MOU | 80 kDa | 0.999365 | 5 | 100% | Q9R1A8 | Cop1 | E3 ubiquiti | 22.08696 | 21.79764 | 20.48077 | 20.93977 | 22.61125 | 21.22947 | 0.404036 | -0.99327 | 1.801346 | 21.52431 | 0.690046 | -6.64952 | 0.513357 | 0.621529 | 0 | 0 |
| 426 | sp\|O35526 | STX1A_MO | 33 kDa | 0.999408 | 6 | 100% | O35526 | Stx1a | Syntaxin-1A | 22.50487 | 21.94678 | 20.98354 | 21.50129 | 21.57472 | 22.62293 | -0.33596 | -1.50554 | 0.833629 | 21.85569 | -0.68549 | -6.65285 | 0.516049 | 0.623949 | 0 | 0 |
| 1278.2 | sp\|Q7TSF1 | DSG1B_MO | 114 kDa | 0.995714 | 2 | 100% | Q7TSF1 | Dsg1b | Desmoglein | 15.88352 | 20.07099 | 16.13897 | 15.54141 | 17.73233 | 16.55523 | -0.80427 | -3.61215 | 2.003609 | 16.98708 | -0.68356 | -6.65426 | 0.517197 | 0.623949 | 0 | 0 |
| 1278.3 | sp\|Q61495 | DSG1A_MO | 115 kDa | 0.995714 | 2 | 100% | Q61495 | Dsg1a | Desmoglein | 15.88352 | 20.07099 | 16.13897 | 15.54141 | 17.73233 | 16.55523 | -0.80427 | -3.61215 | 2.003609 | 16.98708 | -0.68356 | -6.65426 | 0.517197 | 0.623949 | 0 | 0 |
| 1279 | sp\|O89112 | LANC1_MO | 45 kDa | 0.99571 | 2 | 100% | O89112 | Lancl1 | Glutathion | 15.88352 | 20.07099 | 16.13897 | 15.54141 | 17.73233 | 16.55523 | -0.80427 | -3.61215 | 2.003609 | 16.98708 | -0.68356 | -6.65426 | 0.517197 | 0.623949 | 0 | 0 |
| 189 | sp\|Q91VD9 | NDUS1_MO | 80 kDa | 0.999673 | 10 | 100% | Q91VD9 | Ndufs1 | NADH-ubiq | 22.85896 | 22.89416 | 23.25365 | 22.70019 | 22.58183 | 23.94331 | -0.28107 | -1.2689 | 0.706753 | 23.03868 | -0.67903 | -6.65754 | 0.519884 | 0.626586 | 0 | 0 |
| 62 | sp\|P11499 | HS90B_MO | 83 kDa | 0.999751 | 18 | 50% | P11499 | Hsp90ab1 | Heat shock | 24.64623 | 25.0828 | 25.59178 | 25.30348 | 24.47295 | 25.07175 | -0.24902 | -1.12579 | 0.627746 | 25.02817 | -0.67709 | -6.65906 | 0.520933 | 0.626586 | 0 | 0 |
| 438 | sp\|Q9EQK5 | MVP_MOU | 96 kDa | 0.99922 | 6 | 100% | Q9EQK5 | Mvp | Major vault | 22.52034 | 21.75879 | 20.57396 | 21.97124 | 21.34734 | 21.73586 | -0.34142 | -1.54451 | 0.861672 | 21.65125 | -0.67723 | -6.65884 | 0.520955 | 0.626586 | 0 | 0 |
| 285 | sp\|P10630 | IF4A2_MO | 46 kDa | 0.999295 | 8 | 25% | P10630 | Eif4a2 | Eukaryotic | 18.98459 | 18.80082 | 19.41474 | 19.33311 | 19.11248 | 18.76908 | 0.202934 | -0.51125 | 0.917117 | 19.06914 | 0.676313 | -6.6598 | 0.521231 | 0.626586 | 0 | 0 |
| 1072 | sp\|Q9WUC | LY6H_MOU | 15 kDa | 0.999194 | 2 | 100% | Q9WUC3 | Ly6h | Lymphocyt | 23.806 | 24.17645 | 24.31532 | 24.8389 | 23.56914 | 23.51469 | -0.27986 | -1.28051 | 0.720787 | 24.03675 | -0.66744 | -6.66585 | 0.526808 | 0.632729 | 0 | 0 |
| 264 | sp\|Q5DQR4 | STB5L_MO | 132 kDa | 0.999657 | 8 | 88% | Q5DQR4 | Stxbp5l | Syntaxin-bi | 24.28436 | 25.6371 | 24.3766 | 24.87379 | 24.93977 | 20.66351 | 0.80878 | -2.12654 | 3.744094 | 24.12919 | 0.657546 | -6.67285 | 0.532767 | 0.639321 | 0 | 0 |
| 1197 | sp\|Q8C522 | ENDD1_MO | 55 kDa | 0.997966 | 2 | 100% | Q8C522 | Endod1 | Endonuclea | 17.41515 | 17.38588 | 16.37274 | 14.26591 | 16.91452 | 17.34835 | 0.567421 | -1.50203 | 2.636873 | 16.61709 | 0.654336 | -6.6751 | 0.534709 | 0.641084 | 0 | 0 |
| 286 | sp\|O08709 | PRDX6_MO | 25 kDa | 0.999289 | 8 | 100% | O08709 | Prdx6 | Peroxiredo | 23.58778 | 23.54127 | 23.24734 | 23.24801 | 23.38377 | 22.84712 | 0.194162 | -0.52125 | 0.909577 | 23.30922 | 0.645962 | -6.68121 | 0.539537 | 0.6463 | 0 | 0 |
| 354 | sp\|Q61136 | PRP4B_MO | 117 kDa | 0.999238 | 7 | 100% | Q61136 | Prpf4b | Serine/thre | 21.36521 | 20.67229 | 19.84939 | 20.2666 | 21.38417 | 20.73978 | 0.306694 | -0.83044 | 1.443825 | 20.71291 | 0.643642 | -6.68252 | 0.54121 | 0.647732 | 0 | 0 |
| 477 | sp\|Q9Z2T6 | KRT85_MO | 56 kDa | 0.996805 | 6 | 83% | Q9Z2T6 | Krt85 | Keratin, typ | 17.63642 | 21.38056 | 17.75515 | 16.35591 | 17.36254 | 17.35905 | -0.78047 | -3.67913 | 2.118186 | 17.97494 | -0.64255 | -6.68327 | 0.541874 | 0.647955 | 0 | 0 |
| 711 | sp\|P68181 | KAPCB_MO | 41 kDa | 0.998773 | 4 | 50% | P68181 | Prkacb | cAMP-depe | 16.99514 | 17.78472 | 15.17535 | 14.10718 | 17.28583 | 15.64124 | 0.641061 | -1.74457 | 3.026696 | 16.16491 | 0.641278 | -6.68415 | 0.542654 | 0.648316 | 0 | 0 |
| 407 | sp\|Q00623 | APOA1_MO | 31 kDa | 0.9996 | 6 | 100% | Q00623 | Apoa1 | Apolipopro | 23.00534 | 22.96098 | 23.44589 | 23.83148 | 22.93709 | 23.24269 | -0.21561 | -1.02096 | 0.589744 | 23.23724 | -0.63723 | -6.6872 | 0.54488 | 0.650403 | 0 | 0 |
| 1134 | sp\|O35405 | PLD3_MOU | 54 kDa | 0.998581 | 2 | 100% | O35405 | Pld3 | Phospholip | 19.63274 | 18.3577 | 18.81846 | 19.07016 | 18.62872 | 18.92696 | 0.2417 | -0.66462 | 1.148021 | 18.90579 | 0.635753 | -6.68805 | 0.545937 | 0.651091 | 0 | 0 |
| 510 | sp\|P47738 | ALDH2_MO | 57 kDa | 0.99947 | 5 | 100% | P47738 | Aldh2 | Aldehyde d | 22.22193 | 22.29382 | 21.78754 | 21.87387 | 20.9932 | 21.59611 | -0.25371 | -1.21262 | 0.705197 | 21.79441 | -0.63142 | -6.69088 | 0.548701 | 0.653011 | 0 | 0 |
| 1238 | sp\|Q8VE80 | THOC3_MO | 39 kDa | 0.997247 | 2 | 100% | Q8VE80 | Thoc3 | THO compl | 20.55642 | 25.87945 | 20.91534 | 19.5815 | 20.01091 | 19.23768 | -1.07199 | -5.12507 | 2.981099 | 21.03022 | -0.63118 | -6.69104 | 0.548847 | 0.653011 | 0 | 0 |
| 252 | sp\|Q61414 | K1C15_MO | 49 kDa | 0.998201 | 9 | 22% | Q61414 | Krt15 | Keratin, typ | 18.2861 | 19.88792 | 22.21774 | 20.68979 | 19.85278 | 21.6699 | -0.63033 | -3.01446 | 1.753792 | 20.43404 | -0.63095 | -6.6912 | 0.548992 | 0.653011 | 0 | 0 |
| 843 | sp\|Q8BMC | NOP9_MOU | 70 kDa | 0.999222 | 3 | 100% | Q8BMC4 | Nop9 | Nucleolar p | 23.19132 | 23.37565 | 22.84085 | 23.57728 | 23.68111 | 23.34961 | -0.19642 | -0.94447 | 0.55162 | 23.33597 | -0.62498 | -6.6955 | 0.552422 | 0.656516 | 0 | 0 |
| 478 | sp\|Q8BML9 | SYQ_MOUS | 88 kDa | 0.996522 | 6 | 100% | Q8BML9 | Qars | Glutamine- | 20.39094 | 20.29395 | 20.79928 | 20.58554 | 20.48987 | 20.29174 | 0.169619 | -0.48324 | 0.822479 | 20.47522 | 0.61838 | -6.69991 | 0.556514 | 0.660799 | 0 | 0 |
| 755 | sp\|P15920 | VPP2_MOU | 98 kDa | 0.997606 | 4 | 50% | P15920 | Atp6v0a2 | V-type prot | 19.06511 | 19.35388 | 18.35353 | 19.56282 | 19.05765 | 18.33881 | -0.25974 | -1.26715 | 0.747666 | 18.9553 | -0.6153 | -6.70168 | 0.558673 | 0.662783 | 0 | 0 |
| 217 | sp\|Q5ND34 | WDR81_MO | 212 kDa | 0.998748 | 10 | 100% | Q5ND34 | Wdr81 | WD repeat- | 20.87734 | 21.34136 | 19.88104 | 20.54273 | 21.7146 | 21.56276 | -0.32462 | -1.59786 | 0.948614 | 20.98664 | -0.60845 | -6.7062 | 0.562948 | 0.666829 | 0 | 0 |
| 879 | sp\|P35564 | CALX_MOU | 67 kDa | 0.998936 | 3 | 100% | P35564 | Canx | Calnexin | 17.62096 | 17.24647 | 17.71825 | 18.14069 | 18.07511 | 18.83872 | -0.27052 | -1.33189 | 0.790848 | 17.94003 | -0.60826 | -6.70633 | 0.563067 | 0.666829 | 0 | 0 |
| 254 | sp\|P97496 | SMRC1_MO | 123 kDa | 0.996795 | 9 | 11% | P97496 | Smarcc1 | SWI/SNF co | 17.99677 | 17.91573 | 17.32152 | 18.20306 | 18.29381 | 18.10156 | -0.20275 | -1.00526 | 0.59976 | 17.97207 | -0.60133 | -6.7111 | 0.567167 | 0.671098 | 0 | 0 |
| 460 | sp\|Q9DBH5 | LMAN2_MO | 40 kDa | 0.998899 | 6 | 100% | Q9DBH5 | Lman2 | Vesicular ir | 20.93232 | 21.46101 | 22.26293 | 20.61199 | 20.83561 | 21.11186 | 0.281999 | -0.8437 | 1.407697 | 21.20262 | 0.597827 | -6.71311 | 0.569608 | 0.673398 | 0 | 0 |
| 247 | sp\|P62631 | EF1A2_MO | 50 kDa | 0.998875 | 9 | 22% | P62631 | Eef1a2 | Elongation | 19.64305 | 19.51025 | 19.98301 | 20.70349 | 20.27363 | 18.75752 | 0.309476 | -0.93312 | 1.552074 | 19.81183 | 0.594357 | -6.71535 | 0.571794 | 0.675044 | 0 | 0 |
| 734 | sp\|P56959 | FUS_MOUS | 53 kDa | 0.998377 | 4 | 100% | P56959 | Fus | RNA-bindin | 21.80268 | 21.18087 | 19.18505 | 17.89109 | 20.36085 | 20.49338 | 0.59441 | -1.79353 | 2.98235 | 20.15232 | 0.594037 | -6.71555 | 0.571996 | 0.675044 | 0 | 0 |
| 1177.2 | tr\|A0A075 | A0A075B5Y | 13 kDa | 0.998168 | 2 | 50% | A0A075B5Y | Ighv1-74 | Immunoglo | 19.81061 | 20.27385 | 21.1667 | 20.37539 | 19.63427 | 18.94972 | 0.337541 | -1.02403 | 1.699114 | 20.03509 | 0.591611 | -6.71711 | 0.573528 | 0.676264 | 0 | 0 |
| 1127 | sp\|Q920I9 | WDR7_MO | 163 kDa | 0.998654 | 2 | 100% | Q920I9 | Wdr7 | WD repeat- | 20.11111 | 17.67665 | 18.66057 | 21.04512 | 18.16841 | 19.84115 | -0.54095 | -2.73165 | 1.649764 | 19.2505 | -0.58928 | -6.7186 | 0.575004 | 0.677415 | 0 | 0 |
| 318 | sp\|Q00493 | CBPE_MOU | 53 kDa | 0.999646 | 7 | 100% | Q00493 | Cpe | Carboxype | 22.89779 | 24.25512 | 23.72315 | 22.69235 | 23.24539 | 21.80169 | 0.372389 | -1.13855 | 1.883326 | 23.10258 | 0.588167 | -6.7193 | 0.575706 | 0.677654 | 0 | 0 |
| 513 | sp\|Q9DCN2 | NB5R3_MO | 34 kDa | 0.999426 | 5 | 100% | Q9DCN2 | Cyb5r3 | NADH-cyto | 22.0716 | 21.98739 | 23.01042 | 22.38095 | 21.97085 | 22.04763 | 0.212301 | -0.66055 | 1.085149 | 22.24481 | 0.579838 | -6.72466 | 0.580904 | 0.68318 | 0 | 0 |
| 471 | sp\|P62137 | PP1A_MOU | 38 kDa | 0.997925 | 6 | 33% | P62137 | Ppp1ca | Serine/thre | 19.60489 | 18.91981 | 18.94128 | 17.75669 | 19.18296 | 20.03897 | 0.337886 | -1.05636 | 1.732128 | 19.0741 | 0.578339 | -6.72551 | 0.58195 | 0.683816 | 0 | 0 |
| 689 | sp\|P46471 | PRS7_MOU | 49 kDa | 0.999 | 4 | 100% | P46471 | Psmc2 | 26S protea | 20.829 | 21.20927 | 21.50021 | 20.87381 | 20.58285 | 21.41555 | -0.19553 | -1.01909 | 0.628041 | 21.06845 | -0.5651 | -6.73395 | 0.590203 | 0.69237 | 0 | 0 |
| 883 | sp\|Q9CU65 | ZMYM2_MO | 155 kDa | 0.998886 | 3 | 100% | Q9CU65 | Zmym2 | Zinc finger | 17.23808 | 18.42317 | 18.76093 | 18.4447 | 17.7189 | 17.64749 | -0.26582 | -1.3885 | 0.856856 | 18.03888 | -0.56505 | -6.73375 | 0.590455 | 0.69237 | 0 | 0 |
| 1154.2 | tr\|E9PW43 | E9PW43_M | 12 kDa | 0.998394 | 2 | 100% | E9PW43 | Gm10320 | Protein tra | 19.65571 | 20.12876 | 20.76477 | 20.23821 | 19.90625 | 20.58927 | -0.20984 | -1.0971 | 0.677429 | 20.21383 | -0.5638 | -6.73461 | 0.591171 | 0.69237 | 0 | 0 |
| 1071 | sp\|P11531 | DMD_MOU | 426 kDa | 0.999197 | 2 | 100% | P11531 | Dmd | Dystrophin | 20.4741 | 19.62792 | 19.83168 | 19.8011 | 20.29718 | 20.56238 | 0.203854 | -0.65835 | 1.06606 | 20.09906 | 0.563641 | -6.7347 | 0.591272 | 0.69237 | 0 | 0 |
| 126 | sp\|Q91YQ5 | RPN1_MOU | 69 kDa | 0.999763 | 13 | 100% | Q91YQ5 | Rpn1 | Dolichyl-di | 24.3914 | 24.68795 | 24.83312 | 23.73024 | 24.10211 | 24.29589 | 0.204186 | -0.66459 | 1.072964 | 24.34012 | 0.560289 | -6.73675 | 0.593431 | 0.694299 | 0 | 0 |
| 924 | tr\|A0A140 | A0A140T8 | 13 kDa | 0.998347 | 3 | 67% | A0A140T8 | Igkv8-24 | Immunoglo | 19.90754 | 20.00146 | 19.42872 | 19.12867 | 19.88892 | 20.88543 | -0.26346 | -1.40259 | 0.875674 | 19.87346 | -0.55193 | -6.74171 | 0.598916 | 0.700112 | 0 | 0 |
| 102 | tr\|G3X8Q1 | G3X8Q1_M | 243 kDa | 0.999642 | 15 | 100% | G3X8Q1 | Cabin1 | Calcineurin | 22.72226 | 24.03761 | 22.60114 | 22.75351 | 23.67907 | 22.9811 | -0.25658 | -1.38222 | 0.86906 | 23.12911 | -0.54397 | -6.74646 | 0.604091 | 0.70525 | 0 | 0 |
| 13 | sp\|E9Q7G0 | NUMA1_MO | 236 kDa | 0.999659 | 35 | 100% | E9Q7G0 | Numa1 | Nuclear mi | 24.72768 | 24.60314 | 25.0132 | 25.50108 | 24.84293 | 25.00648 | -0.17563 | -0.94514 | 0.593875 | 24.94909 | -0.54324 | -6.74712 | 0.604351 | 0.70525 | 0 | 0 |
| 930.1 | sp\|P60764 | RAC3_MOU | 21 kDa | 0.998224 | 3 | 100% | P60764 | Rac3 | Ras-related | 19.18671 | 19.40853 | 19.42759 | 19.44826 | 19.70052 | 18.98329 | 0.158244 | -0.54281 | 0.859299 | 19.35915 | 0.537251 | -6.75064 | 0.608261 | 0.709202 | 0 | 0 |
| 908 | sp\|Q9Z0P4 | PALM_MOU | 42 kDa | 0.998551 | 3 | 100% | Q9Z0P4 | Palm | Paralemmi | 21.56623 | 21.34025 | 21.07598 | 21.3167 | 21.40037 | 21.87418 | -0.16284 | -0.88662 | 0.560928 | 21.42895 | -0.53551 | -6.75165 | 0.609398 | 0.709917 | 0 | 0 |
| 96 | sp\|Q8VDP4 | CCAR2_MO | 103 kDa | 0.999847 | 15 | 100% | Q8VDP4 | Ccar2 | Cell cycle a | 24.48677 | 24.21227 | 23.42988 | 23.23853 | 24.5259 | 24.26878 | 0.240995 | -0.83517 | 1.317162 | 24.02702 | 0.534416 | -6.75208 | 0.610325 | 0.710387 | 0 | 0 |
| 1267 | sp\|Q80VD1 | FA98B_MO | 45 kDa | 0.996327 | 2 | 100% | Q80VD1 | Fam98b | Protein FAI | 17.64117 | 15.69858 | 19.23325 | 18.91385 | 18.33087 | 19.07164 | 0.507073 | -1.76674 | 2.780888 | 18.14823 | 0.532189 | -6.75337 | 0.611784 | 0.711455 | 0 | 0 |
| 1158 | sp\|P47964 | RL36_MOU | 12 kDa | 0.998336 | 2 | 100% | P47964 | Rpl36 | 60S ribosor | 19.08974 | 20.41369 | 19.74373 | 18.58569 | 18.98256 | 19.63001 | -0.27112 | -1.48992 | 0.947684 | 19.40757 | -0.53086 | -6.75415 | 0.612656 | 0.711455 | 0 | 0 |
| 1149 | sp\|Q5EG47 | AAPK1_MO | 64 kDa | 0.998487 | 2 | 100% | Q5EG47 | Prkaa1 | 5'-AMP-act | 18.92212 | 19.95578 | 19.12417 | 17.8412 | 18.57772 | 19.76345 | -0.31214 | -1.71789 | 1.093611 | 19.03074 | -0.5299 | -6.7547 | 0.613286 | 0.711455 | 0 | 0 |
| 1208 | tr\|A0A0J9 | A0A0J9YUD | 233 kDa | 0.997821 | 2 | 100% | A0A0J9YUD | Nup205 | Nucleopori | 18.11076 | 16.23971 | 17.24764 | 17.96186 | 17.88223 | 18.14149 | 0.29919 | -1.04845 | 1.64683 | 17.59728 | 0.529814 | -6.75475 | 0.613342 | 0.711455 | 0 | 0 |
| 381 | sp\|P16460 | ASSY_MOU | 47 kDa | 0.998214 | 7 | 100% | P16460 | Ass1 | Argininosu | 22.79178 | 22.7732 | 23.17322 | 23.0631 | 22.71076 | 22.34609 | 0.16446 | -0.58166 | 0.910582 | 22.80969 | 0.524627 | -6.75795 | 0.616547 | 0.714563 | 0 | 0 |
| 1120 | sp\|Q8K354 | CBR3_MOU | 31 kDa | 0.998731 | 2 | 50% | Q8K354 | Cbr3 | Carbonyl re | 20.20173 | 21.27588 | 20.73007 | 19.9929 | 19.79225 | 20.15657 | -0.23376 | -1.30283 | 0.835306 | 20.35823 | -0.52182 | -6.75934 | 0.618599 | 0.716328 | 0 | 0 |
| 834 | sp\|Q3THE2 | ML12B_MO | 20 kDa | 0.99929 | 3 | 100% | Q3THE2 | Myl12b | Myosin reg | 21.38875 | 21.27953 | 21.52952 | 22.23487 | 21.48497 | 20.02844 | 0.286799 | -1.02981 | 1.603411 | 21.32435 | 0.519841 | -6.76047 | 0.619907 | 0.716857 | 0 | 0 |
| 434 | sp\|P10649 | GSTM1_MO | 26 kDa | 0.999283 | 6 | 83% | P10649 | Gstm1 | Glutathion | 20.92802 | 21.39024 | 22.893 | 21.88715 | 21.39811 | 21.08552 | 0.285405 | -1.02559 | 1.596403 | 21.59701 | 0.519528 | -6.76065 | 0.620113 | 0.716857 | 0 | 0 |
| 572 | sp\|Q78JE5 | FBX22_MO | 44 kDa | 0.998299 | 5 | 100% | Q78JE5 | Fbxo22 | F-box only | 21.78488 | 20.53452 | 20.9012 | 22.62494 | 21.95091 | 22.44711 | -0.32319 | -1.81428 | 1.167897 | 21.70726 | -0.51726 | -6.76194 | 0.621614 | 0.717935 | 0 | 0 |
| 363 | sp\|Q9ERA6 | TFP11_MO | 96 kDa | 0.99909 | 7 | 100% | Q9ERA6 | Tfip11 | Tuftelin-int | 20.45983 | 18.80913 | 20.29682 | 21.20046 | 19.84115 | 19.64128 | 0.315644 | -1.14272 | 1.774008 | 20.04145 | 0.516515 | -6.76236 | 0.622104 | 0.717935 | 0 | 0 |
| 1247 | sp\|P63085 | MK01_MOU | 41 kDa | 0.99695 | 2 | 100% | P63085 | Mapk1 | Mitogen-ac | 18.798 | 18.84499 | 17.98721 | 18.19567 | 18.00167 | 16.90873 | 0.279164 | -1.01875 | 1.577081 | 18.12271 | 0.51329 | -6.76417 | 0.62424 | 0.719537 | 0 | 0 |
| 86 | sp\|P68372 | TBB4B_MO | 50 kDa | 0.998318 | 17 | 6% | P68372 | Tubb4b | Tubulin bet | 24.69166 | 24.36238 | 24.89946 | 24.5802 | 24.17673 | 24.36194 | 0.154445 | -0.5637 | 0.872587 | 24.51206 | 0.511874 | -6.76517 | 0.62498 | 0.719537 | 0 | 0 |
| 1091 | tr\|A0A140 | A0A140T8 | 13 kDa | 0.999019 | 2 | 100% | A0A140T8 | Igkv9-123 | Immunoglo | 18.58958 | 19.4859 | 19.83884 | 19.02601 | 18.52368 | 19.0933 | -0.21771 | -1.23355 | 0.798142 | 19.09288 | -0.51144 | -6.76521 | 0.625468 | 0.719537 | 0 | 0 |
| 443 | sp\|O35691 | PININ_MO | 82 kDa | 0.999181 | 6 | 100% | O35691 | Pnn | Pinin | 21.90102 | 20.93412 | 20.57801 | 21.31886 | 21.20792 | 20.81976 | 0.204738 | -0.75101 | 1.160491 | 21.12662 | 0.511216 | -6.76534 | 0.625615 | 0.719537 | 0 | 0 |
| 1113 | sp\|P99027 | RLA2_MOU | 12 kDa | 0.998821 | 2 | 100% | P99027 | Rplp2 | 60S acidic r | 18.91857 | 19.18302 | 18.99206 | 19.17815 | 19.04936 | 18.98392 | -0.12836 | -0.73089 | 0.474157 | 19.05085 | -0.50708 | -6.76784 | 0.628168 | 0.721583 | 0 | 0 |
| 964 | sp\|Q9CQ26 | STABP_MO | 49 kDa | 0.997689 | 3 | 100% | Q9CQ26 | Stambp | STAM-bind | 23.84574 | 24.21123 | 23.48265 | 23.34506 | 23.48379 | 23.75128 | -0.16513 | -0.94091 | 0.610646 | 23.68662 | -0.50664 | -6.76808 | 0.628458 | 0.721583 | 0 | 0 |
| 1276 | sp\|Q62465 | VAT1_MOU | 43 kDa | 0.995771 | 2 | 100% | Q62465 | Vat1 | Synaptic ve | 18.39812 | 19.02562 | 18.71781 | 18.04646 | 19.79436 | 21.041 | -0.40093 | -2.29546 | 1.493605 | 19.17056 | -0.50503 | -6.76878 | 0.629727 | 0.722328 | 0 | 0 |
| 226 | sp\|Q8BGC0 | HTSF1_MO | 86 kDa | 0.999777 | 9 | 100% | Q8BGC0 | Htatsf1 | HIV Tat-spe | 22.57438 | 22.12624 | 23.23762 | 22.74238 | 22.81128 | 23.19069 | 0.187993 | -0.70079 | 1.076773 | 22.78043 | 0.504243 | -6.76929 | 0.630172 | 0.722328 | 0 | 0 |
| 529 | sp\|P10605 | CATB_MOU | 37 kDa | 0.999267 | 5 | 100% | P10605 | Ctsb | Cathepsin E | 22.65574 | 22.34416 | 23.29181 | 22.49224 | 21.89326 | 22.40452 | 0.199963 | -0.74959 | 1.149513 | 22.51362 | 0.502553 | -6.77015 | 0.631376 | 0.723096 | 0 | 0 |

| | | | | | | | | | | | | | | | | | | | | | | | | | |
|---|---|---|---|---|---|---|---|---|---|---|---|---|---|---|---|---|---|---|---|---|---|---|---|---|---|
| 855 | sp\|Q9D0T1 | NH2L1_MO | 14 kDa | 0.999084 | 3 | 100% | Q9D0T1 | Snu13 | NHP2-like p | 20.79625 | 21.62489 | 20.97691 | 20.37932 | 21.00473 | 21.33955 | -0.18863 | -1.10265 | 0.725402 | 21.02028 | -0.49209 | -6.77591 | 0.638314 | 0.730181 | 0 | 0 |
| 260 | sp\|P60335 | PCBP1_MO | 37 kDa | 0.99974 | 8 | 62% | P60335 | Pcbp1 | Poly(rC)-bi | 23.66607 | 23.44202 | 23.89618 | 24.30418 | 23.86316 | 23.13573 | 0.181157 | -0.69741 | 1.059726 | 23.71789 | 0.491555 | -6.77622 | 0.638656 | 0.730181 | 0 | 0 |
| 865 | sp\|Q8BGH2 | SAM50_MO | 52 kDa | 0.999029 | 3 | 100% | Q8BGH2 | Samm50 | Sorting and | 21.00546 | 21.3021 | 20.98856 | 20.29583 | 21.17737 | 22.33767 | -0.25474 | -1.49312 | 0.983652 | 21.1845 | -0.49089 | -6.77651 | 0.639178 | 0.730181 | 0 | 0 |
| 828 | sp\|P62320 | SMD3_MOU | 14 kDa | 0.999359 | 3 | 100% | P62320 | Snrpd3 | Small nucle | 18.96889 | 18.69494 | 18.70098 | 18.22903 | 19.91639 | 19.88172 | 0.260191 | -1.01459 | 1.53497 | 19.06532 | 0.487089 | -6.77855 | 0.64173 | 0.731956 | 0 | 0 |
| 132 | sp\|Q9Z1X4 | ILF3_MOUS | 96 kDa | 0.999596 | 13 | 100% | Q9Z1X4 | Ilf3 | Interleukin | 23.28296 | 22.73677 | 22.81955 | 23.62603 | 23.39284 | 23.66521 | -0.17755 | -1.04695 | 0.691842 | 23.25389 | -0.48686 | -6.77874 | 0.641811 | 0.731956 | 0 | 0 |
| 342 | sp\|Q6R0H7 | GNAS1_MO | 122 kDa | 0.999386 | 7 | 86% | Q6R0H7 | Gnas | Guanine nu | 23.17557 | 23.6719 | 23.61081 | 23.30896 | 23.02393 | 23.26202 | -0.14419 | -0.85363 | 0.565236 | 23.3422 | -0.48377 | -6.7805 | 0.643776 | 0.73358 | 0 | 0 |
| 15 | sp\|P39053 | DYN1_MOU | 98 kDa | 0.999724 | 34 | 74% | P39053 | Dnm1 | Dynamin-1 | 26.61206 | 26.34164 | 25.67204 | 26.39923 | 26.76094 | 26.85276 | -0.18286 | -1.08944 | 0.723725 | 26.43978 | -0.48084 | -6.78194 | 0.645865 | 0.735343 | 0 | 0 |
| 1150 | sp\|O35218 | CPSF2_MO | 88 kDa | 0.998475 | 2 | 100% | O35218 | Cpsf2 | Cleavage ar | 19.18184 | 19.45775 | 19.45249 | 19.04598 | 19.69368 | 19.4092 | 0.138357 | -0.54962 | 0.826328 | 19.37349 | 0.478663 | -6.7832 | 0.647224 | 0.736272 | 0 | 0 |
| 1024 | tr\|A0A0B4. | A0A0B4J1I | 13 kDa | 0.999697 | 2 | 100% | A0A0B4J1I | Igkv16-104 | Immunoglo | 21.38847 | 21.58627 | 20.24428 | 21.89224 | 21.79185 | 20.68011 | -0.24467 | -1.47781 | 0.988456 | 21.26387 | -0.47351 | -6.78572 | 0.650892 | 0.738682 | 0 | 0 |
| 1025 | sp\|Q9D032 | SSBP3_MO | 40 kDa | 0.999644 | 2 | 50% | Q9D032 | Ssbp3 | Single-stran | 21.38847 | 21.58627 | 20.24428 | 21.89224 | 21.79185 | 20.68011 | -0.24467 | -1.47781 | 0.988456 | 21.26387 | -0.47351 | -6.78572 | 0.650892 | 0.738682 | 0 | 0 |
| 739 | sp\|Q569Z6 | TR150_MO | 108 kDa | 0.998309 | 4 | 100% | Q569Z6 | Thrap3 | Thyroid ho | 20.76209 | 19.6964 | 20.58644 | 22.44865 | 20.76132 | 18.67997 | 0.428276 | -1.73076 | 2.587308 | 20.48915 | 0.473386 | -6.78578 | 0.650977 | 0.738682 | 0 | 0 |
| 816 | sp\|P01648 | KV5AF_MO | 12 kDa | 0.999423 | 3 | 33% | P01648 | NA | Ig kappa ch | 21.41844 | 22.40012 | 21.30904 | 20.62245 | 20.90759 | 21.29694 | -0.22815 | -1.38337 | 0.927073 | 21.32577 | -0.47131 | -6.78687 | 0.652387 | 0.739662 | 0 | 0 |
| 966 | sp\|Q8JZQ2 | AFG32_MO | 90 kDa | 0.99763 | 3 | 100% | Q8JZQ2 | Afg3l2 | AFG3-like p | 19.27651 | 19.40219 | 20.08141 | 20.42771 | 19.80819 | 19.86487 | -0.17622 | -1.0855 | 0.733066 | 19.81015 | -0.46202 | -6.79171 | 0.658632 | 0.746089 | 0 | 0 |
| 333 | sp\|Q70FJ1 | AKAP9_MO | 436 kDa | 0.99948 | 7 | 100% | Q70FJ1 | Akap9 | A-kinase an | 17.06258 | 19.63767 | 18.42371 | 19.14454 | 20.46672 | 18.3516 | -0.3936 | -2.42958 | 1.642382 | 18.8478 | -0.46135 | -6.79198 | 0.659155 | 0.746089 | 0 | 0 |
| 1241.2 | sp\|Q3THW | H2AV_MOU | 14 kDa | 0.997171 | 2 | 50% | Q3THW5 | H2afv | Histone H2 | 19.58837 | 20.26592 | 21.09828 | 20.96858 | 19.73209 | 19.88424 | -0.23333 | -1.4457 | 0.979036 | 20.25625 | -0.45929 | -6.79303 | 0.660557 | 0.747052 | 0 | 0 |
| 292 | sp\|Q6PE01 | SNR40_MO | 39 kDa | 0.999016 | 8 | 100% | Q6PE01 | Snrnp40 | U5 small nu | 22.10031 | 21.63682 | 21.56255 | 22.63607 | 22.17995 | 20.90991 | 0.220006 | -0.92842 | 1.368431 | 21.8376 | 0.457176 | -6.7941 | 0.662002 | 0.748063 | 0 | 0 |
| 270 | sp\|P63101 | 1433Z_MO | 28 kDa | 0.999587 | 8 | 62% | P63101 | Ywhaz | 14-3-3 prot | 21.68044 | 21.91043 | 22.10508 | 22.03288 | 22.11662 | 21.57364 | 0.128397 | -0.5613 | 0.818094 | 21.90318 | 0.443098 | -6.80125 | 0.671484 | 0.758038 | 0 | 0 |
| 499 | sp\|P61982 | 1433G_MO | 28 kDa | 0.999552 | 5 | 40% | P61982 | Ywhag | 14-3-3 prot | 20.95297 | 21.07012 | 21.48384 | 22.38251 | 21.19212 | 20.79487 | -0.20619 | -1.3178 | 0.905408 | 21.31274 | -0.44267 | -6.80131 | 0.671948 | 0.758038 | 0 | 0 |
| 940 | sp\|P46660 | AINX_MOU | 55 kDa | 0.998031 | 3 | 100% | P46660 | Ina | Alpha-inter | 17.1509 | 16.13965 | 16.3956 | 18.83853 | 19.42203 | 19.41233 | -0.47399 | -3.04597 | 2.097989 | 17.89317 | -0.4398 | -6.80271 | 0.673923 | 0.759634 | 0 | 0 |
| 215 | sp\|Q6PIE5 | AT1A2_MO | 112 kDa | 0.999015 | 10 | 10% | Q6PIE5 | Atp1a2 | Sodium/po | 17.61301 | 18.61843 | 18.79411 | 17.05493 | 16.99542 | 18.52112 | -0.26398 | -1.73244 | 1.204482 | 17.93284 | -0.429 | -6.8079 | 0.681384 | 0.767406 | 0 | 0 |
| 882 | sp\|P01629 | KV2A4_MO | 12 kDa | 0.998891 | 3 | 33% | P01629 | NA | Ig kappa ch | 20.59663 | 20.97024 | 19.3243 | 20.08337 | 21.16943 | 20.70445 | -0.22256 | -1.48174 | 1.036609 | 20.47474 | -0.42181 | -6.81128 | 0.686368 | 0.772378 | 0 | 0 |
| 160 | sp\|O35219 | KCNH2_MO | 127 kDa | 0.999658 | 11 | 100% | O35219 | Kcnh2 | Potassium | 23.00372 | 24.68984 | 23.60599 | 23.95935 | 23.08892 | 21.91815 | -0.28957 | -1.96279 | 1.383647 | 23.37766 | -0.413 | -6.81536 | 0.692504 | 0.778078 | 0 | 0 |
| 1190 | sp\|Q99NB8 | UBQL4_MO | 64 kDa | 0.998054 | 2 | 50% | Q99NB8 | Ubqln4 | Ubiquilin-4 | 17.14193 | 18.76924 | 20.51699 | 20.42141 | 20.19039 | 19.82646 | -0.38927 | -2.63915 | 1.860615 | 19.47774 | -0.41289 | -6.81541 | 0.692582 | 0.778078 | 0 | 0 |
| 528 | sp\|P51655 | GPC4_MOU | 63 kDa | 0.999272 | 5 | 100% | P51655 | Gpc4 | Glypican-4 | 20.94028 | 21.04677 | 21.73373 | 20.69238 | 20.94758 | 22.54985 | -0.22247 | -1.51777 | 1.072824 | 21.31843 | -0.40988 | -6.81679 | 0.694682 | 0.779792 | 0 | 0 |
| 168 | tr\|A8DUK4 | A8DUK4_M | 16 kDa | 0.999378 | 11 | 27% | A8DUK4 | Hbb-bs | Beta-globin | 24.13773 | 24.42798 | 27.41344 | 27.18335 | 22.00268 | 23.74592 | -0.60114 | -4.13889 | 2.936618 | 24.81852 | -0.40551 | -6.81877 | 0.697746 | 0.782112 | 0 | 0 |
| 773 | sp\|P03977 | KV3A4_MO | 12 kDa | 0.995804 | 4 | 25% | P03977 | NA | Ig kappa ch | 19.13728 | 19.53145 | 20.08325 | 19.0562 | 18.47123 | 19.6619 | -0.18593 | -1.28075 | 0.90889 | 19.32355 | -0.40528 | -6.81887 | 0.697902 | 0.782112 | 0 | 0 |
| 779 | sp\|Q9CY27 | TECR_MOU | 36 kDa | 0.994712 | 4 | 100% | Q9CY27 | Tecr | Very-long-c | 20.79924 | 22.84498 | 21.00096 | 17.94505 | 20.66127 | 20.31157 | 0.453287 | -2.22633 | 3.132907 | 20.59385 | 0.403692 | -6.81958 | 0.699016 | 0.782693 | 0 | 0 |
| 873 | sp\|P35279 | RAB6A_MO | 24 kDa | 0.998972 | 3 | 67% | P35279 | Rab6a | Ras-related | 19.85947 | 18.92636 | 18.51753 | 18.52958 | 19.59504 | 19.90064 | 0.205154 | -1.01002 | 1.420331 | 19.22144 | 0.402894 | -6.81994 | 0.699575 | 0.782693 | 0 | 0 |
| 953 | sp\|Q9R190 | MTA2_MOU | 75 kDa | 0.997887 | 3 | 100% | Q9R190 | Mta2 | Metastasis- | 19.05368 | 20.55486 | 21.25122 | 20.84917 | 20.75787 | 20.35336 | -0.23154 | -1.61619 | 1.153108 | 20.47003 | -0.39906 | -6.82165 | 0.702266 | 0.784947 | 0 | 0 |
| 243 | sp\|Q9D361 | SNR48_MO | 39 kDa | 0.999104 | 9 | 100% | Q9D361 | Snrnp48 | U11/U12 sm | 23.8802 | 23.84941 | 22.73898 | 23.57623 | 24.06127 | 23.74127 | -0.16215 | -1.13354 | 0.809227 | 23.64123 | -0.39837 | -6.82195 | 0.702748 | 0.784947 | 0 | 0 |
| 1059 | sp\|P01845 | LAC3_MOU | 11 kDa | 0.999277 | 2 | 50% | P01845 | Iglc3 | Ig lambda-3 | 18.96789 | 18.93317 | 21.05773 | 21.01257 | 20.48814 | 19.71716 | 0.28362 | -1.42314 | 1.990376 | 20.02944 | 0.396566 | -6.82275 | 0.704018 | 0.785719 | 0 | 0 |
| 1213 | sp\|Q9Z2X8 | KEAP1_MO | 70 kDa | 0.997758 | 2 | 100% | Q9Z2X8 | Keap1 | Kelch-like E | 20.26591 | 19.91346 | 21.23667 | 21.64515 | 20.0402 | 20.6109 | -0.20891 | -1.49667 | 1.07885 | 20.61872 | -0.38715 | -6.82686 | 0.710655 | 0.792474 | 0 | 0 |
| 497 | sp\|P14211 | CALR_MOU | 48 kDa | 0.999571 | 5 | 100% | P14211 | Calr | Calreticulin | 23.05235 | 23.38093 | 23.94199 | 23.44173 | 23.01304 | 23.57657 | -0.13062 | -0.94454 | 0.683296 | 23.4011 | -0.38199 | -6.82919 | 0.71416 | 0.795728 | 0 | 0 |
| 214 | sp\|Q80X90 | FLNB_MOU | 278 kDa | 0.999039 | 10 | 90% | Q80X90 | Flnb | Filamin-B | 20.94778 | 21.06141 | 20.71561 | 20.61266 | 20.79619 | 20.45672 | 0.109596 | -0.57533 | 0.794524 | 20.76506 | 0.380846 | -6.82967 | 0.714966 | 0.795971 | 0 | 0 |
| 420 | sp\|P97868 | RBBP6_MO | 200 kDa | 0.99947 | 6 | 100% | P97868 | Rbbp6 | E3 ubiquiti | 17.20791 | 16.80985 | 17.42868 | 18.31085 | 18.05094 | 18.10283 | -0.17866 | -1.32144 | 0.96411 | 17.65184 | -0.3731 | -6.8328 | 0.7206 | 0.801587 | 0 | 0 |
| 205 | sp\|Q78PY7 | SND1_MOU | 102 kDa | 0.99934 | 10 | 100% | Q78PY7 | Snd1 | Staphyloco | 22.07838 | 22.82223 | 23.10942 | 22.07828 | 22.07989 | 22.82135 | -0.15139 | -1.12583 | 0.823043 | 22.49826 | -0.37077 | -6.83377 | 0.722256 | 0.80277 | 0 | 0 |
| 986 | sp\|Q9D2N4 | DTNA_MOU | 84 kDa | 0.997055 | 3 | 100% | Q9D2N4 | Dtna | Dystrobrev | 18.22444 | 18.24875 | 19.0052 | 18.84882 | 18.24069 | 17.96129 | 0.137156 | -0.75513 | 1.029442 | 18.42153 | 0.366442 | -6.83559 | 0.725282 | 0.805473 | 0 | 0 |
| 557 | sp\|P70699 | LYAG_MOU | 106 kDa | 0.998851 | 5 | 100% | P70699 | Gaa | Lysosomal | 21.47067 | 21.69806 | 21.40604 | 20.95197 | 21.27295 | 21.85822 | -0.11953 | -0.90422 | 0.665165 | 21.44299 | -0.36256 | -6.83724 | 0.727971 | 0.807797 | 0 | 0 |
| 1194 | sp\|P50518 | VATE1_MO | 26 kDa | 0.998024 | 2 | 100% | P50518 | Atp6v1e1 | V-type prot | 20.84811 | 19.15862 | 20.85129 | 23.03542 | 20.80973 | 21.2867 | -0.32387 | -2.4718 | 1.824067 | 20.99831 | -0.35983 | -6.83823 | 0.730051 | 0.809307 | 0 | 0 |
| 431 | sp\|P59279 | RAB2B_MO | 24 kDa | 0.999314 | 6 | 17% | P59279 | Rab2b | Ras-related | 18.13569 | 21.64292 | 18.60653 | 16.37499 | 18.13099 | 18.1757 | -0.44014 | -3.36456 | 2.484285 | 18.51114 | -0.35917 | -6.8385 | 0.730524 | 0.809307 | 0 | 0 |
| 1045 | sp\|O35609 | SCAM3_MO | 38 kDa | 0.999387 | 2 | 100% | O35609 | Scamp3 | Secretory c | 15.58442 | 16.91987 | 16.98412 | 13.91799 | 15.90329 | 16.71963 | 0.304782 | -1.73069 | 2.34025 | 16.00489 | 0.357336 | -6.83923 | 0.731834 | 0.809885 | 0 | 0 |
| 875 | sp\|Q9EQH2 | ERAP1_MO | 107 kDa | 0.998959 | 3 | 100% | Q9EQH2 | Erap1 | Endoplasm | 19.74412 | 18.17481 | 18.71048 | 20.06963 | 19.7232 | 19.33663 | 0.198907 | -1.1316 | 1.529412 | 19.29315 | 0.356766 | -6.83946 | 0.732241 | 0.809885 | 0 | 0 |
| 1243 | sp\|Q5RJ54 | ZSC26_MO | 54 kDa | 0.997096 | 2 | 100% | Q5RJ54 | Zscan26 | Zinc finger | 18.91613 | 19.3333 | 19.48355 | 19.29854 | 18.76011 | 18.86824 | -0.11343 | -0.87504 | 0.648183 | 19.10998 | -0.35448 | -6.84047 | 0.733745 | 0.810887 | 0 | 0 |
| 956 | sp\|P62835 | RAP1A_MO | 21 kDa | 0.997817 | 3 | 33% | P62835 | Rap1a | Ras-related | 19.19618 | 19.98074 | 18.56772 | 17.75969 | 18.35197 | 18.99809 | -0.20755 | -1.60825 | 1.193145 | 18.80906 | -0.35362 | -6.8407 | 0.734493 | 0.811052 | 0 | 0 |
| 1092 | sp\|P47754 | CAZA2_MO | 33 kDa | 0.999018 | 2 | 100% | P47754 | Capza2 | F-actin-cap | 20.14143 | 19.07365 | 19.61129 | 20.4151 | 19.4512 | 19.25248 | 0.154229 | -0.88928 | 1.197737 | 19.65753 | 0.352712 | -6.84106 | 0.735142 | 0.811109 | 0 | 0 |
| 946 | sp\|O88291 | ZN326_MO | 65 kDa | 0.997936 | 3 | 100% | O88291 | Znf326 | DBIRD com | 17.16737 | 16.93323 | 17.25425 | 17.47573 | 18.24135 | 17.82308 | 0.143644 | -0.84174 | 1.129031 | 17.4825 | 0.347883 | -6.84295 | 0.738604 | 0.814266 | 0 | 0 |
| 639 | sp\|P50247 | SAHH_MOU | 48 kDa | 0.999404 | 4 | 100% | P50247 | Ahcy | Adenosylho | 22.11378 | 21.10536 | 22.00446 | 21.51085 | 21.09017 | 23.20427 | -0.20402 | -1.65421 | 1.246164 | 21.83815 | -0.33574 | -6.84759 | 0.747335 | 0.823222 | 0 | 0 |
| 857 | sp\|P83870 | PHF5A_MO | 12 kDa | 0.999075 | 3 | 100% | P83870 | Phf5a | PHD finger- | 22.11358 | 22.3509 | 22.38025 | 21.79036 | 22.19039 | 22.25601 | 0.095651 | -0.58634 | 0.777642 | 22.18025 | 0.33382 | -6.84841 | 0.748599 | 0.823945 | 0 | 0 |
| 1160 | sp\|Q91WD | NDUS2_MO | 53 kDa | 0.998328 | 2 | 100% | Q91WD5 | Ndufs2 | NADH dehy | 17.99742 | 17.66516 | 18.6592 | 19.17022 | 17.58113 | 17.89728 | -0.16497 | -1.35618 | 1.026234 | 18.16173 | -0.3305 | -6.84955 | 0.751119 | 0.826048 | 0 | 0 |
| 1231 | sp\|Q9Z321 | TOP3B_MO | 97 kDa | 0.997513 | 2 | 100% | Q9Z321 | Top3b | DNA topois | 20.15416 | 20.45004 | 20.52617 | 20.19876 | 19.85661 | 20.17994 | -0.09726 | -0.8005 | 0.60598 | 20.22761 | -0.32918 | -6.85013 | 0.751948 | 0.826289 | 0 | 0 |
| 1129 | sp\|Q9R0P5 | DEST_MOU | 19 kDa | 0.998633 | 2 | 100% | Q9R0P5 | Dstn | Destrin | 21.1694 | 20.56723 | 20.72556 | 20.27905 | 21.07219 | 21.70453 | 0.13878 | -0.88359 | 1.161147 | 20.91966 | 0.323946 | -6.85195 | 0.75586 | 0.829803 | 0 | 0 |
| 959 | sp\|Q6S5C2 | GNPTG_MO | 34 kDa | 0.99775 | 3 | 100% | Q6S5C2 | Gnptg | N-acetylglu | 21.66111 | 20.92573 | 21.93925 | 22.57333 | 21.90516 | 22.47586 | -0.15646 | -1.3116 | 0.998675 | 21.91341 | -0.32324 | -6.8522 | 0.756369 | 0.829803 | 0 | 0 |
| 889 | sp\|Q921X9 | PDIA5_MO | 59 kDa | 0.998796 | 3 | 100% | Q921X9 | Pdia5 | Protein dis | 17.52168 | 17.37297 | 14.89525 | 17.32488 | 18.33935 | 16.87286 | -0.27148 | -2.29558 | 1.752624 | 17.0545 | -0.32008 | -6.85335 | 0.758664 | 0.831648 | 0 | 0 |
| 369 | sp\|P35700 | PRDX1_MO | 22 kDa | 0.999 | 7 | 86% | P35700 | Prdx1 | Peroxiredo | 24.56835 | 24.25957 | 24.40672 | 24.52826 | 24.31127 | 24.24868 | 0.083278 | -0.54172 | 0.708274 | 24.38714 | 0.31714 | -6.85448 | 0.760676 | 0.83318 | 0 | 0 |
| 122 | sp\|Q9ERD7 | TBB3_MOU | 50 kDa | 0.99869 | 14 | 14% | Q9ERD7 | Tubb3 | Tubulin bet | 20.68687 | 20.29832 | 19.98722 | 20.4519 | 20.1946 | 20.39613 | -0.09255 | -0.79572 | 0.610618 | 20.33584 | -0.31327 | -6.85585 | 0.763487 | 0.835584 | 0 | 0 |
| 130 | sp\|P08113 | ENPL_MOU | 92 kDa | 0.999636 | 13 | 85% | P08113 | Hsp90b1 | Endoplasm | 23.36985 | 23.98179 | 24.60512 | 24.07457 | 23.4087 | 23.70587 | -0.12618 | -1.09585 | 0.843482 | 23.85765 | -0.31055 | -6.85671 | 0.765584 | 0.836705 | 0 | 0 |
| 1240 | sp\|Q9DC28 | KC1D_MOU | 47 kDa | 0.997228 | 2 | 100% | Q9DC28 | Csnk1d | Casein kina | 19.15325 | 19.769 | 19.64491 | 17.68718 | 19.50703 | 20.23264 | 0.205455 | -1.37451 | 1.785418 | 19.33234 | 0.310327 | -6.85679 | 0.765745 | 0.836705 | 0 | 0 |
| 1012 | sp\|Q8R317 | UBQL1_MO | 62 kDa | 0.994329 | 3 | 33% | Q8R317 | Ubqln1 | Ubiquilin-1 | 25.34156 | 20.75504 | 16.76076 | 21.16008 | 18.56516 | 20.62432 | -0.62399 | -5.48539 | 4.237417 | 20.53449 | -0.30631 | -6.85818 | 0.768668 | 0.839222 | 0 | 0 |
| 803 | tr\|A0A075 | A0A075B5I | 11 kDa | 0.9995 | 3 | 33% | A0A075B5I | Igkv3-7 | Immunoglo | 18.92192 | 18.35212 | 20.93875 | 19.58157 | 19.46144 | 20.71307 | 0.225121 | -1.55147 | 2.001712 | 19.66148 | 0.302398 | -6.85952 | 0.771523 | 0.841661 | 0 | 0 |
| 830 | sp\|P58021 | TM9S2_MO | 75 kDa | 0.99935 | 3 | 100% | P58021 | Tm9sf2 | Transmemb | 22.90288 | 23.10182 | 22.57837 | 22.57899 | 22.64582 | 22.7035 | -0.08575 | -0.76369 | 0.592195 | 22.7519 | -0.30105 | -6.86005 | 0.772397 | 0.841938 | 0 | 0 |
| 268 | sp\|Q8BFR5 | EFTU_MOU | 50 kDa | 0.999599 | 8 | 100% | Q8BFR5 | Tufm | Elongation | 22.86184 | 22.51441 | 22.94794 | 22.51967 | 22.49548 | 23.62095 | -0.11659 | -1.04688 | 0.813707 | 22.82671 | -0.29908 | -6.86064 | 0.773948 | 0.84295 | 0 | 0 |
| 1215 | sp\|P97434 | MPRIP_MO | 116 kDa | 0.997743 | 2 | 100% | P97434 | Mprip | Myosin pho | 16.82105 | 15.70851 | 14.69956 | 16.22087 | 15.9545 | 16.0343 | -0.16285 | -1.46885 | 1.143143 | 15.90646 | -0.29758 | -6.86114 | 0.77504 | 0.843462 | 0 | 0 |
| 1107 | sp\|Q9ESX4 | NO40_MOU | 27 kDa | 0.998887 | 2 | 100% | Q9ESX4 | Zcchc17 | Nucleolar p | 21.05791 | 20.94184 | 20.89246 | 21.95751 | 21.29829 | 20.69501 | -0.11524 | -1.06091 | 0.830434 | 21.1405 | -0.2908 | -6.86338 | 0.779999 | 0.847658 | 0 | 0 |
| 555 | sp\|Q9QXK3 | COPG2_MO | 98 kDa | 0.998882 | 5 | 60% | Q9QXK3 | Copg2 | Coatomer s | 19.67779 | 19.60972 | 19.09376 | 18.19907 | 19.55987 | 21.14586 | -0.20774 | -1.91371 | 1.498231 | 19.54768 | -0.2906 | -6.86345 | 0.780145 | 0.847658 | 0 | 0 |
| 522 | sp\|Q9D662 | SC23B_MO | 86 kDa | 0.999327 | 5 | 100% | Q9D662 | Sec23b | Protein tra | 20.96271 | 21.19347 | 20.92166 | 20.48468 | 20.53657 | 21.01095 | -0.08939 | -0.83523 | 0.656451 | 20.85167 | -0.28526 | -6.86525 | 0.783958 | 0.850835 | 0 | 0 |
| 527 | sp\|P60122 | RUVB1_MO | 50 kDa | 0.999281 | 5 | 100% | P60122 | Ruvbl1 | RuvB-like 1 | 21.3495 | 21.68897 | 21.58598 | 21.03236 | 21.17249 | 21.6511 | -0.08815 | -0.82516 | 0.648856 | 21.4134 | -0.28468 | -6.86543 | 0.784385 | 0.850835 | 0 | 0 |
| 1082 | sp\|P27601 | GNA13_MO | 44 kDa | 0.999114 | 2 | 100% | P27601 | Gna13 | Guanine nu | 21.78672 | 21.72878 | 21.39984 | 21.17224 | 21.90926 | 21.91978 | 0.091671 | -0.67685 | 0.860191 | 21.65277 | 0.283909 | -6.86568 | 0.784952 | 0.850835 | 0 | 0 |
| 365 | sp\|P80318 | TCPG_MOU | 61 kDa | 0.999039 | 7 | 100% | P80318 | Cct3 | T-complex | 22.33462 | 22.96928 | 22.87415 | 23.08905 | 24.36377 | 23.99901 | -0.1616 | -1.55849 | 1.235299 | 23.27165 | -0.27607 | -6.86808 | 0.790817 | 0.856508 | 0 | 0 |
| 246 | sp\|P17710 | HXK1_MOU | 108 kDa | 0.99889 | 9 | 100% | P17710 | Hk1 | Hexokinase | 21.72752 | 22.48938 | 22.72211 | 22.1523 | 22.16765 | 22.25295 | -0.09245 | -0.90109 | 0.716193 | 22.25198 | -0.27212 | -6.86936 | 0.793631 | 0.858785 | 0 | 0 |
| 496 | sp\|Q8BX02 | KANK2_MO | 90 kDa | 0.999593 | 5 | 100% | Q8BX02 | Kank2 | KN motif ar | 20.18583 | 20.02936 | 19.595 | 20.42098 | 20.04671 | 19.64596 | -0.08958 | -0.87533 | 0.696164 | 19.98731 | -0.27136 | -6.86959 | 0.794186 | 0.858785 | 0 | 0 |
| 61 | sp\|Q8CIE6 | COPA_MOU | 138 kDa | 0.999781 | 18 | 100% | Q8CIE6 | Copa | Coatomer s | 24.14364 | 23.87813 | 24.11844 | 24.39352 | 24.21512 | 23.9872 | 0.07278 | -0.58208 | 0.727642 | 24.12267 | 0.264523 | -6.87165 | 0.799235 | 0.863556 | 0 | 0 |
| 31 | sp\|P48678 | LMNA_MOU | 74 kDa | 0.999722 | 25 | 96% | P48678 | Lmna | Prelamin-A | 25.54546 | 25.69273 | 25.57772 | 25.66907 | 25.52795 | 25.4869 | -0.06586 | -0.66479 | 0.533071 | 25.5833 | -0.26172 | -6.87248 | 0.801305 | 0.864492 | 0 | 0 |
| 120 | sp\|P58252 | EF2_MOUS | 95 kDa | 0.999506 | 14 | 93% | P58252 | Eef2 | Elongation | 23.5185 | 23.43513 | 23.48735 | 23.49313 | 23.23377 | 23.51272 | -0.06712 | -0.67774 | 0.543501 | 23.44677 | -0.26163 | -6.87251 | 0.801376 | 0.864492 | 0 | 0 |
| 1232 | sp\|Q9JKN1 | ZNT7_MOU | 42 kDa | 0.997459 | 2 | 100% | Q9JKN1 | Slc30a7 | Zinc transp | 19.52019 | 20.74202 | 20.70684 | 20.88422 | 20.84311 | 18.95848 | 0.16181 | -1.33666 | 1.66028 | 20.27581 | 0.257697 | -6.8736 | 0.804379 | 0.867041 | 0 | 0 |
| 618 | sp\|Q7TQI3 | OTUB1_MO | 31 kDa | 0.999546 | 4 | 100% | Q7TQI3 | Otub1 | Ubiquitin t | 19.81449 | 20.59816 | 21.02587 | 20.40909 | 20.44303 | 20.55783 | -0.0939 | -0.96887 | 0.781084 | 20.47475 | -0.25582 | -6.87416 | 0.805728 | 0.867524 | 0 | 0 |
| 1154.1 | sp\|Q9CQS8 | SC61B_MO | 10 kDa | 0.998394 | 2 | 100% | Q9CQS8 | Sec61b | Protein tra | 18.18731 | 20.39654 | 17.13762 | 18.7577 | 22.24006 | 19.39209 | -0.32711 | -3.38408 | 2.72985 | 19.35189 | -0.25536 | -6.87427 | 0.806107 | 0.867524 | 0 | 0 |
| 764 | sp\|P33173 | KIF1A_MO | 192 kDa | 0.997228 | 4 | 100% | P33173 | Kif1a | Kinesin-like | 20.97796 | 21.15958 | 21.30214 | 20.87773 | 21.2574 | 21.29183 | 0.069455 | -0.58482 | 0.72373 | 21.14444 | 0.252664 | -6.8751 | 0.808014 | 0.868887 | 0 | 0 |
| 899.1 | sp\|P27048 | RSMB_MOU | 24 kDa | 0.998705 | 3 | 100% | P27048 | Snrpb | Small nucle | 21.20571 | 21.45506 | 21.00193 | 21.21995 | 21.09475 | 20.8366 | -0.06974 | -0.75092 | 0.611446 | 21.13567 | -0.24368 | -6.87761 | 0.814688 | 0.874768 | 0 | 0 |
| 805 | sp\|P08249 | MDHM_MO | 36 kDa | 0.999491 | 3 | 100% | P08249 | Mdh2 | Malate deh | 19.93401 | 19.91285 | 20.47652 | 20.18746 | 19.62211 | 19.69039 | 0.080649 | -0.70747 | 0.868768 | 19.97055 | 0.243562 | -6.87764 | 0.814773 | 0.874768 | 0 | 0 |
| 366 | sp\|Q9DC48 | PRP17_MO | 65 kDa | 0.999017 | 7 | 100% | Q9DC48 | Cdc40 | Pre-mRNA- | 20.73486 | 19.61313 | 19.34994 | 20.16581 | 21.23264 | 21.10303 | 0.145156 | -1.29068 | 1.580995 | 20.36657 | 0.241257 | -6.87822 | 0.816574 | 0.876009 | 0 | 0 |
| 435 | sp\|Q8CFQ3 | AQR_MOUS | 170 kDa | 0.999277 | 6 | 100% | Q8CFQ3 | Aqr | RNA helicas | 20.99036 | 20.89076 | 20.84026 | 21.96435 | 21.60283 | 20.87386 | -0.0985 | -1.08188 | 0.884874 | 21.19373 | -0.23904 | -6.87882 | 0.818221 | 0.876744 | 0 | 0 |
| 52 | sp\|A2AQ25 | SKT_MOUS | 213 kDa | 0.999096 | 20 | 100% | A2AQ25 | Skt | Sickle tail p | 23.56177 | 23.07745 | 21.95647 | 22.7819 | 24.79154 | 25.0798 | -0.20979 | -2.30809 | 1.888513 | 23.54149 | -0.2386 | -6.87894 | 0.818553 | 0.876744 | 0 | 0 |
| 175 | sp\|P02089 | HBB2_MOU | 16 kDa | 0.999276 | 11 | 45% | P02089 | Hbb-b2 | Hemoglobi | 26.2066 | 26.5462 | 26.29839 | 25.89583 | 26.11172 | 26.38165 | -0.06899 | -0.76249 | 0.624503 | 26.24007 | -0.23679 | -6.87948 | 0.819813 | 0.8774 | 0 | 0 |
| 757 | sp\|Q8BL97 | SRSF7_MO | 31 kDa | 0.997544 | 4 | 100% | Q8BL97 | Srsf7 | Serine/argi | 20.2282 | 20.0828 | 20.98585 | 20.85514 | 20.07208 | 20.08307 | 0.088374 | -0.82505 | 1.001801 | 20.38452 | 0.230704 | -6.88105 | 0.824408 | 0.880985 | 0 | 0 |
| 1183 | tr\|A0A0B4. | A0A0B4J1J | 13 kDa | 0.998116 | 2 | 50% | A0A0B4J1J | Igkv5-43 | Immunoglo | 19.94503 | 19.28247 | 20.14324 | 20.41841 | 20.11446 | 20.24619 | 0.085223 | -0.79573 | 0.966176 | 20.02497 | 0.230621 | -6.88107 | 0.824461 | 0.880985 | 0 | 0 |
| 298 | sp\|Q9JKR6 | HYOU1_MO | 111 kDa | 0.998595 | 8 | 100% | Q9JKR6 | Hyou1 | Hypoxia up | 22.10976 | 22.8192 | 22.38509 | 21.87861 | 22.07534 | 22.10396 | -0.07719 | -0.88006 | 0.725676 | 22.22866 | -0.22884 | -6.88156 | 0.825741 | 0.881657 | 0 | 0 |
| 949 | sp\|Q6ZPZ3 | ZC3H4_MO | 141 kDa | 0.997914 | 3 | 100% | Q6ZPZ3 | Zc3h4 | Zinc finger | 18.39262 | 19.21084 | 19.6217 | 18.60171 | 18.30569 | 18.80158 | -0.09804 | -1.12901 | 0.932924 | 18.82236 | -0.22694 | -6.882 | 0.827239 | 0.882561 | 0 | 0 |
| 476 | sp\|Q91ZW | SMCA5_MO | 122 kDa | 0.997308 | 6 | 100% | Q91ZW3 | Smarca5 | SWI/SNF-re | 19.72855 | 19.11031 | 19.66381 | 20.07715 | 19.55415 | 19.99079 | -0.07725 | -0.89662 | 0.742122 | 19.68746 | -0.22441 | -6.88269 | 0.82905 | 0.883798 | 0 | 0 |
| 261 | sp\|Q8BFZ3 | ACTBL_MO | 42 kDa | 0.999727 | 8 | 12% | Q8BFZ3 | Actbl2 | Beta-actin- | 25.80617 | 25.79862 | 26.06189 | 25.90643 | 25.84889 | 25.8415 | 0.0568 | -0.54988 | 0.663479 | 25.87725 | 0.222838 | -6.88309 | 0.830224 | 0.884355 | 0 | 0 |
| 820 | sp\|Q8BG81 | PDIP3_MO | 46 kDa | 0.9994 | 3 | 100% | Q8BG81 | Poldip3 | Polymerase | 22.7588 | 22.21366 | 21.68589 | 22.03713 | 23.06807 | 22.95655 | 0.101808 | -0.99634 | 1.199958 | 22.45335 | 0.221243 | -6.88345 | 0.831496 | 0.885015 | 0 | 0 |
| 645 | sp\|O70480 | VAMP4_MO | 16 kDa | 0.999362 | 4 | 100% | O70480 | Vamp4 | Vesicle-ass | 21.3392 | 21.90636 | 20.96409 | 21.4667 | 23.09459 | 22.42019 | -0.13179 | -1.56853 | 1.304944 | 21.86519 | -0.21891 | -6.88403 | 0.833243 | 0.885492 | 0 | 0 |
| 540 | sp\|Q9DC69 | NDUA9_MO | 43 kDa | 0.999153 | 5 | 100% | Q9DC69 | Ndufa9 | NADH dehy | 20.95439 | 20.74596 | 21.06156 | 20.8071 | 20.58324 | 21.24013 | -0.06467 | -0.76815 | 0.638813 | 20.89873 | -0.21879 | -6.8841 | 0.833251 | 0.885492 | 0 | 0 |
| 1251.1 | sp\|Q8C181 | MBNL2_MO | 40 kDa | 0.996815 | 2 | 100% | Q8C181 | Mbnl2 | Muscleblin | 19.75808 | 18.58845 | 16.69542 | 17.3453 | 17.72092 | 17.73037 | 0.170101 | -1.71509 | 2.055293 | 17.97309 | 0.215329 | -6.88491 | 0.83592 | 0.887634 | 0 | 0 |
| 265 | sp\|Q9D0I9 | SYRC_MOU | 76 kDa | 0.999616 | 8 | 100% | Q9D0I9 | Rars | Arginine--t | 22.86625 | 22.93493 | 22.67837 | 23.16642 | 23.12159 | 22.35769 | 0.069057 | -0.70049 | 0.838606 | 22.85421 | 0.213586 | -6.88537 | 0.837149 | 0.888242 | 0 | 0 |
| 646 | sp\|Q61171 | PRDX2_MO | 22 kDa | 0.999353 | 4 | 100% | Q61171 | Prdx2 | Peroxiredo | 22.13007 | 21.76013 | 22.65564 | 22.11627 | 22.02764 | 22.70884 | 0.076037 | -0.77682 | 0.928892 | 22.2331 | 0.212494 | -6.88561 | 0.838006 | 0.888457 | 0 | 0 |
| 50 | sp\|E9Q4N7 | ARI1B_MO | 238 kDa | 0.999548 | 20 | 90% | E9Q4N7 | Arid1b | AT-rich inte | 22.91238 | 23.11168 | 22.11225 | 23.12616 | 23.51563 | 22.5675 | -0.08836 | -1.09648 | 0.919763 | 22.89093 | -0.20916 | -6.88639 | 0.840539 | 0.889935 | 0 | 0 |
| 693 | tr\|A0A140 | A0A140T8 | 11 kDa | 0.998986 | 4 | 50% | A0A140T8 | Igkv8-19 | Immunoglo | 20.85234 | 22.0368 | 22.94648 | 22.69191 | 20.61774 | 20.21752 | -0.17655 | -2.19316 | 1.840056 | 21.56047 | -0.20893 | -6.88644 | 0.840713 | 0.889935 | 0 | 0 |
| 941 | sp\|Q9D0F3 | LMAN1_MO | 58 kDa | 0.998022 | 3 | 100% | Q9D0F3 | Lman1 | Protein ER | 17.6207 | 13.26178 | 18.26963 | 21.05394 | 17.06045 | 19.81639 | -0.39378 | -4.9364 | 4.148852 | 17.84715 | -0.20687 | -6.88693 | 0.842261 | 0.89083 | 0 | 0 |
| 288 | tr\|Q8VHM5 | Q8VHM5_M | 71 kDa | 0.999188 | 8 | 100% | Q8VHM5 | Hnrnpr | Heterogene | 23.30095 | 21.90537 | 22.48716 | 23.44388 | 23.13989 | 23.92765 | -0.1163 | -1.46325 | 1.230652 | 23.03415 | -0.20605 | -6.88712 | 0.842872 | 0.89083 | 0 | 0 |
| 1256 | sp\|Q9D1M | FKB11_MO | 22 kDa | 0.996662 | 2 | 100% | Q9D1M7 | Fkbp11 | Peptidyl-pr | 18.64991 | 20.05106 | 19.02298 | 19.2587 | 21.36887 | 20.16775 | -0.14525 | -1.90022 | 1.609723 | 19.75321 | -0.19751 | -6.88907 | 0.849288 | 0.896912 | 0 | 0 |
| 1074 | sp\|Q9CXW | RL11_MOU | 20 kDa | 0.999168 | 2 | 100% | Q9CXW4 | Rpl11 | 60S riboso | 19.20832 | 18.50612 | 20.30768 | 20.06658 | 19.34228 | 20.64254 | -0.11899 | -1.57933 | 1.341359 | 19.67892 | -0.19444 | -6.88975 | 0.851595 | 0.898648 | 0 | 0 |
| 735 | sp\|P12815 | PDCD6_MO | 22 kDa | 0.998357 | 4 | 100% | P12815 | Pdcd6 | Programme | 22.06108 | 22.01147 | 22.57604 | 22.06772 | 21.65422 | 22.02628 | 0.061958 | -0.70127 | 0.825185 | 22.06614 | 0.193216 | -6.89005 | 0.852448 | 0.898849 | 0 | 0 |
| 884 | sp\|Q8BPQ7 | SGSM1_MO | 123 kDa | 0.99885 | 3 | 100% | Q8BPQ7 | Sgsm1 | Small G pro | 16.85289 | 16.09935 | 14.58429 | 16.0955 | 16.90943 | 16.52604 | -0.12476 | -1.6751 | 1.425584 | 16.17792 | -0.19204 | -6.89027 | 0.853405 | 0.899159 | 0 | 0 |
| 1039 | sp\|Q9CZU6 | CISY_MOUS | 52 kDa | 0.999426 | 2 | 100% | Q9CZU6 | Cs | Citrate synt | 18.71935 | 19.08206 | 19.36727 | 18.73393 | 18.53251 | 18.61436 | 0.062927 | -0.72662 | 0.852477 | 18.84158 | 0.189696 | -6.89081 | 0.8551 | 0.900245 | 0 | 0 |
| 219 | sp\|E9Q784 | ZC3HD_MO | 204 kDa | 0.997856 | 10 | 100% | E9Q784 | Zc3h13 | Zinc finger | 21.42312 | 21.17777 | 20.74086 | 21.73705 | 22.0924 | 21.10612 | 0.078479 | -0.91777 | 1.074729 | 21.37955 | 0.18799 | -6.89114 | 0.856453 | 0.90097 | 0 | 0 |
| 1157 | tr\|D3YZZ5 | D3YZZ5_M | 25 kDa | 0.99837 | 2 | 100% | D3YZZ5 | Tmed7 | Transmemb | 19.16088 | 19.91201 | 20.42441 | 19.96342 | 19.02837 | 19.00143 | -0.08773 | -1.24737 | 1.071907 | 19.58175 | -0.18054 | -6.8927 | 0.862066 | 0.905386 | 0 | 0 |
| 276 | sp\|Q5SQX6 | CYFP2_MO | 146 kDa | 0.999473 | 8 | 38% | Q5SQX6 | Cyfip2 | Cytoplasmi | 20.79859 | 21.2019 | 19.66486 | 20.00923 | 20.57171 | 20.07719 | -0.08439 | -1.20517 | 1.036393 | 20.38725 | -0.17968 | -6.89287 | 0.862717 | 0.905386 | 0 | 0 |
| 291 | sp\|P48722 | HS74L_MO | 94 kDa | 0.99902 | 8 | 88% | P48722 | Hspa4l | Heat shock | 22.65517 | 22.68177 | 22.58948 | 22.39452 | 22.79719 | 23.12671 | -0.05372 | -0.76647 | 0.659036 | 22.70747 | -0.17938 | -6.89296 | 0.862884 | 0.905386 | 0 | 0 |
| 746 | sp\|Q01405 | SC23A_MO | 86 kDa | 0.997866 | 4 | 100% | Q01405 | Sec23a | Protein tra | 20.46179 | 20.9745 | 21.20348 | 21.70314 | 22.79839 | 22.13014 | -0.1147 | -1.64534 | 1.415938 | 21.54524 | -0.17883 | -6.89304 | 0.863357 | 0.905386 | 0 | 0 |
| 554 | sp\|A2AIV2 | VIR_MOUSI | 201 kDa | 0.998885 | 5 | 100% | A2AIV2 | Virma | Protein viri | 18.65207 | 20.73825 | 18.01073 | 16.58382 | 19.20584 | 19.08094 | -0.17813 | -2.56635 | 2.210094 | 18.71194 | -0.178 | -6.89321 | 0.863989 | 0.905386 | 0 | 0 |
| 622 | sp\|P09411 | PGK1_MOU | 45 kDa | 0.999493 | 4 | 100% | P09411 | Pgk1 | Phosphogly | 21.27753 | 21.32949 | 21.82823 | 21.64172 | 20.9402 | 20.88428 | 0.063492 | -0.79112 | 0.918099 | 21.31691 | 0.17707 | -6.89341 | 0.864658 | 0.905387 | 0 | 0 |
| 1101 | sp\|Q8VHR5 | P66B_MOU | 65 kDa | 0.998958 | 2 | 100% | Q8VHR5 | Gatad2b | Transcripti | 19.44319 | 20.15276 | 20.86102 | 20.26719 | 20.01486 | 20.1095 | -0.07013 | -1.03132 | 0.891062 | 20.14142 | -0.17411 | -6.89399 | 0.866922 | 0.907057 | 0 | 0 |
| 230 | sp\|Q3UJB9 | EDC4_MOU | 152 kDa | 0.999697 | 9 | 100% | Q3UJB9 | Edc4 | Enhancer o | 23.14227 | 22.16338 | 22.08852 | 22.31154 | 21.60148 | 22.14372 | 0.071208 | -0.95458 | 1.096991 | 22.24182 | 0.165662 | -6.89561 | 0.873312 | 0.913039 | 0 | 0 |
| 519 | sp\|F8VPU2 | FARP1_MO | 119 kDa | 0.999348 | 5 | 100% | F8VPU2 | Farp1 | FERM, ARH | 21.29879 | 21.31905 | 21.6422 | 20.97503 | 21.28754 | 22.11016 | -0.05857 | -0.93514 | 0.817999 | 21.4388 | -0.15929 | -6.8968 | 0.878112 | 0.917338 | 0 | 0 |
| 545 | sp\|Q61316 | HSP74_MO | 94 kDa | 0.999066 | 5 | 80% | Q61316 | Hspa4 | Heat shock | 21.483 | 22.08773 | 22.02101 | 21.20066 | 21.08727 | 21.48338 | -0.06016 | -0.96553 | 0.845201 | 21.56051 | -0.15842 | -6.89696 | 0.878777 | 0.917338 | 0 | 0 |
| 1123 | sp\|Q69ZB0 | LRCC1_MO | 119 kDa | 0.998684 | 2 | 100% | Q69ZB0 | Lrrcc1 | Leucine-ric | 16.72955 | 15.2281 | 17.21865 | 17.44409 | 15.52095 | 16.48037 | 0.105532 | -1.50847 | 1.719536 | 16.43695 | 0.156038 | -6.89737 | 0.880602 | 0.918535 | 0 | 0 |
| 107 | sp\|P68033 | ACTC_MOU | 42 kDa | 0.999399 | 15 | 13% | P68033 | Actc1 | Actin, alph | 24.82825 | 24.95047 | 24.98502 | 25.03139 | 24.82405 | 24.7731 | -0.03921 | -0.6497 | 0.571273 | 24.89871 | -0.15288 | -6.89794 | 0.882938 | 0.920265 | 0 | 0 |
| 970 | sp\|Q07235 | GDN_MOU | 44 kDa | 0.997591 | 3 | 100% | Q07235 | Serpine2 | Glia-derive | 19.53715 | 19.08583 | 20.09918 | 19.9864 | 19.2582 | 19.65896 | 0.054449 | -0.8297 | 0.938595 | 19.60429 | 0.146812 | -6.89896 | 0.887581 | 0.924246 | 0 | 0 |
| 65 | tr\|E9Q0S6 | E9Q0S6_M | 201 kDa | 0.999567 | 18 | 94% | E9Q0S6 | Tns1 | Tensin 1 | 21.95895 | 21.65872 | 20.96663 | 21.55416 | 22.5955 | 22.10686 | 0.067109 | -1.02887 | 1.163089 | 21.8068 | 0.146127 | -6.89907 | 0.888121 | 0.924246 | 0 | 0 |
| 864 | sp\|Q9DCT2 | NDUS3_MO | 30 kDa | 0.999038 | 3 | 100% | Q9DCT2 | Ndufs3 | NADH dehy | 20.36371 | 20.89951 | 20.82638 | 20.18019 | 20.21143 | 20.45857 | -0.04558 | -0.82317 | 0.732004 | 20.48996 | -0.13952 | -6.90016 | 0.893089 | 0.926951 | 0 | 0 |
| 1111 | sp\|Q9CWJ9 | PUR9_MOU | 64 kDa | 0.998832 | 2 | 100% | Q9CWJ9 | Atic | Bifunctiona | 20.32476 | 20.59322 | 20.67541 | 20.10776 | 20.16423 | 20.34214 | 0.040429 | -0.65645 | 0.737307 | 20.36792 | 0.138081 | -6.90038 | 0.894187 | 0.926951 | 0 | 0 |
| 798.2 | sp\|Q91VM | RMXL1_MO | 42 kDa | 0.999544 | 3 | 100% | Q91VM5 | Rbmxl1 | RNA bindin | 23.32623 | 22.79697 | 22.02473 | 23.00488 | 23.438 | 22.81121 | 0.058631 | -0.96831 | 1.085569 | 22.90034 | 0.136249 | -6.90065 | 0.895629 | 0.926951 | 0 | 0 |
| 853 | sp\|D3YVE8 | S35G2_MO | 46 kDa | 0.999109 | 3 | 100% | D3YVE8 | Slc35g2 | Solute carr | 21.33574 | 22.58111 | 21.10183 | 20.10264 | 21.23871 | 20.73697 | 0.085185 | -1.40964 | 1.580013 | 21.18283 | 0.135995 | -6.90069 | 0.895822 | 0.926951 | 0 | 0 |
| 1033.2 | sp\|Q9JJW6 | ALRF2_MO | 24 kDa | 0.999499 | 2 | 100% | Q9JJW6 | Alyref2 | Aly/REF exp | 20.45353 | 20.23386 | 20.84213 | 19.7014 | 20.15506 | 21.32573 | 0.063242 | -1.05543 | 1.181916 | 20.45195 | 0.134913 | -6.90086 | 0.896645 | 0.926951 | 0 | 0 |
| 550 | sp\|P47856 | GFPT1_MO | 79 kDa | 0.99894 | 5 | 100% | P47856 | Gfpt1 | Glutamine- | 19.69151 | 20.05103 | 20.15577 | 19.7354 | 19.58693 | 19.76446 | -0.03889 | -0.73163 | 0.653838 | 19.83085 | -0.13363 | -6.90107 | 0.89757 | 0.926951 | 0 | 0 |
| 916.9 | sp\|C0HKE9 | H2A1P_MO | 14 kDa | 0.998436 | 3 | 33% | C0HKE9 | Hist1h2ap | Histone H2 | 20.40214 | 18.79705 | 18.81104 | 21.67383 | 19.56569 | 18.65782 | -0.11661 | -2.20082 | 1.967598 | 19.65126 | -0.13352 | -6.90107 | 0.897704 | 0.926951 | 0 | 0 |
| 492 | sp\|P26369 | U2AF2_MO | 54 kDa | 0.999676 | 5 | 100% | P26369 | U2af2 | Splicing fac | 21.56485 | 22.2022 | 21.52638 | 21.4041 | 22.07582 | 21.68901 | -0.04275 | -0.83702 | 0.751513 | 21.74373 | -0.12811 | -6.90189 | 0.901778 | 0.926951 | 0 | 0 |
| 375 | sp\|Q91VR2 | ATPG_MOU | 33 kDa | 0.998696 | 7 | 100% | Q91VR2 | Atp5f1c | ATP syntha | 24.74527 | 24.59051 | 24.32571 | 25.39775 | 24.57984 | 23.8282 | -0.05521 | -1.09854 | 0.988111 | 24.57788 | -0.12629 | -6.90213 | 0.903206 | 0.926951 | 0 | 0 |
| 841 | sp\|Q99020 | ROAA_MOU | 31 kDa | 0.999237 | 3 | 67% | Q99020 | Hnrnpab | Heterogene | 21.44213 | 22.40325 | 19.3087 | 19.55064 | 22.88734 | 22.09729 | -0.13767 | -2.74438 | 2.46904 | 21.28156 | -0.12604 | -6.90217 | 0.903401 | 0.926951 | 0 | 0 |
| 867.2 | tr\|A0A140 | A0A140T8 | 13 kDa | 0.999017 | 3 | 100% | A0A140T8 | Igkv6-20 | Immunoglo | 20.59284 | 20.42443 | 20.78981 | 20.59627 | 20.6225 | 20.88238 | 0.034023 | -0.61099 | 0.67904 | 20.65137 | 0.125546 | -6.90226 | 0.903732 | 0.926951 | 0 | 0 |
| 950 | sp\|Q8C052 | MAP1S_MO | 103 kDa | 0.997908 | 3 | 100% | Q8C052 | Map1s | Microtubu | 20.56109 | 20.87354 | 20.92946 | 20.85954 | 21.03078 | 20.6912 | 0.032355 | -0.61833 | 0.683041 | 20.82427 | 0.118349 | -6.90325 | 0.90922 | 0.926951 | 0 | 0 |
| 601 | sp\|P17225 | PTBP1_MO | 56 kDa | 0.999706 | 4 | 100% | P17225 | Ptbp1 | Polypyrimi | 22.61313 | 21.57616 | 21.20093 | 21.83852 | 22.70467 | 22.90967 | 0.064793 | -1.24326 | 1.372841 | 22.14051 | 0.118209 | -6.90326 | 0.909368 | 0.926951 | 0 | 0 |

| | | | | | | | | | | | | | | | | | | | | | | | | | |
|---|---|---|---|---|---|---|---|---|---|---|---|---|---|---|---|---|---|---|---|---|---|---|---|---|---|
| 1198 | sp\|P67984 | RL22_MOU | 15 kDa | 0.997965 | 2 | 100% | P67984 | Rpl22 | 60S ribosor | 19.9585 | 20.06135 | 20.39028 | 20.94424 | 21.58985 | 20.75909 | 0.057984 | -1.12321 | 1.239183 | 20.61722 | 0.117149 | -6.9034 | 0.910177 | 0.926951 | 0 | 0 |
| 340 | sp\|Q8K310 | MATR3_MO | 95 kDa | 0.999425 | 7 | 100% | Q8K310 | Matr3 | Matrin-3 | 22.1656 | 21.47324 | 21.79343 | 22.14264 | 22.12795 | 22.59859 | -0.0425 | -0.90805 | 0.82305 | 22.05024 | -0.11705 | -6.90342 | 0.910234 | 0.926951 | 0 | 0 |
| 153 | sp\|Q8CI94 | PYGB_MOU | 97 kDa | 0.999191 | 12 | 100% | Q8CI94 | Pygb | Glycogen p | 23.31514 | 23.29045 | 23.23935 | 23.33649 | 23.28189 | 23.29564 | -0.02873 | -0.61712 | 0.55965 | 23.29316 | -0.11623 | -6.90353 | 0.910833 | 0.926951 | 0 | 0 |
| 1248 | sp\|O35593 | PSDE_MOU | 35 kDa | 0.99692 | 2 | 100% | O35593 | Psmd14 | 26S protea | 21.12769 | 21.27214 | 21.34948 | 21.16852 | 21.04555 | 21.16986 | -0.02927 | -0.64192 | 0.583385 | 21.18887 | -0.11371 | -6.90386 | 0.912763 | 0.926951 | 0 | 0 |
| 909 | sp\|Q9EQH3 | VPS35_MO | 92 kDa | 0.998549 | 3 | 100% | Q9EQH3 | Vps35 | Vacuolar pr | 20.51883 | 20.37394 | 20.09085 | 20.03742 | 19.4143 | 19.47568 | 0.045646 | -0.9159 | 1.007193 | 19.98517 | 0.113288 | -6.9039 | 0.913123 | 0.926951 | 0 | 0 |
| 910 | sp\|Q64704 | STX3_MOU | 33 kDa | 0.998547 | 3 | 100% | Q64704 | Stx3 | Syntaxin-3 | 20.51883 | 20.37394 | 20.09085 | 20.03742 | 19.4143 | 19.47568 | 0.045646 | -0.9159 | 1.007193 | 19.98517 | 0.113288 | -6.9039 | 0.913123 | 0.926951 | 0 | 0 |
| 911.1 | sp\|P01680 | KV4A1_MO | 14 kDa | 0.998536 | 3 | 67% | P01680 | NA | Ig kappa ch | 20.51883 | 20.37394 | 20.09085 | 20.03742 | 19.4143 | 19.47568 | 0.045646 | -0.9159 | 1.007193 | 19.98517 | 0.113288 | -6.9039 | 0.913123 | 0.926951 | 0 | 0 |
| 911.2 | tr\|A0A075 | A0A075B5 | 10 kDa | 0.998536 | 3 | 67% | A0A075B5 | Igkv4-58 | Immunoglo | 20.51883 | 20.37394 | 20.09085 | 20.03742 | 19.4143 | 19.47568 | 0.045646 | -0.9159 | 1.007193 | 19.98517 | 0.113288 | -6.9039 | 0.913123 | 0.926951 | 0 | 0 |
| 912 | sp\|P62082 | RS7_MOUS | 22 kDa | 0.998535 | 3 | 100% | P62082 | Rps7 | 40S ribosor | 20.51883 | 20.37394 | 20.09085 | 20.03742 | 19.4143 | 19.47568 | 0.045646 | -0.9159 | 1.007193 | 19.98517 | 0.113288 | -6.9039 | 0.913123 | 0.926951 | 0 | 0 |
| 913 | sp\|O70194 | EIF3D_MOU | 64 kDa | 0.998488 | 3 | 100% | O70194 | Eif3d | Eukaryotic | 20.51883 | 20.37394 | 20.09085 | 20.03742 | 19.4143 | 19.47568 | 0.045646 | -0.9159 | 1.007193 | 19.98517 | 0.113288 | -6.9039 | 0.913123 | 0.926951 | 0 | 0 |
| 914 | sp\|O70401 | TSN6_MOU | 27 kDa | 0.998475 | 3 | 100% | O70401 | Tspan6 | Tetraspanir | 20.51883 | 20.37394 | 20.09085 | 20.03742 | 19.4143 | 19.47568 | 0.045646 | -0.9159 | 1.007193 | 19.98517 | 0.113288 | -6.9039 | 0.913123 | 0.926951 | 0 | 0 |
| 915 | sp\|Q8C854 | MYEF2_MO | 63 kDa | 0.998453 | 3 | 100% | Q8C854 | Myef2 | Myelin expr | 20.51883 | 20.37394 | 20.09085 | 20.03742 | 19.4143 | 19.47568 | 0.045646 | -0.9159 | 1.007193 | 19.98517 | 0.113288 | -6.9039 | 0.913123 | 0.926951 | 0 | 0 |
| 916.1 | sp\|C0HKE1 | H2A1B_MO | 14 kDa | 0.998436 | 3 | 33% | C0HKE1 | Hist1h2ab | Histone H2 | 20.51883 | 20.37394 | 20.09085 | 20.03742 | 19.4143 | 19.47568 | 0.045646 | -0.9159 | 1.007193 | 19.98517 | 0.113288 | -6.9039 | 0.913123 | 0.926951 | 0 | 0 |
| 916.2 | sp\|C0HKE2 | H2A1C_MO | 14 kDa | 0.998436 | 3 | 33% | C0HKE2 | Hist1h2ac | Histone H2 | 20.51883 | 20.37394 | 20.09085 | 20.03742 | 19.4143 | 19.47568 | 0.045646 | -0.9159 | 1.007193 | 19.98517 | 0.113288 | -6.9039 | 0.913123 | 0.926951 | 0 | 0 |
| 916.3 | sp\|C0HKE3 | H2A1D_MO | 14 kDa | 0.998436 | 3 | 33% | C0HKE3 | Hist1h2ad | Histone H2 | 20.51883 | 20.37394 | 20.09085 | 20.03742 | 19.4143 | 19.47568 | 0.045646 | -0.9159 | 1.007193 | 19.98517 | 0.113288 | -6.9039 | 0.913123 | 0.926951 | 0 | 0 |
| 916.4 | sp\|C0HKE4 | H2A1E_MO | 14 kDa | 0.998436 | 3 | 33% | C0HKE4 | Hist1h2ae | Histone H2 | 20.51883 | 20.37394 | 20.09085 | 20.03742 | 19.4143 | 19.47568 | 0.045646 | -0.9159 | 1.007193 | 19.98517 | 0.113288 | -6.9039 | 0.913123 | 0.926951 | 0 | 0 |
| 916.5 | sp\|C0HKE5 | H2A1G_MO | 14 kDa | 0.998436 | 3 | 33% | C0HKE5 | Hist1h2ag | Histone H2 | 20.51883 | 20.37394 | 20.09085 | 20.03742 | 19.4143 | 19.47568 | 0.045646 | -0.9159 | 1.007193 | 19.98517 | 0.113288 | -6.9039 | 0.913123 | 0.926951 | 0 | 0 |
| 916.6 | sp\|C0HKE6 | H2A1I_MO | 14 kDa | 0.998436 | 3 | 33% | C0HKE6 | Hist1h2ai | Histone H2 | 20.51883 | 20.37394 | 20.09085 | 20.03742 | 19.4143 | 19.47568 | 0.045646 | -0.9159 | 1.007193 | 19.98517 | 0.113288 | -6.9039 | 0.913123 | 0.926951 | 0 | 0 |
| 916.7 | sp\|C0HKE7 | H2A1N_MO | 14 kDa | 0.998436 | 3 | 33% | C0HKE7 | Hist1h2an | Histone H2 | 20.51883 | 20.37394 | 20.09085 | 20.03742 | 19.4143 | 19.47568 | 0.045646 | -0.9159 | 1.007193 | 19.98517 | 0.113288 | -6.9039 | 0.913123 | 0.926951 | 0 | 0 |
| 155 | sp\|P17426 | AP2A1_MO | 108 kDa | 0.998576 | 12 | 75% | P17426 | Ap2a1 | AP-2 compl | 22.93295 | 22.47298 | 22.6362 | 22.01365 | 21.94942 | 22.90079 | 0.043716 | -0.87866 | 0.966088 | 22.48433 | 0.113079 | -6.90393 | 0.913279 | 0.926951 | 0 | 0 |
| 1151 | sp\|Q3ULD5 | MCCB_MOU | 61 kDa | 0.998431 | 2 | 100% | Q3ULD5 | Mccc2 | Methylcrot | 21.55498 | 22.09253 | 22.86161 | 22.41022 | 21.89145 | 21.93979 | -0.04483 | -1.00743 | 0.917765 | 22.1251 | -0.11115 | -6.90418 | 0.914755 | 0.927755 | 0 | 0 |
| 1073 | sp\|Q8BYR5 | CAPS2_MO | 148 kDa | 0.999194 | 2 | 100% | Q8BYR5 | Cadps2 | Calcium-de | 17.76841 | 18.70472 | 17.65086 | 17.46606 | 17.77377 | 16.86584 | 0.052138 | -1.10626 | 1.210536 | 17.70494 | 0.10741 | -6.90464 | 0.91761 | 0.929955 | 0 | 0 |
| 885 | sp\|Q9D0F9 | PGM1_MO | 61 kDa | 0.998848 | 3 | 100% | Q9D0F9 | Pgm1 | Phosphoglu | 18.09085 | 15.92527 | 18.3743 | 19.68268 | 18.78224 | 19.33598 | 0.101155 | -2.2151 | 2.417412 | 18.36522 | 0.10422 | -6.90502 | 0.920047 | 0.931728 | 0 | 0 |
| 1116 | sp\|Q7M6Y3 | PICAL_MOU | 72 kDa | 0.998771 | 2 | 100% | Q7M6Y3 | Picalm | Phosphatid | 20.13643 | 19.17647 | 21.37808 | 20.471 | 19.70109 | 21.37022 | 0.065972 | -1.53927 | 1.671217 | 20.37222 | 0.098077 | -6.90573 | 0.924741 | 0.935783 | 0 | 0 |
| 745 | tr\|A0A140T | A0A140T8M | 13 kDa | 0.997869 | 4 | 25% | A0A140T8M | Igkv12-44 | Immunoglo | 19.83415 | 19.31605 | 19.20688 | 18.51155 | 19.08573 | 20.16884 | 0.043438 | -1.09724 | 1.184112 | 19.35387 | 0.090877 | -6.9065 | 0.930248 | 0.940654 | 0 | 0 |
| 976 | sp\|P63323 | RS12_MOU | 15 kDa | 0.997401 | 3 | 100% | P63323 | Rps12 | 40S ribosor | 19.49148 | 19.14987 | 16.10126 | 18.24643 | 19.09579 | 17.5199 | -0.07589 | -2.31001 | 2.158234 | 18.26745 | -0.08106 | -6.90746 | 0.937762 | 0.947544 | 0 | 0 |
| 251 | sp\|Q7TMB8 | CYFP1_MO | 145 kDa | 0.998228 | 9 | 44% | Q7TMB8 | Cyfip1 | Cytoplasmi | 21.39762 | 20.75995 | 19.23383 | 20.25249 | 20.90249 | 20.3841 | 0.045797 | -1.3236 | 1.415198 | 20.48841 | 0.079811 | -6.90758 | 0.93872 | 0.947729 | 0 | 0 |
| 223 | sp\|Q8R1B4 | EIF3C_MOU | 106 kDa | 0.99983 | 9 | 100% | Q8R1B4 | Eif3c | Eukaryotic | 21.64697 | 21.43479 | 22.32186 | 21.69111 | 21.79317 | 22.73647 | -0.03346 | -1.04412 | 0.97721 | 21.9374 | -0.079 | -6.90765 | 0.939342 | 0.947729 | 0 | 0 |
| 1186 | tr\|A0A075 | A0A075B5V | 11 kDa | 0.998102 | 2 | 50% | A0A075B5V | Ighv8-6 | Immunoglo | 19.61395 | 20.03651 | 18.98044 | 18.48211 | 20.13247 | 20.3272 | -0.03966 | -1.39342 | 1.314111 | 19.59545 | -0.06991 | -6.90842 | 0.946309 | 0.952636 | 0 | 0 |
| 39 | sp\|P11679 | K2C8_MOU | 55 kDa | 0.999223 | 23 | 78% | P11679 | Krt8 | Keratin, typ | 24.76468 | 24.61796 | 24.25378 | 24.79579 | 25.13428 | 24.80522 | -0.02208 | -0.7797 | 0.735551 | 24.72862 | -0.06935 | -6.90847 | 0.946711 | 0.952636 | 0 | 0 |
| 446 | sp\|P62301 | RS13_MOU | 17 kDa | 0.999096 | 6 | 100% | P62301 | Rps13 | 40S ribosor | 23.85364 | 23.37366 | 24.21973 | 23.1666 | 23.2309 | 24.87424 | -0.03674 | -1.30458 | 1.231096 | 23.78646 | -0.06916 | -6.90848 | 0.94688 | 0.952636 | 0 | 0 |
| 139 | sp\|Q6PIC6 | AT1A3_MO | 112 kDa | 0.999076 | 13 | 31% | Q6PIC6 | Atp1a3 | Sodium/po | 20.23796 | 20.39139 | 21.50874 | 21.785 | 20.56224 | 20.24407 | -0.03717 | -1.32307 | 1.248727 | 20.78823 | -0.06898 | -6.90849 | 0.947016 | 0.952636 | 0 | 0 |
| 502 | sp\|O08784 | TCOF_MOU | 135 kDa | 0.999531 | 5 | 100% | O08784 | Tcof1 | Treacle pro | 20.91253 | 20.42715 | 21.2026 | 22.96105 | 23.16259 | 21.7215 | 0.056006 | -1.90819 | 2.020203 | 21.73123 | 0.068046 | -6.90857 | 0.947736 | 0.952654 | 0 | 0 |
| 461 | sp\|Q9DC16 | ERGI1_MO | 33 kDa | 0.99886 | 6 | 100% | Q9DC16 | Ergic1 | Endoplasm | 21.9127 | 21.74401 | 22.29723 | 21.89614 | 21.43066 | 22.0419 | -0.01382 | -0.77446 | 0.746813 | 21.88711 | -0.04325 | -6.91014 | 0.966747 | 0.970325 | 0 | 0 |
| 564 | sp\|P30415 | NKTR_MOU | 163 kDa | 0.998614 | 5 | 100% | P30415 | Nktr | NK-tumor r | 18.89173 | 18.71571 | 18.58952 | 19.10442 | 19.91397 | 19.63225 | -0.01905 | -1.07998 | 1.041882 | 19.14127 | -0.04285 | -6.91016 | 0.967073 | 0.970325 | 0 | 0 |
| 4 | sp\|Q8VDD5 | MYH9_MOU | 226 kDa | 0.999708 | 54 | 78% | Q8VDD5 | Myh9 | Myosin-9 | 25.63779 | 25.28698 | 25.19296 | 25.61598 | 25.52883 | 25.49176 | -0.01171 | -0.67088 | 0.647447 | 25.45905 | -0.0423 | -6.91018 | 0.967477 | 0.970325 | 0 | 0 |
| 1192 | sp\|Q921G7 | ETFD_MOU | 68 kDa | 0.998029 | 2 | 100% | Q921G7 | Etfdh | Electron tra | 16.08342 | 16.00759 | 16.94603 | 16.22071 | 16.29947 | 17.15236 | -0.01725 | -1.01128 | 0.976783 | 16.4516 | -0.04141 | -6.91022 | 0.968178 | 0.970325 | 0 | 0 |
| 1182 | sp\|P12656 | TSHB_MOU | 15 kDa | 0.998122 | 2 | 100% | P12656 | Tshb | Thyrotropi | 18.9127 | 18.78192 | 19.13725 | 18.57023 | 18.65812 | 19.32253 | 0.01113 | -0.74749 | 0.769754 | 18.89713 | 0.034919 | -6.91051 | 0.97315 | 0.974588 | 0 | 0 |
| 925 | sp\|P40142 | TKT_MOUS | 68 kDa | 0.998322 | 3 | 100% | P40142 | Tkt | Transketola | 20.65241 | 20.75414 | 20.7934 | 20.09826 | 19.95131 | 20.52276 | 0.00732 | -0.82558 | 0.840215 | 20.46204 | 0.020918 | -6.91095 | 0.983913 | 0.98464 | 0 | 0 |
| 1260 | sp\|Q9D023 | MPC2_MOU | 14 kDa | 0.99645 | 2 | 100% | Q9D023 | Mpc2 | Mitochond | 20.59037 | 20.82323 | 21.61562 | 22.10251 | 21.26301 | 20.56059 | -0.00578 | -1.19682 | 1.185272 | 21.15922 | -0.01157 | -6.91112 | 0.991103 | 0.991103 | 0 | 0 |
| 1130 | sp\|P60766 | CDC42_MO | 21 kDa | 0.998611 | 2 | 100% | P60766 | Cdc42 | Cell divisio | NA | NA | NA | NA | 19.23984 | NA | NA | NA | NA | 19.23984 | NA | NA | NA | NA | NA | NA |
| 1228 | sp\|P11031 | TCP4_MOU | 14 kDa | 0.99755 | 2 | 100% | P11031 | Sub1 | Activated R | 14.74753 | NA | NA | NA | 13.88403 | NA | NA | NA | NA | 14.31578 | NA | NA | NA | NA | NA | NA |
| 1258 | sp\|P48774 | GSTM5_MO | 27 kDa | 0.996561 | 2 | 50% | P48774 | Gstm5 | Glutathion | 17.06121 | NA | 17.08049 | NA | 16.39204 | NA | NA | NA | NA | 16.84458 | NA | NA | NA | NA | NA | NA |
| 1292 | sp\|Q9WUA | PFKAP_MO | 85 kDa | 0.994644 | 2 | 100% | Q9WUA3 | Pfkp | ATP-dependent 6-phosphofructokinase, platelet type | | | | | | | | | | | | | | | | |
| 1293 | sp\|Q99JX7 | NXF1_MOU | 70 kDa | 0.994523 | 2 | 100% | Q99JX7 | Nxf1 | Nuclear RNA export factor 1 | | | | | | | | | | | | | | | | |
| 1294.1 | sp\|P83741 | WNK1_MO | 251 kDa | 0.994185 | 2 | 50% | P83741 | Wnk1 | Serine/threonine-protein kinase WNK1 | | | | | | | | | | | | | | | | |
| 1294.2 | sp\|Q80UE6 | WNK4_MO | 132 kDa | 0.994185 | 2 | 50% | Q80UE6 | Wnk4 | Serine/threonine-protein kinase WNK4 | | | | | | | | | | | | | | | | |
| 1295.1 | sp\|Q497I4 | KRT35_MO | 51 kDa | 0.99394 | 2 | 50% | Q497I4 | Krt35 | Keratin, type I cuticular Ha5 | | | | | | | | | | | | | | | | |
| 1295.2 | sp\|Q61765 | K1H1_MOU | 47 kDa | 0.99394 | 2 | 50% | Q61765 | Krt31 | Keratin, type I cuticular Ha1 | | | | | | | | | | | | | | | | |
| 1295.3 | sp\|Q61897 | KT33B_MO | 46 kDa | 0.99394 | 2 | 50% | Q61897 | Krt33b | Keratin, type I cuticular Ha3-II | | | | | | | | | | | | | | | | |
| 1296 | tr\|Q68FG2 | Q68FG2_M | 271 kDa | 0.993774 | 2 | 50% | Q68FG2 | Sptbn2 | Spectrin beta chain | | | | | | | | | | | | | | | | |
| 1297 | sp\|Q8CI43 | MYL6B_MO | 23 kDa | 0.993631 | 2 | 50% | Q8CI43 | Myl6b | Myosin light chain 6B | | | | | | | | | | | | | | | | |
| 1298 | sp\|Q05793 | PGBM_MO | 398 kDa | 0.993589 | 2 | 100% | Q05793 | Hspg2 | Basement membrane-specific heparan sulfate proteoglycan core protein | | | | | | | | | | | | | | | | |
| 1299 | sp\|Q8C006 | TRI35_MOU | 57 kDa | 0.993579 | 2 | 100% | Q8C006 | Trim35 | Tripartite motif-containing protein 35 | | | | | | | | | | | | | | | | |
| 1300 | sp\|Q3TWW | SRSF6_MOU | 39 kDa | 0.993577 | 2 | 50% | Q3TWW8 | Srsf6 | Serine/arginine-rich splicing factor 6 | | | | | | | | | | | | | | | | |
| 1301 | sp\|A1L317 | K1C24_MO | 54 kDa | 0.993541 | 2 | 50% | A1L317 | Krt24 | Keratin, type I cytoskeletal 24 | | | | | | | | | | | | | | | | |
| 1302 | sp\|Q9JMH6 | TRXR1_MO | 67 kDa | 0.993513 | 2 | 100% | Q9JMH6 | Txnrd1 | Thioredoxin reductase 1, cytoplasmic | | | | | | | | | | | | | | | | |
| 1303 | sp\|Q91W4 | KLC3_MOU | 56 kDa | 0.993453 | 2 | 100% | Q91W40 | Klc3 | Kinesin light chain 3 | | | | | | | | | | | | | | | | |
| 1304.1 | sp\|P63271 | SPT4A_MO | 13 kDa | 0.993287 | 2 | 100% | P63271 | Supt4h1a | Transcription elongation factor SPT4-A | | | | | | | | | | | | | | | | |
| 1304.2 | sp\|Q9Z199 | SPT4B_MO | 13 kDa | 0.993287 | 2 | 100% | Q9Z199 | Supt4h1b | Transcription elongation factor SPT4-B | | | | | | | | | | | | | | | | |
| 1305 | sp\|Q5HZG4 | TAF3_MOU | 105 kDa | 0.993207 | 2 | 100% | Q5HZG4 | Taf3 | Transcription initiation factor TFIID subunit 3 | | | | | | | | | | | | | | | | |
| 1306 | sp\|Q9QUM | PSA6_MOU | 27 kDa | 0.99316 | 2 | 100% | Q9QUM9 | Psma6 | Proteasome subunit alpha type-6 | | | | | | | | | | | | | | | | |

### Supplementary Table 2

| Protein Annotation | | | | | | | | | | Log2 Cyclic Loess Normalized Intensities | | | | | | Musashi_vs_IgG | | | | | | | | | |
|---|---|---|---|---|---|---|---|---|---|---|---|---|---|---|---|---|---|---|---|---|---|---|---|---|---|
| id | rotein Nam | ession Num | lecular Wei | ein Group S | ied Peptide | Exclusivity | UniProt ID | Gene name | Description | E1_MSI12 | E1_IgG | E2_MSI12 | E2_IgG | E3_MSI12 | E3_IgG | logFC | CI.L | CI.R | AveExpr | t | B | P.Value | adj.P.Val | sig.PVal | sig.FDR |
| 677 | sp\|Q80WC | UBN2_MOU | 142 kDa | 0.999085 | 4 | 1 | Q80WC1 | Ubn2 | Ubinuclein- | 19.17213 | 13.26706 | 17.38307 | 5.625675 | 21.08303 | 14.01973 | 8.241925 | 2.886949 | 13.5969 | 15.09178 | 3.673007 | -2.73929 | 0.008574 | 0.036911 | 1 | 1 |
| 778 | sp\|P01811 | HVM41_MC | 13 kDa | 0.995278 | 4 | 0.25 | P01811 | NA | Ig heavy ch | 21.20257 | 14.84614 | 21.92793 | 15.22002 | 20.71727 | 15.79102 | 5.996865 | 4.988373 | 7.005358 | 18.28416 | 14.19065 | 5.325797 | 3.03E-06 | 0.000427 | 1 | 1 |
| 305 | sp\|Q60749 | KHDR1_MC | 48 kDa | 0.999885 | 7 | 0.86 | Q60749 | Khdrbs1 | KH domain- | 26.58346 | 21.36758 | 25.47485 | 20.85161 | 27.02601 | 20.11857 | 5.582188 | 4.353376 | 6.811 | 23.57035 | 10.84101 | 3.721701 | 1.72E-05 | 0.001058 | 1 | 1 |
| 1061 | sp\|O09108 | LSHB_MOU | 15 kDa | 0.999248 | 2 | 1 | O09108 | Lhb | Lutropin su | 19.02823 | NA | 18.45906 | 13.25611 | 19.67979 | 13.71208 | 5.571598 | 4.430488 | 6.712707 | 16.82705 | 12.10273 | 3.290413 | 2.8E-05 | 0.001518 | 1 | 1 |
| 1015.2 | tr\|E9Q035 | E9Q035_M | 108 kDa | 0.99986 | 2 | 1 | E9Q035 | Gm20425 | Predicted ge | 24.19942 | 18.47307 | 24.71601 | 18.60751 | 23.98793 | 19.52478 | 5.432666 | 4.499976 | 6.365356 | 21.58479 | 13.90037 | 5.207377 | 3.46E-06 | 0.000427 | 1 | 1 |
| 611 | sp\|Q9CYZ8 | SSBP2_MO | 38 kDa | 0.999605 | 4 | 0.75 | Q9CYZ8 | Ssbp2 | Single-stran | 23.22381 | 17.48517 | 23.58941 | 19.2692 | 22.92989 | 18.70953 | 4.759738 | 3.573528 | 5.945949 | 20.86784 | 9.575731 | 2.949857 | 3.77E-05 | 0.001704 | 1 | 1 |
| 1 | sp\|Q68FD5 | CLH1_MOU | 192 kDa | 0.999927 | 86 | 1 | Q68FD5 | Cltc | Clathrin hea | 33.45827 | 28.44516 | 32.44402 | 27.97304 | 33.37165 | 28.92852 | 4.642404 | 3.66433 | 5.620478 | 30.77011 | 11.32718 | 3.990771 | 1.3E-05 | 0.000976 | 1 | 1 |
| 89 | sp\|P60879 | SNP25_MO | 23 kDa | 0.999799 | 16 | 0.94 | P60879 | Snap25 | Synaptosor | 28.13807 | 22.63727 | 27.60588 | 23.42141 | 27.93553 | 23.73367 | 4.629044 | 3.744928 | 5.513159 | 25.57864 | 12.48178 | 4.611365 | 6.68E-06 | 0.000697 | 1 | 1 |
| 346 | sp\|P70662 | LDB1_MOU | 47 kDa | 0.999359 | 7 | 0.86 | P70662 | Ldb1 | LIM domain | 26.67237 | 21.65568 | 26.89319 | 21.90304 | 26.29582 | 22.43223 | 4.62348 | 3.83559 | 5.41137 | 24.30872 | 13.96704 | 5.331501 | 3.01E-06 | 0.000427 | 1 | 1 |
| 150 | sp\|Q6IRU5 | CLCB_MOU | 25 kDa | 0.999464 | 12 | 0.92 | Q6IRU5 | Cltb | Clathrin ligh | 28.38085 | 23.80329 | 28.405 | 24.14557 | 28.20823 | 23.45666 | 4.529515 | 3.830334 | 5.228696 | 26.0666 | 15.41923 | 5.895842 | 1.57E-06 | 0.000425 | 1 | 1 |
| 465 | sp\|O08585 | CLCA_MOU | 26 kDa | 0.998487 | 6 | 0.83 | O08585 | Clta | Clathrin ligh | 26.99857 | 21.91196 | 26.85232 | 23.2787 | 26.83168 | 22.39684 | 4.365024 | 3.419942 | 5.310106 | 24.71168 | 11.02218 | 3.823632 | 1.54E-05 | 0.001047 | 1 | 1 |
| 1006 | sp\|Q8R1F6 | HID1_MOU | 89 kDa | 0.995228 | 3 | 1 | Q8R1F6 | Hid1 | Protein HID | 21.51472 | 18.28828 | 21.00575 | 17.14254 | 22.13313 | 16.37584 | 4.282311 | 2.959029 | 5.605593 | 19.41004 | 7.722831 | 1.594312 | 0.000143 | 0.004042 | 1 | 1 |
| 51 | sp\|Q8BZ98 | DYN3_MOU | 97 kDa | 0.999347 | 20 | 0.65 | Q8BZ98 | Dnm3 | Dynamin-3 | 25.00528 | 21.075 | 23.35635 | 19.66307 | 25.03548 | 20.22352 | 4.145173 | 2.745449 | 5.544897 | 22.39312 | 7.067259 | 1.03641 | 0.000245 | 0.004931 | 1 | 1 |
| 131 | sp\|Q61474 | MSI1H_MO | 39 kDa | 0.999636 | 13 | 0.92 | Q61474 | Msi1 | RNA-bindin | 28.00456 | 24.28889 | 27.83918 | 23.26068 | 27.77549 | 23.72374 | 4.115307 | 3.306103 | 4.924512 | 25.81542 | 12.10503 | 4.477346 | 7.7E-06 | 0.000746 | 1 | 1 |
| 25 | sp\|O35245 | PKD2_MOU | 109 kDa | 0.999878 | 26 | 1 | O35245 | Pkd2 | Polycystin-2 | 26.92774 | 23.18959 | 26.6901 | 22.84655 | 26.98119 | 22.60046 | 3.98748 | 3.303276 | 4.671684 | 24.8726 | 13.87119 | 5.291385 | 3.15E-06 | 0.000427 | 1 | 1 |
| 45 | sp\|P28740 | KIF2A_MOU | 80 kDa | 0.999855 | 21 | 1 | P28740 | Kif2a | Kinesin-like | 27.45602 | 23.07958 | 26.23326 | 23.07752 | 27.59822 | 23.29447 | 3.94531 | 2.947576 | 4.943043 | 25.12318 | 9.436634 | 2.85807 | 4.13E-05 | 0.001752 | 1 | 1 |
| 35 | sp\|Q6NZC7 | S23IP_MOU | 111 kDa | 0.999696 | 23 | 1 | Q6NZC7 | Sec23ip | SEC23-inte | 28.70469 | 24.29238 | 27.76828 | 24.26505 | 28.49033 | 24.75437 | 3.883829 | 3.049447 | 4.718211 | 26.37918 | 11.07943 | 3.933021 | 1.37E-05 | 0.000976 | 1 | 1 |
| 164 | sp\|Q920Q6 | MSI2H_MO | 37 kDa | 0.999498 | 11 | 0.91 | Q920Q6 | Msi2 | RNA-bindin | 30.20478 | 26.58534 | 29.58572 | 24.87125 | 30.10317 | 27.15205 | 3.761676 | 2.325269 | 5.198084 | 28.08372 | 6.249632 | 0.272013 | 0.000506 | 0.006933 | 1 | 1 |
| 202 | sp\|Q61235 | SNTB2_MO | 56 kDa | 0.99945 | 10 | 0.9 | Q61235 | Sntb2 | Beta-2-synt | 23.90047 | 20.00991 | 23.21943 | 18.6166 | 23.04242 | 20.27871 | 3.752367 | 2.538014 | 4.966719 | 21.51126 | 7.374136 | 1.303293 | 0.000189 | 0.004586 | 1 | 1 |
| 415 | sp\|P42669 | PURA_MOU | 35 kDa | 0.999562 | 6 | 1 | P42669 | Pura | Transcriptio | 27.2554 | 23.15779 | 26.17552 | 23.10798 | 27.42142 | 23.38845 | 3.732709 | 2.792645 | 4.672772 | 25.08443 | 9.475831 | 2.884084 | 4.03E-05 | 0.001752 | 1 | 1 |
| 329 | sp\|P06879 | PRL_MOUS | 25 kDa | 0.999529 | 7 | 1 | P06879 | Prl | Prolactin | 30.24519 | 26.1948 | 30.41172 | 26.66332 | 29.35326 | 26.185 | 3.655683 | 2.766844 | 4.544523 | 28.17555 | 9.804813 | 3.125505 | 3.15E-05 | 0.001631 | 1 | 1 |
| 788 | sp\|P16125 | LDHB_MOU | 37 kDa | 0.999654 | 3 | 0.67 | P16125 | Ldhb | L-lactate de | 21.37743 | 17.29409 | 19.77552 | 17.52084 | 22.76833 | 18.14281 | 3.654516 | 1.893459 | 5.415572 | 19.47984 | 4.952303 | -1.12114 | 0.001877 | 0.014463 | 1 | 1 |
| 185 | sp\|Q7TSH6 | SCAF4_MO | 129 kDa | 0.999718 | 10 | 0.7 | Q7TSH6 | Scaf4 | SR-related a | 26.36227 | 22.25709 | 25.69055 | 23.08979 | 26.06716 | 21.89685 | 3.625415 | 2.680733 | 4.570096 | 24.22728 | 9.158468 | 2.670024 | 4.99E-05 | 0.001989 | 1 | 1 |
| 317 | sp\|Q8BP99 | CA216_MO | 25 kDa | 0.999646 | 7 | 1 | Q8BP99 | NA | UPF0500 p | 22.22913 | 17.70089 | 22.15634 | 19.02669 | 22.08825 | 19.02444 | 3.573903 | 2.569788 | 4.578018 | 20.37096 | 8.493954 | 2.195138 | 7.97E-05 | 0.002702 | 1 | 1 |
| 631 | sp\|Q924W | RBTN1_MO | 18 kDa | 0.999432 | 4 | 0.25 | Q924W9 | Lmo1 | Rhombotin | 20.35864 | 15.1481 | 20.83886 | 17.96954 | 19.56281 | 16.95335 | 3.563107 | 1.795188 | 5.331027 | 18.47189 | 4.809689 | -1.28917 | 0.002197 | 0.016125 | 1 | 1 |
| 1019 | sp\|P07759 | SPA3K_MO | 47 kDa | 0.999735 | 2 | 1 | P07759 | Serpina3k | Serine prote | 18.43752 | 14.04793 | 19.3601 | NA | 18.39439 | 16.31902 | 3.547192 | 1.705851 | 5.388533 | 17.31179 | 4.775089 | -1.56869 | 0.003536 | 0.021945 | 1 | 1 |
| 222 | sp\|O09044 | SNP23_MO | 23 kDa | 0.999882 | 9 | 0.89 | O09044 | Snap23 | Synaptosor | 24.79802 | 21.10309 | 24.14536 | 20.64175 | 24.43624 | 21.11742 | 3.505791 | 2.767889 | 4.243692 | 22.70698 | 11.30807 | 4.061053 | 1.2E-05 | 0.000976 | 1 | 1 |
| 137 | sp\|Q3UZ01 | RNPC3_MO | 58 kDa | 0.999248 | 13 | 1 | Q3UZ01 | Rnpc3 | RNA-bindin | 25.61458 | 20.83318 | 24.21064 | 22.35387 | 25.69598 | 21.84229 | 3.497288 | 2.152342 | 4.842235 | 23.42509 | 6.205503 | 0.22841 | 0.000527 | 0.007135 | 1 | 1 |
| 955 | sp\|Q9DBP5 | KCY_MOUS | 22 kDa | 0.99784 | 3 | 1 | Q9DBP5 | Cmpk1 | UMP-CMP k | 17.69648 | 16.82647 | 17.40337 | 14.87972 | 18.18634 | 11.29231 | 3.429234 | 0.354809 | 6.503659 | 16.04745 | 2.661851 | -4.17299 | 0.03373 | 0.087525 | 1 | 0 |
| 1058 | sp\|Q5SUR0 | PUR4_MOU | 145 kDa | 0.999277 | 2 | 1 | Q5SUR0 | Pfas | Phosphorib | 18.74609 | 17.12656 | 18.66624 | 12.98109 | 19.0687 | 16.2118 | 3.387192 | 0.985407 | 5.788977 | 17.13341 | 3.365552 | -3.16379 | 0.012811 | 0.047575 | 1 | 1 |
| 240 | sp\|Q9WTY1 | PDCD7_MO | 54 kDa | 0.999289 | 9 | 1 | Q9WTY1 | Pdcd7 | Programme | 24.95165 | 20.8272 | 23.6166 | 21.2312 | 25.04715 | 21.50362 | 3.351132 | 2.257519 | 4.444745 | 22.8629 | 7.312712 | 1.250699 | 0.000199 | 0.004622 | 1 | 1 |
| 1277 | sp\|Q9D287 | SPF27_MO | 26 kDa | 0.99577 | 2 | 1 | Q9D287 | Bcas2 | Pre-mRNA-s | 18.70835 | 13.9885 | 18.59317 | 17.27421 | 18.81194 | 15.04786 | 3.267629 | 1.385691 | 5.149567 | 17.07067 | 4.143602 | -2.1152 | 0.004767 | 0.026134 | 1 | 1 |
| 1278.1 | sp\|Q7TSF0 | DSG1C_MC | 101 kDa | 0.995714 | 2 | 1 | Q7TSF0 | Dsg1c | Desmoglein | 18.70835 | 13.9885 | 18.59317 | 17.27421 | 18.81194 | 15.04786 | 3.267629 | 1.385691 | 5.149567 | 17.07067 | 4.143602 | -2.1152 | 0.004767 | 0.026134 | 1 | 1 |
| 180 | sp\|Q80U40 | RIMB2_MO | 118 kDa | 0.998865 | 11 | 1 | Q80U40 | Rimbp2 | RIMS-bindi | 22.50782 | 19.61629 | 22.41039 | 18.36859 | 23.01815 | 20.31138 | 3.213367 | 1.9648 | 4.461934 | 21.03877 | 6.14185 | 0.165072 | 0.00056 | 0.007302 | 1 | 1 |
| 18 | sp\|P63017 | HSP7C_MO | 71 kDa | 0.999881 | 31 | 0.65 | P63017 | Hspa8 | Heat shock | 27.85382 | 24.30508 | 27.7775 | 24.49842 | 27.83724 | 25.23323 | 3.143945 | 2.361162 | 3.926729 | 26.25088 | 9.559479 | 3.007402 | 3.52E-05 | 0.001704 | 1 | 1 |
| 1184 | sp\|Q8K386 | RAB15_MO | 24 kDa | 0.998107 | 2 | 0.5 | Q8K386 | Rab15 | Ras-related | 18.95745 | 15.91514 | 19.35923 | 16.2916 | 19.33877 | 16.0425 | 3.135401 | 2.470756 | 3.800046 | 17.65078 | 11.22805 | 4.017113 | 1.25E-05 | 0.000976 | 1 | 1 |
| 1002.2 | sp\|Q80VQ0 | AL3B1_MO | 52 kDa | 0.995458 | 3 | 1 | Q80VQ0 | Aldh3b1 | Aldehyde d | 20.88722 | 19.67256 | 23.96654 | 17.32303 | 21.72996 | 20.42034 | 3.05593 | 0.571543 | 5.540317 | 20.66661 | 2.935451 | -3.77595 | 0.02298 | 0.067424 | 1 | 0 |
| 817 | tr\|A0A0B4J | A0A0B4J1N | 11 kDa | 0.999419 | 3 | 0.33 | A0A0B4J1M | Ighv1-77 | Immunoglo | 19.53749 | 16.75791 | 19.94923 | 16.80292 | 19.96815 | 16.76371 | 3.043447 | 2.402736 | 3.684159 | 18.29657 | 11.30587 | 4.059855 | 1.2E-05 | 0.000976 | 1 | 1 |
| 1034 | tr\|A0A075E | A0A075B6 | 10 kDa | 0.999481 | 2 | 1 | A0A075B6 | Igkv4-53 | Immunoglo | 22.30582 | 18.99434 | 21.35155 | 19.49426 | 22.41964 | 18.67668 | 2.970574 | 2.010291 | 3.930856 | 20.54038 | 7.382304 | 1.310256 | 0.000188 | 0.004586 | 1 | 1 |
| 32 | sp\|Q6PDG5 | SMRC2_MC | 133 kDa | 0.999852 | 24 | 0.67 | Q6PDG5 | Smarcc2 | SWI/SNF co | 25.9951 | 22.71468 | 25.24442 | 22.74158 | 26.21342 | 23.12373 | 2.957648 | 2.127008 | 3.788287 | 24.33882 | 8.47531 | 2.239788 | 7.54E-05 | 0.002622 | 1 | 1 |
| 224 | sp\|Q3UH66 | WNK2_MOU | 228 kDa | 0.999818 | 9 | 0.89 | Q3UH66 | Wnk2 | Serine/threo | 21.59838 | 18.0727 | 21.81789 | 19.49968 | 21.87457 | 18.92093 | 2.932511 | 1.959063 | 3.90596 | 20.29736 | 7.189146 | 1.143654 | 0.000221 | 0.004622 | 1 | 1 |
| 401 | sp\|Q9CX30 | YIF1B_MOU | 34 kDa | 0.999669 | 6 | 1 | Q9CX30 | Yif1b | Protein YIF1 | 25.40531 | 22.23832 | 25.71674 | 23.02212 | 25.38173 | 22.52984 | 2.904495 | 2.157756 | 3.651234 | 24.04901 | 9.257689 | 2.803678 | 4.32E-05 | 0.001774 | 1 | 1 |
| 227 | sp\|Q9QZM0 | UBQL2_MO | 67 kDa | 0.999751 | 9 | 0.78 | Q9QZM0 | Ubqln2 | Ubiquilin-2 | 25.1567 | 22.82399 | 26.28683 | 22.33402 | 24.68113 | 22.3235 | 2.881051 | 1.784037 | 3.978064 | 23.93436 | 6.267431 | 0.289529 | 0.000498 | 0.006888 | 1 | 1 |
| 336 | sp\|Q8VCM | FIBG_MOUS | 49 kDa | 0.999459 | 7 | 1 | Q8VCM7 | Fgg | Fibrinogen g | 24.21079 | 21.07079 | 23.39936 | 21.09125 | 23.81612 | 20.64447 | 2.873253 | 2.09987 | 3.646637 | 22.37213 | 8.842596 | 2.511556 | 5.77E-05 | 0.002206 | 1 | 1 |
| 1022 | sp\|P00687 | AMY1_MOU | 58 kDa | 0.999718 | 2 | 1 | P00687 | Amy1 | Alpha-amyl | 22.9301 | 20.18507 | 21.64963 | 18.50636 | 23.00559 | 20.3898 | 2.834698 | 1.34859 | 4.320806 | 21.11109 | 4.55205 | -1.6006 | 0.002942 | 0.019244 | 1 | 1 |
| 790 | sp\|P03976 | KV2A5_MO | 12 kDa | 0.999638 | 3 | 0.33 | P03976 | NA | Ig kappa ch | 21.81688 | 19.43468 | 20.28002 | 18.3122 | 21.67842 | 17.72153 | 2.768972 | 1.348592 | 4.189351 | 19.87395 | 4.652269 | -1.47825 | 0.002623 | 0.018057 | 1 | 1 |
| 325 | sp\|P46062 | SIPA1_MOU | 112 kDa | 0.999569 | 7 | 1 | P46062 | Sipa1 | Signal-indu | 22.8695 | 19.37953 | 21.36005 | 19.55026 | 22.09384 | 19.09797 | 2.765208 | 1.743169 | 3.787247 | 20.72519 | 6.456705 | 0.473304 | 0.000418 | 0.006259 | 1 | 1 |
| 825 | tr\|A0A075E | A0A075B5V | 11 kDa | 0.999375 | 3 | 0.33 | A0A075B5V | Ighv1-56 | Immunoglo | 23.47433 | 20.17281 | 22.34031 | 20.90487 | 23.77537 | 20.27581 | 2.745506 | 1.665334 | 3.825678 | 21.82392 | 6.065691 | 0.088595 | 0.000602 | 0.00764 | 1 | 1 |
| 77 | sp\|Q91XU0 | WRIP1_MO | 72 kDa | 0.999602 | 17 | 1 | Q91XU0 | Wrnip1 | ATPase WR | 25.96271 | 22.74806 | 24.0297 | 22.65917 | 26.33713 | 22.96971 | 2.650867 | 1.199294 | 4.10244 | 24.11775 | 4.358126 | -1.84173 | 0.003688 | 0.022641 | 1 | 1 |
| 981 | sp\|Q6NSU3 | GL8D1_MO | 42 kDa | 0.997254 | 3 | 1 | Q6NSU3 | Glt8d1 | Glycosyltra | 18.38342 | 16.25019 | 15.80608 | 13.96295 | 19.09967 | 15.22438 | 2.617219 | 0.330419 | 4.904018 | 16.45445 | 2.731257 | -4.07193 | 0.030577 | 0.081781 | 1 | 0 |
| 869 | sp\|Q61543 | GSLG1_MO | 134 kDa | 0.998993 | 3 | 1 | Q61543 | Glg1 | Golgi appar | 22.33651 | 19.98973 | 22.35442 | 19.45564 | 22.00334 | 19.49077 | 2.58604 | 1.888317 | 3.283763 | 20.9384 | 8.821714 | 2.496481 | 5.86E-05 | 0.002206 | 1 | 1 |
| 1068 | sp\|Q9DCR2 | AP3S1_MO | 22 kDa | 0.999217 | 2 | 1 | Q9DCR2 | Ap3s1 | AP-3 compl | 21.64431 | 17.63543 | 19.06564 | 18.13848 | 21.87576 | 19.16112 | 2.550227 | 0.605097 | 4.495357 | 19.58679 | 3.128822 | -3.49846 | 0.017617 | 0.056254 | 1 | 0 |
| 1242 | sp\|P21278 | GNA11_MC | 42 kDa | 0.997167 | 2 | 1 | P21278 | Gna11 | Guanine nu | 16.86335 | 14.97846 | 18.33561 | 17.22075 | 17.6544 | 13.02014 | 2.544668 | 0.102554 | 4.986782 | 16.34545 | 2.486656 | -4.42848 | 0.043297 | 0.105893 | 1 | 0 |
| 113 | tr\|Q3UJB0 | Q3UJB0_M | 98 kDa | 0.999862 | 14 | 1 | Q3UJB0 | Sf3b2 | Splicing fac | 24.56331 | 21.89792 | 23.82033 | 22.09233 | 24.76927 | 21.53842 | 2.541412 | 1.698843 | 3.383981 | 23.1136 | 7.184315 | 1.174536 | 0.000212 | 0.004622 | 1 | 1 |
| 487 | sp\|P29699 | FETUA_MO | 37 kDa | 0.99975 | 5 | 1 | P29699 | Ahsg | Alpha-2-HS | 25.11261 | 22.42983 | 25.37017 | 22.68068 | 24.46618 | 22.30354 | 2.511637 | 1.720119 | 3.303154 | 23.72717 | 7.552614 | 1.505597 | 0.000154 | 0.004094 | 1 | 1 |
| 7 | sp\|P70336 | ROCK2_MC | 161 kDa | 0.99975 | 49 | 1 | P70336 | Rock2 | Rho-associ | 27.49656 | 24.87803 | 28.27345 | 24.56162 | 26.65873 | 25.46058 | 2.509501 | 1.362953 | 3.656049 | 26.22149 | 5.223309 | -0.81026 | 0.001403 | 0.012336 | 1 | 1 |
| 614 | sp\|O70439 | STX7_MOU | 30 kDa | 0.999563 | 4 | 1 | O70439 | Stx7 | Syntaxin-7 | 24.47173 | 21.79934 | 24.09237 | 22.1601 | 24.28612 | 21.40442 | 2.495454 | 1.75852 | 3.232387 | 23.03568 | 8.059757 | 1.919958 | 0.000103 | 0.003176 | 1 | 1 |
| 171 | sp\|Q99KN9 | EPN4_MOU | 69 kDa | 0.999326 | 11 | 1 | Q99KN9 | Clint1 | Clathrin inte | 24.55638 | 21.62096 | 24.17584 | 21.75137 | 23.7416 | 21.67437 | 2.47571 | 1.745107 | 3.206314 | 22.92009 | 8.065269 | 1.924321 | 0.000103 | 0.003176 | 1 | 1 |
| 1046 | sp\|Q80TT2 | BAIP3_MOU | 127 kDa | 0.999382 | 2 | 1 | Q80TT2 | Baiap3 | BAI1-assoc | 18.3657 | 15.61973 | 18.83519 | 17.41713 | 19.38739 | 16.12937 | 2.474014 | 1.205602 | 3.742425 | 17.62575 | 4.654709 | -1.47529 | 0.002616 | 0.018057 | 1 | 1 |
| 760 | sp\|Q8R3N6 | THOC1_MC | 75 kDa | 0.997356 | 4 | 1 | Q8R3N6 | Thoc1 | THO compl | 21.25154 | 18.81515 | 20.96518 | 18.36662 | 21.37676 | 19.01436 | 2.465782 | 1.747415 | 3.184149 | 19.96494 | 8.169757 | 2.006487 | 9.47E-05 | 0.003058 | 1 | 1 |
| 47 | sp\|Q9QX47 | SON_MOUS | 266 kDa | 0.999742 | 20 | 1 | Q9QX47 | Son | Protein SON | 23.98495 | 21.62267 | 23.9458 | 21.34901 | 24.5157 | 22.22077 | 2.417995 | 1.5945 | 3.24149 | 22.93982 | 6.989019 | 1.012516 | 0.000247 | 0.004931 | 1 | 1 |
| 812 | sp\|Q61753 | SERA_MOU | 57 kDa | 0.999441 | 3 | 1 | Q61753 | Phgdh | D-3-phosph | 18.26501 | 15.45136 | 18.07564 | 14.6353 | 17.04925 | 16.12053 | 2.394237 | 1.1893 | 3.599173 | 16.59952 | 4.741913 | -1.37011 | 0.002371 | 0.017008 | 1 | 1 |
| 191 | sp\|Q3U0V1 | FUBP2_MO | 77 kDa | 0.999648 | 10 | 0.9 | Q3U0V1 | Khsrp | Far upstrea | 24.72273 | 21.66067 | 24.14036 | 21.8274 | 23.23654 | 21.44241 | 2.389714 | 1.380986 | 3.398441 | 22.83835 | 5.653567 | -0.33875 | 0.000901 | 0.009848 | 1 | 1 |
| 836.1 | sp\|P01638 | KV5A6_MO | 13 kDa | 0.999277 | 3 | 1 | P01638 | NA | Ig kappa ch | 17.54744 | 14.73293 | 15.63094 | 16.03972 | 17.94353 | 13.2049 | 2.381449 | 0.29432 | 4.468578 | 15.84991 | 2.722969 | -4.08399 | 0.030937 | 0.082417 | 1 | 0 |
| 12 | sp\|Q8CGB6 | TNS2_MOU | 152 kDa | 0.999844 | 38 | 0.97 | Q8CGB6 | Tns2 | Tensin-2 | 27.48814 | 25.23302 | 27.2899 | 24.28074 | 27.30562 | 25.42792 | 2.380662 | 1.497997 | 3.263327 | 26.17089 | 6.429779 | 0.463807 | 0.00042 | 0.006259 | 1 | 1 |
| 235 | sp\|Q9Z2N8 | ACL6A_MO | 47 kDa | 0.999542 | 9 | 0.89 | Q9Z2N8 | Actl6a | Actin-like pr | 24.21382 | 21.73671 | 23.18929 | 21.35708 | 24.51204 | 21.71926 | 2.367363 | 1.399225 | 3.335502 | 22.78803 | 5.8355 | -0.14725 | 0.000752 | 0.008839 | 1 | 1 |
| 826 | sp\|Q9CQN | TRAP1_MO | 80 kDa | 0.999366 | 3 | 1 | Q9CQN1 | Trap1 | Heat shock | 21.91922 | 18.94401 | 19.9797 | 19.23083 | 22.06275 | 18.69712 | 2.363236 | 0.965036 | 3.761437 | 20.13894 | 4.033558 | -2.2582 | 0.005452 | 0.027897 | 1 | 1 |
| 511 | sp\|Q3TWN | CNNM2_M | 97 kDa | 0.999457 | 5 | 1 | Q3TWN3 | Cnnm2 | Metal trans | 20.05472 | 17.92137 | 19.41725 | 17.7696 | 20.64281 | 17.37158 | 2.350741 | 1.425285 | 3.276196 | 18.86289 | 6.060599 | 0.086212 | 0.000603 | 0.00764 | 1 | 1 |
| 831 | sp\|Q0VBL3 | RBM15_MC | 106 kDa | 0.999348 | 3 | 1 | Q0VBL3 | Rbm15 | RNA-bindin | 22.4191 | 20.36229 | 20.53938 | 18.73121 | 21.96914 | 18.78609 | 2.34934 | 0.798737 | 3.899944 | 20.46787 | 3.615728 | -2.81743 | 0.009231 | 0.038395 | 1 | 1 |
| 1259 | sp\|Q9CU62 | SMC1A_MC | 143 kDa | 0.996524 | 2 | 1 | Q9CU62 | Smc1a | Structural m | 19.69182 | 17.35776 | 20.37165 | 18.03578 | 19.11579 | 16.78584 | 2.333295 | 1.22361 | 3.44298 | 18.55977 | 5.017885 | -1.0449 | 0.001748 | 0.013699 | 1 | 1 |
| 209 | tr\|Q6PHQ9 | Q6PHQ9_M | 72 kDa | 0.999289 | 10 | 0.8 | Q6PHQ9 | Pabpc4 | Polyadenyl | 22.35413 | 19.69856 | 21.93965 | 19.30669 | 21.99438 | 20.32371 | 2.319733 | 1.48701 | 3.152457 | 20.93619 | 6.630695 | 0.680866 | 0.000339 | 0.00576 | 1 | 1 |
| 281 | sp\|Q61103 | REQU_MOU | 44 kDa | 0.999382 | 8 | 0.88 | Q61103 | Dpf2 | Zinc finger p | 24.10588 | 21.64723 | 23.5233 | 21.22568 | 24.4496 | 22.29356 | 2.304103 | 1.347004 | 3.261203 | 22.87421 | 5.74507 | -0.24186 | 0.000822 | 0.009368 | 1 | 1 |
| 485 | sp\|Q9D0L4 | ADCK1_MC | 60 kDa | 0.999797 | 5 | 1 | Q9D0L4 | Adck1 | Uncharacte | 22.14353 | 19.38703 | 21.67331 | 19.53578 | 22.35148 | 20.44388 | 2.267211 | 1.346956 | 3.187467 | 20.9225 | 5.877701 | -0.09952 | 0.000718 | 0.008616 | 1 | 1 |
| 1014 | sp\|P12961 | 7B2_MOUS | 24 kDa | 0.999876 | 2 | 1 | P12961 | Scg5 | Neuroendo | 18.97902 | 17.09026 | 18.34253 | 15.80426 | 19.94018 | 17.59953 | 2.255898 | 0.82642 | 3.685375 | 17.9593 | 3.766108 | -2.61326 | 0.007614 | 0.034782 | 1 | 1 |
| 148 | sp\|Q9WU2 | MTHR_MOU | 75 kDa | 0.999556 | 12 | 1 | Q9WU20 | Mthfr | Methylenet | 25.13436 | 23.70929 | 24.14165 | 22.34666 | 25.26627 | 21.77208 | 2.238083 | 0.863195 | 3.61297 | 23.72839 | 3.884718 | -2.45451 | 0.006557 | 0.031417 | 1 | 1 |
| 100.1 | sp\|P17879 | HS71B_MO | 70 kDa | 0.999699 | 15 | 0.73 | P17879 | Hspa1b | Heat shock | 24.99751 | 22.79349 | 24.59787 | 22.00965 | 24.82205 | 22.90713 | 2.23572 | 1.426672 | 3.044768 | 23.68795 | 6.577565 | 0.630384 | 0.000356 | 0.00576 | 1 | 1 |
| 100.2 | sp\|Q61696 | HS71A_MO | 70 kDa | 0.999699 | 15 | 0.73 | Q61696 | Hspa1a | Heat shock | 24.99751 | 22.79349 | 24.59787 | 22.00965 | 24.82205 | 22.90713 | 2.23572 | 1.426672 | 3.044768 | 23.68795 | 6.577565 | 0.630384 | 0.000356 | 0.00576 | 1 | 1 |
| 10 | sp\|Q9EPU0 | RENT1_MO | 124 kDa | 0.999794 | 40 | 1 | Q9EPU0 | Upf1 | Regulator o | 26.88991 | 24.32191 | 26.69588 | 24.45162 | 26.57692 | 24.69033 | 2.232949 | 1.592188 | 2.873711 | 25.60443 | 8.294372 | 2.10313 | 8.62E-05 | 0.002851 | 1 | 1 |
| 553 | sp\|P62141 | PP1B_MOU | 37 kDa | 0.998911 | 5 | 0.4 | P62141 | Ppp1cb | Serine/threo | 22.1122 | 19.15572 | 21.42328 | 20.32593 | 22.05817 | 19.44851 | 2.221164 | 1.258729 | 3.183599 | 20.75397 | 5.507569 | -0.49578 | 0.001044 | 0.010776 | 1 | 1 |
| 176 | sp\|Q9Z204 | HNRPC_MC | 34 kDa | 0.999252 | 11 | 1 | Q9Z204 | Hnrnpc | Heterogene | 25.77075 | 23.35799 | 24.80237 | 23.26727 | 26.29288 | 23.58338 | 2.21912 | 1.21016 | 3.228079 | 24.51244 | 5.248771 | -0.78161 | 0.001365 | 0.012261 | 1 | 1 |
| 967 | sp\|Q99LX0 | PARK7_MO | 20 kDa | 0.997612 | 3 | 1 | Q99LX0 | Park7 | Protein/nuc | 19.73811 | 17.72553 | 20.29779 | 17.60711 | 19.48512 | 17.55125 | 2.212375 | 1.473917 | 2.950833 | 18.73415 | 7.130727 | 1.140447 | 0.000219 | 0.004622 | 1 | 1 |
| 133 | sp\|Q8C3F2 | F120C_MO | 120 kDa | 0.999553 | 13 | 0.85 | Q8C3F2 | Fam120c | Constitutive | 24.45541 | 21.95124 | 24.25257 | 21.71945 | 23.83938 | 22.35075 | 2.175304 | 1.422157 | 2.928451 | 23.0948 | 6.874494 | 0.909007 | 0.000273 | 0.005213 | 1 | 1 |
| 5 | sp\|Q8R0W | EPIPL_MOU | 723 kDa | 0.999889 | 53 | 0.96 | Q8R0W0 | Eppk1 | Epiplakin | 26.42704 | 23.51274 | 24.96372 | 24.08294 | 26.96435 | 24.25857 | 2.166955 | 0.852678 | 3.481232 | 25.03489 | 3.934718 | -2.3882 | 0.00616 | 0.030487 | 1 | 1 |
| 328 | sp\|Q8BRM2 | GORAB_MC | 41 kDa | 0.999541 | 7 | 1 | Q8BRM2 | Gorab | RAB6-intera | 24.13988 | 21.38348 | 22.70421 | 21.12528 | 23.93925 | 21.7973 | 2.159091 | 1.084359 | 3.233824 | 22.5149 | 4.794256 | -1.30754 | 0.002236 | 0.016125 | 1 | 1 |
| 437 | sp\|O55203 | LDB2_MOU | 43 kDa | 0.999263 | 6 | 0.83 | O55203 | Ldb2 | LIM domain | 23.60646 | 21.88231 | 23.91088 | 20.79993 | 23.29214 | 21.68058 | 2.148885 | 1.242273 | 3.055498 | 22.52872 | 5.650481 | -0.32866 | 0.000888 | 0.009789 | 1 | 1 |
| 652 | sp\|Q9R226 | KHDR3_MC | 39 kDa | 0.99932 | 4 | 0.75 | Q9R226 | Khdrbs3 | KH domain- | 23.03737 | 20.64425 | 21.33682 | 19.77296 | 22.90573 | 20.54607 | 2.105547 | 0.835703 | 3.375391 | 21.37387 | 3.956992 | -2.35877 | 0.005992 | 0.029873 | 1 | 1 |
| 928 | sp\|P01831 | THY1_MOU | 18 kDa | 0.998266 | 3 | 1 | P01831 | Thy1 | Neuroendo | 18.01021 | 16.79774 | 18.45987 | 15.83689 | 18.24632 | 15.81 | 2.090589 | 1.221915 | 2.959262 | 17.19351 | 5.73728 | -0.23635 | 0.000814 | 0.009355 | 1 | 1 |
| 891 | sp\|Q8BG32 | PSD11_MO | 47 kDa | 0.998773 | 3 | 1 | Q8BG32 | Psmd11 | 26S protea | 22.34552 | 19.62844 | 19.25061 | 18.66524 | 22.13427 | 19.29053 | 2.048736 | 0.048762 | 4.04871 | 20.2191 | 2.444625 | -4.48977 | 0.045988 | 0.109594 | 1 | 0 |
| 892 | sp\|Q80WC | AGFG2_MO | 49 kDa | 0.99877 | 3 | 1 | Q80WC7 | Agfg2 | Arf-GAP dor | 22.34552 | 19.62844 | 19.25061 | 18.66524 | 22.13427 | 19.29053 | 2.048736 | 0.048762 | 4.04871 | 20.2191 | 2.444625 | -4.48977 | 0.045988 | 0.109594 | 1 | 0 |
| 98 | sp\|Q80U23 | SNPH_MOU | 54 kDa | 0.999775 | 15 | 0.87 | Q80U23 | Snph | Syntaphilin | 24.56258 | 20.65571 | 23.42978 | 21.67479 | 24.41434 | 23.97825 | 2.032649 | 0.019173 | 4.046125 | 23.11924 | 2.409166 | -4.54144 | 0.048392 | 0.112169 | 1 | 0 |
| 330 | sp\|Q9Z0H3 | SNF5_MOU | 44 kDa | 0.999526 | 7 | 1 | Q9Z0H3 | Smarcb1 | SWI/SNF-re | 24.42627 | 22.33792 | 23.4462 | 21.24587 | 24.44884 | 22.66562 | 2.023966 | 0.867448 | 3.180484 | 23.09512 | 4.176393 | -2.07294 | 0.004581 | 0.025802 | 1 | 1 |
| 128 | sp\|Q8CJG0 | AGO2_MOU | 97 kDa | 0.999715 | 13 | 0.85 | Q8CJG0 | Ago2 | Protein argo | 23.84731 | 21.8418 | 23.43332 | 21.78657 | 23.93438 | 21.52956 | 2.019025 | 1.344763 | 2.693288 | 22.72882 | 7.127113 | 1.137234 | 0.000219 | 0.004622 | 1 | 1 |
| 534 | sp\|Q91YN9 | BAG2_MOU | 23 kDa | 0.999217 | 5 | 1 | Q91YN9 | Bag2 | BAG family | 22.41719 | 20.56972 | 22.06817 | 20.48345 | 22.55459 | 19.98094 | 2.001945 | 1.276328 | 2.727562 | 21.34568 | 6.566671 | 0.620783 | 0.000359 | 0.00576 | 1 | 1 |
| 21 | sp\|O08810 | U5S1_MOU | 109 kDa | 0.999725 | 28 | 0.96 | O08810 | Eftud2 | 116 kDa U5 | 25.76133 | 24.08064 | 24.68752 | 22.86079 | 26.05939 | 23.60689 | 1.986638 | 0.821078 | 3.152199 | 24.50943 | 4.067567 | -2.21381 | 0.005229 | 0.027525 | 1 | 1 |
| 1081 | sp\|Q3TJZ6 | FA98A_MO | 55 kDa | 0.999131 | 2 | 1 | Q3TJZ6 | Fam98a | Protein FAM | 26.18688 | 23.61623 | 25.0779 | 23.13685 | 26.61336 | 25.17665 | 1.982799 | 0.452619 | 3.51298 | 24.96798 | 3.092337 | -3.55056 | 0.018516 | 0.058038 | 1 | 0 |
| 888 | sp\|Q9CYK2 | QPCT_MOU | 41 kDa | 0.998803 | 3 | 1 | Q9CYK2 | Qpct | Glutaminyl- | 22.55832 | 20.3088 | 22.12631 | 20.06781 | 21.77316 | 20.14065 | 1.98018 | 1.250027 | 2.710333 | 21.16251 | 6.454927 | 0.513303 | 0.000398 | 0.006063 | 1 | 1 |
| 505 | sp\|Q9CQF3 | CPSF5_MO | 26 kDa | 0.999523 | 5 | 1 | Q9CQF3 | Nudt21 | Cleavage an | 23.55598 | 21.18985 | 22.98271 | 21.12811 | 23.0107 | 21.40734 | 1.94136 | 1.245526 | 2.637194 | 22.21245 | 6.640507 | 0.690958 | 0.000336 | 0.00576 | 1 | 1 |
| 127 | sp\|Q8BHS8 | SYBU_MOU | 73 kDa | 0.999745 | 13 | 0.85 | Q8BHS8 | Sybu | Syntabulin | 24.71025 | 23.33591 | 23.35876 | 20.98313 | 24.62368 | 22.5543 | 1.939781 | 0.322297 | 3.557264 | 23.26101 | 2.861959 | -3.88219 | 0.025454 | 0.072058 | 1 | 0 |
| 331 | sp\|Q99KP6 | PRP19_MO | 55 kDa | 0.999488 | 7 | 1 | Q99KP6 | Prpf19 | Pre-mRNA-p | 24.82743 | 22.70567 | 24.27015 | 23.05885 | 25.17441 | 22.69926 | 1.93607 | 1.146878 | 2.725263 | 23.78929 | 5.839006 | -0.10786 | 0.000716 | 0.008616 | 1 | 1 |
| 1280 | sp\|Q8K0L9 | ZBT20_MOU | 81 kDa | 0.995702 | 2 | 1 | Q8K0L9 | Zbtb20 | Zinc finger a | 20.26318 | 18.44949 | 22.24132 | 20.89245 | 21.43402 | 18.80873 | 1.929284 | 0.076287 | 3.782281 | 20.3482 | 2.48469 | -4.43135 | 0.043419 | 0.105893 | 1 | 0 |
| 621 | sp\|Q9ER00 | STX12_MO | 31 kDa | 0.999496 | 4 | 1 | Q9ER00 | Stx12 | Syntaxin-12 | 23.70315 | 22.0989 | 23.50824 | 20.65139 | 23.50365 | 22.18932 | 1.925143 | 0.830354 | 3.019932 | 22.60911 | 4.196461 | -2.04716 | 0.004472 | 0.025501 | 1 | 1 |
| 990 | sp\|Q3THK7 | GUAA_MOU | 77 kDa | 0.99685 | 3 | 1 | Q3THK7 | Gmps | GMP syntha | 19.43538 | 18.31149 | 19.37049 | 17.27847 | 19.24929 | 16.69458 | 1.92354 | 0.870142 | 2.976938 | 18.38995 | 4.357721 | -1.84224 | 0.00369 | 0.022641 | 1 | 1 |
| 1122 | tr\|B2RXQ2 | B2RXQ2_M | 143 kDa | 0.998694 | 2 | 1 | B2RXQ2 | Ppfia1 | Ppfia1 prote | 20.92404 | 19.22578 | 20.93676 | 18.99896 | 20.78955 | 18.725 | 1.900201 | 1.251107 | 2.549294 | 19.93335 | 6.967754 | 0.994115 | 0.000252 | 0.004945 | 1 | 1 |
| 1261 | sp\|P51660 | DHB4_MOU | 79 kDa | 0.996427 | 2 | 1 | P51660 | Hsd17b4 | Peroxisoma | 20.02291 | 18.00302 | 19.22207 | 17.81911 | 20.25533 | 17.99475 | 1.894478 | 1.067929 | 2.721027 | 18.8862 | 5.455607 | -0.52104 | 0.001057 | 0.010776 | 1 | 1 |
| 428 | sp\|P70333 | HNRH2_MC | 49 kDa | 0.999352 | 6 | 0.33 | P70333 | Hnrnph2 | Heterogene | 22.53427 | 20.68878 | 23.45805 | 20.71131 | 23.04984 | 22.05715 | 1.861641 | 0.730894 | 2.992388 | 22.08323 | 3.928993 | -2.39577 | 0.006204 | 0.030593 | 1 | 1 |
| 121 | sp\|O70133 | DHX9_MOU | 149 kDa | 0.999414 | 14 | 1 | O70133 | Dhx9 | ATP-depend | 24.84958 | 22.61174 | 24.1873 | 22.58325 | 24.62655 | 22.89707 | 1.857126 | 1.147273 | 2.566979 | 23.62592 | 6.226926 | 0.289132 | 0.000492 | 0.006878 | 1 | 1 |
| 786 | tr\|A0A075E | A0A075B6 | 11 kDa | 0.999674 | 3 | 0.33 | A0A075B6 | Ighv1-78 | Immunoglo | 21.42798 | 18.9608 | 20.20741 | 20.11767 | 21.60376 | 18.59672 | 1.854654 | 0.546871 | 3.162437 | 20.15239 | 3.384371 | -3.13746 | 0.012495 | 0.047066 | 1 | 1 |
| 1056.2 | tr\|Q63ZW6 | Q63ZW6_M | 162 kDa | 0.999285 | 2 | 1 | Q63ZW6 | Col4a5 | Col4a5 pro | 19.54683 | 18.24412 | 19.82987 | 18.05429 | 19.42494 | 16.95742 | 1.848606 | 0.881459 | 2.815752 | 18.67624 | 4.561446 | -1.58906 | 0.00291 | 0.019158 | 1 | 1 |
| 863 | sp\|P60867 | RS20_MOU | 13 kDa | 0.99904 | 3 | 1 | P60867 | Rps20 | 40S ribosor | 22.76709 | 20.86503 | 23.24579 | 21.0859 | 22.65141 | 21.19691 | 1.838817 | 1.141489 | 2.536146 | 21.96869 | 6.276274 | 0.338213 | 0.00047 | 0.006775 | 1 | 1 |
| 178 | sp\|Q68FH0 | PKP4_MOU | 132 kDa | 0.999035 | 11 | 1 | Q68FH0 | Pkp4 | Plakophilin | 22.72108 | 20.14036 | 21.18809 | 19.96626 | 21.97705 | 20.28417 | 1.83181 | 0.814718 | 2.848901 | 21.04617 | 4.298044 | -1.91761 | 0.00396 | 0.02377 | 1 | 1 |
| 699 | sp\|P33267 | CP2F2_MO | 56 kDa | 0.998901 | 4 | 1 | P33267 | Cyp2f2 | Cytochrom | 19.63472 | 17.50575 | 19.24635 | 17.57778 | 19.99771 | 18.32354 | 1.823905 | 0.969059 | 2.678751 | 18.71431 | 5.085218 | -0.95373 | 0.001596 | 0.013037 | 1 | 1 |
| 163 | sp\|Q8BPB5 | FBLN3_MO | 55 kDa | 0.999504 | 11 | 1 | Q8BPB5 | Efemp1 | EGF-contai | 25.46593 | 23.13289 | 24.44394 | 23.06345 | 25.39581 | 23.70309 | 1.802079 | 0.881151 | 2.723007 | 24.20085 | 4.668441 | -1.45602 | 0.002566 | 0.017983 | 1 | 1 |
| 777 | sp\|Q62159 | RHOC_MOU | 22 kDa | 0.99532 | 4 | 0.25 | Q62159 | Rhoc | Rho-related | 23.62226 | 22.13382 | 23.3445 | 21.15459 | 23.28133 | 21.55788 | 1.8006 | 0.994191 | 2.607008 | 22.51573 | 5.314763 | -0.67783 | 0.001225 | 0.011522 | 1 | 1 |
| 405 | sp\|P62754 | RS6_MOUS | 29 kDa | 0.999619 | 6 | 1 | P62754 | Rps6 | 40S ribosor | 25.81162 | 24.16112 | 26.02414 | 23.63338 | 25.38644 | 24.0423 | 1.795134 | 1.055764 | 2.534505 | 24.84316 | 5.778772 | -0.17132 | 0.00076 | 0.008839 | 1 | 1 |
| 464 | sp\|Q6ZPR4 | KCNT1_MO | 138 kDa | 0.998579 | 6 | 1 | Q6ZPR4 | Kcnt1 | Potassium c | 19.77841 | 17.82771 | 18.95494 | 16.91251 | 20.32247 | 18.98341 | 1.777396 | 0.328975 | 3.225818 | 18.79657 | 2.928464 | -3.78604 | 0.023204 | 0.067424 | 1 | 0 |
| 592 | sp\|Q9D1G1 | RAB1B_MO | 22 kDa | 0.999802 | 4 | 0.25 | Q9D1G1 | Rab1b | Ras-related | 19.76854 | 18.12916 | 20.66956 | 18.64778 | 19.87042 | 18.24636 | 1.761741 | 0.926592 | 2.596891 | 19.22197 | 5.02111 | -1.0141 | 0.00168 | 0.01351 | 1 | 1 |
| 63 | sp\|Q9ESU6 | BRD4_MOU | 156 kDa | 0.999719 | 18 | 1 | Q9ESU6 | Brd4 | Bromodom | 25.06452 | 23.9049 | 23.7937 | 22.30319 | 25.51418 | 22.88037 | 1.761315 | 0.351272 | 3.171358 | 23.91014 | 2.980954 | -3.71037 | 0.021578 | 0.065165 | 1 | 0 |
| 1200 | sp\|O35226 | PSMD4_MC | 41 kDa | 0.997946 | 2 | 1 | O35226 | Psmd4 | 26S protea | 19.63834 | 18.04042 | 19.6948 | 17.72898 | 19.696 | 17.97699 | 1.760916 | 1.148333 | 2.373499 | 18.79592 | 6.841865 | 0.87899 | 0.000281 | 0.00529 | 1 | 1 |
| 741 | sp\|P49615 | CDK5_MOU | 33 kDa | 0.9981 | 4 | 1 | P49615 | Cdk5 | Cyclin-depe | 20.73597 | 19.32378 | 21.71029 | 19.15244 | 20.60443 | 19.31223 | 1.754077 | 0.880937 | 2.627218 | 20.13986 | 4.789152 | -1.30366 | 0.002218 | 0.016125 | 1 | 1 |
| 162 | sp\|P47911 | RL6_MOUS | 34 kDa | 0.999527 | 11 | 1 | P47911 | Rpl6 | 60S ribosor | 25.64704 | 24.0894 | 25.82642 | 24.06906 | 25.67027 | 23.73458 | 1.750233 | 1.118812 | 2.381655 | 24.83946 | 6.597471 | 0.650136 | 0.000349 | 0.00576 | 1 | 1 |
| 158 | sp\|P27659 | RL3_MOUS | 46 kDa | 0.999783 | 11 | 1 | P27659 | Rpl3 | 60S ribosor | 25.38364 | 23.54433 | 25.15199 | 23.32146 | 25.12776 | 23.56829 | 1.7431 | 1.121216 | 2.364985 | 24.34958 | 6.671347 | 0.720072 | 0.000327 | 0.00576 | 1 | 1 |
| 201 | sp\|P12970 | RL7A_MOU | 30 kDa | 0.999456 | 10 | 1 | P12970 | Rpl7a | 60S ribosor | 24.97129 | 23.25296 | 25.23147 | 23.27719 | 24.75899 | 23.24403 | 1.729186 | 1.091552 | 2.36682 | 24.12266 | 6.454625 | 0.51301 | 0.000398 | 0.006063 | 1 | 1 |
| 72 | sp\|Q9D8E6 | RL4_MOUS | 47 kDa | 0.999669 | 17 | 1 | Q9D8E6 | Rpl4 | 60S ribosor | 25.47186 | 23.96585 | 25.47296 | 23.57612 | 25.53497 | 23.76307 | 1.724916 | 1.102302 | 2.347531 | 24.63081 | 6.594014 | 0.646847 | 0.00035 | 0.00576 | 1 | 1 |
| 439 | sp\|Q6NZF1 | ZC11A_MO | 86 kDa | 0.999213 | 6 | 1 | Q6NZF1 | Zc3h11a | Zinc finger C | 24.71797 | 22.96025 | 23.5417 | 22.63279 | 25.27682 | 22.80947 | 1.711325 | 0.589156 | 2.833494 | 23.6565 | 3.639359 | -2.78514 | 0.008953 | 0.03794 | 1 | 1 |
| 195 | sp\|O54941 | SMCE1_MC | 47 kDa | 0.999567 | 10 | 1 | O54941 | Smarce1 | SWI/SNF-re | 24.90503 | 23.47545 | 23.76077 | 22.6846 | 25.01228 | 22.45604 | 1.687331 | 0.587594 | 2.787067 | 23.71569 | 3.661527 | -2.75491 | 0.008702 | 0.03734 | 1 | 1 |
| 616 | sp\|P62821 | RAB1A_MO | 23 kDa | 0.999554 | 4 | 0.25 | P62821 | Rab1A | Ras-related | 19.98859 | 18.9982 | 20.65602 | 18.91909 | 20.51377 | 18.20002 | 1.680356 | 0.845515 | 2.515197 | 19.54595 | 4.790927 | -1.28672 | 0.00217 | 0.016076 | 1 | 1 |
| 144 | sp\|Q3UTJ2 | SRBS2_MO | 132 kDa | 0.999691 | 12 | 1 | Q3UTJ2 | Sorbs2 | Sorbin and | 23.3373 | 20.86913 | 21.87923 | 20.29638 | 23.36335 | 22.3805 | 1.677959 | 0.105736 | 3.250182 | 22.02098 | 2.546935 | -4.34056 | 0.039721 | 0.098467 | 1 | 0 |
| 482 | sp\|O35295 | PURB_MOU | 34 kDa | 0.99993 | 5 | 1 | O35295 | Purb | Transcriptio | 22.71308 | 20.3186 | 21.9463 | 20.82005 | 22.24894 | 20.75416 | 1.671833 | 0.901958 | 2.441707 | 21.46685 | 5.168606 | -0.84306 | 0.00143 | 0.012407 | 1 | 1 |
| 620 | sp\|Q9CR57 | RL14_MOU | 24 kDa | 0.999496 | 4 | 1 | Q9CR57 | Rpl14 | 60S ribosor | 25.19446 | 23.55456 | 25.51297 | 23.76991 | 25.40258 | 23.79248 | 1.664354 | 1.038076 | 2.290632 | 24.53783 | 6.325271 | 0.386636 | 0.000449 | 0.006612 | 1 | 1 |

| | | | | | | | | | | | | | | | | | | | | | | | | |
|---|---|---|---|---|---|---|---|---|---|---|---|---|---|---|---|---|---|---|---|---|---|---|---|---|
| 59 | sp\|Q3TKT4 | SMCA4_MO | 181 kDa | 0.999626 | 19 | 0.58 | Q3TKT4 | Smarca4 | Transcriptio | 23.64992 | 21.92278 | 22.80156 | 22.20405 | 24.01612 | 21.35074 | 1.663339 | 0.66639 | 2.660288 | 22.65753 | 3.981607 | -2.32635 | 0.005812 | 0.029083 | 1 | 1 |
| 119 | sp\|Q8VEK3 | HNRPU_MO | 88 kDa | 0.999549 | 14 | 1 | Q8VEK3 | Hnrnpu | Heterogene | 26.3786 | 24.6574 | 26.38156 | 24.48236 | 26.37727 | 25.01479 | 1.660958 | 1.008343 | 2.313572 | 25.54866 | 6.057632 | 0.118341 | 0.000578 | 0.007469 | 1 | 1 |
| 1210 | sp\|Q8R326 | PSPC1_MO | 59 kDa | 0.99779 | 2 | 1 | Q8R326 | Pspc1 | Paraspeckle | 19.0285 | 17.98653 | 18.45406 | 16.52011 | 18.40393 | 16.43615 | 1.647899 | 0.491239 | 2.80456 | 17.80488 | 3.399973 | -3.11566 | 0.01224 | 0.046361 | 1 | 1 |
| 321 | sp\|O88665 | BRD7_MOU | 74 kDa | 0.999593 | 7 | 1 | O88665 | Brd7 | Bromodom | 22.8287 | 20.91494 | 21.83513 | 20.89656 | 23.0596 | 20.983 | 1.642973 | 0.737377 | 2.54857 | 21.75299 | 4.325039 | -1.87532 | 0.003791 | 0.022951 | 1 | 1 |
| 590 | sp\|Q6P542 | ABCF1_MO | 95 kDa | 0.999828 | 4 | 1 | Q6P542 | Abcf1 | ATP-binding | 20.71838 | 17.61124 | 18.74682 | 17.92869 | 18.71617 | 17.72178 | 1.639884 | 0.27567 | 3.004098 | 18.57385 | 2.868674 | -3.87247 | 0.025217 | 0.071535 | 1 | 0 |
| 1234 | tr\|Q9JHF5\| | Q9JHF5_MO | 93 kDa | 0.997355 | 2 | 0.5 | Q9JHF5 | Tcirg1 | V-type prot | 23.00079 | 21.08417 | 23.83329 | 21.73598 | 22.65635 | 21.76223 | 1.636017 | 0.675878 | 2.596156 | 22.34547 | 4.066346 | -2.2154 | 0.005237 | 0.027525 | 1 | 1 |
| 685 | sp\|Q6ZWU9 | RS27_MOU | 9 kDa | 0.999015 | 4 | 0.5 | Q6ZWU9 | Rps27 | 40S ribosor | 21.56806 | 19.69359 | 21.59014 | 20.16681 | 21.07589 | 19.46714 | 1.635521 | 0.873956 | 2.397085 | 20.59361 | 5.111524 | -0.90868 | 0.001521 | 0.012812 | 1 | 1 |
| 566 | sp\|Q4FK66 | PR38A_MO | 37 kDa | 0.998592 | 5 | 1 | Q4FK66 | Prpf38a | Pre-mRNA-s | 21.39541 | 18.91557 | 20.33887 | 19.80857 | 21.57198 | 19.67731 | 1.634938 | 0.582709 | 2.687168 | 20.28462 | 3.708016 | -2.69176 | 0.008198 | 0.036481 | 1 | 1 |
| 186 | sp\|P62242 | RS8_MOUS | 24 kDa | 0.999717 | 10 | 1 | P62242 | Rps8 | 40S ribosor | 26.12299 | 25.33812 | 26.36842 | 24.02069 | 25.86389 | 24.09249 | 1.634663 | 0.617928 | 2.651397 | 25.3011 | 3.836817 | -2.51838 | 0.006963 | 0.032784 | 1 | 1 |
| 710 | sp\|P08226 | APOE_MOU | 36 kDa | 0.998824 | 4 | 1 | P08226 | Apoe | Apolipopro | 21.16605 | 19.66038 | 21.29676 | 19.95859 | 21.53739 | 19.48436 | 1.632292 | 0.96341 | 2.301174 | 20.51726 | 5.808305 | -0.14014 | 0.000738 | 0.008781 | 1 | 1 |
| 782 | sp\|Q9JKK1 | STX6_MOU | 29 kDa | 0.999725 | 3 | 1 | Q9JKK1 | Stx6 | Syntaxin-6 | 21.04916 | 19.62749 | 19.84004 | 19.1177 | 21.38642 | 18.66022 | 1.623402 | 0.459569 | 2.787235 | 19.94684 | 3.328788 | -3.21535 | 0.013454 | 0.04801 | 1 | 1 |
| 824 | tr\|A0A140T | A0A140T8N | 13 kDa | 0.999377 | 3 | 1 | A0A140T8N | Igkv13-84 | Immunoglo | 23.25403 | 22.29525 | 23.27321 | 21.35865 | 23.14747 | 21.18339 | 1.61248 | 0.747207 | 2.477753 | 22.41867 | 4.442578 | -1.72742 | 0.0033 | 0.021108 | 1 | 1 |
| 315 | sp\|Q8R081 | HNRPL_MO | 64 kDa | 0.999659 | 7 | 1 | Q8R081 | Hnrnpl | Heterogene | 23.52547 | 21.8458 | 23.09379 | 21.48003 | 23.40808 | 21.87588 | 1.608544 | 0.935155 | 2.281933 | 22.53818 | 5.685484 | -0.27059 | 0.000835 | 0.009433 | 1 | 1 |
| 1133 | sp\|Q9ERU9 | RBP2_MOU | 341 kDa | 0.998587 | 2 | 1 | Q9ERU9 | Ranbp2 | E3 SUMO-p | 20.797 | 19.66775 | 21.43199 | 19.14767 | 20.93915 | 19.54311 | 1.603206 | 0.861802 | 2.34461 | 20.25445 | 5.146776 | -0.8681 | 0.001464 | 0.012567 | 1 | 1 |
| 142 | sp\|Q8BG05 | ROA3_MOU | 40 kDa | 0.999911 | 12 | 0.75 | Q8BG05 | Hnrnpa3 | Heterogene | 25.09629 | 23.31561 | 24.26269 | 23.01576 | 24.88287 | 23.11078 | 1.599899 | 0.839059 | 2.360738 | 23.94733 | 5.004955 | -1.03247 | 0.001709 | 0.013629 | 1 | 1 |
| 277 | sp\|P62281 | RS11_MOU | 18 kDa | 0.999465 | 8 | 1 | P62281 | Rps11 | 40S ribosor | 24.70272 | 23.4546 | 24.58657 | 22.66491 | 24.48314 | 22.88109 | 1.590609 | 0.853852 | 2.327365 | 23.79551 | 5.135543 | -0.87756 | 0.001477 | 0.012599 | 1 | 1 |
| 104 | sp\|Q6A068 | CDC5L_MO | 92 kDa | 0.9996 | 15 | 0.27 | Q6A068 | Cdc5l | Cell division | 22.38783 | 20.79496 | 22.74659 | 20.95253 | 22.79606 | 21.42558 | 1.585804 | 0.865296 | 2.306312 | 21.85059 | 5.238554 | -0.76332 | 0.001327 | 0.012077 | 1 | 1 |
| 452.1 | sp\|P62270 | RS18_MOU | 18 kDa | 0.999008 | 6 | 1 | P62270 | Rps18 | 40S ribosor | 24.85654 | 23.21343 | 24.95486 | 23.24909 | 24.33298 | 22.92725 | 1.584867 | 0.875634 | 2.294099 | 23.92236 | 5.318691 | -0.67285 | 0.001219 | 0.011522 | 1 | 1 |
| 452.2 | tr\|A0A1Y7V | A0A1Y7VKY | 18 kDa | 0.999008 | 6 | 1 | A0A1Y7VKY | Gm11361 | MCG11667 | 24.85654 | 23.21343 | 24.95486 | 23.24909 | 24.33298 | 22.92725 | 1.584867 | 0.875634 | 2.294099 | 23.92236 | 5.318691 | -0.67285 | 0.001219 | 0.011522 | 1 | 1 |
| 452.3 | tr\|F6YVP7\| | F6YVP7_MO | 18 kDa | 0.999008 | 6 | 1 | F6YVP7 | Gm10260 | Predicted g | 24.85654 | 23.21343 | 24.95486 | 23.24909 | 24.33298 | 22.92725 | 1.584867 | 0.875634 | 2.294099 | 23.92236 | 5.318691 | -0.67285 | 0.001219 | 0.011522 | 1 | 1 |
| 170 | sp\|P48962 | ADT1_MOU | 33 kDa | 0.999351 | 11 | 0.55 | P48962 | Slc25a4 | ADP/ATP tra | 24.79143 | 23.2315 | 24.44439 | 22.75918 | 24.49906 | 23.01531 | 1.576296 | 0.909271 | 2.243321 | 23.79015 | 5.624665 | -0.33597 | 0.000888 | 0.009789 | 1 | 1 |
| 1040 | sp\|P62267 | RS23_MOU | 16 kDa | 0.99942 | 2 | 1 | P62267 | Rps23 | 40S ribosor | 24.25166 | 22.62279 | 24.47838 | 22.71259 | 24.08159 | 22.75469 | 1.573851 | 0.969897 | 2.200804 | 23.48362 | 5.974877 | 0.033478 | 0.000627 | 0.007788 | 1 | 1 |
| 111 | sp\|Q62095 | DDX3Y_MO | 73 kDa | 0.999675 | 15 | 0.13 | Q62095 | Ddx3y | ATP-depen | 19.10992 | 17.52939 | 19.08762 | 16.97791 | 18.737 | 17.74815 | 1.559697 | 0.805989 | 2.313406 | 18.19833 | 4.925359 | -1.12604 | 0.001865 | 0.014455 | 1 | 1 |
| 770 | sp\|Q9WUA2 | SYFB_MOU | 66 kDa | 0.996919 | 4 | 1 | Q9WUA2 | Farsb | Phenylalan | 21.28569 | 18.8359 | 22.92422 | 20.56716 | 21.21848 | 20.46881 | 1.518838 | 0.263461 | 2.774214 | 21.05004 | 2.887274 | -3.84556 | 0.024571 | 0.070145 | 1 | 0 |
| 23 | sp\|Q61768 | KINH_MOU | 110 kDa | 0.999746 | 27 | 0.81 | Q61768 | Kif5b | Kinesin-1 h | 24.23015 | 22.50157 | 24.08387 | 22.4242 | 24.27546 | 23.13422 | 1.509827 | 0.785926 | 2.233728 | 23.44158 | 4.964192 | -1.08027 | 0.001787 | 0.013927 | 1 | 1 |
| 8 | sp\|Q6P4T2 | U520_MOU | 245 kDa | 0.999706 | 46 | 1 | Q6P4T2 | Snrnp200 | U5 small n | 26.14711 | 24.35526 | 25.19469 | 24.83136 | 26.5794 | 24.24679 | 1.495931 | 0.486917 | 2.504946 | 25.22577 | 3.538059 | -2.92409 | 0.010209 | 0.040629 | 1 | 1 |
| 267 | sp\|P62806 | H4_MOUSE | 11 kDa | 0.999599 | 8 | 1 | P62806 | Hist4h4 | Histone H4 | 28.31263 | 27.3346 | 27.27886 | 25.73448 | 28.04698 | 26.0879 | 1.49383 | 0.282503 | 2.705158 | 27.13258 | 2.943 | -3.76506 | 0.022741 | 0.067396 | 1 | 0 |
| 399 | sp\|Q3B7Z2 | OSBP1_MO | 89 kDa | 0.999671 | 6 | 1 | Q3B7Z2 | Osbp | Oxysterol-b | 22.60444 | 20.02101 | 21.7037 | 21.84946 | 22.58158 | 20.61257 | 1.468892 | 0.193964 | 2.743819 | 21.56212 | 2.749507 | -4.04539 | 0.029801 | 0.080406 | 1 | 0 |
| 382 | sp\|Q99PU8 | DHX30_MO | 137 kDa | 0.99821 | 7 | 1 | Q99PU8 | Dhx30 | ATP-depen | 20.1292 | 18.1591 | 19.60263 | 17.52875 | 19.21206 | 18.85229 | 1.467917 | 0.430368 | 2.505465 | 18.914 | 3.376321 | -3.14872 | 0.01263 | 0.047309 | 1 | 1 |
| 559 | sp\|Q80YR5 | SAFB2_MO | 112 kDa | 0.99883 | 5 | 1 | Q80YR5 | Safb2 | Scaffold att | 23.31818 | 21.69918 | 22.26191 | 21.08378 | 23.0486 | 21.47517 | 1.456853 | 0.568295 | 2.345411 | 22.1478 | 3.908624 | -2.41626 | 0.006302 | 0.030671 | 1 | 1 |
| 1027 | sp\|P04247 | MYG_MOUS | 17 kDa | 0.999598 | 2 | 1 | P04247 | Mb | Myoglobin | 19.8928 | 18.89333 | 20.06902 | 18.93971 | 20.65781 | 18.52598 | 1.453536 | 0.631476 | 2.275597 | 19.47977 | 4.208661 | -2.01249 | 0.004288 | 0.02506 | 1 | 1 |
| 385 | sp\|P17427 | AP2A2_MO | 104 kDa | 0.997759 | 7 | 0.57 | P17427 | Ap2a2 | AP-2 comp | 21.497 | 19.59318 | 21.49084 | 19.78374 | 20.31035 | 19.56154 | 1.45324 | 0.512366 | 2.394115 | 20.37278 | 3.686009 | -2.72162 | 0.008433 | 0.03672 | 1 | 1 |
| 627.1 | sp\|P10853 | H2B1F_MO | 14 kDa | 0.999468 | 4 | 1 | P10853 | Hist1h2bn | Histone H2 | 27.60808 | 26.65783 | 26.88069 | 25.35859 | 27.35263 | 25.46878 | 1.452069 | 0.411749 | 2.492388 | 26.55443 | 3.330972 | -3.21228 | 0.013415 | 0.047997 | 1 | 1 |
| 627.2 | sp\|P10854 | H2B1M_MO | 14 kDa | 0.999468 | 4 | 1 | P10854 | Hist1h2bm | Histone H2 | 27.60808 | 26.65783 | 26.88069 | 25.35859 | 27.35263 | 25.46878 | 1.452069 | 0.411749 | 2.492388 | 26.55443 | 3.330972 | -3.21228 | 0.013415 | 0.047997 | 1 | 1 |
| 627.3 | sp\|Q6ZWY9 | H2B1C_MO | 14 kDa | 0.999468 | 4 | 1 | Q6ZWY9 | Hist1h2bg | Histone H2 | 27.60808 | 26.65783 | 26.88069 | 25.35859 | 27.35263 | 25.46878 | 1.452069 | 0.411749 | 2.492388 | 26.55443 | 3.330972 | -3.21228 | 0.013415 | 0.047997 | 1 | 1 |
| 627.4 | sp\|Q8CGP1 | H2B1K_MO | 14 kDa | 0.999468 | 4 | 1 | Q8CGP1 | Hist1h2bk | Histone H2 | 27.60808 | 26.65783 | 26.88069 | 25.35859 | 27.35263 | 25.46878 | 1.452069 | 0.411749 | 2.492388 | 26.55443 | 3.330972 | -3.21228 | 0.013415 | 0.047997 | 1 | 1 |
| 627.5 | sp\|Q8CGP2 | H2B1P_MO | 14 kDa | 0.999468 | 4 | 1 | Q8CGP2 | Hist1h2bp | Histone H2 | 27.60808 | 26.65783 | 26.88069 | 25.35859 | 27.35263 | 25.46878 | 1.452069 | 0.411749 | 2.492388 | 26.55443 | 3.330972 | -3.21228 | 0.013415 | 0.047997 | 1 | 1 |
| 627.6 | sp\|Q64475 | H2B1B_MO | 14 kDa | 0.999468 | 4 | 1 | Q64475 | Hist1h2bb | Histone H2 | 27.60808 | 26.65783 | 26.88069 | 25.35859 | 27.35263 | 25.46878 | 1.452069 | 0.411749 | 2.492388 | 26.55443 | 3.330972 | -3.21228 | 0.013415 | 0.047997 | 1 | 1 |
| 627.7 | sp\|Q64478 | H2B1H_MO | 14 kDa | 0.999468 | 4 | 1 | Q64478 | Hist1h2bh | Histone H2 | 27.60808 | 26.65783 | 26.88069 | 25.35859 | 27.35263 | 25.46878 | 1.452069 | 0.411749 | 2.492388 | 26.55443 | 3.330972 | -3.21228 | 0.013415 | 0.047997 | 1 | 1 |
| 627.8 | sp\|Q64525 | H2B2B_MO | 14 kDa | 0.999468 | 4 | 1 | Q64525 | Hist2h2bb | Histone H2 | 27.60808 | 26.65783 | 26.88069 | 25.35859 | 27.35263 | 25.46878 | 1.452069 | 0.411749 | 2.492388 | 26.55443 | 3.330972 | -3.21228 | 0.013415 | 0.047997 | 1 | 1 |
| 489 | sp\|P47963 | RL13_MOU | 24 kDa | 0.99974 | 5 | 1 | P47963 | Rpl13 | 60S ribosor | 25.0958 | 23.73636 | 25.33176 | 23.65812 | 24.96917 | 23.67106 | 1.443731 | 0.824413 | 2.06305 | 24.41036 | 5.548471 | -0.41861 | 0.00096 | 0.010246 | 1 | 1 |
| 441 | sp\|P62717 | RL18A_MO | 21 kDa | 0.999204 | 6 | 1 | P62717 | Rpl18a | 60S ribosor | 23.75002 | 22.17157 | 23.98249 | 22.46415 | 23.73768 | 22.51228 | 1.440731 | 0.805961 | 2.0755 | 23.10303 | 5.402163 | -0.57961 | 0.001117 | 0.011262 | 1 | 1 |
| 657 | sp\|Q9QUI0 | RHOA_MO | 22 kDa | 0.999264 | 4 | 0.25 | Q9QUI0 | Rhoa | Transformir | 19.77692 | 17.76794 | 20.3614 | 18.64351 | 19.62169 | 19.0265 | 1.436683 | 0.448794 | 2.432572 | 19.19966 | 3.46622 | -3.02345 | 0.011216 | 0.043453 | 1 | 1 |
| 112 | sp\|P06880 | SOMA_MO | 25 kDa | 0.999893 | 14 | 1 | P06880 | Gh1 | Somatotrop | 30.46798 | 29.19401 | 31.5654 | 30.12217 | 31.10104 | 29.50956 | 1.436224 | 0.468745 | 2.403703 | 30.32669 | 3.542676 | -2.91772 | 0.010148 | 0.040629 | 1 | 1 |
| 394 | sp\|Q9CZT8 | RAB3B_MO | 25 kDa | 0.999744 | 6 | 0.33 | Q9CZT8 | Rab3b | Ras-related | 22.3992 | 21.13097 | 22.84088 | 20.58854 | 22.00084 | 21.22692 | 1.4315 | 0.611413 | 2.251587 | 21.69789 | 4.154831 | -2.08221 | 0.004578 | 0.025802 | 1 | 1 |
| 28 | sp\|A2BH40 | ARI1A_MO | 242 kDa | 0.999848 | 25 | 0.92 | A2BH40 | Arid1a | AT-rich inter | 23.41653 | 22.00725 | 22.88232 | 21.53515 | 23.66679 | 22.13474 | 1.429503 | 0.633365 | 2.225641 | 22.60713 | 4.273634 | -1.92856 | 0.003962 | 0.02377 | 1 | 1 |
| 532 | sp\|Q6NVF9 | CPSF6_MO | 59 kDa | 0.999221 | 5 | 1 | Q6NVF9 | Cpsf6 | Cleavage ar | 23.60424 | 22.58169 | 23.01529 | 21.44957 | 23.17718 | 21.52069 | 1.414919 | 0.466095 | 2.363744 | 22.55811 | 3.558742 | -2.8956 | 0.009938 | 0.040469 | 1 | 1 |
| 916.11 | sp\|Q8CGP6 | H2A1F_MO | 14 kDa | 0.998436 | 3 | 0.33 | Q8CGP5 | Hist1h2af | Histone H2 | 21.70109 | 19.53103 | 20.35261 | 19.26683 | 20.57721 | 19.59098 | 1.414022 | 0.431684 | 2.396359 | 20.16996 | 3.435153 | -3.06662 | 0.011684 | 0.044756 | 1 | 1 |
| 1075 | sp\|Q61033 | LAP2A_MO | 75 kDa | 0.999163 | 2 | 1 | Q61033 | Tmpo | Lamina-ass | 17.217 | 16.49091 | 18.70633 | 16.7869 | 17.75882 | 16.16876 | 1.411857 | 0.366835 | 2.456478 | 17.18812 | 3.224155 | -3.36294 | 0.015481 | 0.051893 | 1 | 1 |
| 969 | sp\|Q08460 | KCMA1_MO | 134 kDa | 0.997601 | 3 | 1 | Q08460 | Kcnma1 | Calcium-ac | 17.678 | 15.88339 | 18.00248 | 16.26589 | 17.57318 | 16.89396 | 1.403468 | 0.57301 | 2.233927 | 17.04948 | 4.022597 | -2.25532 | 0.005387 | 0.027767 | 1 | 1 |
| 1152 | sp\|P01844 | LAC2_MOU | 11 kDa | 0.99843 | 2 | 0.5 | P01844 | Iglc2 | Ig lambda-2 | 21.79314 | 19.61536 | 20.98702 | 18.86239 | 20.24305 | 19.26072 | 1.401577 | 0.589401 | 2.213754 | 19.94695 | 4.107604 | -2.14373 | 0.00485 | 0.026309 | 1 | 1 |
| 776 | sp\|Q9D0M5 | DYL2_MOU | 10 kDa | 0.995494 | 4 | 0.5 | Q9D0M5 | Dynll2 | Dynein ligh | 22.52327 | 20.83074 | 22.79541 | 21.85832 | 22.67702 | 21.11447 | 1.39739 | 0.573796 | 2.220984 | 21.96654 | 4.038558 | -2.23429 | 0.005282 | 0.027546 | 1 | 1 |
| 613 | sp\|P62918 | RL8_MOUS | 28 kDa | 0.99957 | 4 | 1 | P62918 | Rpl8 | 60S ribosor | 23.91971 | 21.96292 | 25.35297 | 23.62357 | 24.659 | 23.16656 | 1.392877 | 0.367367 | 2.418387 | 23.94745 | 3.241333 | -3.33863 | 0.015127 | 0.051539 | 1 | 1 |
| 894 | sp\|P21107 | TPM3_MO | 33 kDa | 0.998754 | 3 | 0.33 | P21107 | Tpm3 | Tropomyos | 23.42961 | 22.32265 | 24.02872 | 22.19628 | 23.23099 | 22.03522 | 1.378388 | 0.630675 | 2.126101 | 22.87391 | 4.387702 | -1.78339 | 0.003457 | 0.021602 | 1 | 1 |
| 811 | sp\|Q62186 | SSRD_MO | 19 kDa | 0.999444 | 3 | 1 | Q62186 | Ssr4 | Transloco | 23.63197 | 22.07744 | 23.67203 | 21.73563 | 23.51822 | 22.88704 | 1.374038 | 0.512621 | 2.235455 | 22.92039 | 3.802587 | -2.55813 | 0.007203 | 0.033449 | 1 | 1 |
| 141 | sp\|Q9DBG3 | AP2B1_MO | 105 kDa | 0.997806 | 13 | 0.46 | Q9DBG3 | Ap2b1 | AP-2 comp | 23.15598 | 21.37204 | 23.25725 | 21.8094 | 21.75426 | 20.86725 | 1.372933 | 0.193299 | 2.552567 | 22.03603 | 2.77749 | -4.00472 | 0.028611 | 0.078964 | 1 | 0 |
| 662 | sp\|P63024 | VAMP3_MO | 11 kDa | 0.999219 | 4 | 0.5 | P63024 | Vamp3 | Vesicle-ass | 22.11123 | 20.79044 | 20.86191 | 19.33916 | 21.63336 | 20.36416 | 1.370912 | 0.178921 | 2.562904 | 20.85004 | 2.744651 | -4.05245 | 0.030005 | 0.080729 | 1 | 0 |
| 944 | sp\|Q8K019 | BCLF1_MO | 106 kDa | 0.998011 | 3 | 1 | Q8K019 | Bclaf1 | Bcl-2-assoc | 21.51691 | 20.60907 | 21.94028 | 20.36035 | 22.17413 | 20.55908 | 1.367607 | 0.669924 | 2.065289 | 21.1933 | 4.665565 | -1.43814 | 0.0025 | 0.017844 | 1 | 1 |
| 508 | sp\|Q8K0E8 | FIBB_MOU | 55 kDa | 0.999491 | 5 | 1 | Q8K0E8 | Fgb | Fibrinogen | 22.61871 | 20.39334 | 21.63383 | 21.20591 | 22.10401 | 20.67798 | 1.361281 | 0.463388 | 2.259174 | 21.43871 | 3.614243 | -2.81408 | 0.00917 | 0.038262 | 1 | 1 |
| 524 | sp\|Q9CZM2 | RL15_MOU | 24 kDa | 0.999297 | 5 | 1 | Q9CZM2 | Rpl15 | 60S ribosor | 24.96063 | 23.24249 | 24.88872 | 23.63036 | 24.9648 | 23.87104 | 1.356754 | 0.680046 | 2.033462 | 24.25967 | 4.772004 | -1.30903 | 0.002215 | 0.016125 | 1 | 1 |
| 136 | sp\|P14148 | RL7_MOUS | 31 kDa | 0.99931 | 13 | 1 | P14148 | Rpl7 | 60S ribosor | 25.84312 | 24.28867 | 26.35916 | 24.82055 | 25.8448 | 24.87377 | 1.354696 | 0.607028 | 2.102364 | 25.33834 | 4.312547 | -1.87881 | 0.003781 | 0.022951 | 1 | 1 |
| 574 | sp\|P35276 | RAB3D_MO | 24 kDa | 0.998133 | 5 | 0.2 | P35276 | Rab3d | Ras-related | 23.05199 | 21.57264 | 22.76164 | 21.55718 | 22.76453 | 21.38654 | 1.353929 | 0.73204 | 1.975819 | 22.18242 | 5.181839 | -0.82792 | 0.00141 | 0.012336 | 1 | 1 |
| 400 | sp\|Q6ZWV3 | RL10_MOU | 25 kDa | 0.99967 | 6 | 1 | Q6ZWV3 | Rpl10 | 60S ribosor | 23.70277 | 22.76863 | 24.73022 | 22.67266 | 23.89147 | 22.84036 | 1.347603 | 0.519987 | 2.175219 | 23.43375 | 3.875745 | -2.45063 | 0.006475 | 0.031134 | 1 | 1 |
| 109 | sp\|P62702 | RS4X_MOU | 30 kDa | 0.999356 | 15 | 1 | P62702 | Rps4x | 40S ribosor | 24.89577 | 23.38949 | 24.89357 | 23.57903 | 24.68421 | 23.46402 | 1.346508 | 0.738328 | 1.95567 | 24.15102 | 5.267271 | -0.7308 | 0.001287 | 0.011873 | 1 | 1 |
| 88 | sp\|Q8CHC4 | SYNJ1_MO | 173 kDa | 0.999829 | 16 | 1 | Q8CHC4 | Synj1 | Synaptojan | 23.53908 | 21.82992 | 22.44211 | 21.43725 | 23.00188 | 21.69697 | 1.339644 | 0.488383 | 2.190904 | 22.32454 | 3.750784 | -2.62676 | 0.007678 | 0.034939 | 1 | 1 |
| 849 | sp\|Q6ZPJ3 | UBE2O_MO | 141 kDa | 0.999172 | 3 | 1 | Q6ZPJ3 | Ube2o | (E3-indepe | 21.22238 | 20.05973 | 20.16797 | 19.47169 | 21.10626 | 18.95748 | 1.335902 | 0.304726 | 2.367077 | 20.16425 | 3.091666 | -3.55152 | 0.018533 | 0.058038 | 1 | 0 |
| 679.1 | sp\|P02301 | H3C_MOUS | 15 kDa | 0.999062 | 4 | 1 | P02301 | H3f3c | Histone H3 | 26.84963 | 25.79016 | 26.23383 | 25.38899 | 27.05354 | 24.9619 | 1.331984 | 0.472108 | 2.191861 | 26.04634 | 3.69281 | -2.70671 | 0.008286 | 0.036481 | 1 | 1 |
| 679.2 | sp\|P68433 | H31_MOUS | 15 kDa | 0.999062 | 4 | 1 | P68433 | Hist1h3i | Histone H3 | 26.84963 | 25.79016 | 26.23383 | 25.38899 | 27.05354 | 24.9619 | 1.331984 | 0.472108 | 2.191861 | 26.04634 | 3.69281 | -2.70671 | 0.008286 | 0.036481 | 1 | 1 |
| 679.3 | sp\|P84228 | H32_MOUS | 15 kDa | 0.999062 | 4 | 1 | P84228 | Hist2h3c2 | Histone H3 | 26.84963 | 25.79016 | 26.23383 | 25.38899 | 27.05354 | 24.9619 | 1.331984 | 0.472108 | 2.191861 | 26.04634 | 3.69281 | -2.70671 | 0.008286 | 0.036481 | 1 | 1 |
| 679.4 | sp\|P84244 | H33_MOUS | 15 kDa | 0.999062 | 4 | 1 | P84244 | H3f3b | Histone H3 | 26.84963 | 25.79016 | 26.23383 | 25.38899 | 27.05354 | 24.9619 | 1.331984 | 0.472108 | 2.191861 | 26.04634 | 3.69281 | -2.70671 | 0.008286 | 0.036481 | 1 | 1 |
| 596 | sp\|Q99K48 | NONO_MO | 55 kDa | 0.999738 | 4 | 0.75 | Q99K48 | Nono | Non-POU d | 20.89474 | 19.68457 | 21.03482 | 19.68744 | 20.48027 | 19.06889 | 1.322976 | 0.562791 | 2.083162 | 20.14179 | 4.142221 | -2.09826 | 0.004647 | 0.025929 | 1 | 1 |
| 626 | sp\|Q9D1M0 | SEC13_MO | 36 kDa | 0.999481 | 4 | 1 | Q9D1M0 | Sec13 | Protein SEC | 22.36954 | 20.35718 | 21.84705 | 20.1058 | 21.82239 | 21.61808 | 1.319306 | 0.227413 | 2.4112 | 21.35334 | 2.883474 | -3.85105 | 0.024702 | 0.070369 | 1 | 0 |
| 536 | sp\|Q922Q8 | LRC59_MO | 35 kDa | 0.999211 | 5 | 1 | Q922Q8 | Lrrc59 | Leucine-ric | 22.67913 | 21.96097 | 22.66218 | 21.11112 | 22.70258 | 21.02209 | 1.316565 | 0.514377 | 2.118754 | 22.02301 | 3.90631 | -2.40942 | 0.006227 | 0.030594 | 1 | 1 |
| 543 | sp\|P21956 | MFGM_MO | 51 kDa | 0.999127 | 5 | 1 | P21956 | Mfge8 | Lactadherin | 20.43569 | 19.30064 | 21.21015 | 19.72779 | 20.78354 | 19.4737 | 1.309081 | 0.558832 | 2.059331 | 20.15525 | 4.152997 | -2.08424 | 0.004586 | 0.025802 | 1 | 1 |
| 169 | sp\|Q99JR8 | SMRD2_MO | 59 kDa | 0.999372 | 11 | 0.91 | Q99JR8 | Smarcd2 | SWI/SNF-re | 23.43968 | 20.97937 | 22.3396 | 22.52246 | 23.50122 | 21.8694 | 1.303092 | 0.0767 | 2.529483 | 22.44195 | 2.535692 | -4.35696 | 0.040364 | 0.099878 | 1 | 0 |
| 87 | sp\|P61979 | HNRPK_MO | 51 kDa | 0.999832 | 16 | 1 | P61979 | Hnrnpk | Heterogene | 25.68902 | 24.44393 | 25.63567 | 24.37872 | 25.64861 | 24.2448 | 1.301951 | 0.705108 | 1.898794 | 25.00679 | 5.192008 | -0.8163 | 0.001395 | 0.012336 | 1 | 1 |
| 607 | sp\|P62751 | RL23A_MO | 18 kDa | 0.999674 | 4 | 1 | P62751 | Rpl23a | 60S ribosor | 24.08806 | 22.72058 | 24.21192 | 22.90146 | 24.01876 | 22.82157 | 1.291708 | 0.688635 | 1.89478 | 23.46039 | 5.097951 | -0.92435 | 0.001544 | 0.012892 | 1 | 1 |
| 961 | sp\|Q64378 | FKBP5_MO | 51 kDa | 0.997747 | 3 | 1 | Q64378 | Fkbp5 | Peptidyl-pr | 21.81719 | 19.52478 | 21.06636 | 20.02997 | 21.8661 | 21.34804 | 1.282288 | 0.028515 | 2.536062 | 20.94207 | 2.440716 | -4.49547 | 0.046246 | 0.109825 | 1 | 0 |
| 174 | sp\|P51881 | ADT2_MOU | 33 kDa | 0.999284 | 11 | 0.55 | P51881 | Slc25a5 | ADP/ATP tra | 24.52507 | 23.05248 | 23.82315 | 22.66325 | 24.18888 | 22.97536 | 1.282003 | 0.555055 | 2.008951 | 23.53803 | 4.197458 | -2.02661 | 0.004344 | 0.025066 | 1 | 1 |
| 951 | sp\|Q8K1M6 | DNM1L_MO | 83 kDa | 0.997901 | 3 | 1 | Q8K1M6 | Dnm1l | Dynamin-1 | 20.22039 | 18.34894 | 20.94661 | 19.36239 | 20.20156 | 19.83991 | 1.27244 | 0.176408 | 2.368472 | 19.81997 | 2.770543 | -4.01481 | 0.028932 | 0.079416 | 1 | 0 |
| 962 | sp\|P20357 | MTAP2_MO | 199 kDa | 0.999708 | 3 | 1 | P20357 | Map2 | Microtubule | 19.9232 | 19.08077 | 19.92721 | 17.82748 | 19.56382 | 18.68972 | 1.272088 | 0.347611 | 2.196565 | 19.1687 | 3.283124 | -3.27884 | 0.014283 | 0.049787 | 1 | 1 |
| 717 | sp\|Q9CYH2 | PXL2A_MO | 24 kDa | 0.998714 | 4 | 1 | Q9CYH2 | Prxl2a | Peroxiredox | 20.74462 | 19.6203 | 21.02182 | 19.35665 | 20.32793 | 19.31563 | 1.267262 | 0.552984 | 1.981541 | 20.06449 | 4.222793 | -1.9939 | 0.004213 | 0.024944 | 1 | 1 |
| 931 | sp\|Q3TXS7 | PSMD1_MO | 106 kDa | 0.998211 | 3 | 1 | Q3TXS7 | Psmd1 | 26S protea | 21.5493 | 20.55774 | 22.5378 | 20.84932 | 21.24385 | 20.13167 | 1.264069 | 0.261182 | 2.266957 | 21.11495 | 3.00794 | -3.67155 | 0.020789 | 0.063065 | 1 | 0 |
| 483 | sp\|O08547 | SC22B_MO | 25 kDa | 0.999894 | 5 | 1 | O08547 | Sec22b | Vesicle-traf | 23.79215 | 22.57444 | 24.05767 | 22.73716 | 23.46862 | 22.24778 | 1.25302 | 0.538889 | 1.967152 | 23.1463 | 4.176194 | -2.05413 | 0.004458 | 0.025501 | 1 | 1 |
| 880 | sp\|Q9CW03 | SMC3_MOU | 142 kDa | 0.998919 | 3 | 1 | Q9CW03 | Smc3 | Structural n | 21.30276 | 20.49528 | 22.70691 | 21.01078 | 22.34168 | 21.09511 | 1.25045 | 0.220391 | 2.280509 | 21.49228 | 2.897043 | -3.83143 | 0.02424 | 0.06949 | 1 | 0 |
| 1287 | sp\|P26638 | SYSC_MOU | 58 kDa | 0.995287 | 2 | 1 | P26638 | Sars | Serine--tRN | 17.36066 | 16.74451 | 17.39737 | 16.11356 | 16.99057 | 15.13956 | 1.250322 | 0.183512 | 2.317132 | 16.62437 | 2.796955 | -3.97645 | 0.027878 | 0.077783 | 1 | 0 |
| 167 | sp\|Q91VR5 | DDX1_MOU | 82 kDa | 0.999436 | 11 | 1 | Q91VR5 | Ddx1 | ATP-depen | 23.30122 | 21.80544 | 22.70698 | 21.38053 | 22.84285 | 21.92862 | 1.245489 | 0.50796 | 1.983019 | 22.32761 | 4.019401 | -2.25922 | 0.005406 | 0.027767 | 1 | 1 |
| 271 | sp\|Q9CXY6 | ILF2_MOUS | 43 kDa | 0.999564 | 8 | 1 | Q9CXY6 | Ilf2 | Interleukin | 22.558 | 21.56879 | 22.00128 | 21.26223 | 22.60948 | 20.60511 | 1.24421 | 0.382711 | 2.105708 | 21.76748 | 3.442967 | -3.05101 | 0.011475 | 0.04408 | 1 | 1 |
| 702 | sp\|Q9WV55 | VAPA_MOU | 28 kDa | 0.998865 | 4 | 0.75 | Q9WV55 | Vapa | Vesicle-ass | 21.03147 | 20.30019 | 21.34536 | 19.71504 | 21.1408 | 19.77819 | 1.241406 | 0.543529 | 1.939283 | 20.55184 | 4.233854 | -1.97965 | 0.004157 | 0.024829 | 1 | 1 |
| 774 | tr\|A0A075B | A0A075B5I | 13 kDa | 0.995633 | 4 | 0.25 | A0A075B5I | Ighv7-3 | Immunoglo | 22.6625 | 21.39578 | 22.08317 | 21.19738 | 22.67557 | 21.21905 | 1.203013 | 0.50692 | 1.899106 | 21.87224 | 4.113428 | -2.15579 | 0.004813 | 0.026213 | 1 | 1 |
| 9 | sp\|Q9PV0 | PRP8_MOU | 274 kDa | 0.999619 | 45 | 1 | Q9PV0 | Prpf8 | Pre-mRNA-p | 26.51989 | 24.63566 | 25.40605 | 25.26912 | 26.96499 | 25.38132 | 1.201608 | 0.081871 | 2.321345 | 25.69617 | 2.560927 | -4.32015 | 0.038935 | 0.097052 | 1 | 0 |
| 608 | sp\|Q99M28 | RNPS1_MO | 34 kDa | 0.999648 | 4 | 1 | Q99M28 | Rnps1 | RNA-bindin | 23.17282 | 21.85916 | 22.4934 | 21.54962 | 22.90669 | 21.61598 | 1.182714 | 0.472782 | 1.892646 | 22.26628 | 3.965189 | -2.33098 | 0.005784 | 0.029047 | 1 | 1 |
| 379 | sp\|Q9QZQ8 | H2AY_MOU | 40 kDa | 0.999802 | 7 | 1 | Q9QZQ8 | H2afy | Core histon | 23.14974 | 22.48891 | 22.8055 | 21.74509 | 23.35294 | 21.52854 | 1.18188 | 0.337856 | 2.025905 | 22.51178 | 3.335293 | -3.19822 | 0.013161 | 0.047997 | 1 | 1 |
| 323 | sp\|P35980 | RL18_MOU | 22 kDa | 0.999574 | 7 | 1 | P35980 | Rpl18 | 60S ribosor | 25.86043 | 25.50647 | 26.16436 | 24.52851 | 25.80528 | 24.252 | 1.181033 | 0.245946 | 2.11612 | 25.35284 | 3.01402 | -3.66267 | 0.020613 | 0.062671 | 1 | 0 |
| 348 | sp\|P23249 | MOV10_MO | 114 kDa | 0.999342 | 7 | 1 | P23249 | Mov10 | Putative hel | 21.27676 | 19.93984 | 21.55647 | 20.3593 | 20.98829 | 19.98992 | 1.177482 | 0.475892 | 1.879071 | 20.6851 | 3.994585 | -2.29202 | 0.005575 | 0.028171 | 1 | 1 |
| 781 | tr\|Q3U4B1 | Q3U4B1_M | 37 kDa | 0.999788 | 3 | 0.67 | Q3U4B1 | Ssbp4 | Single stran | 20.36239 | 19.84935 | 21.6409 | 19.64033 | 20.25046 | 19.25902 | 1.168347 | 0.111184 | 2.225511 | 20.16707 | 2.637428 | -4.20858 | 0.034919 | 0.089678 | 1 | 0 |
| 468 | sp\|O70228 | ATP9A_MO | 119 kDa | 0.998178 | 6 | 0.67 | O70228 | Atp9a | Probable p | 20.46668 | 19.03661 | 20.53157 | 19.15316 | 20.25761 | 19.58721 | 1.165597 | 0.481043 | 1.838212 | 19.83881 | 4.067386 | -2.19606 | 0.005094 | 0.027196 | 1 | 1 |
| 730 | sp\|Q9CYN9 | RENR_MOU | 39 kDa | 0.998453 | 4 | 1 | Q9CYN9 | Atp6ap2 | Renin recep | 21.33995 | 20.01686 | 20.3396 | 19.81024 | 21.11157 | 19.49734 | 1.15556 | 0.302683 | 2.008438 | 20.35259 | 3.229248 | -3.35088 | 0.015248 | 0.05179 | 1 | 0 |
| 80 | sp\|Q3TEA8 | HP1B3_MO | 61 kDa | 0.999402 | 17 | 1 | Q3TEA8 | Hp1bp3 | Heterochro | 24.89401 | 23.54595 | 25.31159 | 24.88947 | 25.29301 | 23.61559 | 1.149199 | 0.123926 | 2.174472 | 24.5916 | 2.674892 | -4.15399 | 0.033112 | 0.086848 | 1 | 0 |
| 19 | sp\|Q8BTI8 | SRRM2_MO | 295 kDa | 0.999729 | 29 | 1 | Q8BTI8 | Srrm2 | Serine/argin | 25.50006 | 24.34192 | 25.24684 | 24.57461 | 26.15066 | 24.54117 | 1.146622 | 0.369694 | 1.923551 | 25.05921 | 3.512693 | -2.94644 | 0.010341 | 0.040881 | 1 | 1 |
| 236 | sp\|Q00PI9 | HNRL2_MO | 85 kDa | 0.999487 | 9 | 1 | Q00PI9 | Hnrnpul2 | Heterogene | 23.46053 | 21.80072 | 23.27767 | 21.56276 | 22.95646 | 22.90901 | 1.140717 | 0.141444 | 2.13999 | 22.66119 | 2.724233 | -4.08215 | 0.030882 | 0.082417 | 1 | 0 |
| 406 | sp\|P14869 | RLA0_MOU | 34 kDa | 0.999608 | 6 | 1 | P14869 | Rplp0 | 60S acidic r | 22.43835 | 21.56258 | 23.14436 | 21.87303 | 22.72625 | 21.48823 | 1.128374 | 0.39992 | 1.856827 | 22.20547 | 3.686818 | -2.70623 | 0.008239 | 0.036481 | 1 | 1 |
| 681 | sp\|Q9D0W5 | PPIL1_MOU | 18 kDa | 0.999051 | 4 | 1 | Q9D0W5 | Ppil1 | Peptidyl-pr | 21.37567 | 20.23097 | 20.71761 | 20.84909 | 21.91929 | 19.55584 | 1.125558 | 0.019056 | 2.232059 | 20.77474 | 2.427541 | -4.51467 | 0.04713 | 0.110037 | 1 | 0 |
| 907 | sp\|Q68FF6 | GIT1_MOU | 85 kDa | 0.998593 | 3 | 1 | Q68FF6 | Git1 | ARF GTPase | 20.14337 | 19.68871 | 20.58869 | 18.81858 | 21.31187 | 20.16075 | 1.125295 | 0.003098 | 2.247492 | 20.11866 | 2.39303 | -4.56494 | 0.049528 | 0.114084 | 1 | 0 |
| 653 | sp\|P42225 | STAT1_MO | 87 kDa | 0.999283 | 4 | 1 | P42225 | Stat1 | Signal trans | 20.35865 | 19.5651 | 20.85073 | 19.40968 | 20.54185 | 19.40103 | 1.12514 | 0.475323 | 1.774957 | 20.02117 | 4.121132 | -2.12573 | 0.004768 | 0.026134 | 1 | 1 |
| 110 | sp\|Q7TQG1 | PKHA6_MO | 131 kDa | 0.999124 | 15 | 1 | Q7TQG1 | Plekha6 | Pleckstrin h | 23.65664 | 21.53064 | 22.01761 | 22.06882 | 23.09271 | 21.79914 | 1.122787 | 0.023842 | 2.221731 | 22.36093 | 2.438216 | -4.49911 | 0.046413 | 0.109911 | 1 | 0 |
| 41 | sp\|Q9R0I7 | YLPM1_MO | 155 kDa | 0.999814 | 22 | 1 | Q9R0I7 | Ylpm1 | YLP motif-c | 26.61066 | 24.99479 | 25.31427 | 25.07381 | 26.83904 | 25.33052 | 1.121616 | 0.053378 | 2.189854 | 25.69385 | 2.505688 | -4.40073 | 0.042133 | 0.103689 | 1 | 0 |
| 343 | sp\|P68254 | 1433T_MO | 28 kDa | 0.99937 | 7 | 0.71 | P68254 | Ywhaq | 14-3-3 prot | 22.78802 | 21.55871 | 22.96714 | 21.86973 | 22.28267 | 21.25707 | 1.11744 | 0.350156 | 1.884724 | 22.12056 | 3.46632 | -3.01109 | 0.010994 | 0.043027 | 1 | 1 |
| 656 | sp\|O70404 | VAMP8_MO | 11 kDa | 0.999267 | 4 | 1 | O70404 | Vamp8 | Vesicle-ass | 25.10347 | 23.49243 | 24.69223 | 23.33267 | 24.8207 | 24.45309 | 1.112737 | 0.215775 | 2.009699 | 24.31577 | 2.957416 | -3.74119 | 0.022164 | 0.066639 | 1 | 0 |
| 832.5 | tr\|A0A140T | A0A140T8C | 13 kDa | 0.999312 | 3 | 0.33 | A0A140T8C | Igkv3-4 | Immunoglo | 23.40676 | 22.58069 | 23.36312 | 21.70752 | 23.16474 | 22.31016 | 1.112086 | 0.347275 | 1.876898 | 22.7555 | 3.460868 | -3.0187 | 0.011074 | 0.043027 | 1 | 1 |
| 422 | sp\|O08538 | ANGP1_MO | 58 kDa | 0.999433 | 6 | 1 | O08538 | Angpt1 | Angiopoieti | 21.66208 | 20.81513 | 22.21574 | 21.05985 | 21.15419 | 19.82284 | 1.111399 | 0.041083 | 2.181715 | 21.12164 | 2.478041 | -4.44105 | 0.043835 | 0.106715 | 1 | 0 |
| 624 | sp\|Q9QYH6 | MAGD1_MO | 86 kDa | 0.999491 | 4 | 1 | Q9QYH6 | Maged1 | Melanoma- | 20.44559 | 18.32565 | 20.25479 | 19.83698 | 20.48935 | 19.70369 | 1.107803 | 0.035008 | 2.180599 | 19.84268 | 2.464316 | -4.46106 | 0.044706 | 0.108018 | 1 | 0 |
| 196 | sp\|Q80WJ7 | LYRIC_MO | 64 kDa | 0.999538 | 10 | 1 | Q80WJ7 | Mtdh | Protein LYR | 24.06499 | 23.15208 | 24.28559 | 22.94461 | 24.34649 | 23.33512 | 1.088419 | 0.447462 | 1.729377 | 23.68815 | 4.041735 | -2.22978 | 0.005258 | 0.02753 | 1 | 1 |
| 957 | sp\|Q99P58 | RB27B_MO | 25 kDa | 0.997785 | 3 | 0.33 | Q99P58 | Rab27b | Ras-related | 21.76208 | 21.26309 | 22.24039 | 20.23237 | 21.23287 | 20.50384 | 1.078678 | 0.103477 | 2.05388 | 21.20578 | 2.639661 | -4.20533 | 0.034808 | 0.089564 | 1 | 0 |
| 1003 | sp\|Q8BX10 | PGAM5_MO | 32 kDa | 0.995432 | 3 | 1 | Q8BX10 | Pgam5 | Serine/thre | 20.61479 | 19.25946 | 21.03657 | 20.16825 | 20.80334 | 19.797 | 1.076663 | 0.285382 | 1.867945 | 20.2799 | 3.238543 | -3.3324 | 0.014922 | 0.051357 | 1 | 0 |
| 1142.2 | tr\|W4VSN7 | W4VSN7_M | 18 kDa | 0.998518 | 2 | 1 | W4VSN7 | Gm3550 | 60S ribosor | 21.53802 | 19.7664 | 22.0227 | 20.84075 | 21.50734 | 21.23563 | 1.075095 | 0.031224 | 2.118966 | 21.15181 | 2.457824 | -4.47053 | 0.045125 | 0.108491 | 1 | 0 |
| 1143 | sp\|P41105 | RL28_MOU | 16 kDa | 0.998509 | 2 | 1 | P41105 | Rpl28 | 60S ribosor | 21.53802 | 19.7664 | 22.0227 | 20.84075 | 21.50734 | 21.23563 | 1.075095 | 0.031224 | 2.118966 | 21.15181 | 2.457824 | -4.47053 | 0.045125 | 0.108491 | 1 | 0 |
| 740 | sp\|Q9Z108 | STAU1_MO | 54 kDa | 0.998308 | 4 | 1 | Q9Z108 | Stau1 | Double-stra | 20.90502 | 19.94529 | 21.08713 | 19.76023 | 20.53657 | 19.60812 | 1.071692 | 0.391062 | 1.752321 | 20.30706 | 3.747658 | -2.62327 | 0.007618 | 0.034782 | 1 | 1 |
| 671 | sp\|Q922H4 | GMPPA_MO | 46 kDa | 0.999115 | 4 | 1 | Q922H4 | Gmppa | Mannose-1 | 21.88209 | 20.61914 | 21.71688 | 21.04773 | 21.85652 | 20.61797 | 1.056882 | 0.407984 | 1.70578 | 21.29006 | 3.876601 | -2.44919 | 0.006465 | 0.031134 | 1 | 1 |
| 418 | sp\|P70372 | ELAV1_MO | 36 kDa | 0.999519 | 6 | 1 | P70372 | Elavl1 | ELAV-like p | 23.48833 | 22.39918 | 23.45239 | 22.14754 | 23.07889 | 22.30028 | 1.055871 | 0.408458 | 1.703284 | 22.81194 | 3.881775 | -2.44225 | 0.006423 | 0.031104 | 1 | 1 |
| 486 | sp\|P61358 | RL27_MOU | 16 kDa | 0.999784 | 5 | 1 | P61358 | Rpl27 | 60S ribosor | 24.21536 | 22.86925 | 24.38481 | 23.44313 | 24.006 | 23.12912 | 1.054886 | 0.36499 | 1.744781 | 23.67461 | 3.639342 | -2.77132 | 0.008762 | 0.037479 | 1 | 1 |
| 36 | sp\|E1U8D0 | SOGA1_MO | 159 kDa | 0.99968 | 23 | 0.91 | E1U8D0 | Soga1 | Protein SOG | 23.38885 | 22.81849 | 22.73312 | 22.3781 | 23.91569 | 21.72737 | 1.037901 | -0.00087 | 2.076668 | 22.82694 | 2.384452 | -4.57743 | 0.050143 | 0.114541 | 1 | 0 |
| 159 | sp\|P62908 | RS3_MOUS | 27 kDa | 0.999714 | 11 | 1 | P62908 | Rps3 | 40S ribosor | 26.7488 | 25.49396 | 26.55744 | 25.63845 | 26.84899 | 25.94945 | 1.024457 | 0.369736 | 1.679178 | 26.20618 | 3.724245 | -2.65514 | 0.007851 | 0.035516 | 1 | 1 |
| 463 | sp\|Q9DCD2 | SYF1_MOU | 100 kDa | 0.998711 | 6 | 1 | Q9DCD2 | Xab2 | Pre-mRNA-s | 20.7791 | 20.10436 | 20.67987 | 20.2501 | 21.74623 | 19.77823 | 1.024172 | 0.131058 | 1.917286 | 20.55631 | 2.733759 | -4.06589 | 0.030322 | 0.081418 | 1 | 0 |
| 85 | sp\|Q9CWF2 | TBB2B_MO | 50 kDa | 0.998684 | 17 | 0.06 | Q9CWF2 | Tubb2b | Tubulin bet | 20.83364 | 19.967 | 21.31004 | 20.26951 | 20.2755 | 19.1449 | 1.012591 | -0.00103 | 2.026213 | 20.3001 | 2.384014 | -4.57807 | 0.050175 | 0.114541 | 1 | 0 |
| 157 | sp\|Q9D0E1 | HNRPM_MO | 78 kDa | 0.999898 | 11 | 1 | Q9D0E1 | Hnrnpm | Heterogene | 23.81454 | 23.01013 | 23.27448 | 22.16757 | 23.58094 | 22.45551 | 1.011259 | 0.218618 | 1.80588 | 23.05053 | 3.035775 | -3.62314 | 0.019704 | 0.060451 | 1 | 0 |
| 583 | sp\|P84091 | AP2M1_MO | 50 kDa | 0.997211 | 5 | 1 | P84091 | Ap2m1 | AP-2 comp | 23.81037 | 22.32336 | 23.57002 | 22.19383 | 22.90525 | 22.73735 | 1.010369 | 0.186655 | 1.834082 | 22.92336 | 2.919616 | -3.79146 | 0.023167 | 0.067424 | 1 | 0 |
